# Determinants of chromosomal rearrangements in holocentric *Leptidea* butterflies

**DOI:** 10.64898/2026.02.26.708211

**Authors:** Filip Thörn, Jean-Loup Claret, Niclas Backström, Jesper Boman

## Abstract

Chromosomes can undergo large-scale rearrangements such as fissions and fusions. Occasional rearrangements can be common, especially in organisms with holocentric chromosomes such as butterflies. However, high rates of fissions and fusions have only been observed in a few taxonomic groups. One such group is the Palearctic *Leptidea* butterflies, where fissions and fusions have resulted in considerable inter- and intraspecific variation in chromosome numbers. The large number of rearrangements in *Leptidea*, provides a rare opportunity to study the mutational determinants of chromosomal rearrangements within a statistical framework. Using nine chromosome-level genome assemblies and 138 whole-genome re-sequenced individuals, we mapped evolutionary breakpoint regions and quantified the association between annotation features and rearrangements. Evolutionary breakpoint regions were significantly depleted in protein-coding genes and the majority resided in repetitive regions. However, rearrangements were only weakly associated with transposable elements. Instead, the strongest sequence predictors were large clusters of satellite DNA, ribosomal DNA and segmental duplications, with differing patterns among rearrangement types. Copy-number variation was observed in evolutionary breakpoint regions and lineages dominated by fissions or fusions were associated respectively with genome expansion and reduction. The results give novel insights into the mechanistic basis of interchromosomal rearrangements.

## Introduction

Interchromosomal rearrangements such as fissions and fusions represent major reorganisations of the genome with repercussions on health, adaptation and speciation [1–3]. Fissions and fusions of chromosomes lead to changes in chromosome number and have been studied for more than a century [4–6]. Still, our understanding of the mechanistic underpinnings of chromosomal rearrangements is limited. Some so-called ‘fragile sites’ in human cancers have been shown to overlap evolutionary breakpoints, giving some insights into the origin of fissions [7–9]. Recent investigations of two Robertsonian fusions in humans indicate that repetitive regions with inverted orientation on different chromosomes may mediate chromosome fusions via ectopic recombination [10,11]. While more evidence is required for certainty, it has been suggested that this model could be applicable for the many fusions in mice to men and even butterflies [12].

Chromosomes of Lepidoptera (butterflies and moths) are fundamentally different from mammalian chromosomes. While both have telomeres, only mammalian chromosomes have large regional monocentromeres, often characterised by arrays of satellite DNA [13]. Lepidoptera are holocentric, meaning that centromere activity is spread across a large portion of the chromosome [14]. This has widespread implications on differences in chromosome evolution between both systems. A mammalian chromosome without a centromere is expected to eventually be lost because it cannot be pulled to one of the centrosomes during cell division [15]. In holocentric organisms on the other hand, fissioned chromosomes can segregate correctly [16–18]. Regardless of this fundamental difference, chromosome fissions and fusions are observed in both holocentric and monocentric organisms, but it is perhaps not surprising that one of the extremes of fusions and fissions is found in a holocentric butterfly.

The cryptic wood white butterflies constitute a group of three species (*Leptidea sinapis, L. reali* and *L. juvernica*) with large inter- and intraspecific variation in chromosome numbers; 2n = 51-55 in *L. reali* and 76-91 in *L. juvernica* [19–21]. *Leptidea sinapis* shows the most extreme intraspecific chromosome number variation of any diploid animal with for example 2n = 57, 58 in Sweden to 2n = 106-108 in Catalonia, Spain. Karyotype differences between the species are believed to mainly have occurred through chromosome fissions and fusions [22]. While occasional fusions are rather common in Lepidoptera, large numbers of fusions, and especially fissions, are more rarely observed in both cytogenetic and comparative genomic analysis [23,24]. Despite generation of chromosome-level genome assemblies for more than a thousand Lepidoptera species, the drivers of chromosomal rearrangements are poorly understood [25,26]. In some cases, associations with specific transposable element (TE) superfamilies (including in *Leptidea*) and interstitial telomere sequences (in *Polyommatus*) have been observed [22,26,27]. However, detailed association studies are needed to further our understanding of the mechanistic underpinnings of chromosomal rearrangements. We know nothing of the associations between segmental duplications and chromosomal rearrangements in holocentric Lepidoptera, while such associations have frequently been seen in monocentric mammals [27–29]. In addition, ribosomal- and satellite DNA are associated with two Robertsonian fusions in humans [10]. Satellite DNA occurs but is generally rare in butterflies and moths [28]. *Leptidea* butterflies, are an exception in this aspect too, since they have experienced a recent expansion of satellite DNA [29]. It is therefore tempting to speculate upon a connection between satellite DNA proliferation and chromosomal rearrangements in general, but such an association has not been tested in *Leptidea* or any other lepidopteran taxon as far as we are aware. Here, we perform an in-depth investigation of evolutionary breakpoint regions (EBRs) in *Leptidea* butterflies and show that satellite DNA, ribosomal DNA repeat clusters and segmental duplications are significantly associated with chromosome fissions and fusions in this holocentric group.

## Results

### Mapping evolutionary breakpoint regions

We used nine chromosome-level assemblies of *Leptidea sinapis, L. reali* and *L. juvernica*, including a high-quality reference genome of an Asturian *L. sinapis*, to reconstruct chromosome evolution in this clade (Fig. 1A). We identified ∼66 fusions, ∼32 fissions and ∼35 putative translocations (Fig. S1). Note that if both fusions and translocations are mediated by ectopic recombination, they represent the same type of molecular event. *L. juvernica* was used as an outgroup and therefore we could not polarise events on the branch leading to *L. sinapis* + *L. reali*. Rates of interchromosomal rearrangements varied from 0.0023-0.13 per substitution for fusions, to 0.0015-0.04 for fissions and 0.003-0.09 for translocations. We defined evolutionary breakpoint regions (EBRs) as the region between pairs of BUSCO genes, in which a rearrangement had been called based on ancestral gene order reconstructions with AGORA. The EBRs that could be polarised varied in length from 1,361 to 3,146,840 bp (Fig. S1).

**Figure 1.**
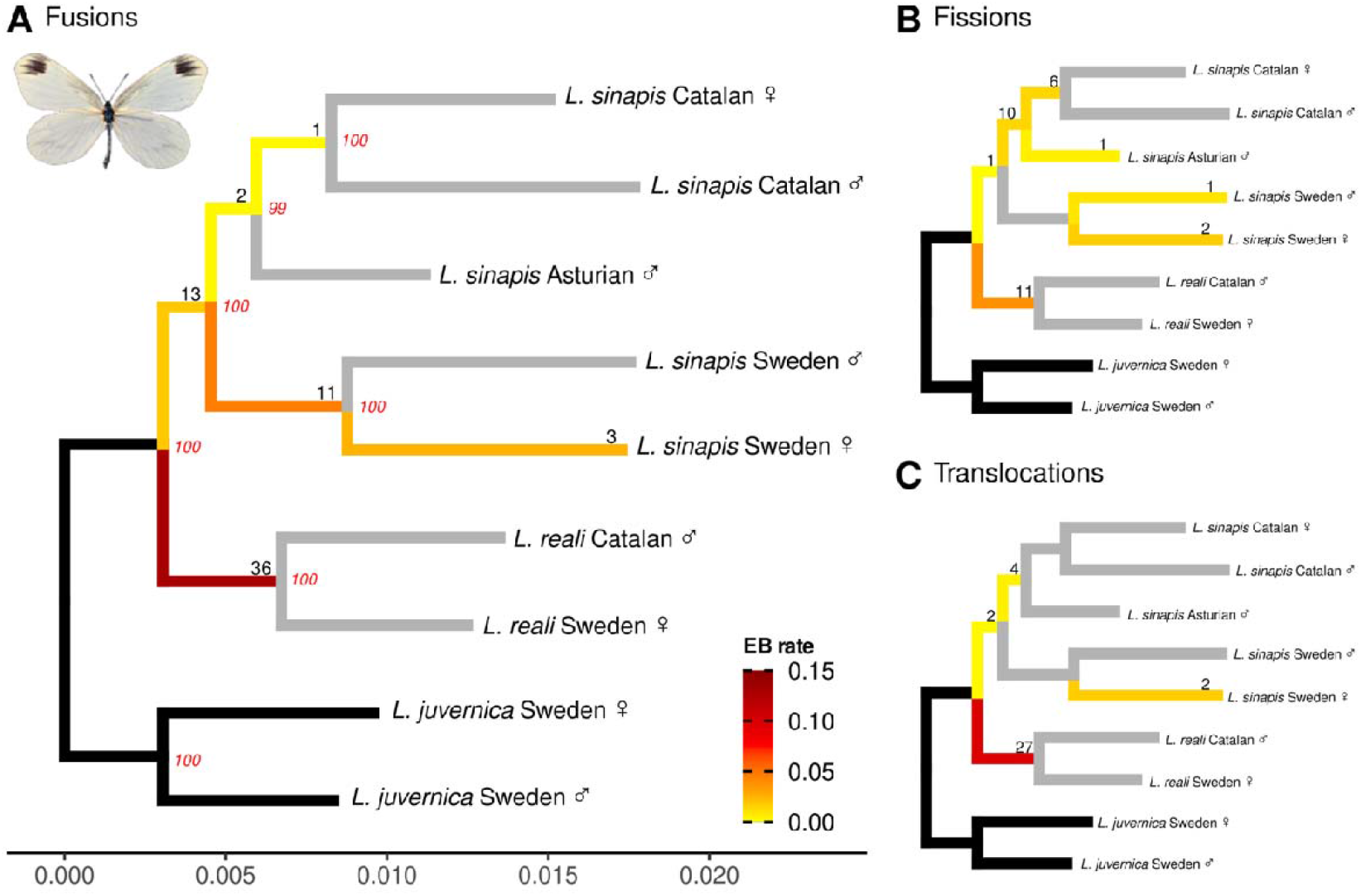
Interchromosomal rearrangements in the cryptic wood whites. Numbers and rates of fusions (A), fissions (B) and translocations (C). Average number of events per branch are given in black. All evolutionary breakpoint (EB) rates were calculated per branch as the mean number of events multiplied by the number of amino acid substitutions per site. Events on black branches could not be polarised. Grey branches represent an EB rate = 0. Picture in (A) depicts a mounted female *L. sinapis* photographed by Niclas Backström.

### The diverse repeat set of the *Leptidea* genome

Previous studies of determinants of rearrangements in butterflies have focused on TEs [22,26,30,31]. Given observations from other clades [32,33], we wanted to investigate the content of EBRs using a more comprehensive set of repeats. We gathered previously available data on the locations of satellite DNA and protein-coding genes in the *L. sinapis* reference genome [29,34]. In addition, we inferred the locations of ribosomal DNA (rDNA) repeat clusters and annotated TEs and other shorter repeat elements (such as telomeres). We observed two main clusters of rDNA units (Fig. S2). In exploratory analysis we noted that a tetramer of LepSat01-100 (monomer size 100 bp), the by-far most common satellite in the genome, was annotated as a long interspersed nuclear element (LINE) by RepeatModeler. We used CENSOR to search the RepBase database for support of this annotation, but we only found partial hits (∼50 bp) to much longer *Academ* DNA transposons, annotated in the bivalve, *Archivesica marissinica* [35] (Table S1). Such partial hits are common when manually curating repeats and do not suffice for identification [36,37]. We therefore removed all hits in the RepeatMasker output that overlapped with the satellite DNA annotation, to reduce the risk of erroneous annotation, which may have affected a previous observed association between LINEs and fusion EBRs in *Leptidea* [22]. In addition, we performed all-vs-all within-chromosome alignments in 2 kb windows and could observe both large clusters of satellite DNA (previously mapped [29]) and many previously undescribed clusters of large segmental duplications (Additional file 1, Fig. S3-S4). In total, large clusters of segmental duplications covered more than 5% of the *Leptidea sinapis* genome, highlighting a repeat class rarely studied in butterflies on a genome-wide scale [38]. On chromosome 12, for example, the reference genome has three copies of a segmental duplication with a monomer size ∼400 kb, but our read-depth analysis of 138 re-sequenced individuals revealed a copy number higher than three in most individuals (Fig. S5). Curiously, two EBRs cover this region: a fission ancestral to Catalan *L. sinapis* and an unpolarisable fission/fusion between *L. juvernica* and *L. sinapis* + *L. reali* (Fig. S1).

### Determinants of chromosomal rearrangements

We observed two EBRs overlapping the segmental duplication cluster on chromosome 12, but this observation alone does not verify that segmental duplications or other annotation features are *significantly* associated with chromosomal rearrangements. In most studies of chromosomal rearrangements, too few large-scale rearrangements have occurred to systematically test for associations. The large number of rearrangements within the cryptic wood whites provides a unique opportunity to statistically test for potential associations between annotation features and rearrangements. To do this, we calculated odds ratios of EBRs and annotation tracks, where a value close to 1 means that the two features overlap as frequently as expected by chance (Fig. 2). We used 1,000 permutations of the genomic positions of the annotation tracks and a two-tailed test to assess the significance of the overlap. All types of EBRs had around half the number of protein-coding genes as expected. In addition, almost all TE classes where underrepresented. Instead, fusion- and translocation EBRs where significantly associated with ribosomal- and satellite DNA (*p <* 0.001). All EBR types were also significantly enriched in segmental duplications. Fission EBRs were also enriched in satellite DNA (*p <* 0.001), but less so than the other rearrangement types. Instead, fission EBRs had a unique association with simple repeats (i.e. mini- and microsatellites; *p <* 0.001). Telomere repeat arrays (TTAGG)_n_ were not significantly associated with EBRs, but had a high odds-ratio for fissions (Fig. 2). *Leptidea* (like other Lepidoptera) have LINE elements in their telomeres, which could explain the significant but weak enrichment of LINEs in fission EBRs. We also explored the associations between small structural variants and EBRs but observed no significant associations (Table S2), with the limitation that short-read structural variants calling is uncertain in satellite regions (Table S3).

**Figure 2.**
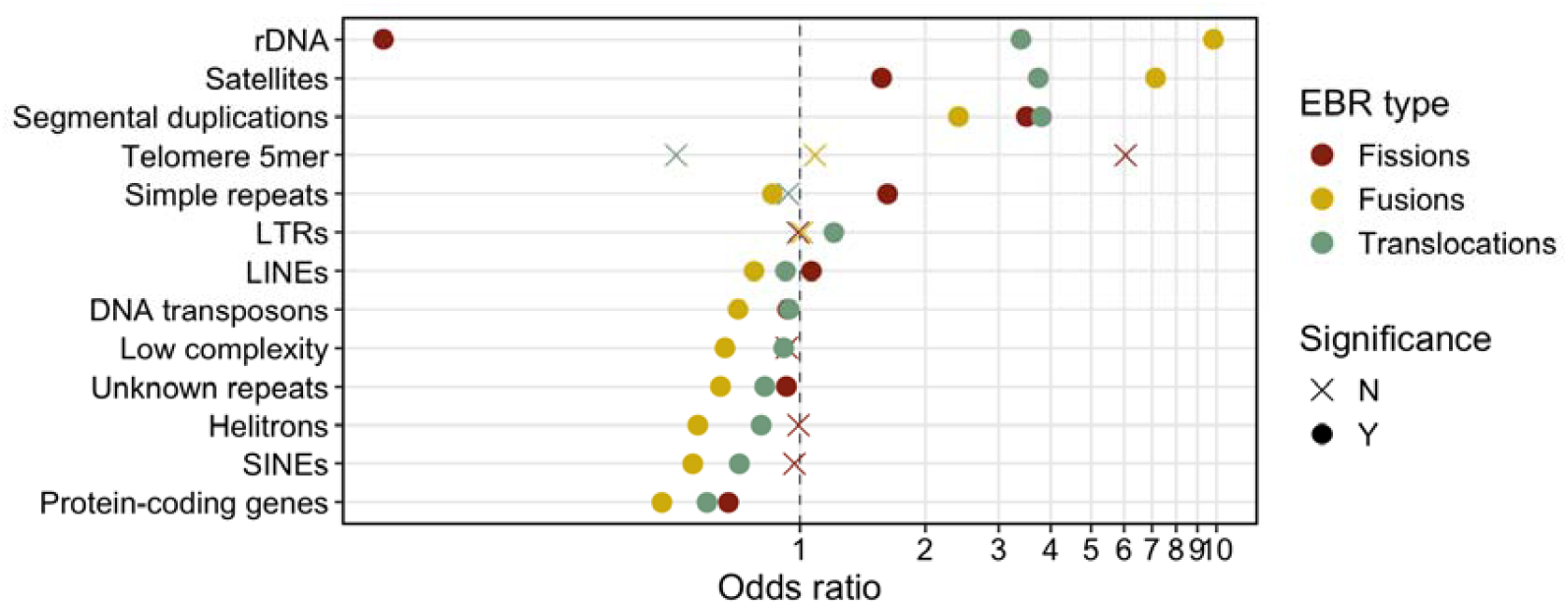
Observed odds ratios and permutation test results for the association between EBRs and annotated features. Shapes indicate significance after Benjamini-Hochberg correction per EBR type to a false discovery rate < 0.05.

### Sequence change associated with chromosomal rearrangements

In *Polyommatini* butterflies, cytogenetic analyses have shown that fissioned chromosomes are larger than homologous chromosomes in related species with the ancestral arrangement [39]. Conversely, fusions are believed to mainly occur through non-homologous- or ectopic recombination, which potentially leads to a loss of DNA [10]. Therefore, for these classes of EBRs, we have contrasting predictions for expansion and reduction of DNA. We tested this by studying copy number variation in 10 kb windows overlapping EBRs. To minimise the impact of post-rearrangement evolution of repeat DNA, we focused on two groups of recently evolved rearrangements: fissions on the Catalan *L. sinapis* branch and fusions on the Swedish *L. sinapis* branch. For fissions, we observed that in three out of six cases, Catalan *L. sinapis* had the highest average copy number (*p <* 0.05, determined by non-overlapping 95% confidence intervals [40]), in line with expectations (Fig. 3A). For fusions, Swedish *L. sinapis* had significantly lower copy numbers in 8 out 14 cases (Fig 3B). Thus, *L. sinapis* fissions and fusions follow the expectations overall and indicate that fissions and fusions can modulate genome size.

**Figure 3.**
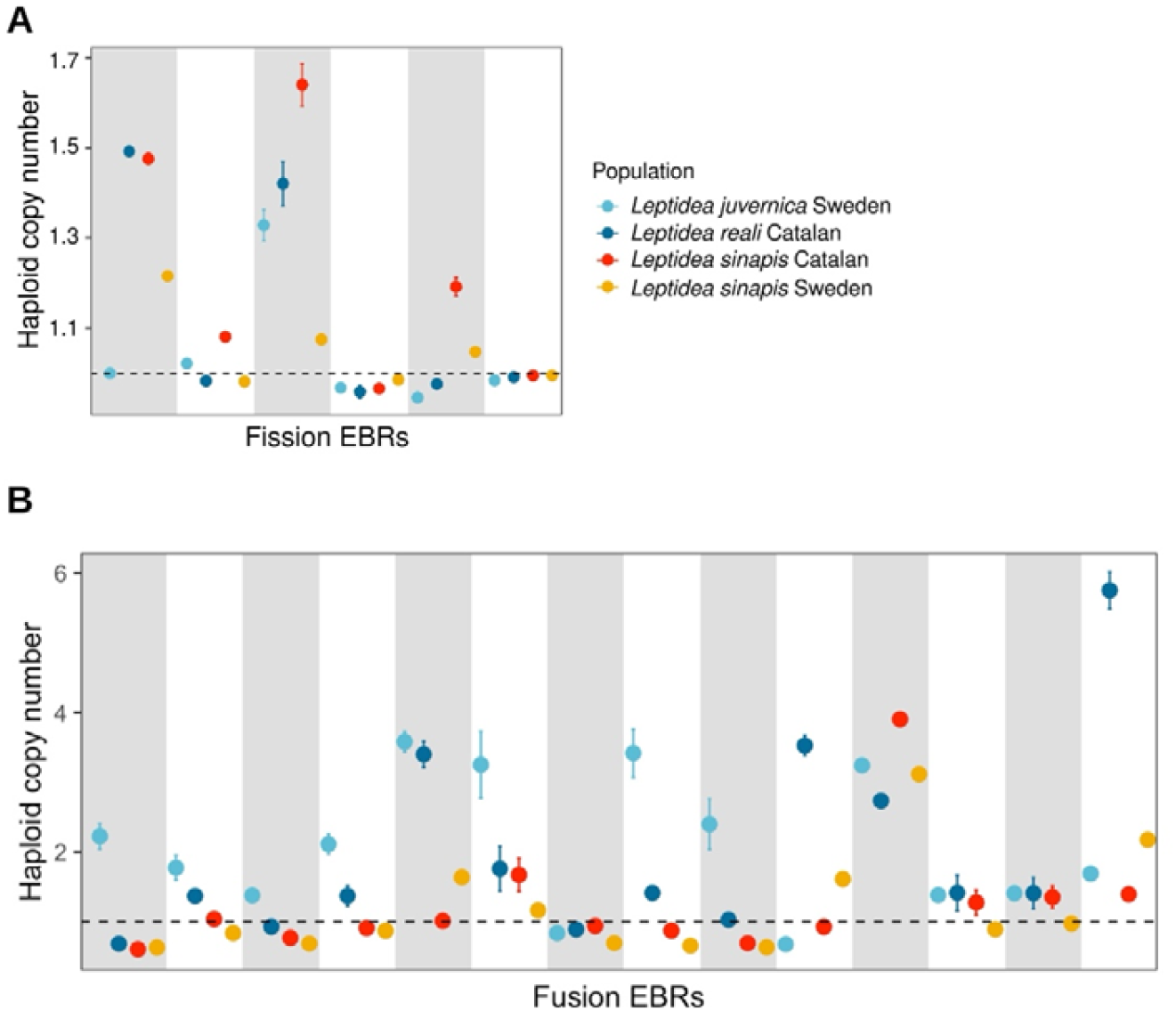
Evolution of copy number variation at EBRs. (A) Average haploid copy number of fission EBRs in the Catalan *L. sinapis* lineage. (B) Average haploid copy number of fusion EBRs in the Swedish *L. sinapis* lineage. Bars represent 95% confidence intervals and are often too short to be visible. Each rectangle represents one separate EBR. Haploid copy number for each population was estimated from read depth data of 12-96 individuals per population (see *Methods*).

## Discussion

Cryptic wood white butterflies have one of the highest rates of chromosome fissions and fusions observed in any clade [41]. We here leveraged this property to study the determinants of rearrangements in this clade. Most rearrangements in *Leptidea* are found in repetitive regions of the genome, but with only weak associations with TEs. Instead, satellite DNA is the overall strongest predictor of rearrangements. This may sound surprising for holocentric organisms, which generally have a more homogenous content of repeats in the genome compared to monocentric organisms, which have centromeres composed of large satellite DNA arrays [42]. Even so, satellite DNA often occurs in large arrays (>1 Mb) in *L. sinapis* [29], somewhat reminiscent of the chromosome structure of monocentric organisms. Associations between satellite DNA and fissions and fusions, respectively, have also been observed in holocentric *Rhynchospora* sedges, though they have smaller satellite clusters (3-16 kb) and a more homogenous repeat structure – typical of holocentric species [33]. This hints at some commonalities in large-scale rearrangements between monocentric- and holocentric taxa, with satellite DNA involvement, regardless of array size.

It is likely that a considerable part of the association between rearrangements and annotation features is driven by mutation propensity, though we acknowledge that fixation of rearrangements could have been influenced by non-neutral processes. This would confound statements about mutation with the effects of selection and meiotic drive. The latter effect has been inferred from transmission distortion patterns observed in F_2_ offspring between Swedish and Catalan *L. sinapis*, where the most consistent effect was a bias towards the Catalan unfused state at derived fusions in the Swedish population [24]. The strong association between satellite DNA and fusion EBRs suggests that this sequence type may contribute to the proposed meiotic drive.

Support for a mutational interpretation of the enrichment signals we observed comes from similarities in patterns observed in other taxa. In line with studies of human Robertsonian fusions [10], we observe associations between satellite DNA, ribosomal DNA and fusions. For humans, it is believed that the role of rDNA is indirect. Ribosomal clusters are present on two chromosomes undergoing fusion and their colocalisation in the nucleolus is believed to increase the opportunity for ectopic recombination between inverted segments of satellite DNA [10]. For *Leptidea*, the two main rDNA clusters in the reference genome are present on chromosomes 34 and 45, but these have not fused with one another. While there could be unassembled clusters on other chromosomes, fluorescent *in-*situ hybridisation of rDNA in *Leptidea* chromosome spreads suggests that only two main clusters are present [21]. Therefore, the involvement of rDNA clusters in rearrangements in *Leptidea* is still an open question and rearrangements could instead be mediated by repeats found in for example intergenic spacers. Translocation EBRs also showed an enrichment in rDNA clusters and segmental duplications. As previously said, it is possible that they are equivalent to fusions but with the retention of one DNA fragment. However, we cannot exclude that some putative translocations represent sequences of fission and fusion events occurring on the same branch, especially given the high rate of rearrangements in *Leptidea*.

Less is known about the mechanisms generating fissions compared to fusions. In general, a model of random breakage has been superseded by the fragile site breakage model: simply put, some regions are more prone to break than others [7,32,43,44]. The enrichment of segmental duplications, satellite- and the smaller microsatellite DNA (simple repeats) at fission EBRs in *Leptidea* supports this model, since rare fragile sites are associated with these repeat types in other systems [7,9,45,46]. However, it cannot be excluded that some portion of these enrichments has more to do with the survival of the rearrangement after the mutational event. We observed an increase in copy number of DNA in recent fission EBRs, possibly since telomere reformation takes time (>40 years in a line of silk moths, *Bombyx mori*), which poses a threat for degradation of genes in subtelomeric regions (without substrate for telomerase binding, the chromosomes get shorter for each cell division) [17,47]. Consequently, successful fissions can perhaps only form if there are longer stretches of spacer DNA that can buffer more vital DNA before proper telomere reformation.

Future experimental scoring of *de novo* fusions, fissions and translocations will help answer outstanding questions on the mutational determinants of rearrangements in *Leptidea* and other taxa. For example, fusions and fissions may occur frequently in gene-rich regions, but have negative fitness consequences and therefore not reach frequencies that are detectable in small sample sets. Rapidly decreasing costs for sequencing methods that provide long-range phase information suggests that such approaches soon will be accessible also for studies of non-model organisms [48]. That said, we do provide an unprecedented view of on the associations between annotation features and chromosomal rearrangements in butterflies, revealing striking commonalities across both deeply divergent branches of the tree of life and fundamentally different chromosome structures.

## Materials and Methods

### Genome assemblies

We used nine previously generated chromosome-level assemblies in this study. One male and female assembly each of Swedish and Catalan *L. sinapis*, Catalan *L. reali* and Swedish *L. juvernica* [22]. These assemblies were built using 10X linked read sequencing as backbone and scaffolded with Dovetail Hi-C long-range contact maps. In addition, we used a high-quality reference genome based on a male *L. sinapis* from Asturias in northwest Spain, sequenced with PacBio HiFi and scaffolded with Arima Hi-C [34].

### Inference of chromosomal rearrangements and evolutionary breakpoint regions

Evolutionary breakpoint regions (EBRs) were inferred based on ancestral gene order reconstructions at internal nodes of the species tree with single-copy BUSCO genes. Single-copy BUSCOs were recovered from all nine assemblies using BUSCO v5.7.1 with the lepidoptera_odb10 database [49]. Orthologous gene sets at internal nodes of the species tree were identified using OrthoFinder v2.5.5 [50]. Ancestral gene order was reconstructed for each internal node using the Algorithm of Gene Order Reconstruction in Ancestors (AGORA v3.1; [51]). We applied AGORA’s basic algorithm to the single-copy BUSCOs together with the inferred ortholog sets. Following ancestral reconstruction, pairwise gene alignments between extant genomes and internal nodes were generated using the auxiliary AGORA script misc.compareGenomes.py with mode=printOrthologuesList. Evolutionary breakpoint regions were identified per-species, per-chromosome synteny comparisons between present-day chromosomes and reconstructed ancestral karyotypes at each internal node. An EBR was inferred when a present-day chromosome shared synteny with multiple ancestral karyotypes or when an ancestral karyotype shared synteny with multiple present-day chromosomes. In such cases, a breakpoint was defined between two consecutive single-copy BUSCO genes where the ancestral karyotype or present-day chromosome identity switched. When both BUSCO genes were located on the same ancestral karyotype but on different present-day chromosomes, the event was annotated as a split, whereas when the BUSCO genes were located on different ancestral karyotypes but on the same present-day chromosome, the event was annotated as a merger. Splits and mergers represent fissions and fusions events except when both occur on the same branch, which we instead called as putative translocations. Splits and mergers occurring on the same branch could represent two events in a specific order: first a fission then a fusion or only a single molecular event in the case of a translocation (e.g. a double-strand break repaired by an ectopic template on another chromosome). We chose the latter interpretation since we consider it more parsimonious. For more details, see the Supplementary Methods.

### Phylogenetic reconstruction

We inferred a phylogeny of all nine genome assembly samples using autosomal single-copy BUSCO genes present in all samples (n = 3454). Individual amino acid sequences of the filtered set of single-copy BUSCO genes were aligned using the mafft-linsi algorithm (v.7.526; [52]). These alignments were then trimmed using trimAl (v.1.5.0; [53]) using a gap threshold of 0.2. Following trimming, alignments were concatenated and the full alignment was visually inspected in AliView (v.1.30; [54]). Single-gene alignments with more than 25% missing data were excluded. We reconstructed a phylogeny based on the concatenated alignment with IQ-TREE (v.3.0.1; [55]). For more details on model selection, see the Supplementary Methods. We calculated the rate of interchromosomal rearrangement events along the branches of the phylogeny, normalising by the amino acid substitution rate per branch. We excluded the branch to the outgroup (*L. juvernica*) as EBRs along this branch are unpolarisable.

### Sequence tracks and segmental duplications

We obtained the coordinates of protein-coding genes from Ensemble gene annotation (update 2021-11). Satellite DNA was obtained from REF: [29], who used CHRISMAPP on the Asturian *L. sinapis* reference genome. We *de novo* predicted repeat consensus sequences by first running RepeatModeler v2.0.4 [56]. We then masked the Asturian reference genome using RepeatMasker v4.1.5 [57]. We obtained separate sets of TEs based on class (or sub-class) after intersection with predicted satellites, since the most common satellite was incorrectly inferred as LINE repeat (see *Results*). Ribosomal DNA was inferred using Barrnap v0.9 [58]. We first predicted segmental duplications in a hardmasked version of the Asturian reference genome using BISER v1.4 [59]. In addition, we used StainedGlass v0.6 [60], employing within-chromosome all-vs-all alignments in 2 kb windows. By visual comparison we noted that BISER missed or underestimated the extent of most large-scale segmental duplications which were beyond-doubt clearly visible in the StainedGlass output (Additional file 1). Therefore, we developed an algorithm to define regions containing large within-chromosome segmental duplications from the StainedGlass alignments. The main idea of the algorithm was to identify stretches of pairwise alignments that are characteristic of segmental duplications (Fig. S4-5). For more details see the Supplementary Methods.

### Statistical association between annotation features and EBRs

We calculated odds ratios and performed permutation tests similar to Boman *et al*. [1], using custom scripts. Briefly, overlap was quantified based on the number of base pair overlap. Odds ratios were calculated based on the observed overlap between an EBR set and an annotation set, normalising by how much of the genome these two sets occupy. A permutation was made by randomly permuting the ranges of the annotation set (keeping the EBR set constant), and computing the random overlap. We performed 1000 permutations and calculated a two-tailed *p*-value [1]. The false discovery rate of the *p*-values was corrected separately per EBR set (fissions, fusions and translocations), using the Benjamini-Hochberg method.

### Inference of structural variants from short-read sequence data

Structural variants (SVs) were identified in 10 male each of Swedish and Catalan *L. sinapis*, Catalan *L. reali* and Swedish *L. juvernica* using the Parliament2 (v0.1.11; [61]) pipeline. For more details see the Supplementary Methods.

### Inference of copy number variants based on read depth

We inferred larger copy number variants to study sequence evolution at fissions in the Catalan *L. sinapis* lineage and fusions in the Swedish *L. sinapis* lineage. To infer copy number variants, we used the simple principle that a copy more or less frequent than the reference sequence will show a read-depth congruent with its copy number. We used whole-genome sequence data from 96 Swedish *L. sinapis*, 12 Catalan *L. sinapis*, 12 Swedish *L. juvernica* and 12 Catalan *L. reali* [22,62,63]. Included in these numbers are, 10X data from the assemblies of each population. In addition, we used outgroup data from 2 individuals each of *L. morsei, L. amurensis* and *L. lactea* [64], from which no genome assemblies exist. Raw sequence reads were trimmed using TrimGalore v0.6.1 [65], a wrapper for CutAdapt v4.0 [66]. Filtered reads were mapped to the Asturian reference genome using bwa *mem* v0.7.17 [67]. We inferred CNVs in 10 kb windows ControlFREEC v11.6 [68]. This method does not identify particular SVs but instead uses the read depth information in larger windows, making it suitable for our purpose here. We chose 10 kb windows since this size is larger than most single TEs in the genome. Note that the segmentation of the reference genome in read depths that underlies the CNV calling in Control-FREEC is not restricted to 10 kb windows and allows sharing of read-depth information across larger scales, which allows for more precise CNV calls. Thus 10 kb should be seen as a minimal window size. See Additional file 2 for CNV maps across all chromosomes.

## Acknowledgements

We thank Mahwash Jamy for advice on phylogenomics and Diogo Cavalcanti Cabral-de-Mello for sharing satellite DNA annotation files. The computations were enabled by resources in project UPPMAX 2025/2-462 provided by the National Academic Infrastructure for Supercomputing in Sweden (NAISS) at UPPMAX, funded by the Swedish Research

Council through grant agreement no. 2022-06725. This project was funded by a research grant from the Swedish Research Council to NB (grant no. 2019-04791). JB acknowledges support from The Birgitta Sintring Foundation, Lennanders Foundation, Royal Physiographic Society Lund (grant no. 45881) and the Swedish Research Council (grant no. 2025-00450).

## Data Accessibility

Previously published data is available on the European Nucleotide Archive under accession numbers: PRJEB66418, PRJEB21838, PRJEB56690 and PRJEB58697. Analysis code will be available on the GitHub repository: https://github.com/FilipThorn/Repeats-and-rearrangements-in-holocentric-butterflies and as a static copy on Zenodo (upon final manuscript version).

## Authors’ Contributions

Conceptualisation, F.T., and J.B.; data analysis, F.T., J.-L.C., J.B; writing – original draft, J.B.; writing – review & editing, F.T., J.-L.C., N.B., and J.B.

## Declaration of AI use

AI-assisted technologies have been used as a tool when scripting computer code.

