## Supplementary Information for "Determinants of chromosomal rearrangements in holocentric *Leptidea* butterflies"

Supplementary Methods

**Further details on the mapping of evolutionary breakpoint regions**

We excluded synteny blocks with fewer than three BUSCO genes supporting an identity switch. Due to its higher assembly quality, the Asturian *L. sinapis* reference genome was used as the reference coordinates for breakpoint inference. Single-copy BUSCO genes served as anchor points to transfer coordinates between assemblies and the Asturian reference genome. For each EBR, we inferred the genomic interval between the first BUSCO and the subsequent BUSCO in the Asturian *L. sinapis* coordinate system, as well as the interval between the second BUSCO and its preceding BUSCO, resulting in 1-2 genomic ranges per EBR corresponding to the regions flanking the breakpoint in the original coordinate systems. EBRs between within-population pairs on external branches were inferred directly based on aligned BUSCO genes, as ancestral reconstruction at these nodes did not include the Asturian reference genome. We excluded synteny blocks with fewer than five BUSCO genes for these switches, deciding on a more conservative filter were appropriate without the model of AGORA informing these directly. These EBRs were polarised based on EBRs that shared the same position identified on the branch leading to the external branches, which were identified with on the ancestral reconstruction.

**Model selection and branch support for the phylogenetic reconstruction**

IQ-TREE model selection was used to pick the best fitting substitution model based on a set of Le och Gascuel models (LG) with a gamma distribution (G) in combination with different sets amino acid profiles (C10-C60) to account for heterotachy and compositional heterogeneity of amino acids. The best fitting model was the LG+G4 substitution model. Branch support was calculated based on 1000 bootstraps.

**Description of the algorithm used to identify large within-chromosome segmental duplications**

We filtered for 2 kb pairwise alignments, at least 10 kb distant (“offset”) with an identity larger than 90%. To reduce noise from other repeats the algorithm moves sequentially along the sequence to find pairs of pairwise alignments with high identity and less than 3 kb absolute difference in offset. If for example there is a tandem duplication with a 100 kb monomer, then position 0 and 2000 in the beginning of the first repeat should both align to the start of the second repeat at 100,000 and 102,000 respectively, allowing for minor discrepancies. This filters away some true positive regions within larger stretches of segmental duplications momentarily, but they are salvaged later. We then take the remaining 2 kb alignments, their offsets and offset differences among pairs, and cluster them using the DBscan v1.2-0 package in *R* v4.4.2 [1,2]. This can create separate clusters at different offset levels for tandem repeats larger than dimers. Therefore, we include a subsequent step where we merged all alignment regions within 20 kb from each other, using BEDtools v2.31.1 *merge* [3]. Finally, we excluded ranges in which the sequence consisted of more than 50% satellite DNA, to define large segmental duplications with large monomer sizes specifically (Fig. S3).

**Structural variant calling from short-read data**

We went beyond annotation features such as (segmental duplications, TEs, rDNA and satellite DNA) and tested whether inter-chromosomal rearrangements were associated with smaller structural variants (SVs) since it is possible that structural heterozygosity could promote e.g. ectopic recombination, leading to fusions or translocations [41]. We identified structural variants using Parliament2 (v0.1.11; [4]) pipeline, running all its integrated callers: BreakDancer [5], Manta [6], CNVnator [7], LUMPY [8], and DELLY [9]. To account for potential cross-species mapping, stringent filtering criteria were applied at both the caller and sample levels. Only variants passing internal filters were retained, with additional thresholds: Manta, DELLY, and LUMPY calls required ≥25% of local depth in supporting reads, while BreakDancer and CNVnator calls were retained if their absolute length exceeded one kb and five kb, respectively. Filtered calls were then merged at the sample level using Jasmine (v1.1.5; [10]), retaining variant types and merging variants of the same type located within 1 kb. This step ensured that only variants supported by a minimum of two callers were retained for downstream analysis. Finally, a cohort-level VCF was generated by merging all single-sample VCFs using SURVIVOR (v1.0.7; [11]), applying the same merging criteria as in the sample-level step, merging variants of the same type located within 1 kb.

**Figure S1**: Map of polarised EBRs across the Asturian *L. sinapis* reference genome.

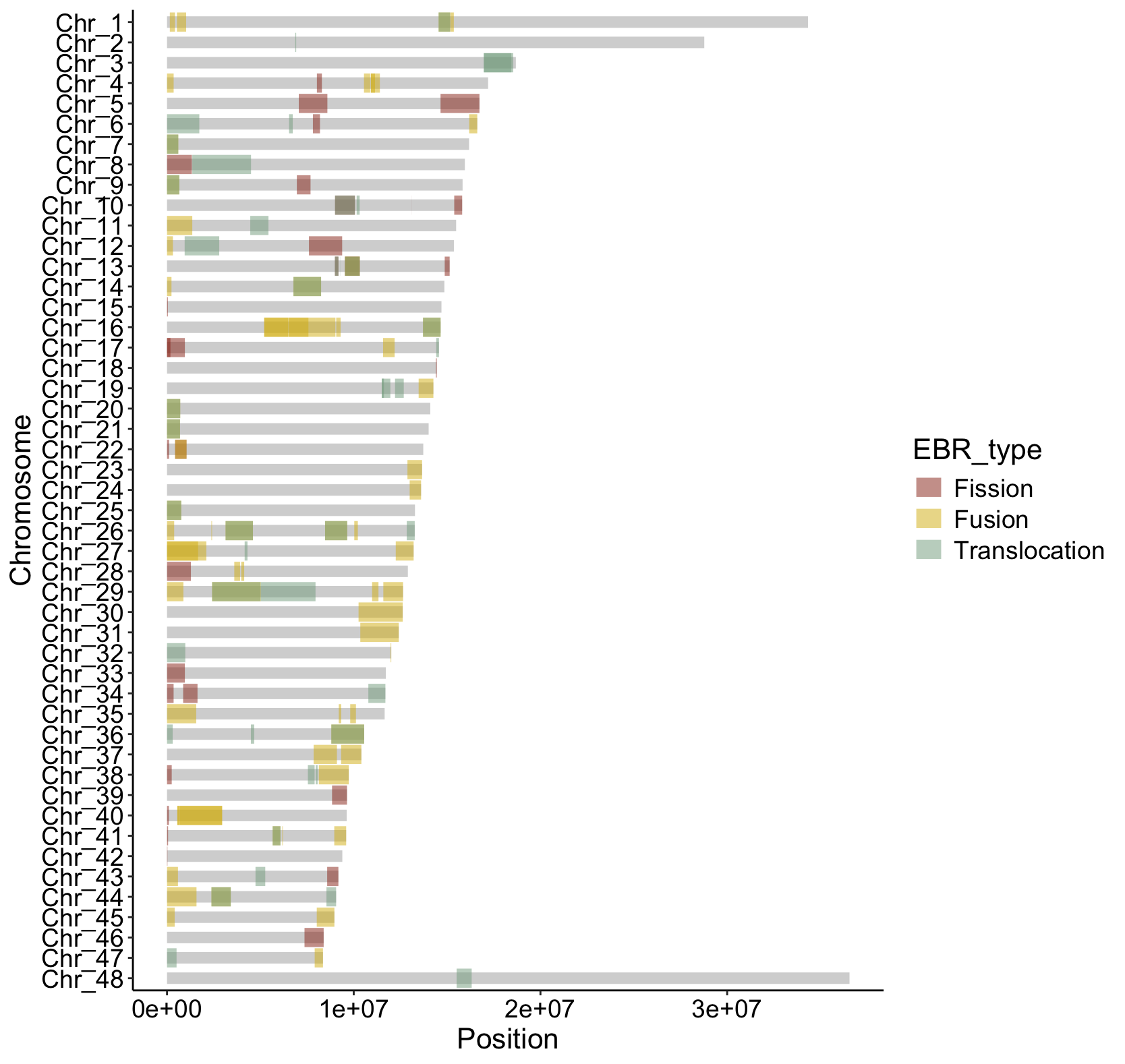

**Figure S2**: Map of rDNA annotations from barrnap. In the *Leptidea sinapis* reference genome there are two main clusters of rDNA, located on chromosomes 34 and 45. The rDNA unit is generally ca 10 kbp long with a ca 80-88 bp long (TAAG)_n_ microsatellite as the most common intergenic spacer, similar to what has been observed in the hepialid moth *Phymatopus californicus* [12]**.**

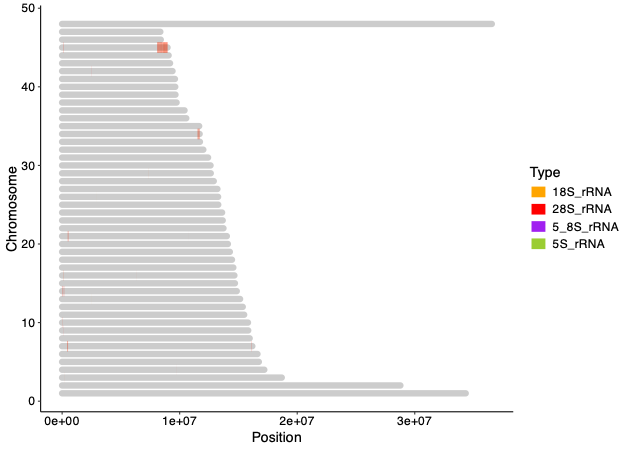

**Figure S3:** Genome-wide map of segmental duplications inferred from StainedGlass within-chromosome pairwise alignments in 2 kb windows. Our method to characterise segmental duplications from these alignments discovers most larger segmental duplications within chromosomes as determined by visual inspection of the alignment output (see Additional file 1). Using this method, ~5.4% of the reference genome is part of a segmental duplication. The method does not the characterise the monomers specifically and will therefore miss segmental duplications which are single-copy in a particular chromosome.

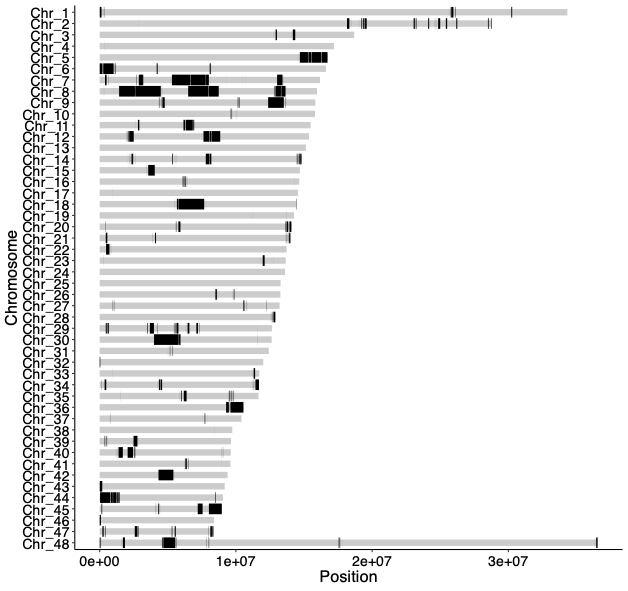

**Figure S4:** Examples of repeat structures in the Asturian *L. sinapis* reference genome. We generated within-chromosome all-vs-all pairwise alignments in 2 kb windows using StainedGlass. See Additional File 1 for whole-chromosome plots. The following plots were generated by using only the shown regions of chromosomes 8 and 19. (A) A large region (~2 Mb) with seven tandem copies of a segmental duplication. (B) A large (~2 Mb) cluster of satellite DNA (LepSat01-100) with higher order repeat structure. The smaller cluster close to position 12 Mb consists mainly of the satellite LepSat04−30. See Supplementary Figure 2 of Cavalcanti Cabral-de-Mello et al. [13], for full maps of satellite DNA across the genome.

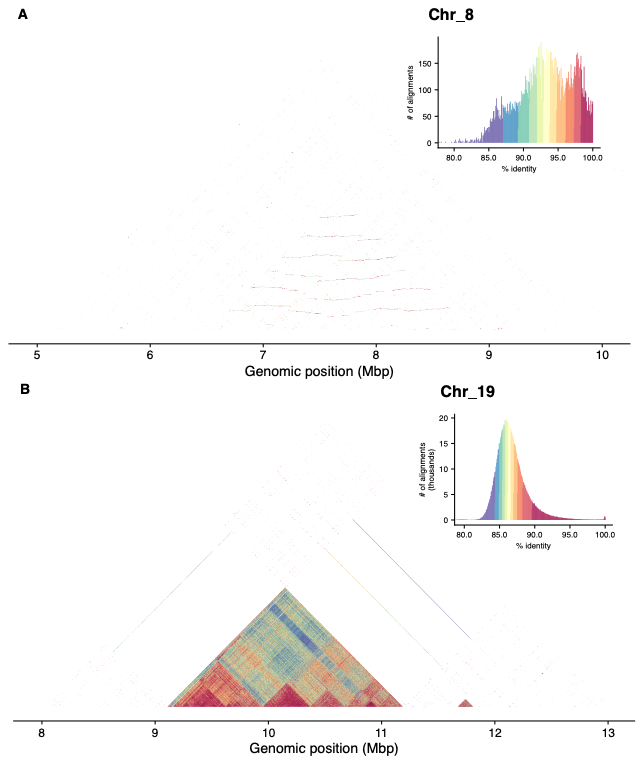

**Figure S5:** Detailed look at repeat structure and copy number variation at the evolutionary breakpoint region on chromosome 12 (from 7.5 to 9 Mb). The rearrangements here are a fission shared by the Catalan *L. sinapis* assemblies and a breakpoint on the branch to *L. juvernica* with unknown polarisation. (A) Detailed look at repeat structure shows that the assembly contains three tandem copies of a ca 400 kb monomer segmental duplication. Pairwise self-alignment was done using MAFFT with standard settings (<https://mafft.cbrc.jp/alignment/server/>, accessed 2025-10-06). Shown is the LAST plot. (B) Copy number variation in the same region in 10 kb windows from Control-FREEC. Grey means normal diploid copy number.

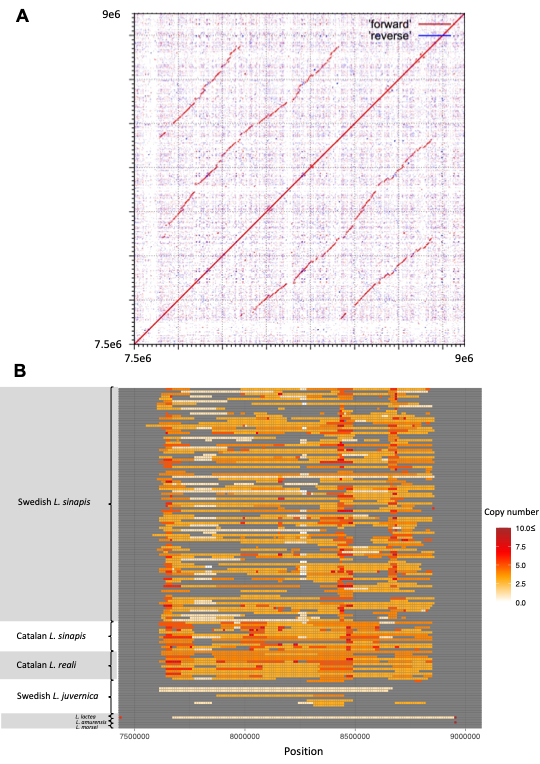

**Table S1.** RepBase hits for the RepeatModeler consensus sequence: rnd-1_family-0#LINE/RTE-BovB. With a consensus length of 400 nucleotides, encompassing 4 monomers of lepSat01-100. The hits are to the terminal inverted repeat region of *Academ* DNA transposons annotated in the bivalve, *Archivesica marissinica*. Such short hits are common to Repbase and are not long enough to make homology certain. We also blasted the rnd-1_family-0#LINE/RTE-BovB sequence to the NCBI conserved domain database and obtained no hits. We therefore continue to consider lepSat01-100 as a satellite sequence with unknown origin and not a LINE nor a DNA transposon.

| Query start | Query end | Reference | Ref. start | Ref. end | Similarity |
| --- | --- | --- | --- | --- | --- |
| 115 | 168 | Academ-9B_ArMa | 110 | 158 | 82.35% |
| 218 | 275 | Academ-80_ArMa | 104 | 153 | 83.02% |
| 318 | 375 | Academ-80_ArMa | 104 | 153 | 83.02% |

**Table S2.** Permutation analysis for short structural variants inferred using 10 males each of *L. sinapis* from Catalonia and Sweden, *L. reali* from Catalonia and *L. juvernica* from Sweden. P-values were adjusted according to the Benjamini-Hochberg method per EBR type. Structural variants were tested separately per population.

| **Odds ratio** | ***p*-value** | **Adjusted *p-*value** | **Comparison** |
| --- | --- | --- | --- |
| 0.998993 | 0.986 | 0.986 | Fission EBRs vs. SV.Ljuvernica_SWE |
| 1.53313 | 0.264 | 0.352 | Fission EBRs vs. SV.Lreali_CAT |
| 0.51063 | 0.19 | 0.352 | Fission EBRs vs. SV.Lsinapis_CAT |
| 1.30656 | 0.184 | 0.352 | Fission EBRs vs. SV.Lsinapis_SWE |
| 0.683807 | 0.038 | 0.108 | Fusion EBRs vs. SV.Ljuvernica_SWE |
| 0.908652 | 0.79 | 0.790 | Fusion EBRs vs. SV.Lreali_CAT |
| 0.566614 | 0.054 | 0.108 | Fusion EBRs vs. SV.Lsinapis_CAT |
| 0.810712 | 0.168 | 0.224 | Fusion EBRs vs. SV.Lsinapis_SWE |
| 0.778059 | 0.208 | 0.266 | Translocation EBRs vs. SV.Ljuvernica_SWE |
| 1.39184 | 0.266 | 0.266 | Translocation EBRs vs. SV.Lreali_CAT |
| 0.50171 | 0.066 | 0.140 | Translocation EBRs vs. SV.Lsinapis_CAT |
| 0.724438 | 0.07 | 0.140 | Translocation EBRs vs. SV.Lsinapis_SWE |

**Table S3.** Permutation analysis for SVs and satellite DNA. We hypothesised that short-read SV calling would struggle in highly repetitive regions and therefore largely be depauperate in satellite DNA regions. To confirm this, we compared satellite DNA and SVs, and all SV types were significantly absent (*p <* 0.001) in satellite DNA with odds ratios ranging from 0.03-0.11.

| **Odds ratio** | ***p*-value** | **Adjusted *p-*value** | **Comparison** |
| --- | --- | --- | --- |
| 0.0560161 | 0 | 0 | Satellite DNA vs. SV.Ljuvernica_SWE |
| 0.032761 | 0 | 0 | Satellite DNA vs. SV.Lreali_CAT |
| 0.112611 | 0 | 0 | Satellite DNA vs. SV.Lsinapis_CAT |
| 0.0860769 | 0 | 0 | Satellite DNA vs. SV.Lsinapis_SWE |

**Additional file 1.** StainedGlass output for all chromosomes of the Asturian *L. sinapis* reference genome.

**Additional file 2.** ControlFREEC output in 10 kb for all chromosomes and individuals. Copy number variation (dosage) is capped at 10x.
