## Additional File 1 for "Determinants of chromosomal rearrangements in holocentric *Leptidea* butterflies"

### Chr_1

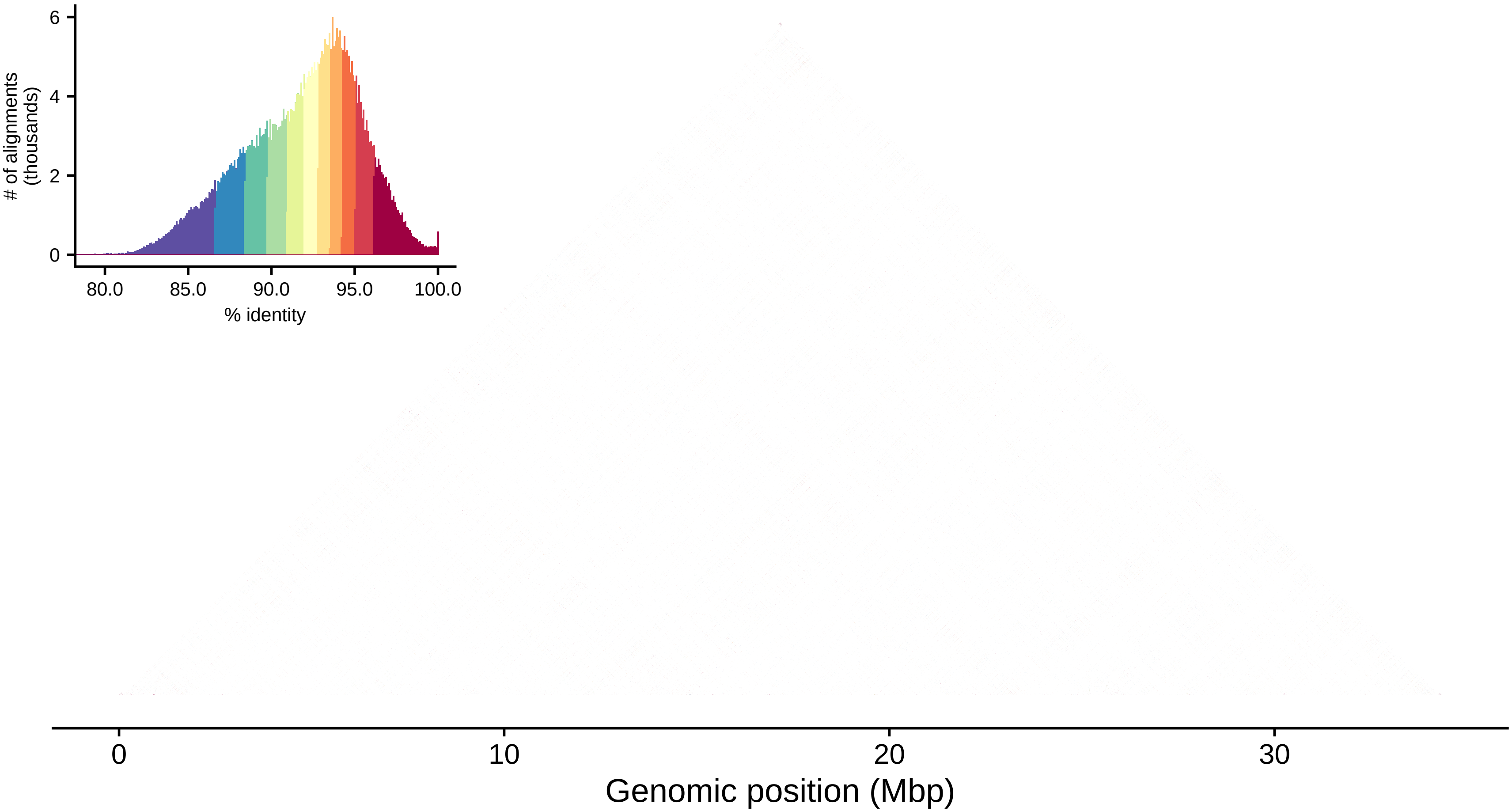

6
4
2
0
### of alignments
(thousands)
80.0
85.0
90.0
95.0
100.0
% identity
0
10
20
30
Genomic position (Mbp)

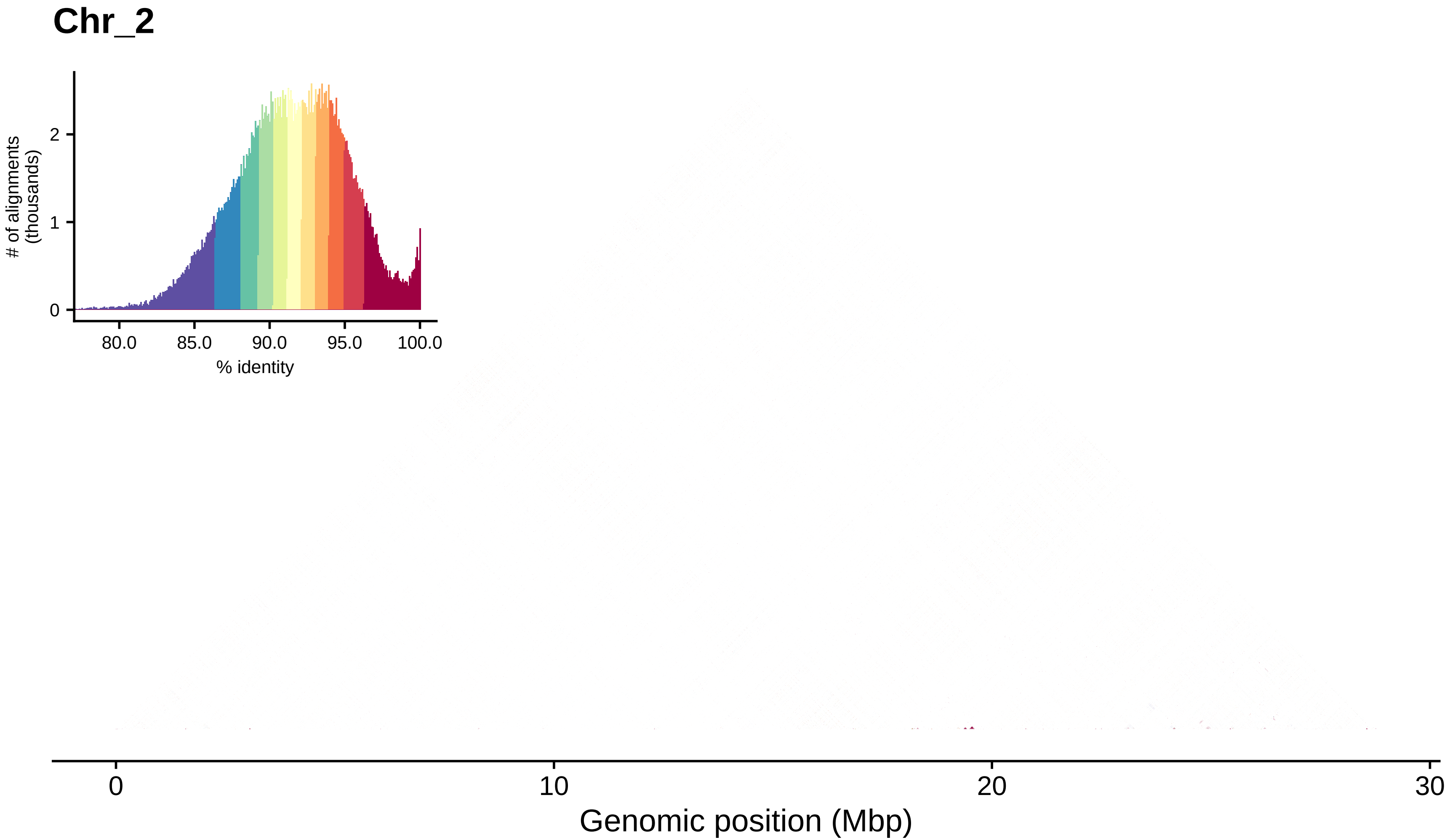
Chr_2
### of alignments (thousands)
0
1
2
80.0
85.0
90.0
95.0
100.0
% identity
0
10
20
30
Genomic position (Mbp)

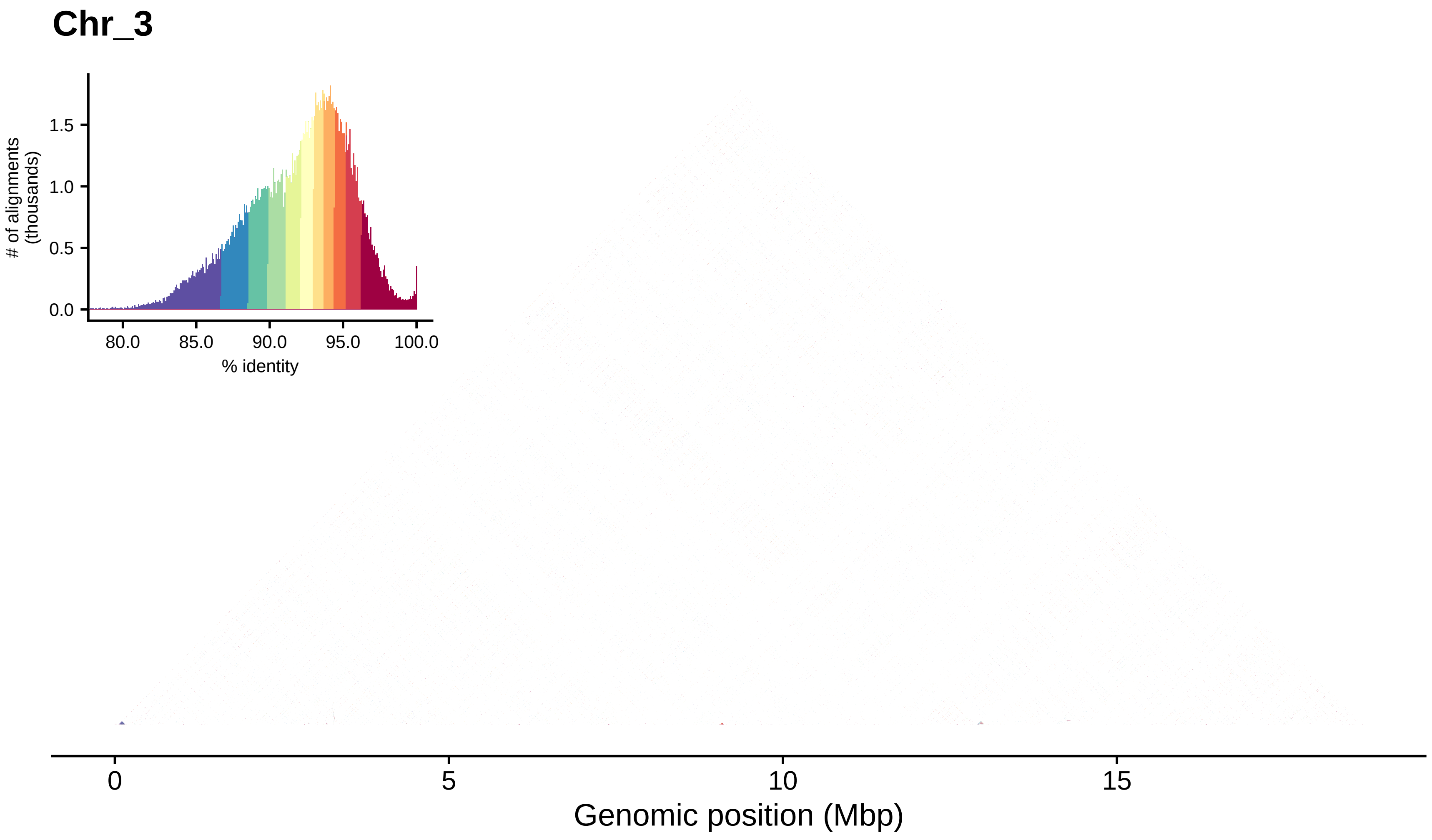

Chr_3
### of alignments (thousands)
0.0
0.5
1.0
1.5
80.0
85.0
90.0
95.0
100.0
% identity
0
5
10
15
Genomic position (Mbp)

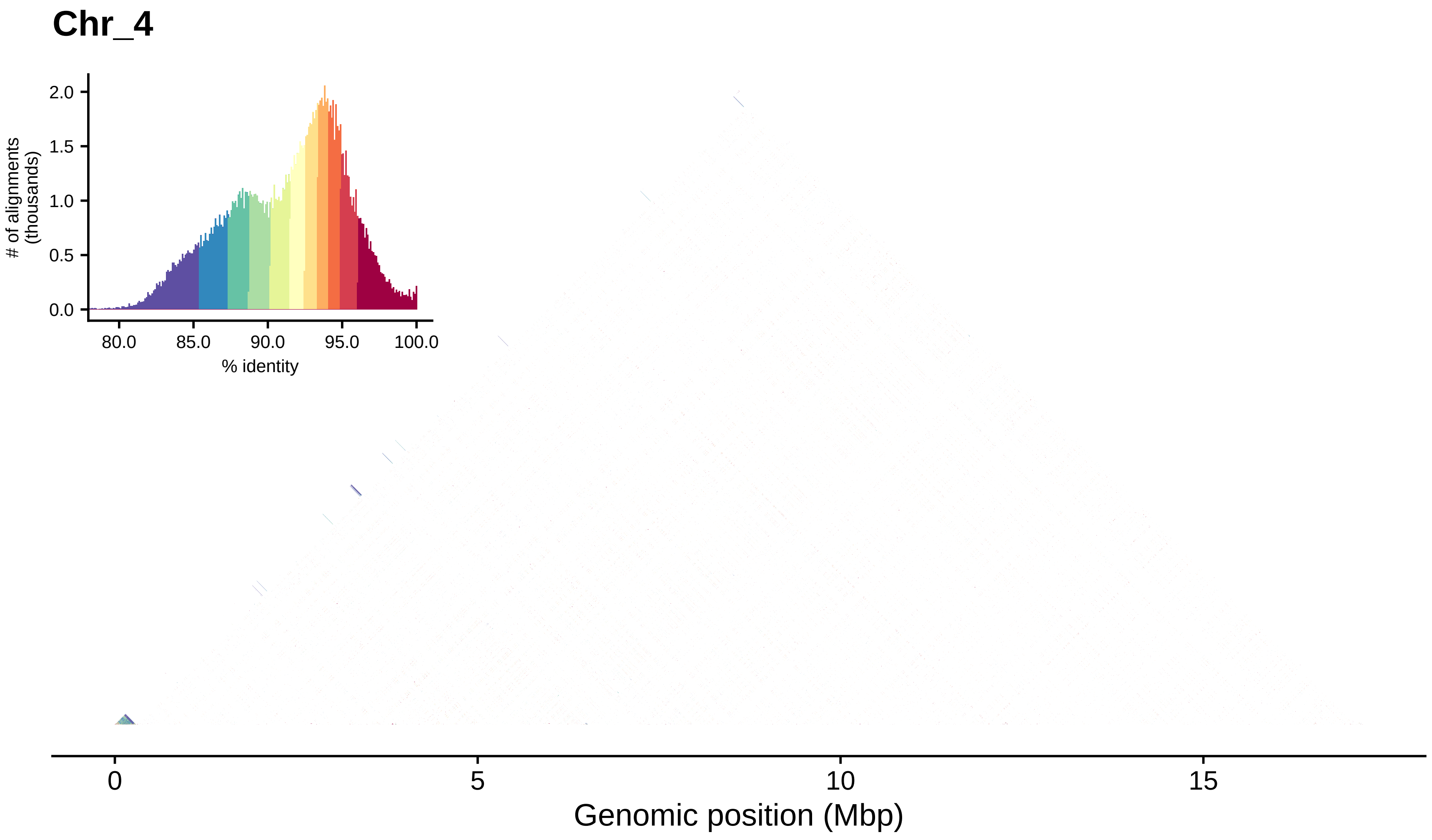
Chr_4
### of alignments (thousands)
2.0
1.5
1.0
0.5
0.0
80.0
85.0
90.0
95.0
100.0
% identity
0
5
10
15
Genomic position (Mbp)

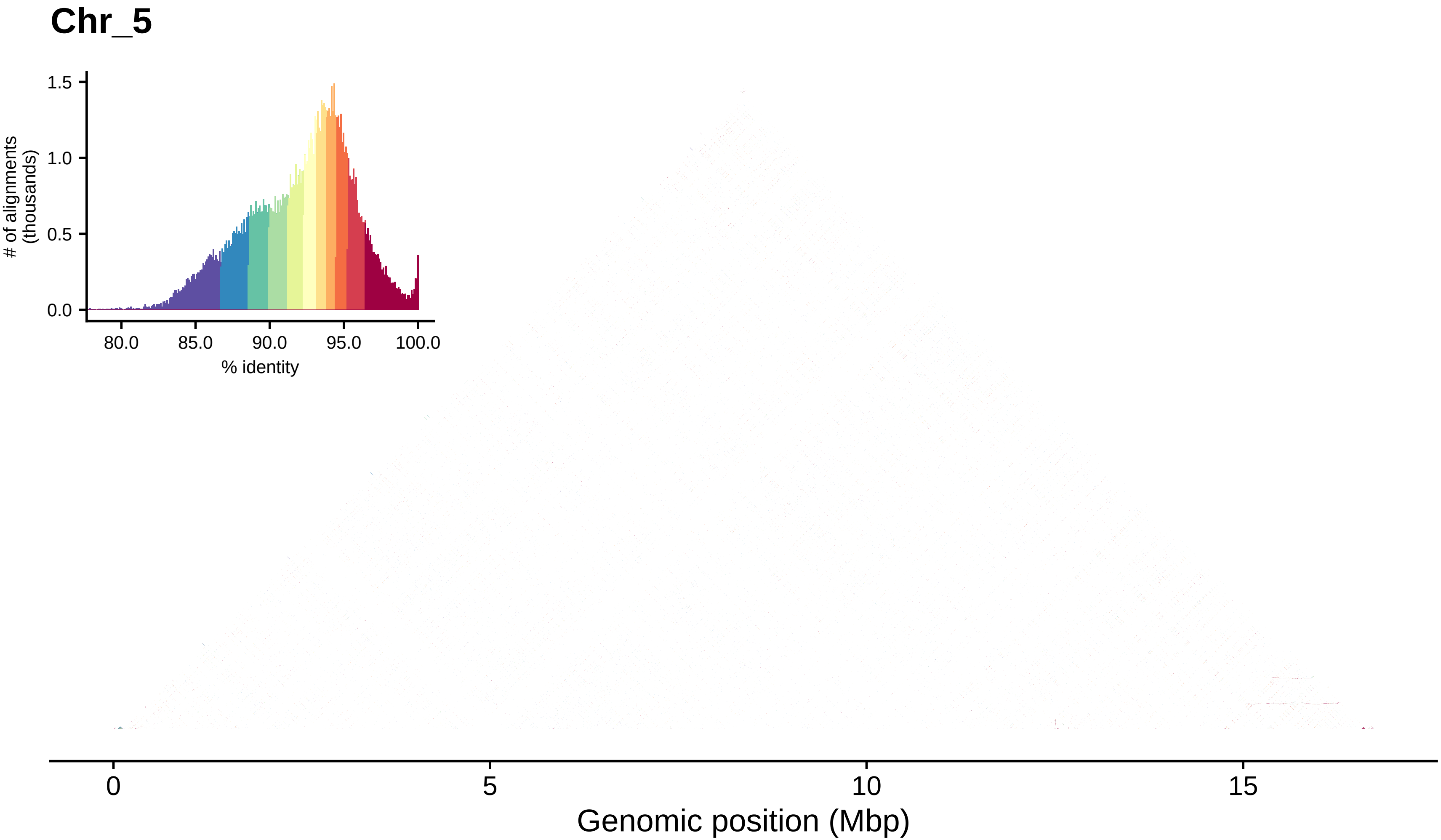
Chr_5
### of alignments (thousands)
1.5
1.0
0.5
0.0
80.0
85.0
90.0
95.0
100.0
% identity
0
5
10
15
Genomic position (Mbp)

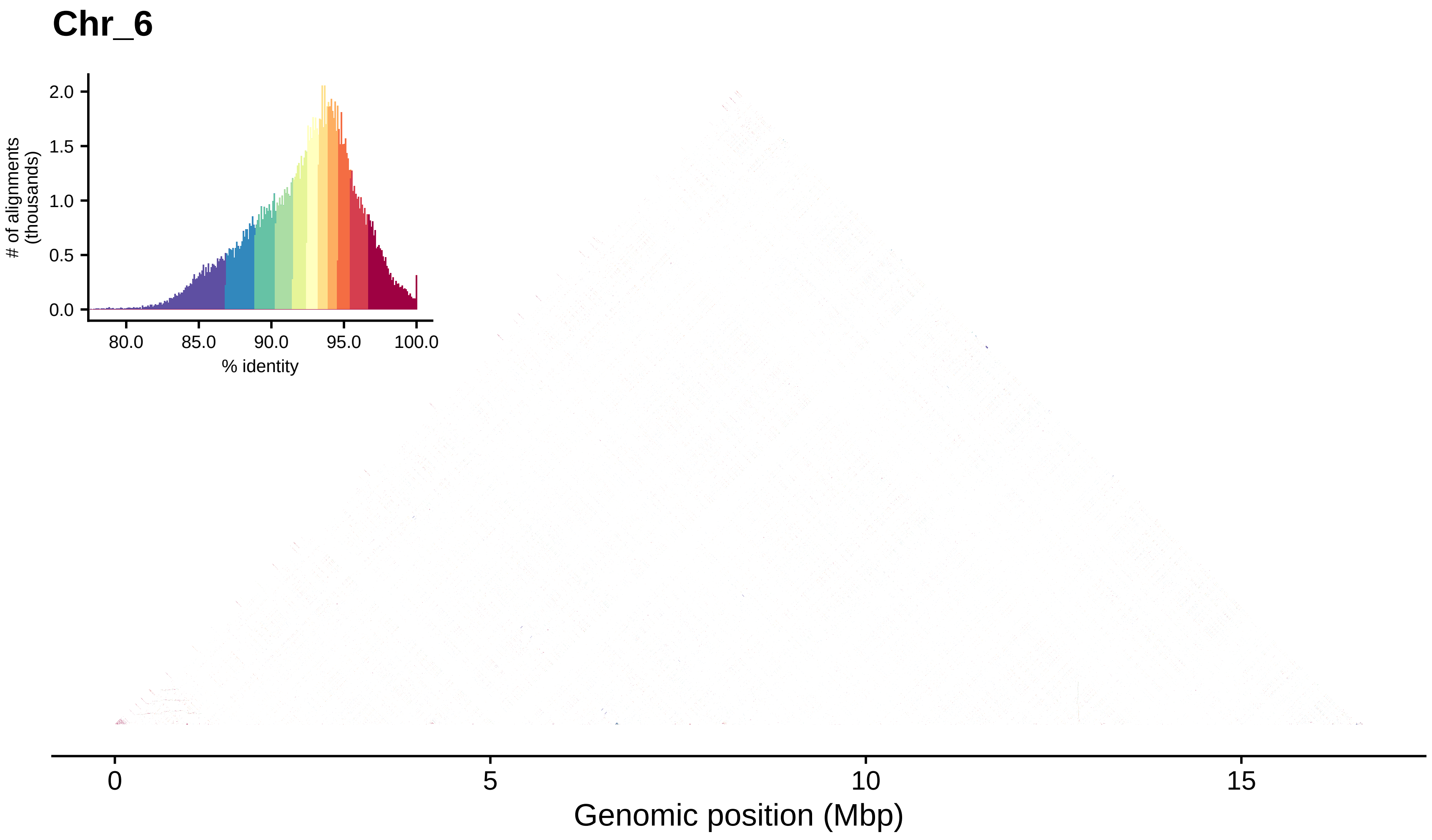
Chr_6
### of alignments (thousands)
2.0
1.5
1.0
0.5
0.0
80.0
85.0
90.0
95.0
100.0
% identity
0
5
10
15
Genomic position (Mbp)

#### Chr_7

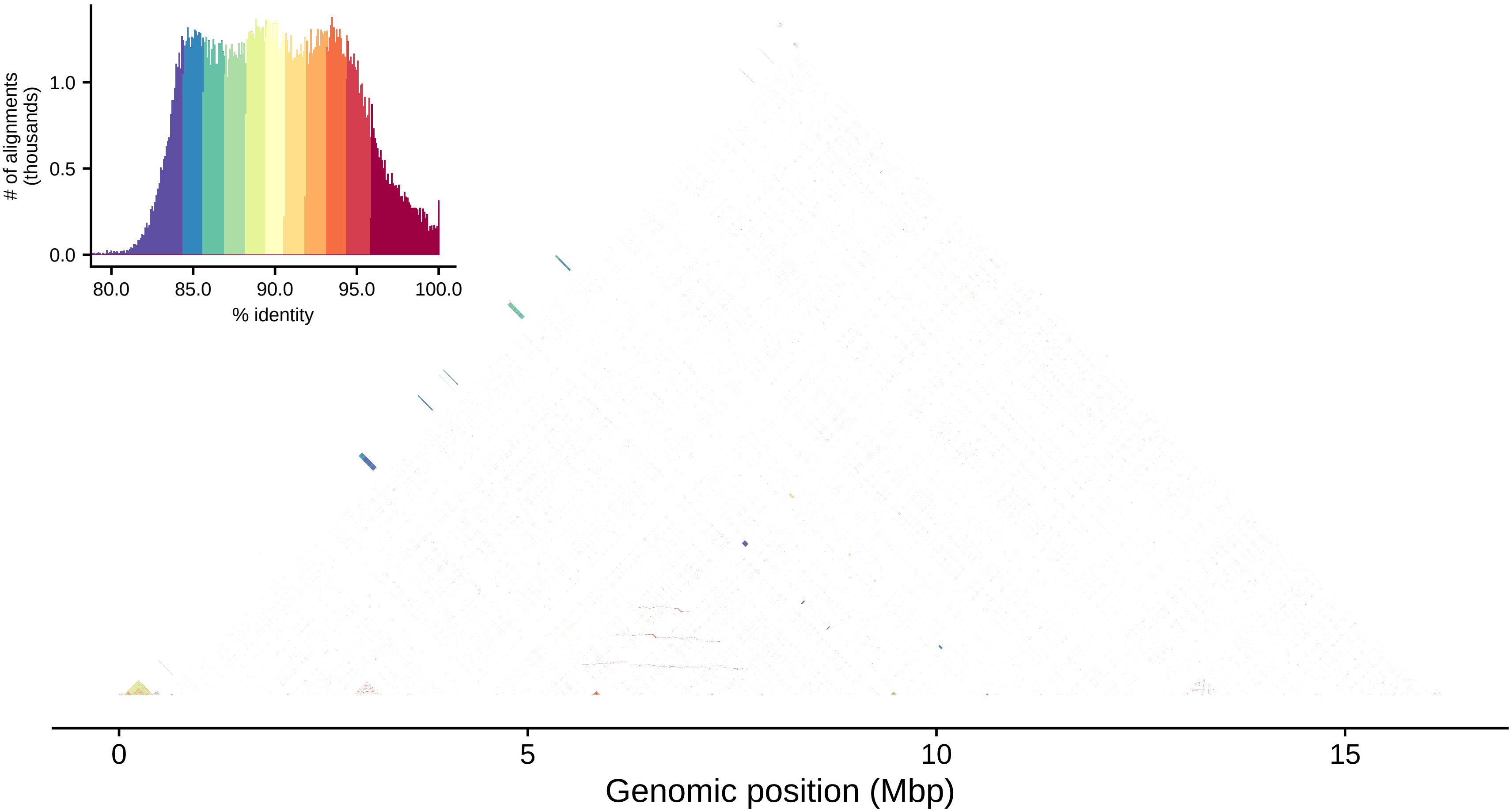

### of alignments (thousands)
0.0
0.5
1.0
80.0
85.0
90.0
95.0
100.0
% identity
0
5
10
15
Genomic position (Mbp)

#### Chr_8

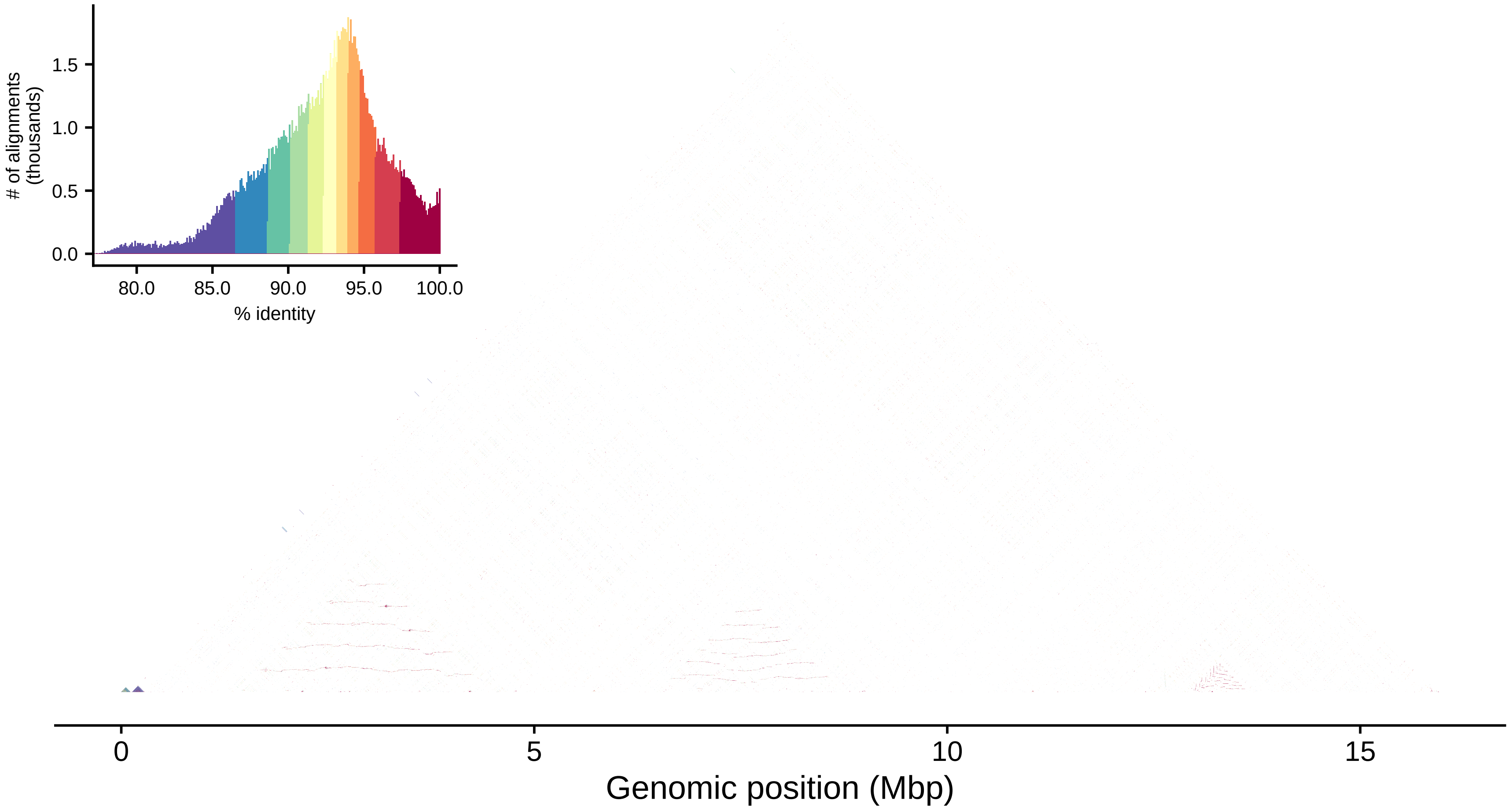

### of alignments (thousands)
0.0
0.5
1.0
1.5
80.0
85.0
90.0
95.0
100.0
% identity
0
5
10
15
Genomic position (Mbp)

#### Chr_9

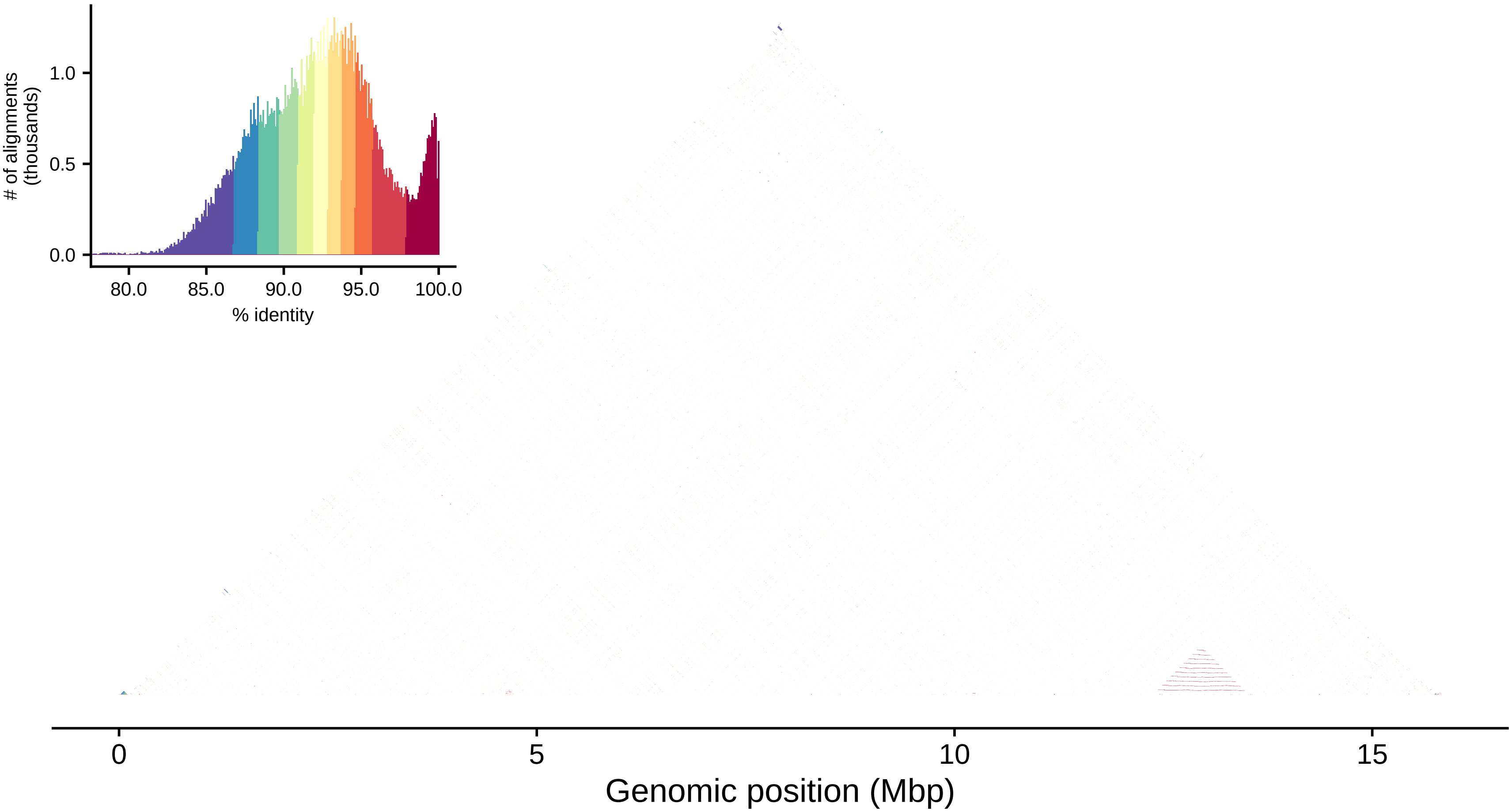

### of alignments
(thousands)
1.0
0.5
0.0
80.0
85.0
90.0
95.0
100.0
% identity
0
5
10
15
Genomic position (Mbp)

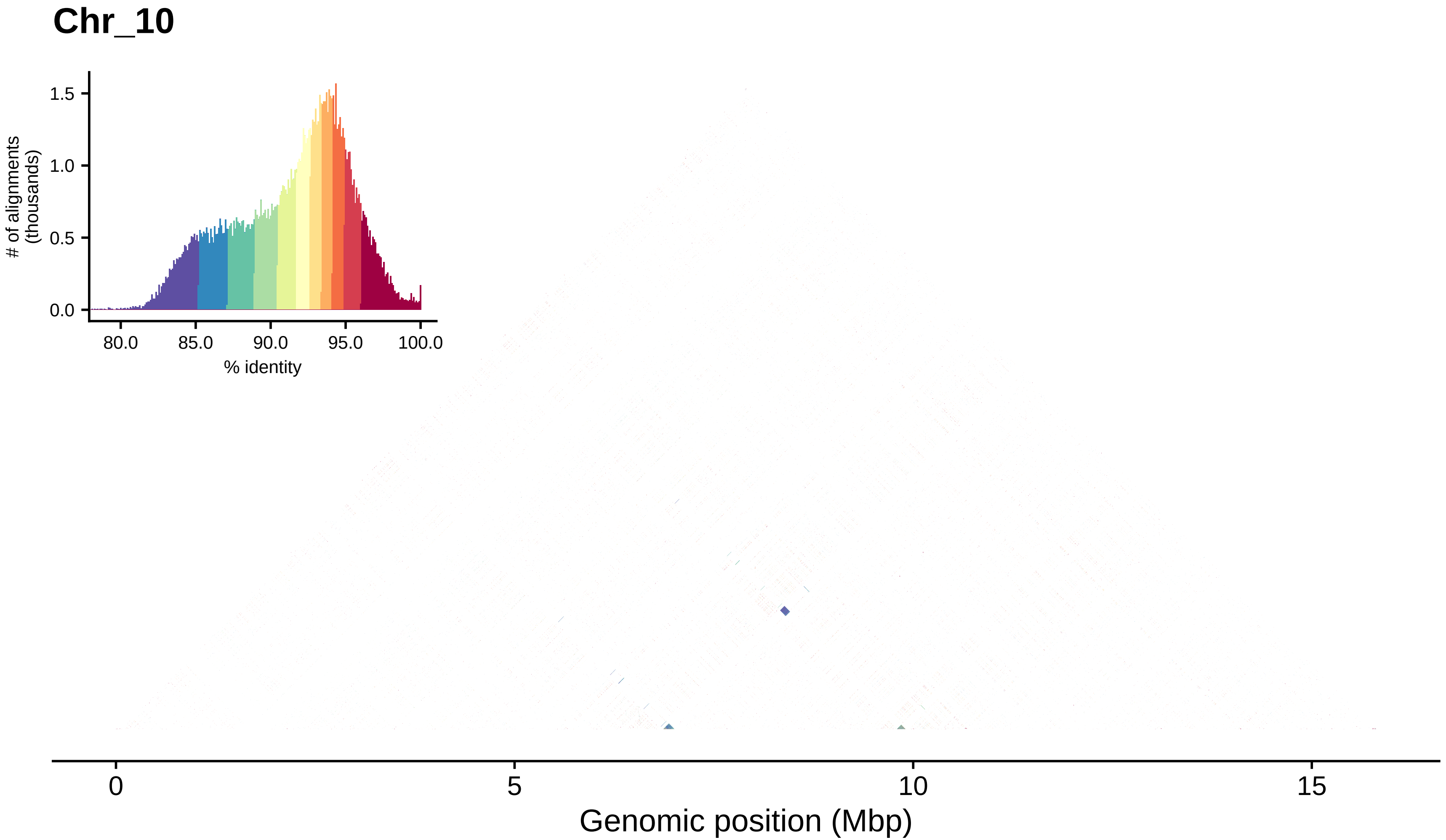

Chr_10
1.5
1.0
0.5
0.0
### of alignments
(thousands)
80.0
85.0
90.0
95.0
100.0
% identity
0
5
10
15
Genomic position (Mbp)

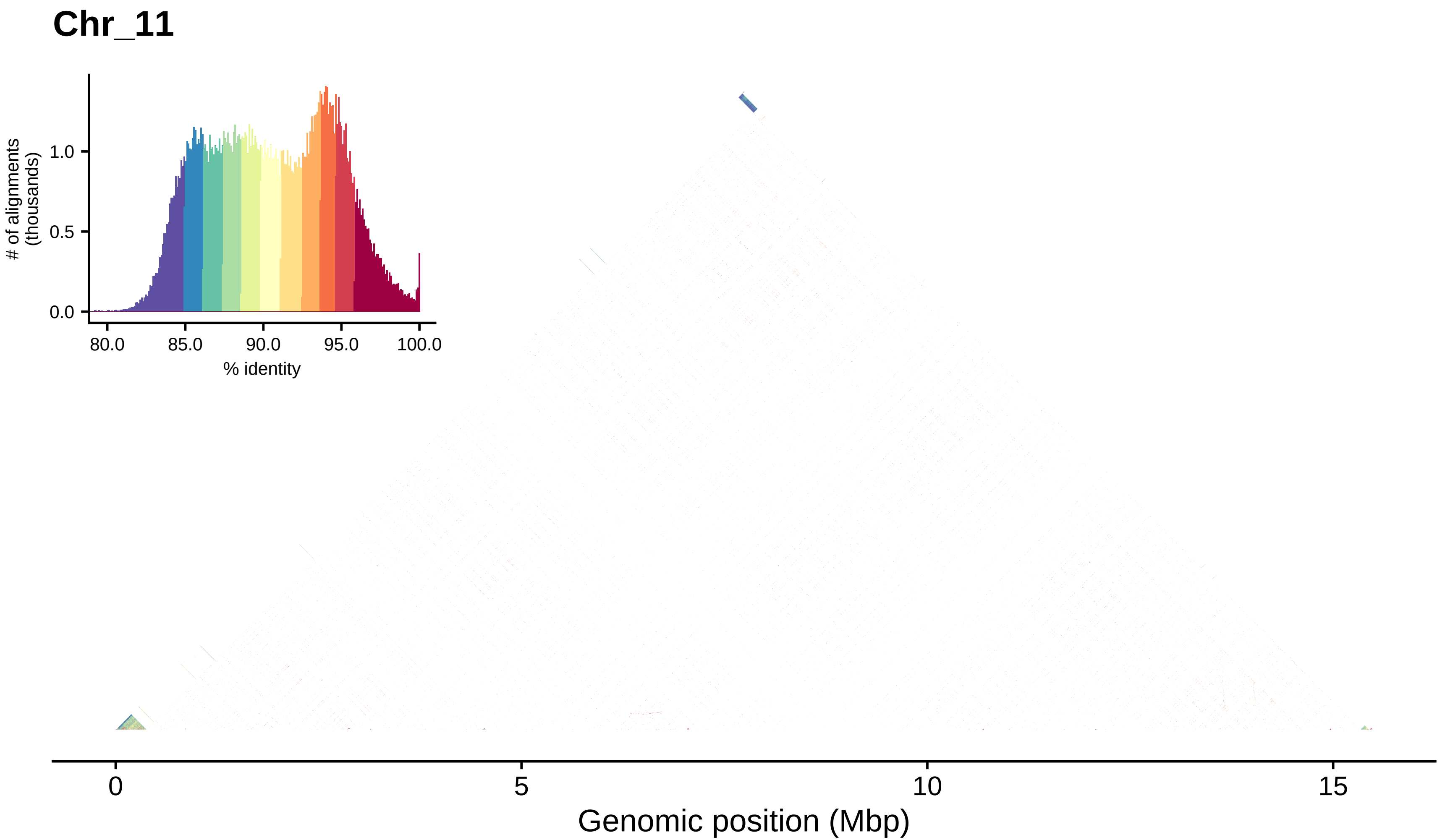
Chr_11
### of alignments (thousands)
1.0
0.5
0.0
80.0
85.0
90.0
95.0
100.0
% identity
0
5
10
15
Genomic position (Mbp)

### Chr_12

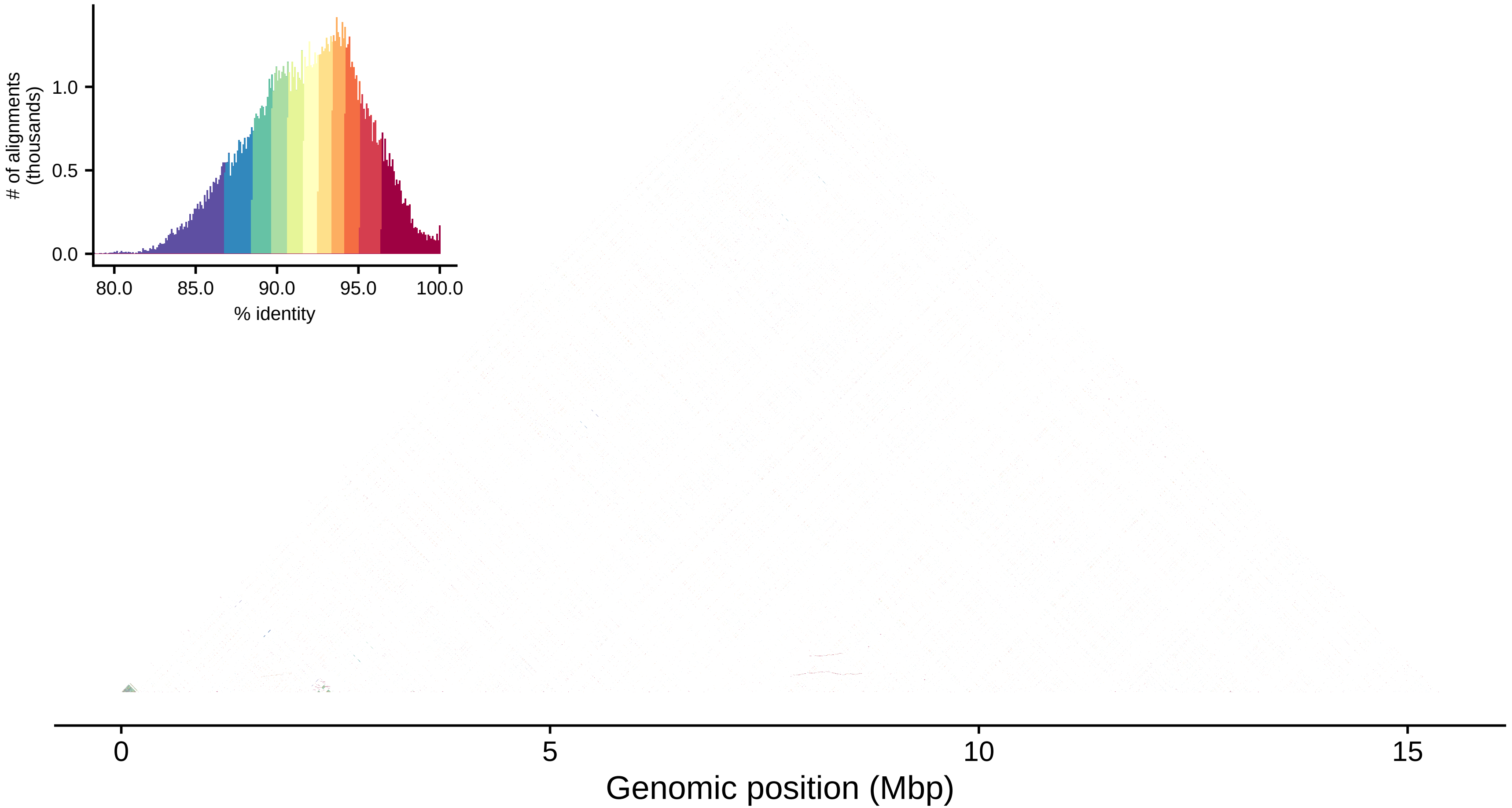

### of alignments (thousands)
0.0
0.5
1.0
80.0
85.0
90.0
95.0
100.0
% identity
0
5
10
15
Genomic position (Mbp)

### Chr_13

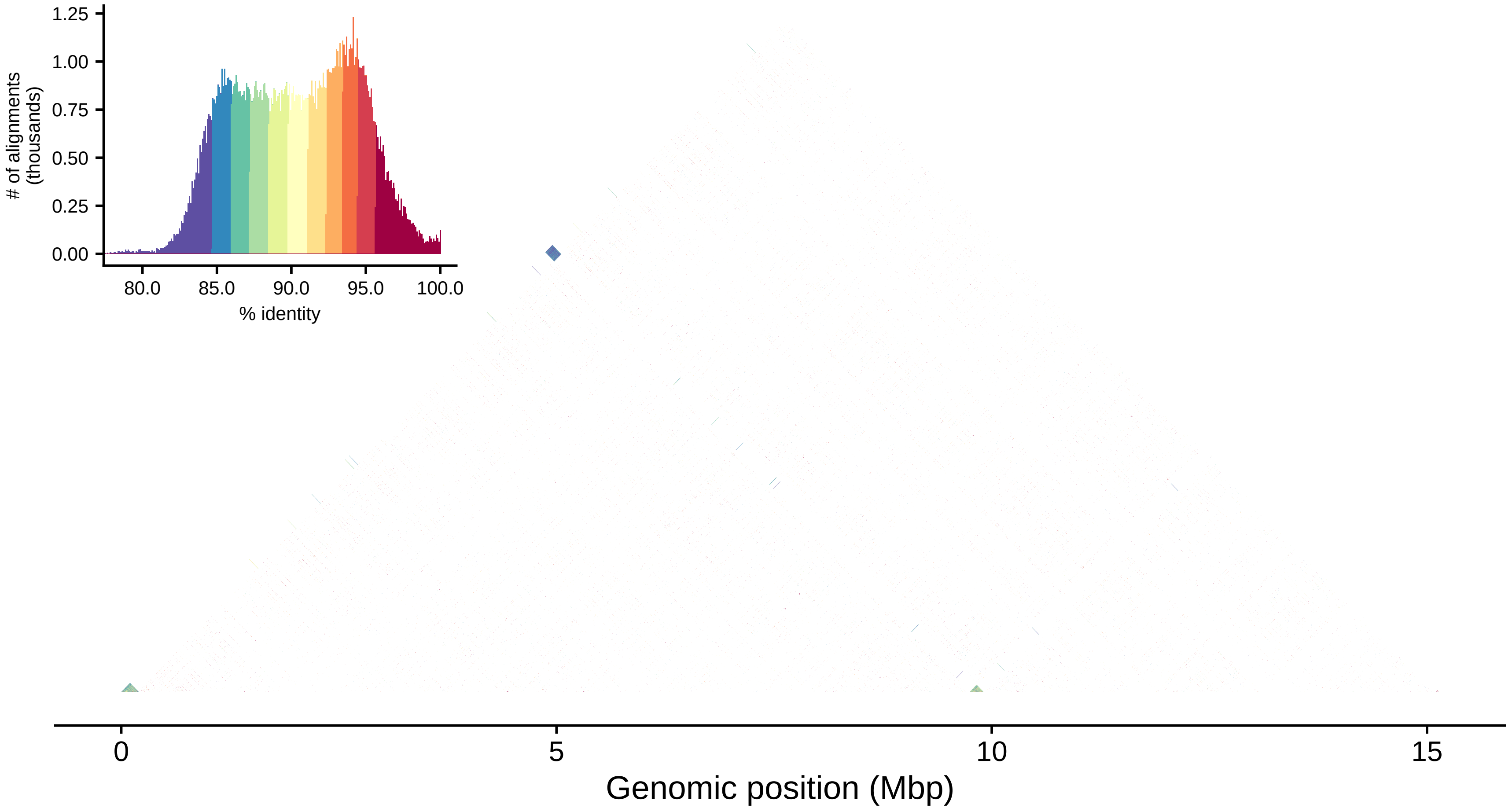

1.25
1.00
0.75
0.50
0.25
0.00
### of alignments (thousands)
80.0
85.0
90.0
95.0
100.0
% identity
0
5
10
15
Genomic position (Mbp)

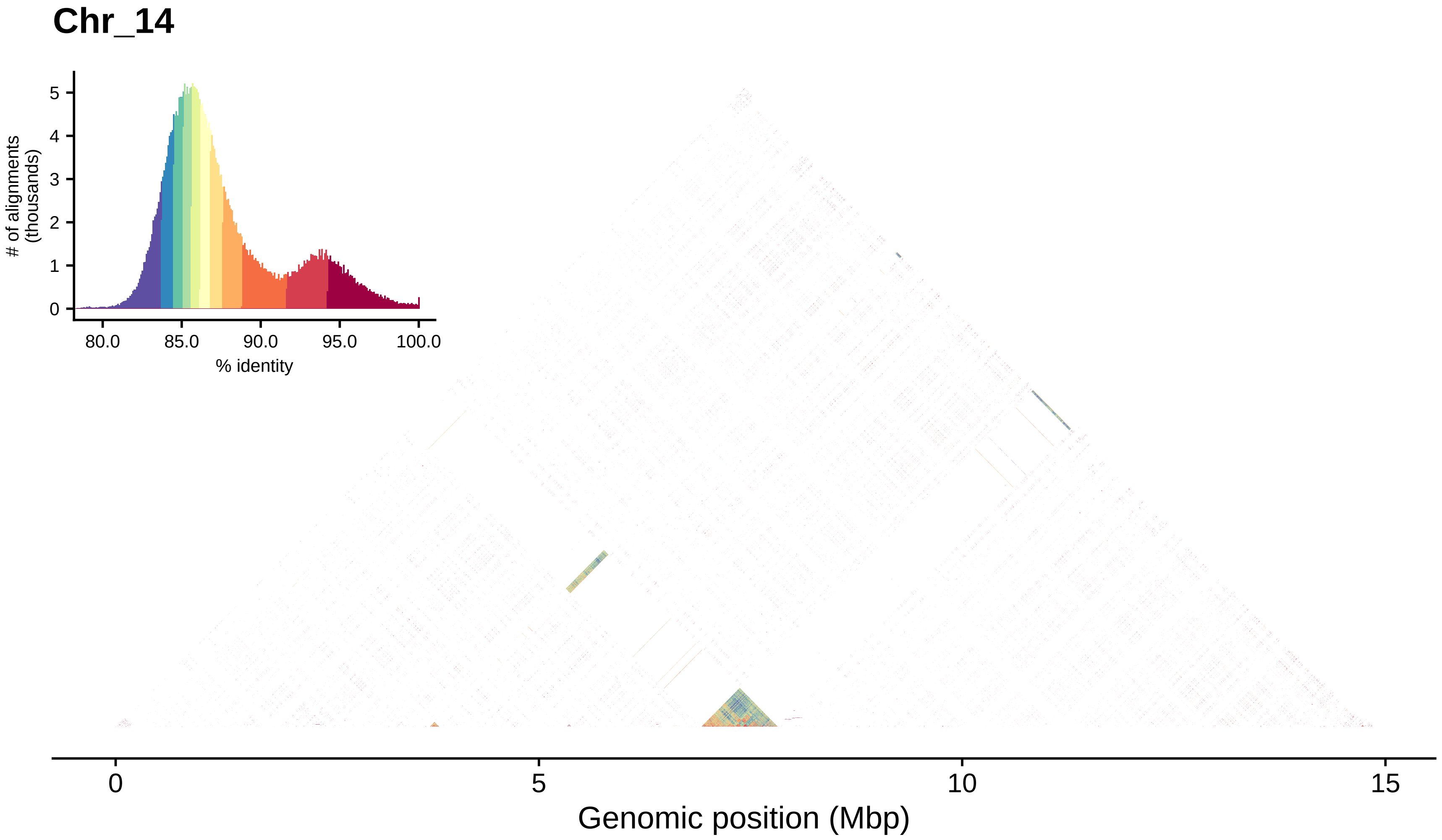

Chr_14
### of alignments
(thousands)
0
1
2
3
4
5
80.0
85.0
90.0
95.0
100.0
% identity
0
5
10
15
Genomic position (Mbp)

#### Chr_15

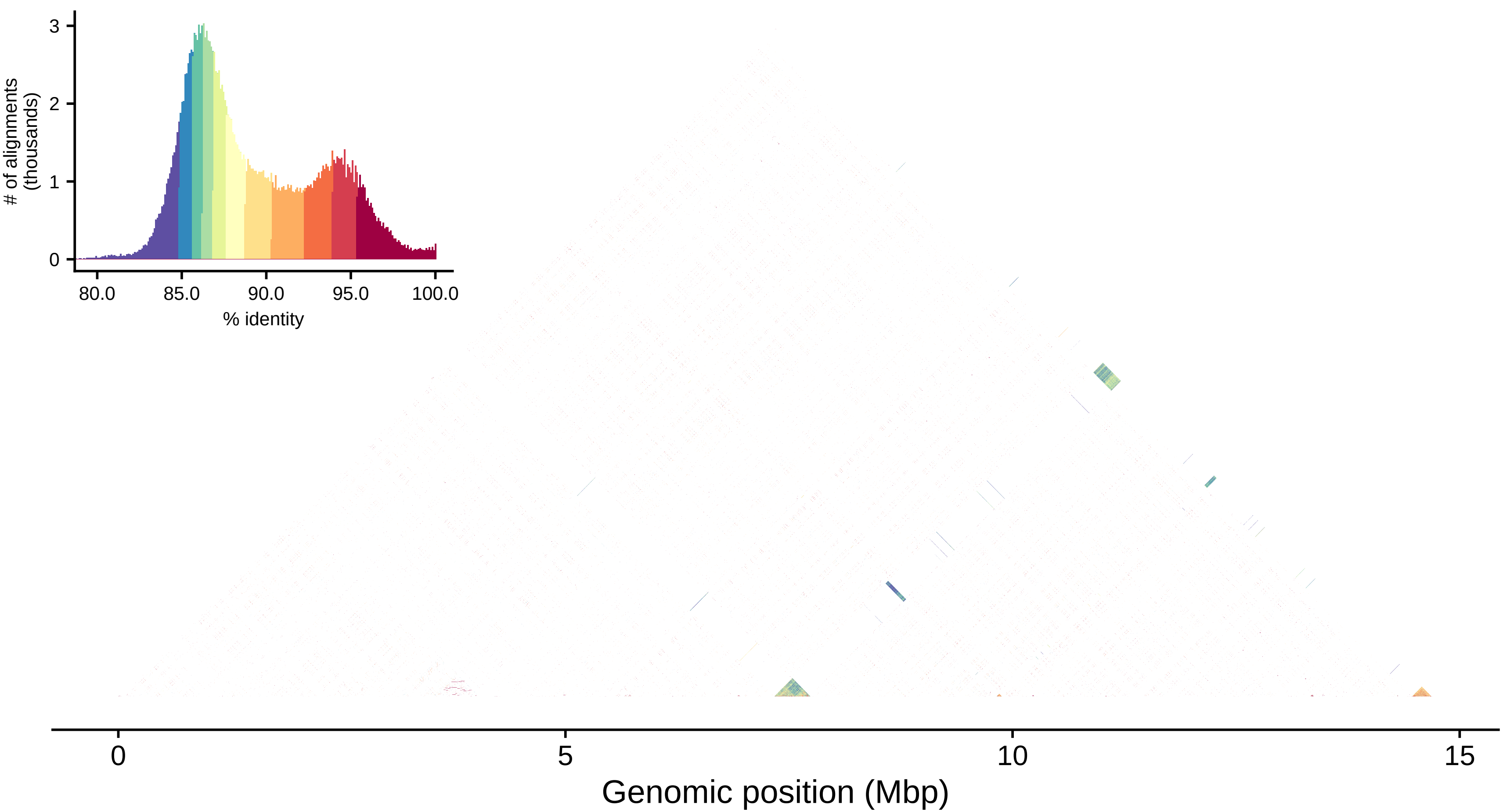

Chr_15
### of alignments (thousands)
0
1
2
3
80.0
85.0
90.0
95.0
100.0
% identity
0
5
10
15
Genomic position (Mbp)

### Chr_16

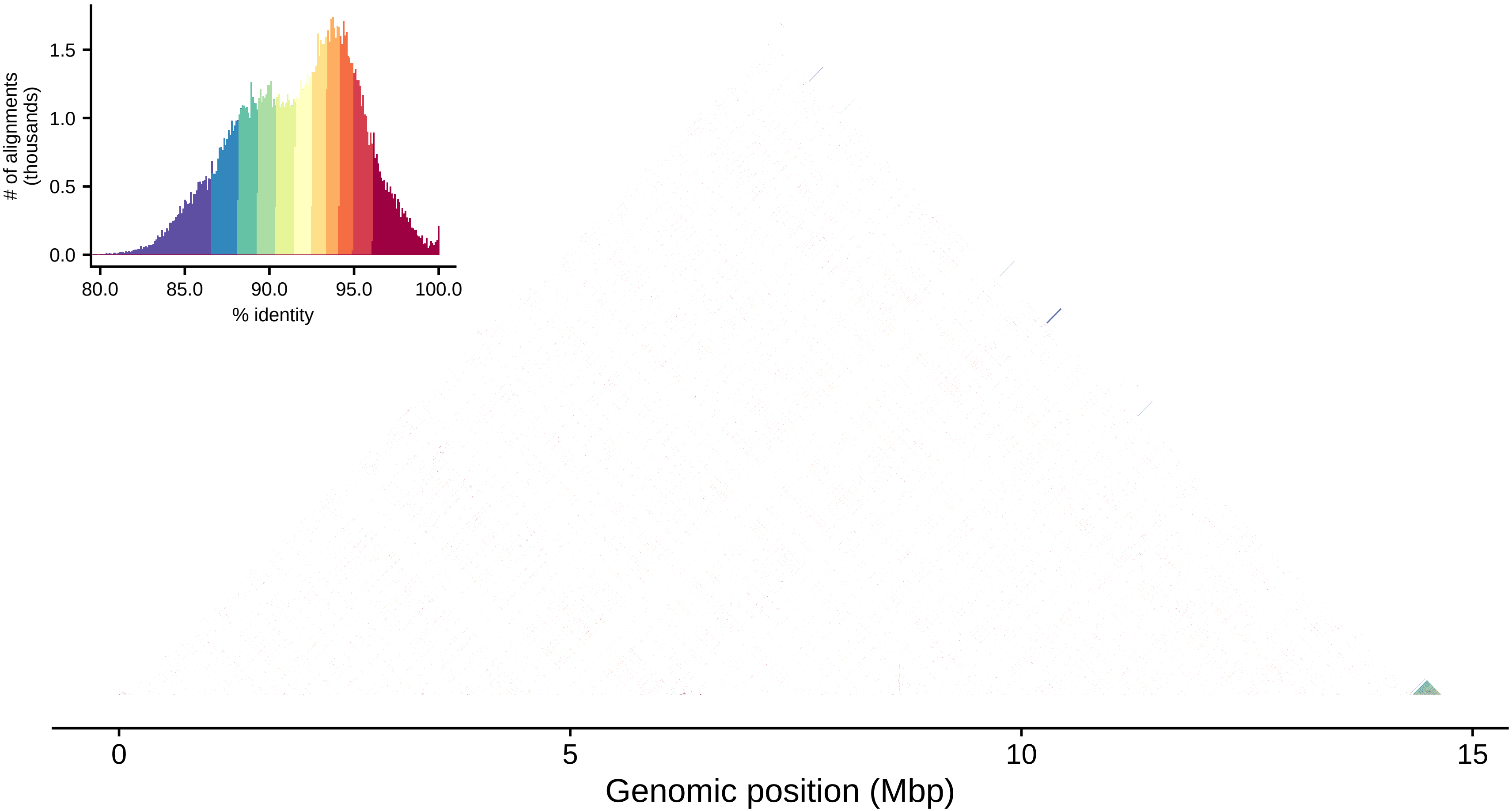

1.5
1.0
0.5
0.0
### of alignments
(thousands)
80.0
85.0
90.0
95.0
100.0
% identity
0
5
10
15
Genomic position (Mbp)

#### Chr_17

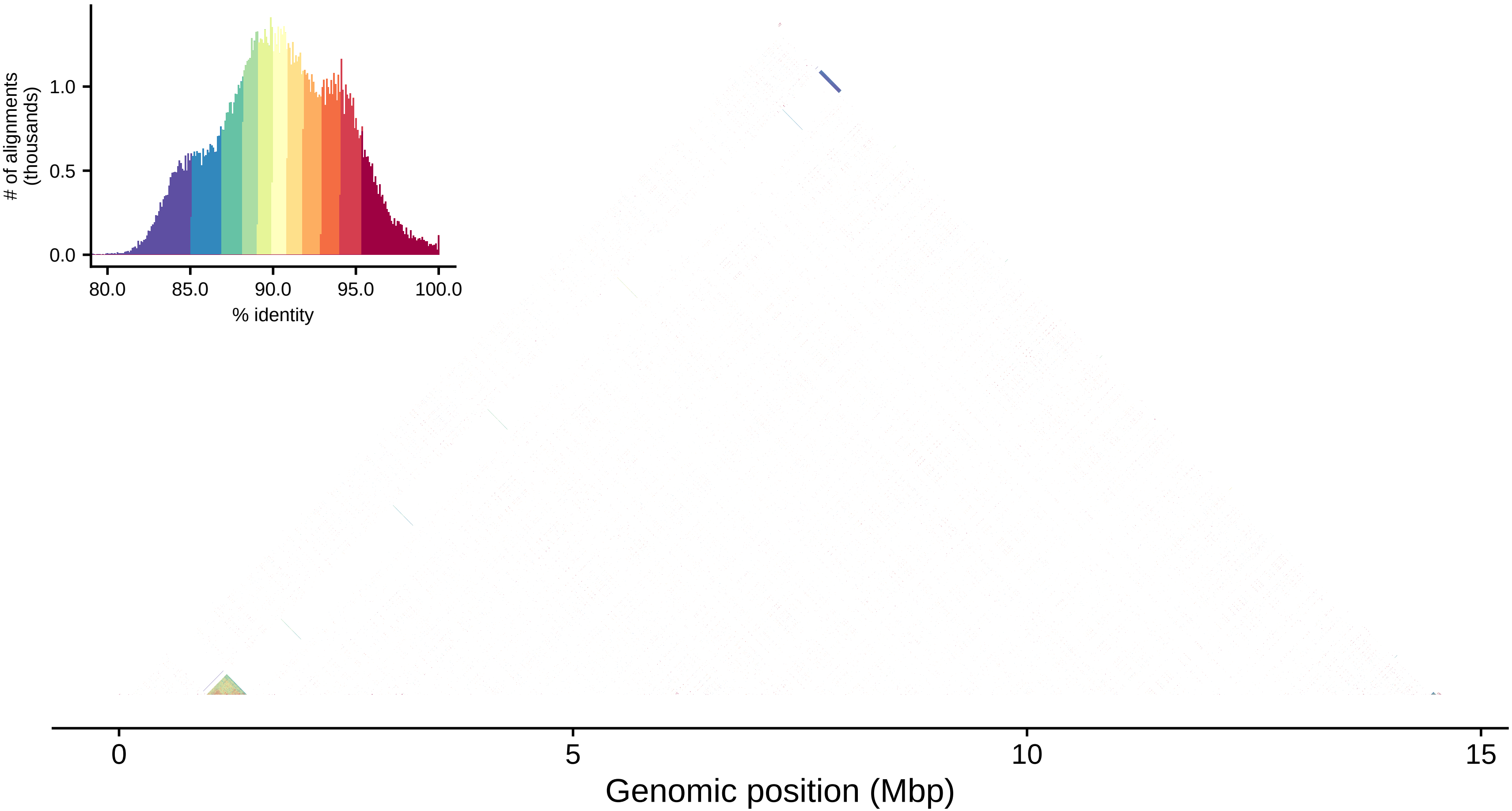

### of alignments (thousands)
1.0
0.5
0.0
80.0
85.0
90.0
95.0
100.0
% identity
0
5
10
15
Genomic position (Mbp)

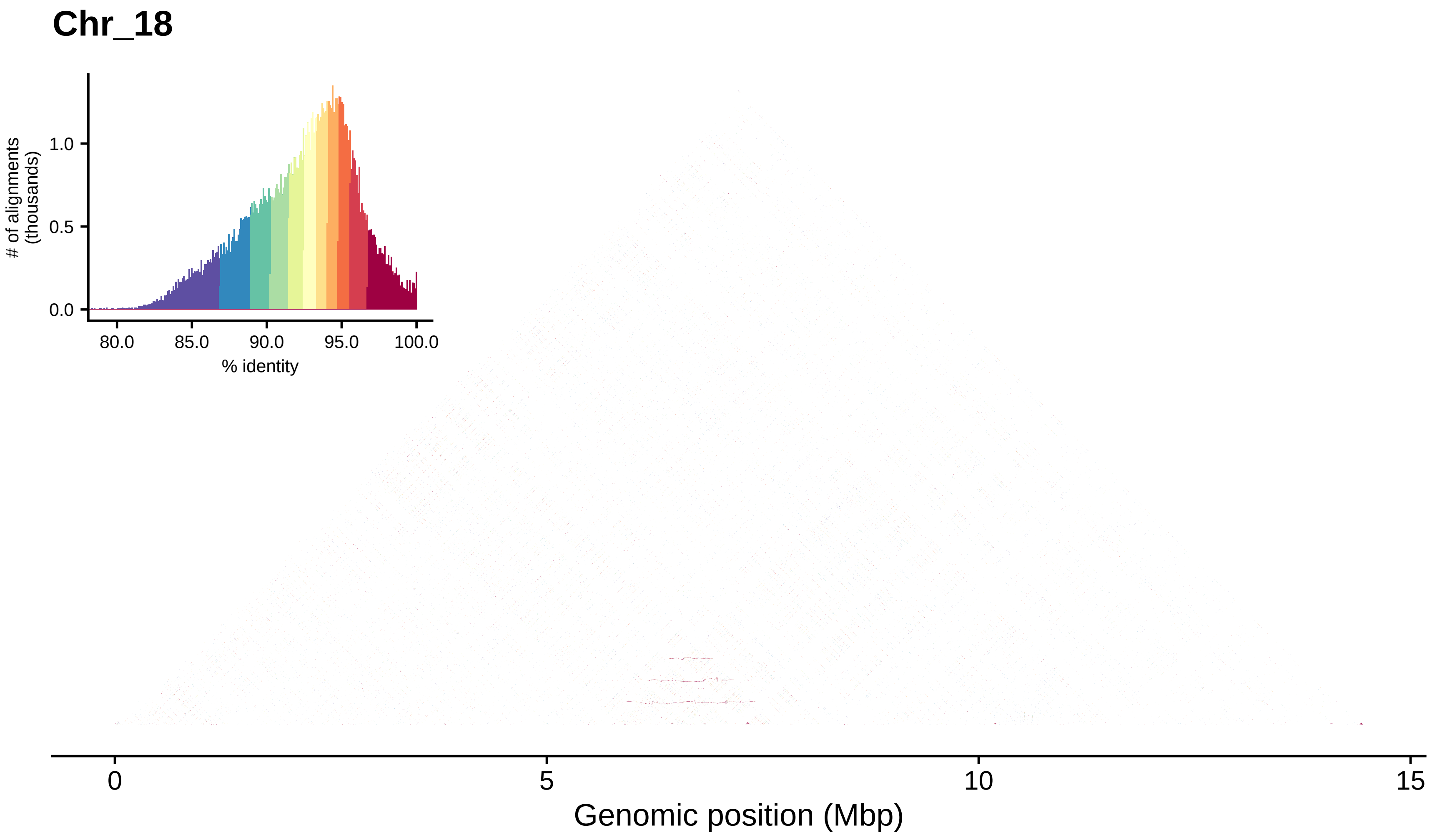
Chr_18
### of alignments
(thousands)
1.0
0.5
0.0
80.0
85.0
90.0
95.0
100.0
% identity
0
5
10
15
Genomic position (Mbp)

#### Chr_19

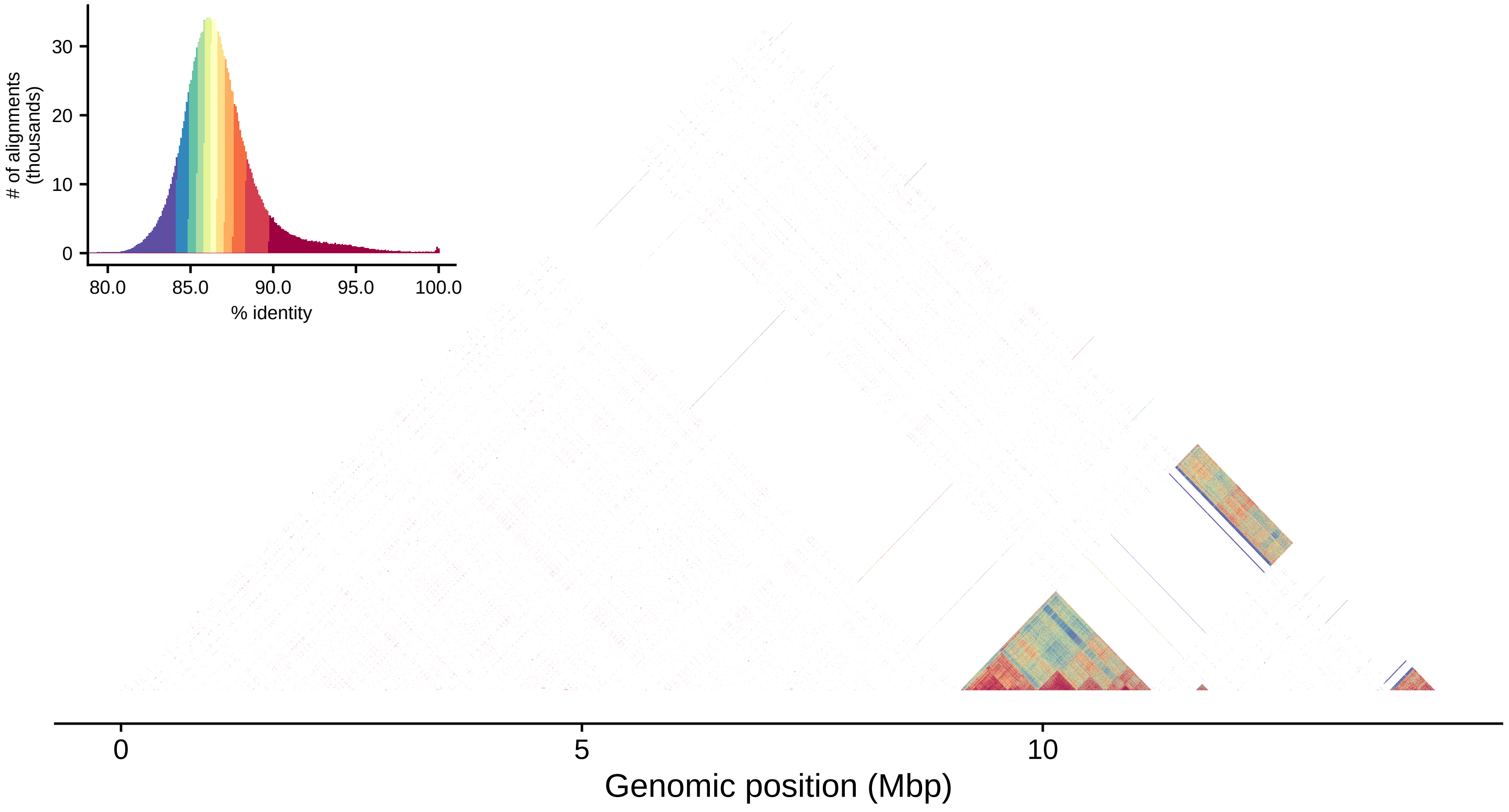

### of alignments
(thousands)
30
20
10
0
80.0
85.0
90.0
95.0
100.0
% identity
0
5
10
Genomic position (Mbp)

### Chr_20

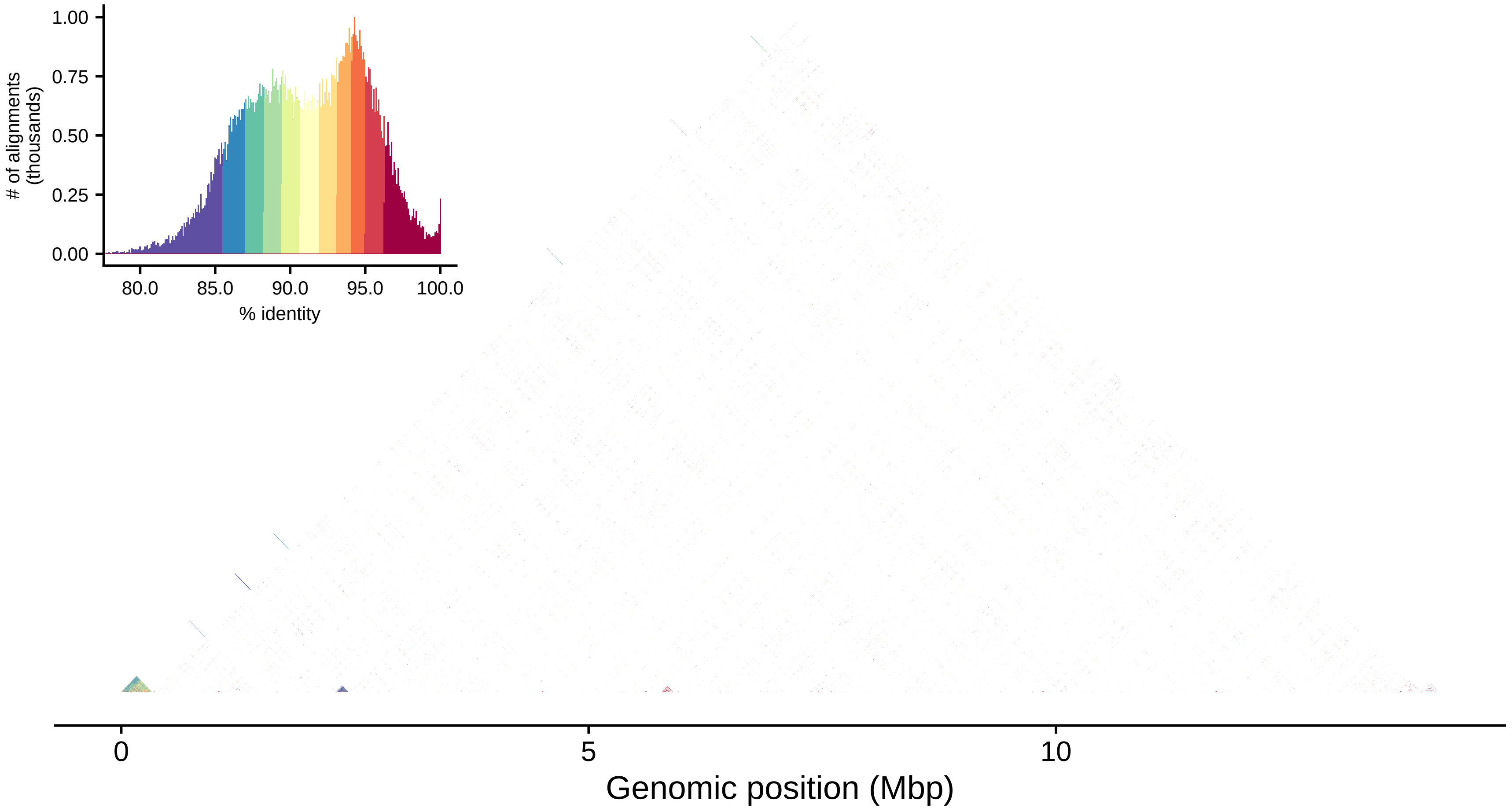

1.00
0.75
0.50
0.25
0.00
### of alignments (thousands)
80.0
85.0
90.0
95.0
100.0
% identity
0
5
10
Genomic position (Mbp)

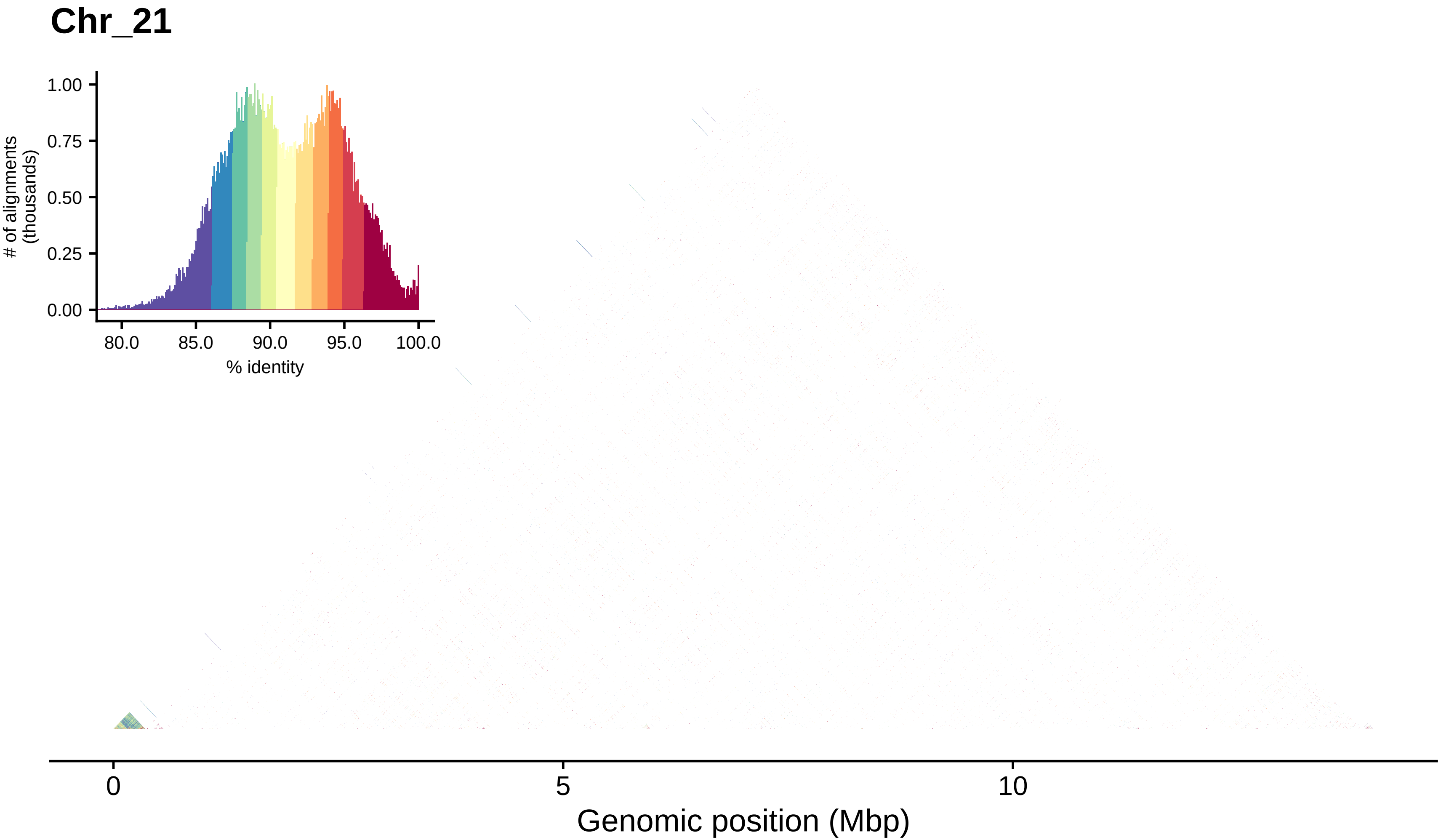

Chr_21
1.00
0.75
0.50
0.25
0.00
### of alignments
(thousands)
80.0
85.0
90.0
95.0
100.0
% identity
0
5
10
Genomic position (Mbp)

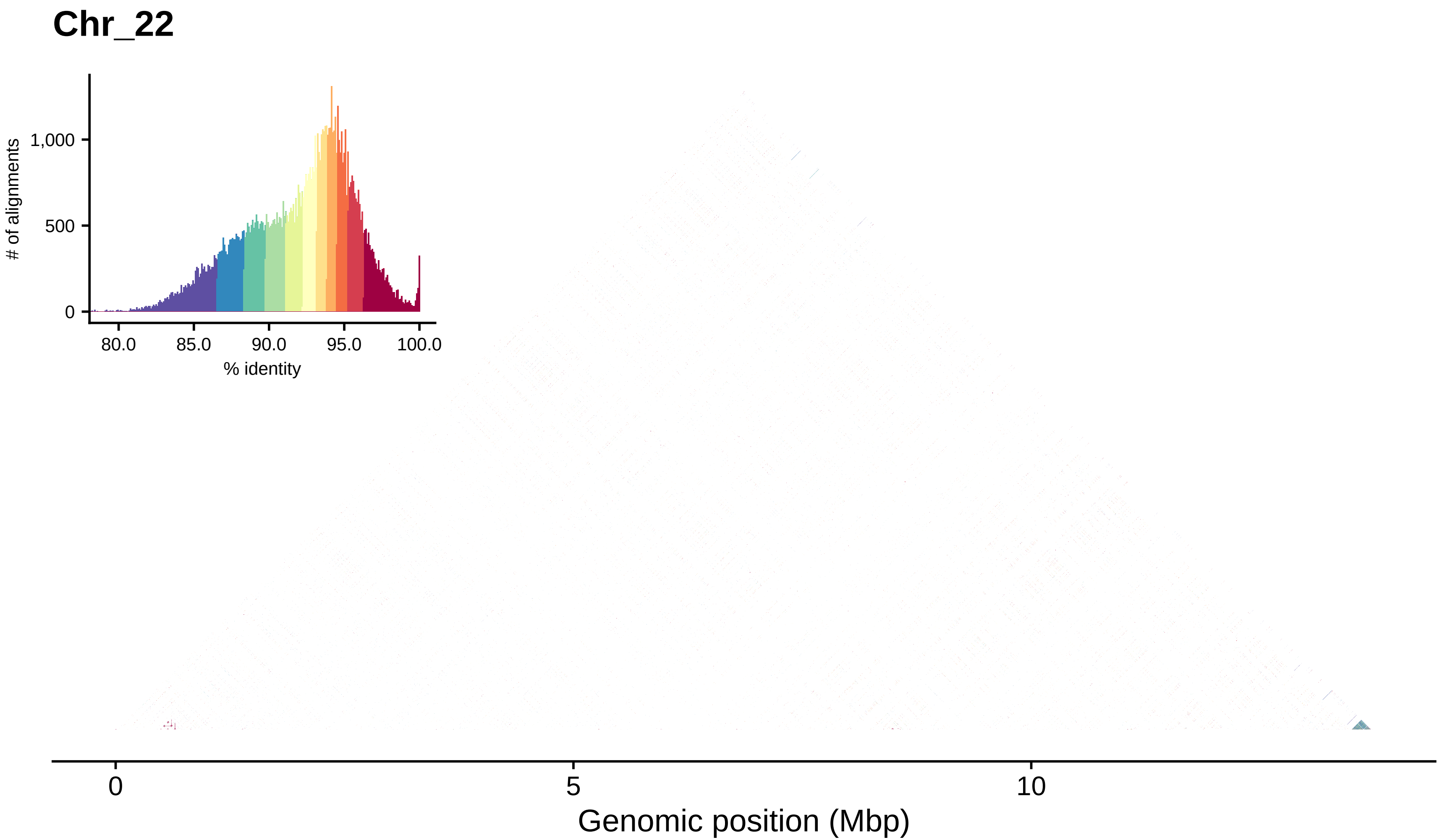
Chr_22
### of alignments
1,000
500
0
80.0
85.0
90.0
95.0
100.0
% identity
0
5
10
Genomic position (Mbp)

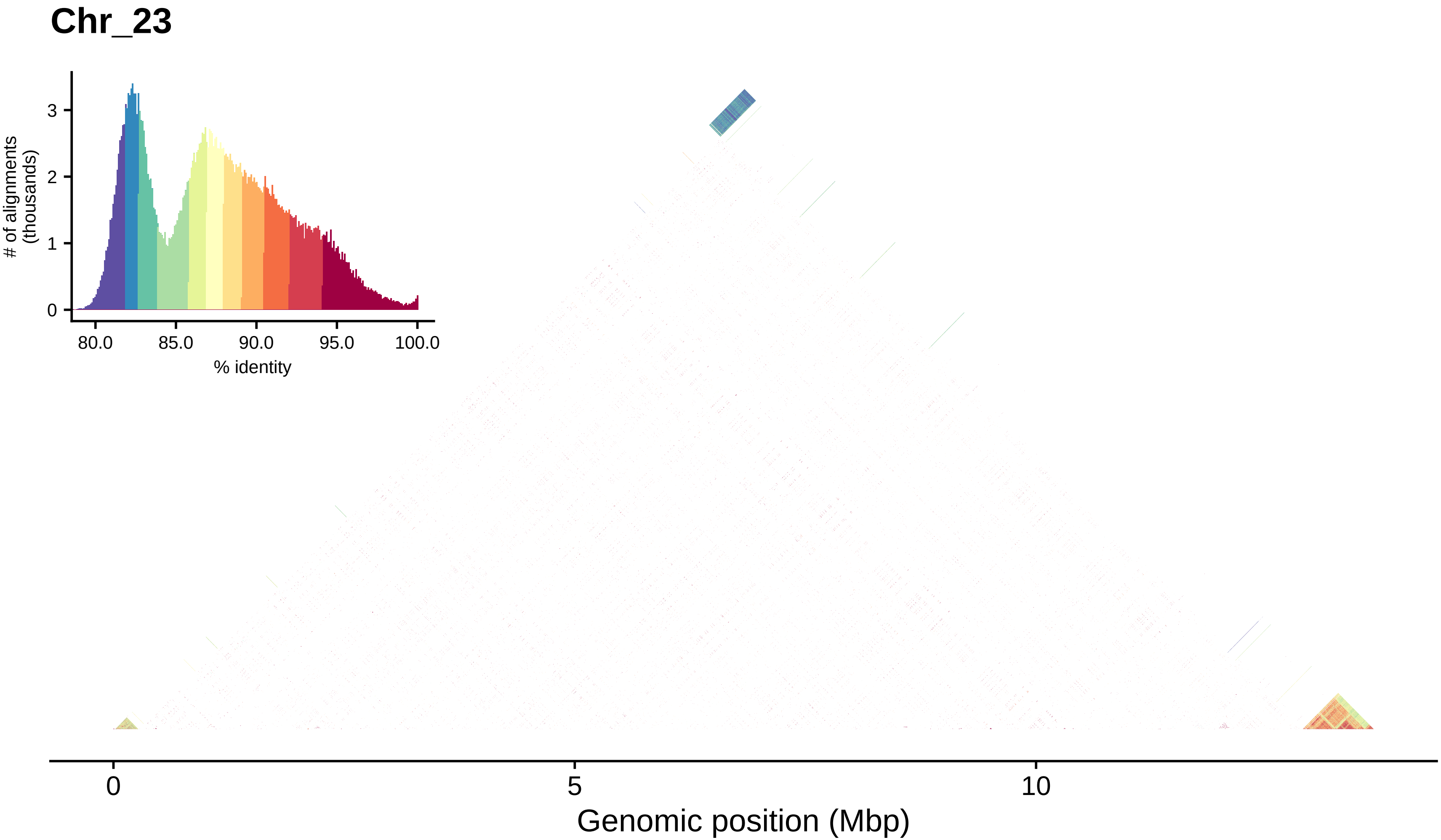
Chr_23
### of alignments (thousands)
0
1
2
3
80.0
85.0
90.0
95.0
100.0
% identity
0
5
10
Genomic position (Mbp)

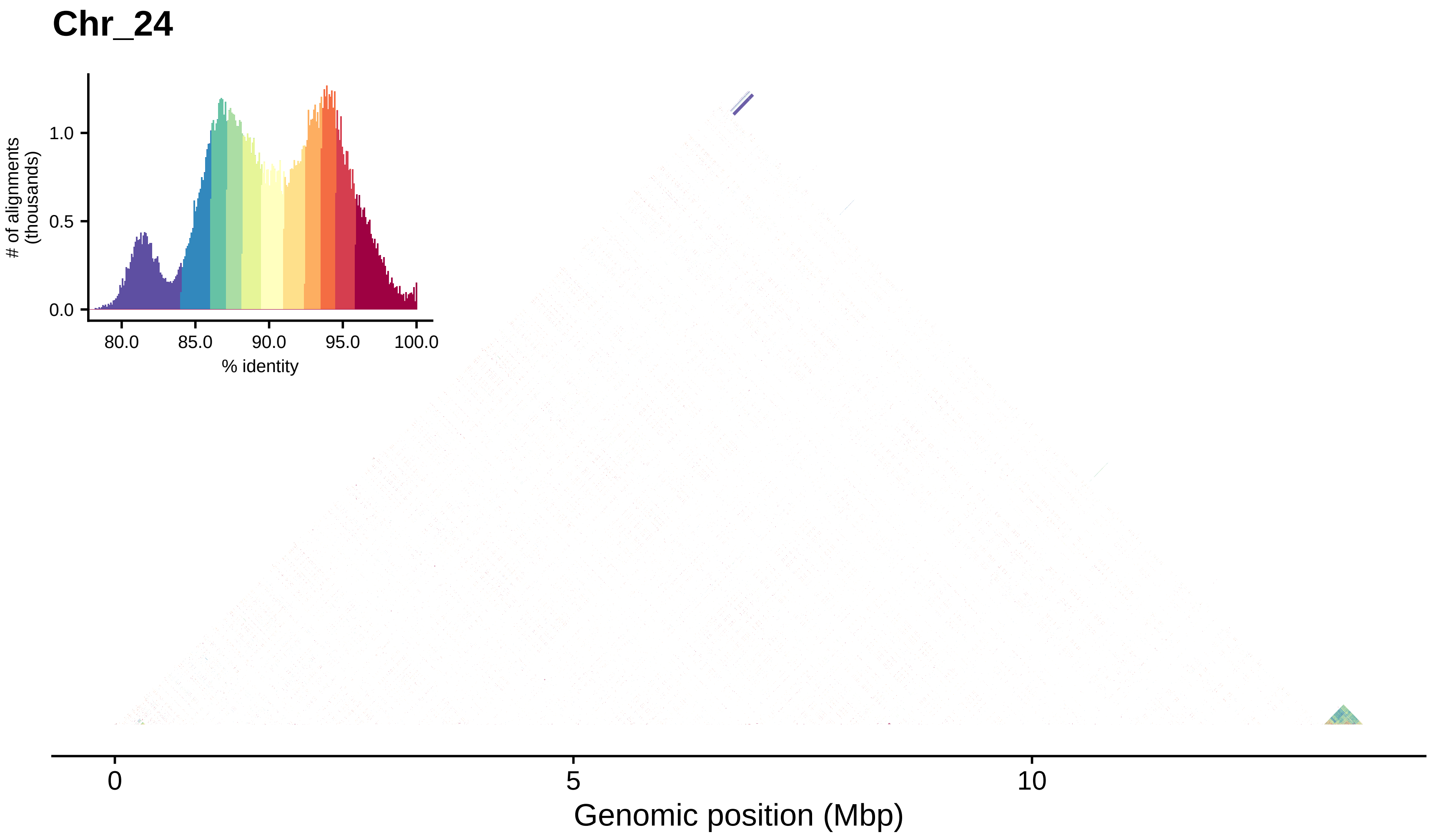
Chr_24
### of alignments (thousands)
1.0
0.5
0.0
80.0
85.0
90.0
95.0
100.0
% identity
0
5
10
Genomic position (Mbp)

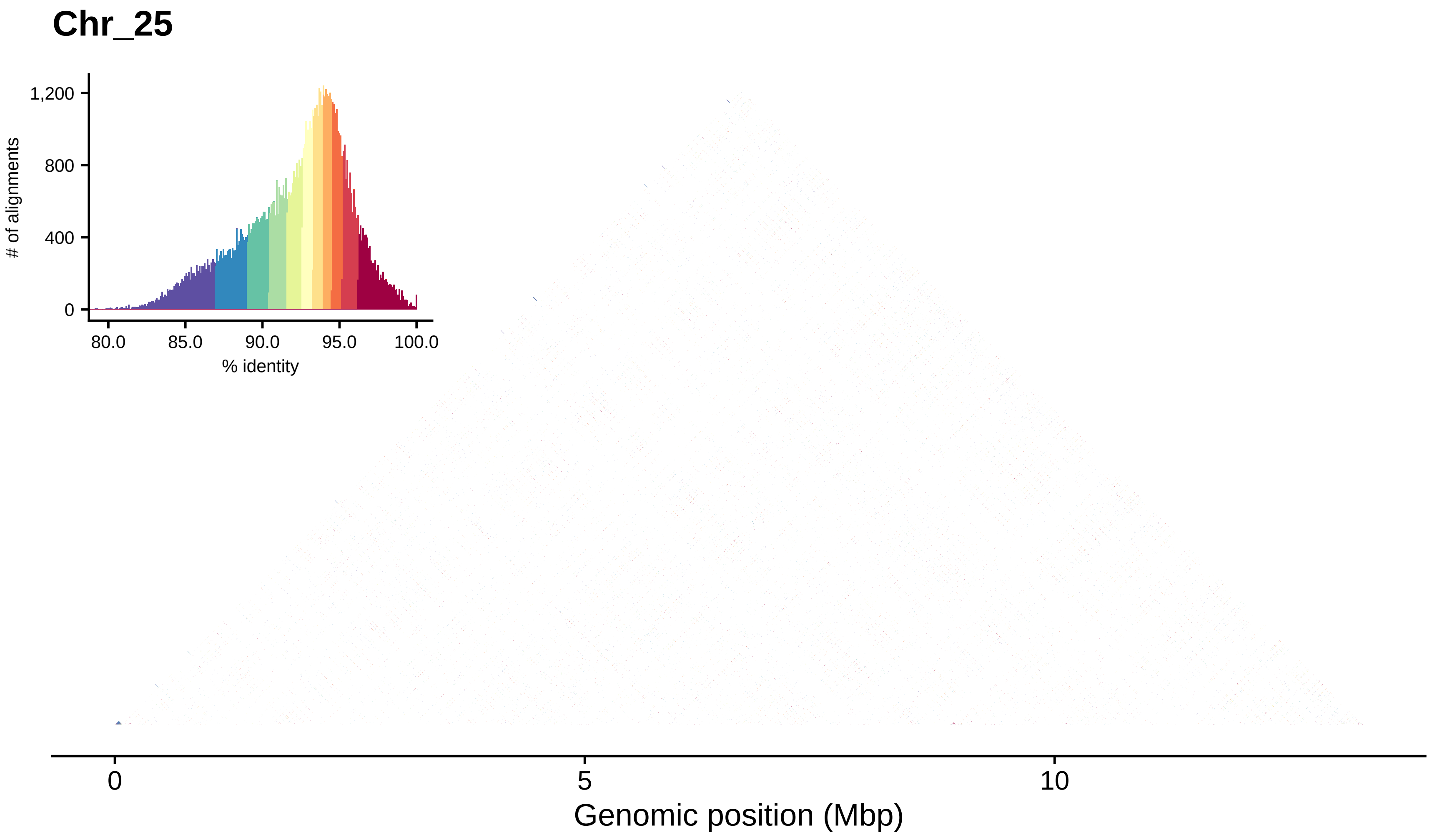
Chr_25
### of alignments
1,200
800
400
0
80.0
85.0
90.0
95.0
100.0
% identity
0
5
10
Genomic position (Mbp)

#### Chr_26

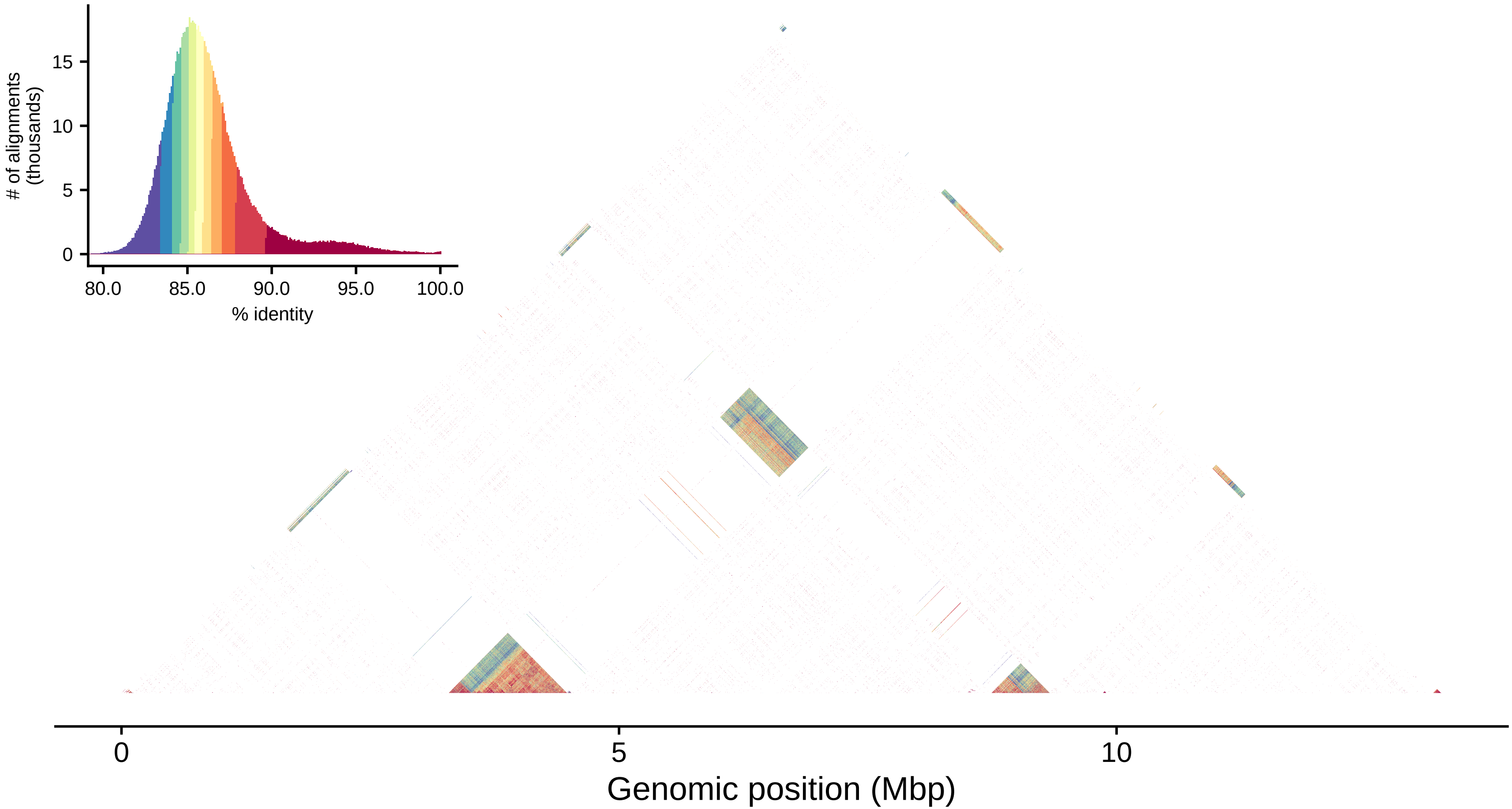

### of alignments (thousands)
15
10
5
0
80.0
85.0
90.0
95.0
100.0
% identity
0
5
10
Genomic position (Mbp)

#### Chr_27

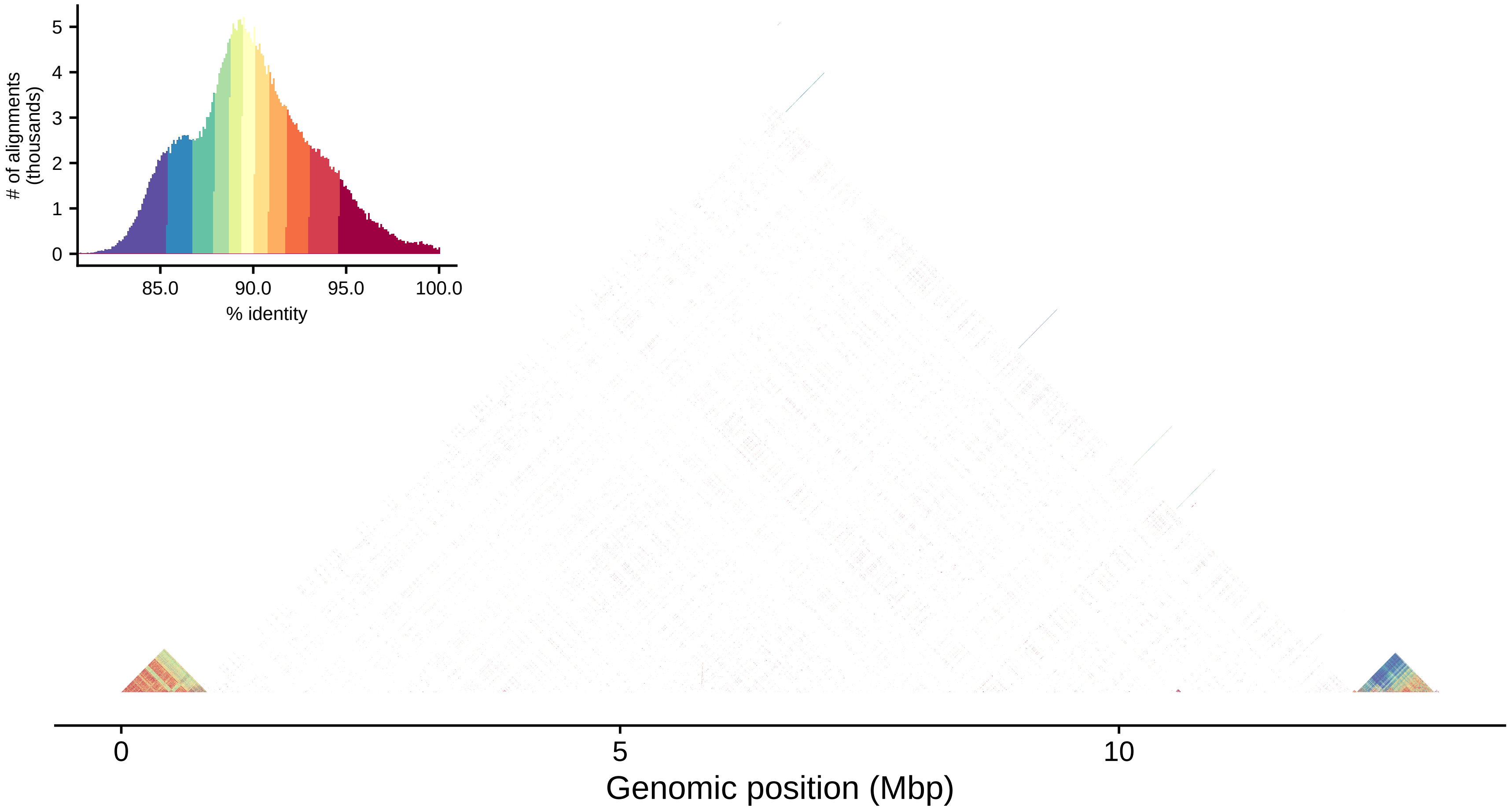

### of alignments (thousands)
5
4
3
2
1
0
85.0
90.0
95.0
100.0
% identity
0
5
10
Genomic position (Mbp)

#### Chr_28

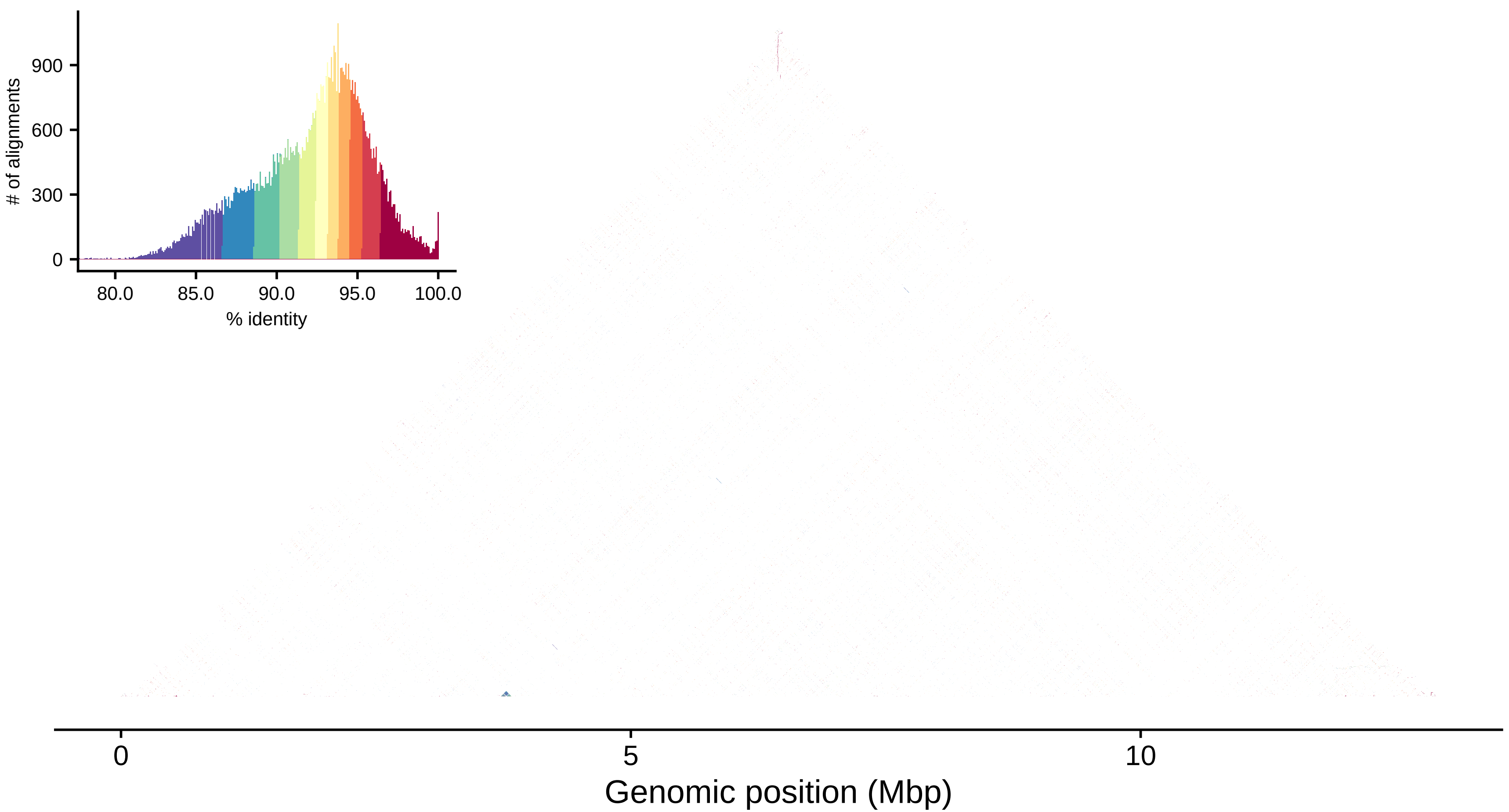

900
600
300
0
### of alignments
80.0
85.0
90.0
95.0
100.0
% identity
0
5
10
Genomic position (Mbp)

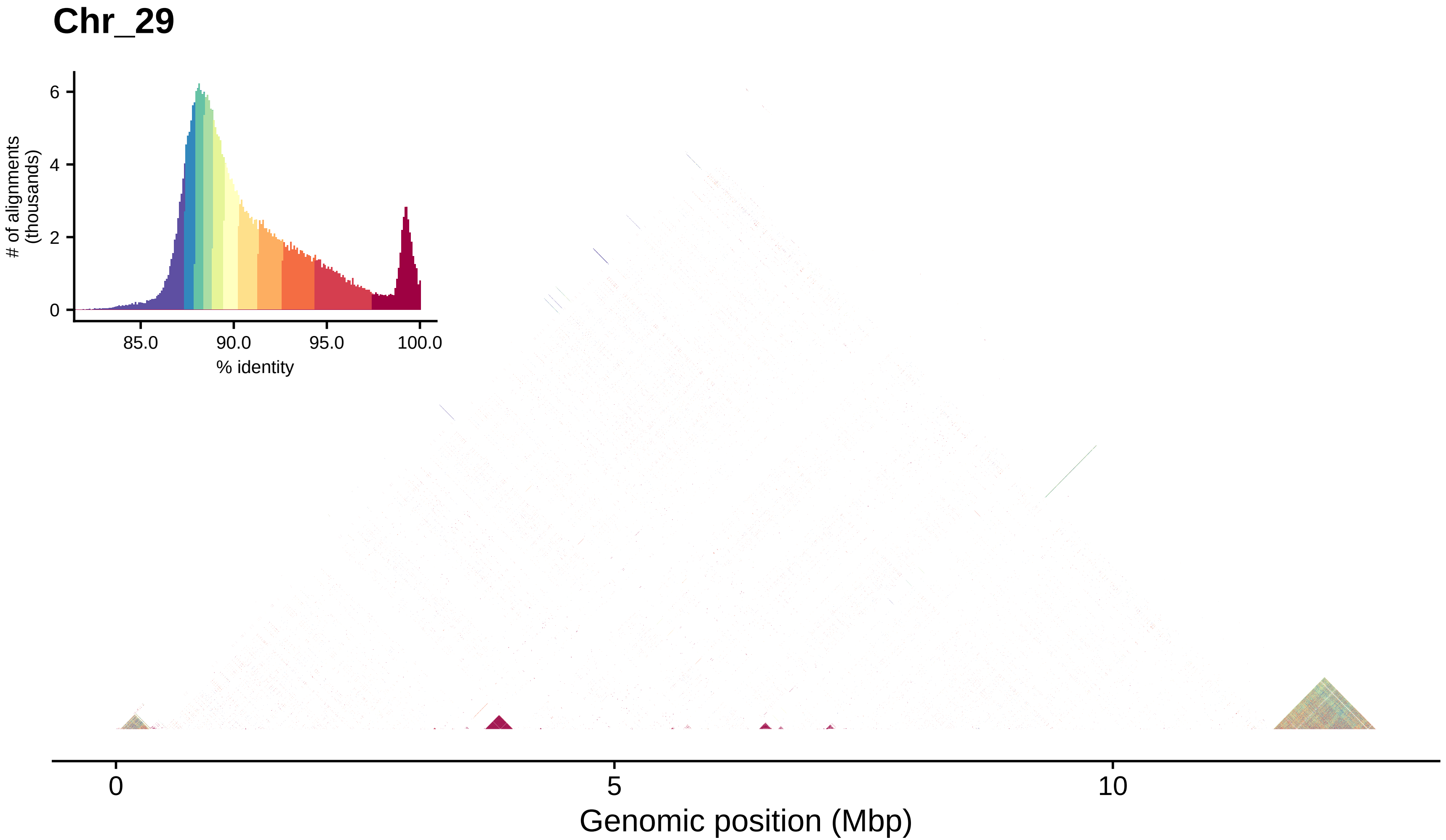
Chr_29
### of alignments
(thousands)
6
4
2
0
85.0
90.0
95.0
100.0
% identity
0
5
10
Genomic position (Mbp)

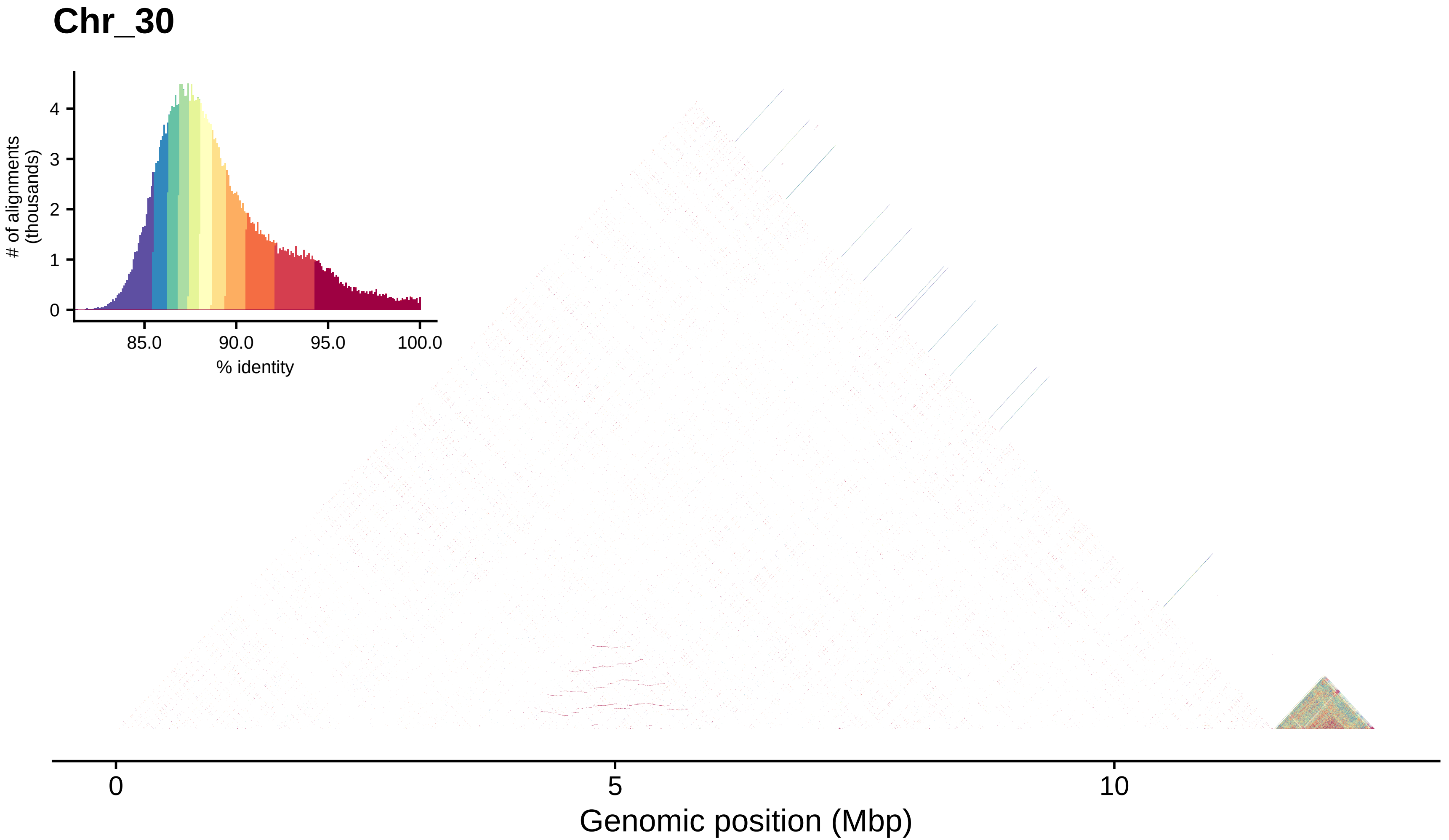

Chr_30
### of alignments (thousands)
0
1
2
3
4
85.0
90.0
95.0
100.0
% identity
0
5
10
Genomic position (Mbp)

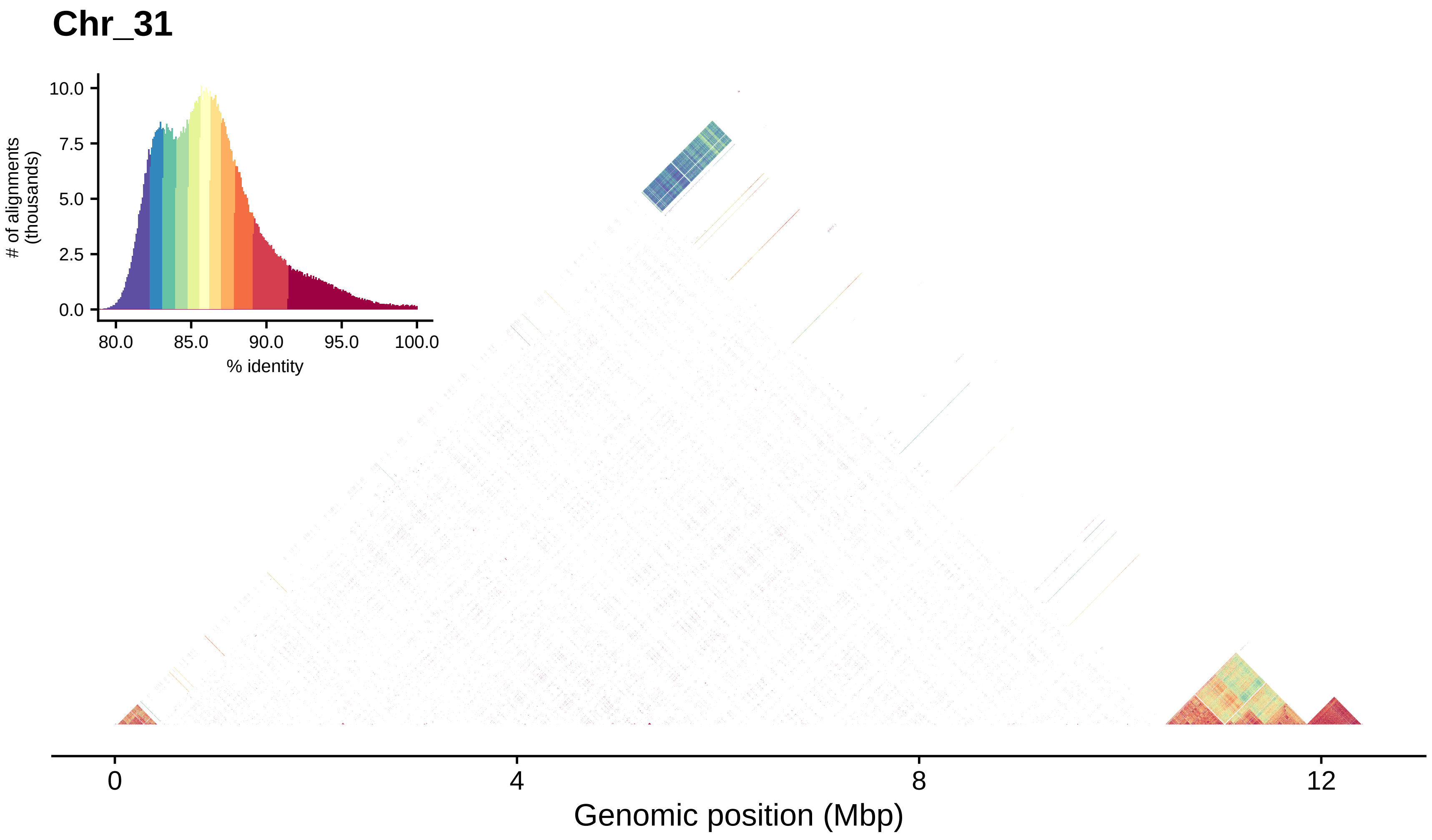
Chr_31
### of alignments
(thousands)
10.0
7.5
5.0
2.5
0.0
80.0
85.0
90.0
95.0
100.0
% identity
0
4
8
12
Genomic position (Mbp)

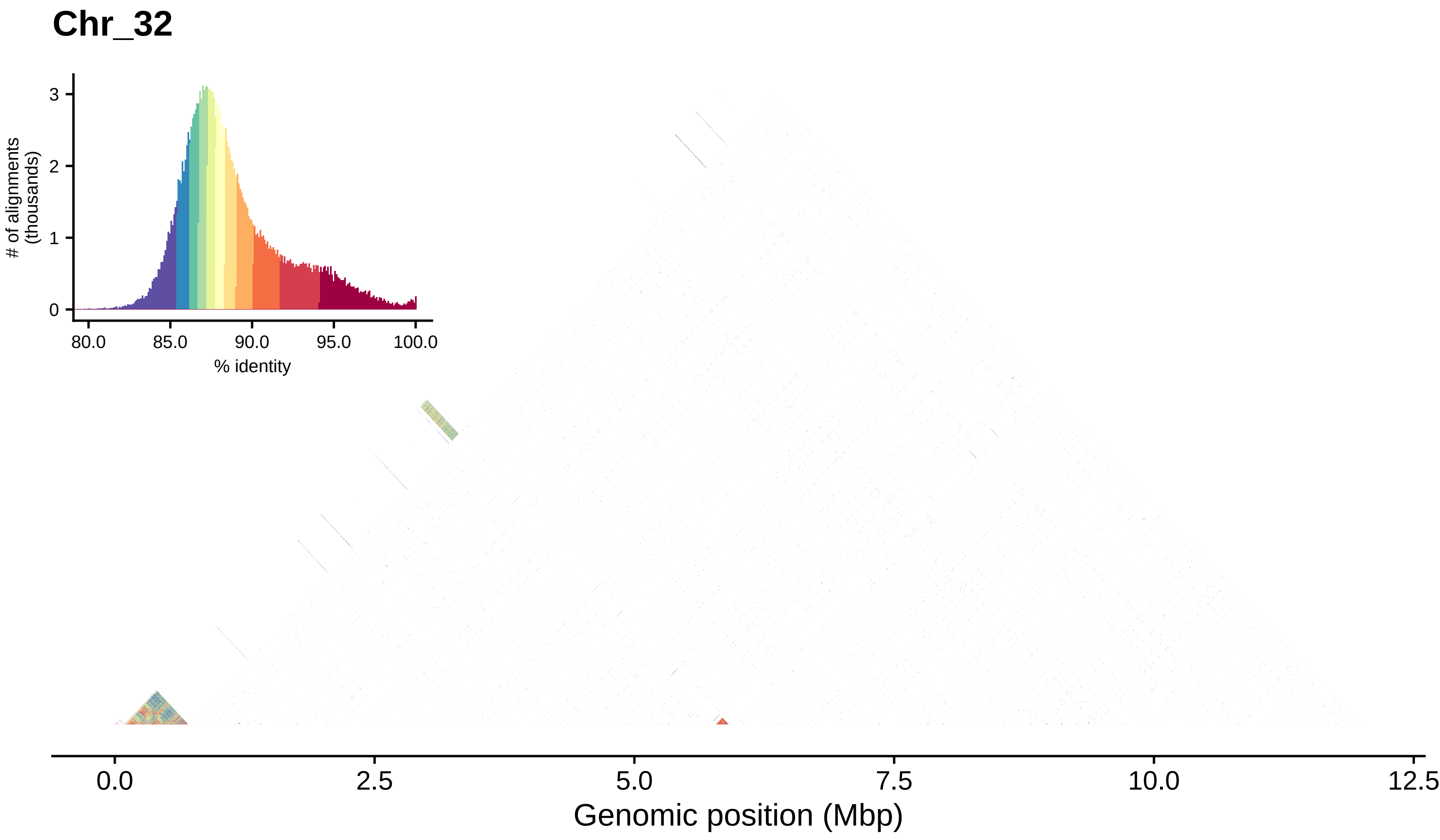

Chr_32
### of alignments (thousands)
0
1
2
3
80.0
85.0
90.0
95.0
100.0
% identity
0.0
2.5
5.0
7.5
10.0
12.5
Genomic position (Mbp)

#### Chr_33

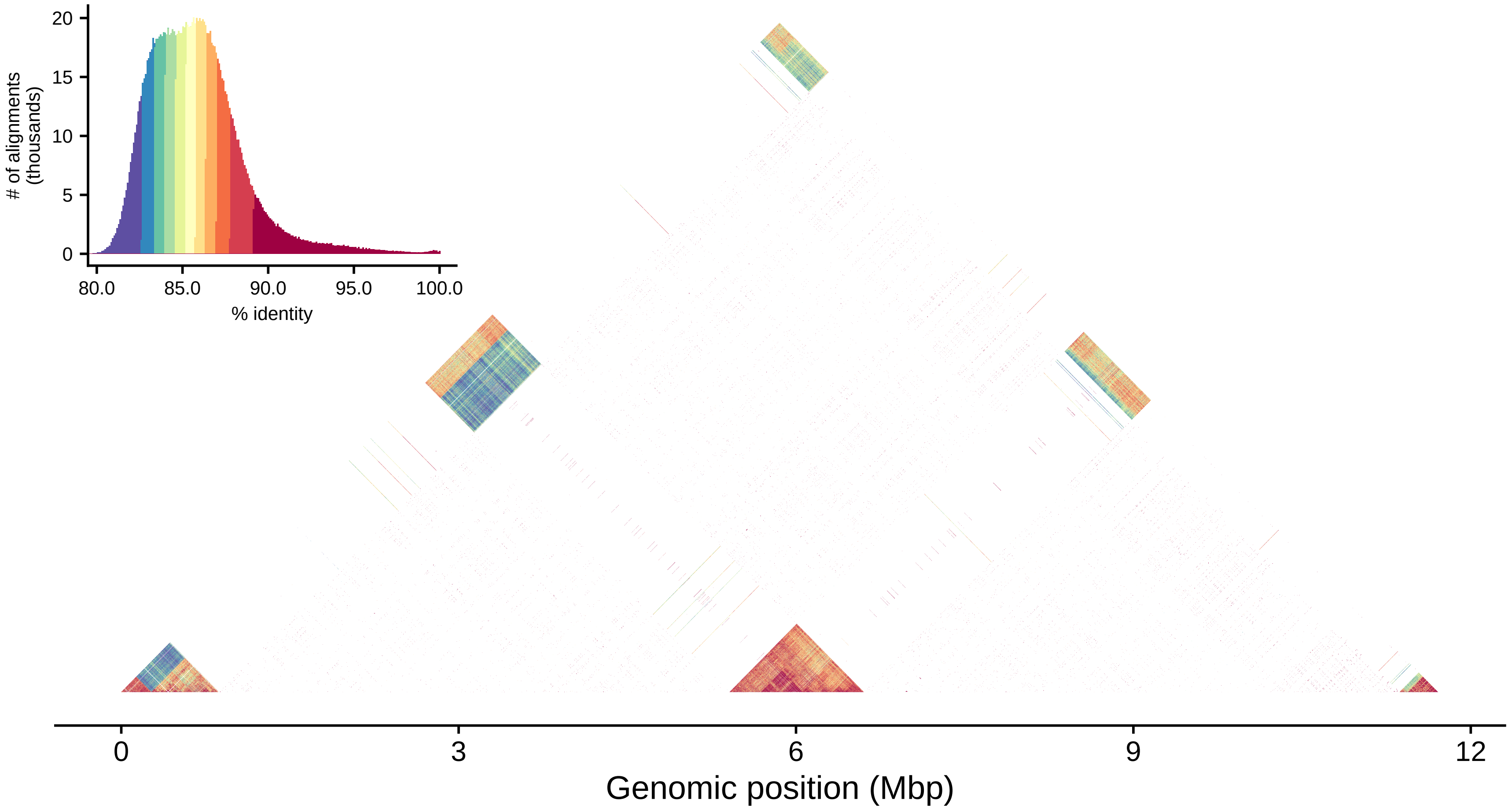

### of alignments (thousands)
20
15
10
5
0
80.0
85.0
90.0
95.0
100.0
% identity
0
3
6
9
12
Genomic position (Mbp)

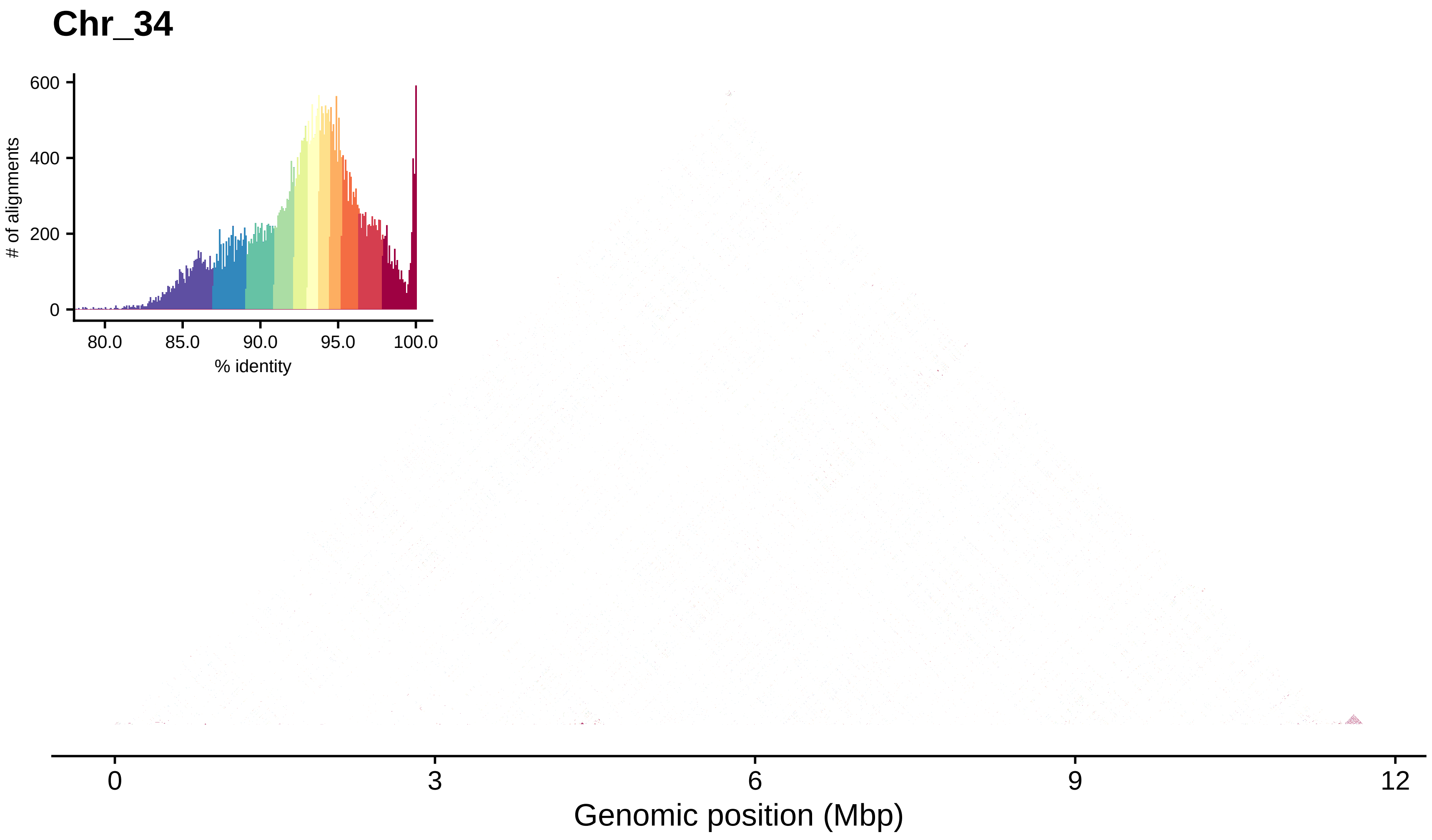

Chr_34
600
400
200
0
### of alignments
80.0
85.0
90.0
95.0
100.0
% identity
0
3
6
9
12
Genomic position (Mbp)

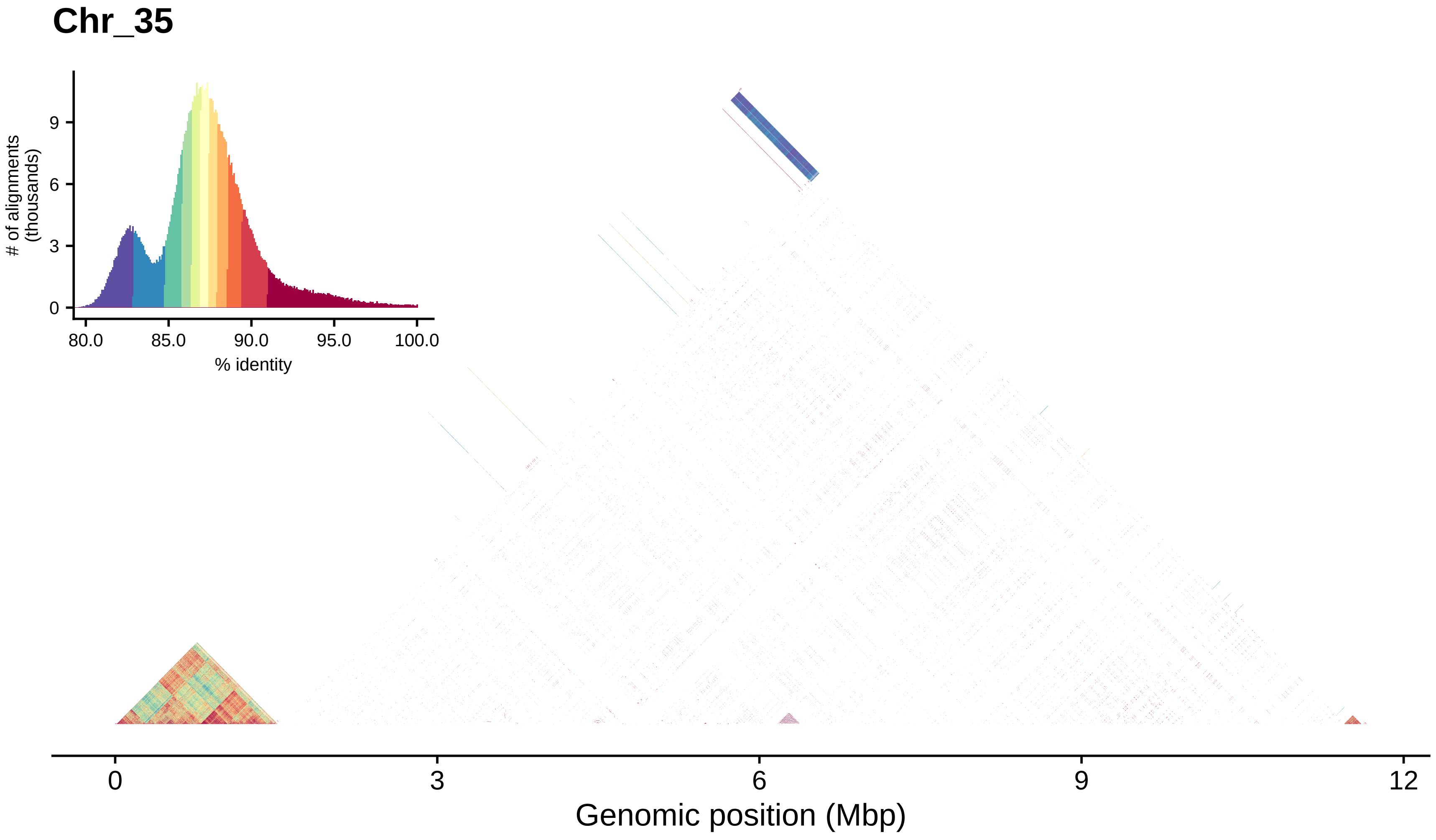
Chr_35
### of alignments
(thousands)
0
3
6
9
80.0
85.0
90.0
95.0
100.0
% identity
0
3
6
9
12
Genomic position (Mbp)

#### Chr_36

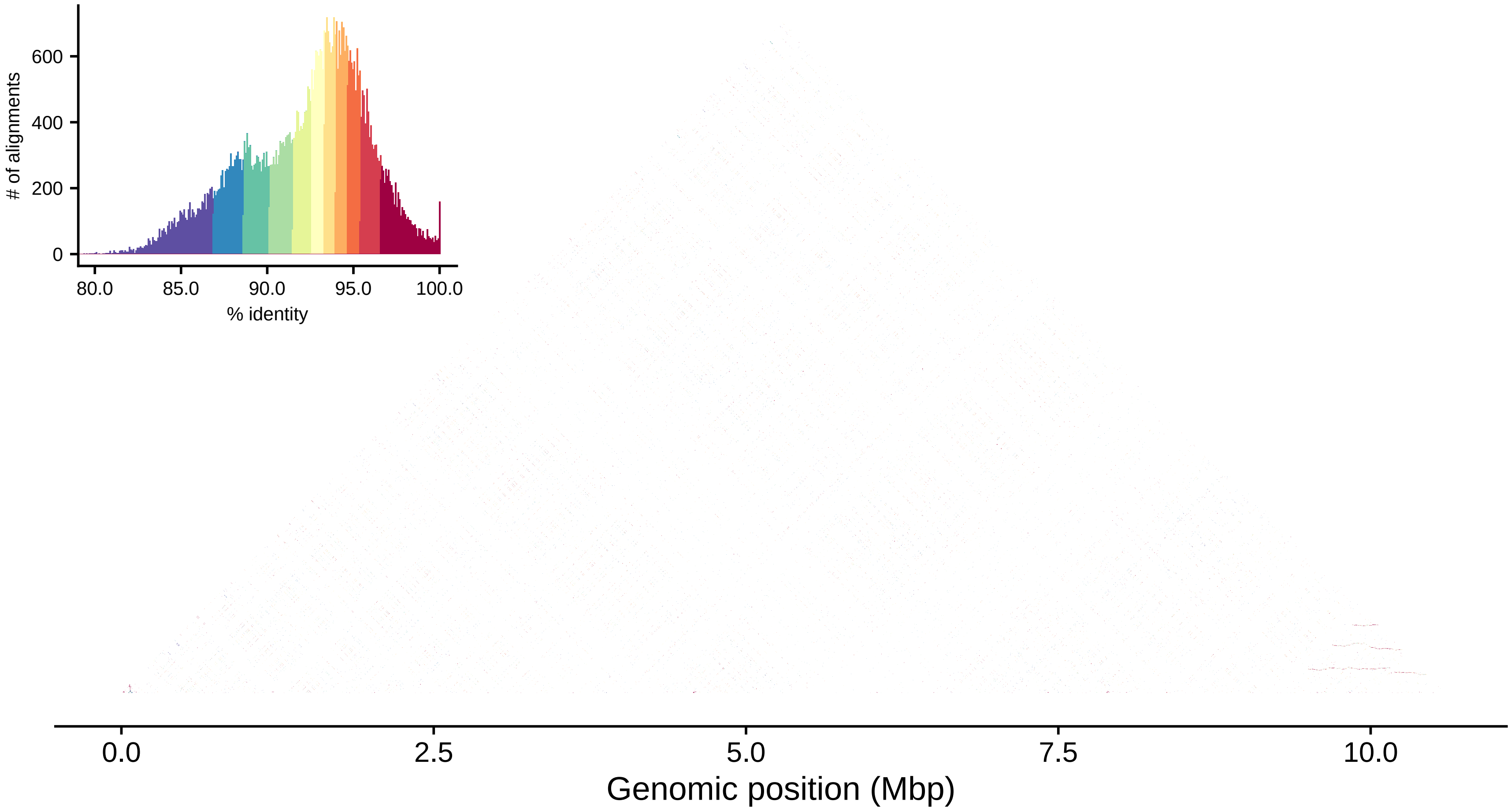

### of alignments
600
400
200
0
80.0
85.0
90.0
95.0
100.0
% identity
0.0
2.5
5.0
7.5
10.0
Genomic position (Mbp)

#### Chr_37

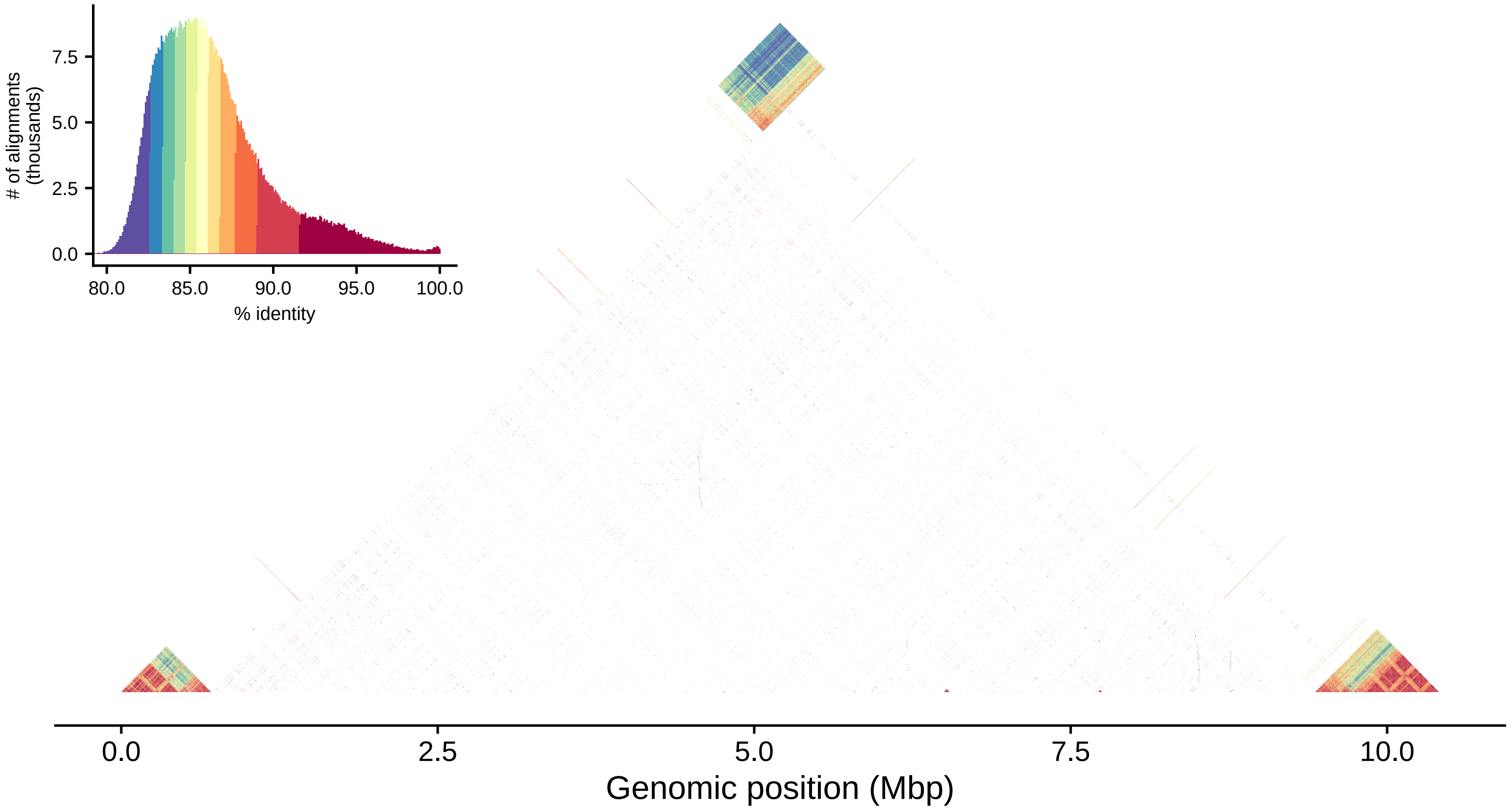

### of alignments (thousands)
7.5
5.0
2.5
0.0
80.0
85.0
90.0
95.0
100.0
% identity
0.0
2.5
5.0
7.5
10.0
Genomic position (Mbp)

### Chr_38

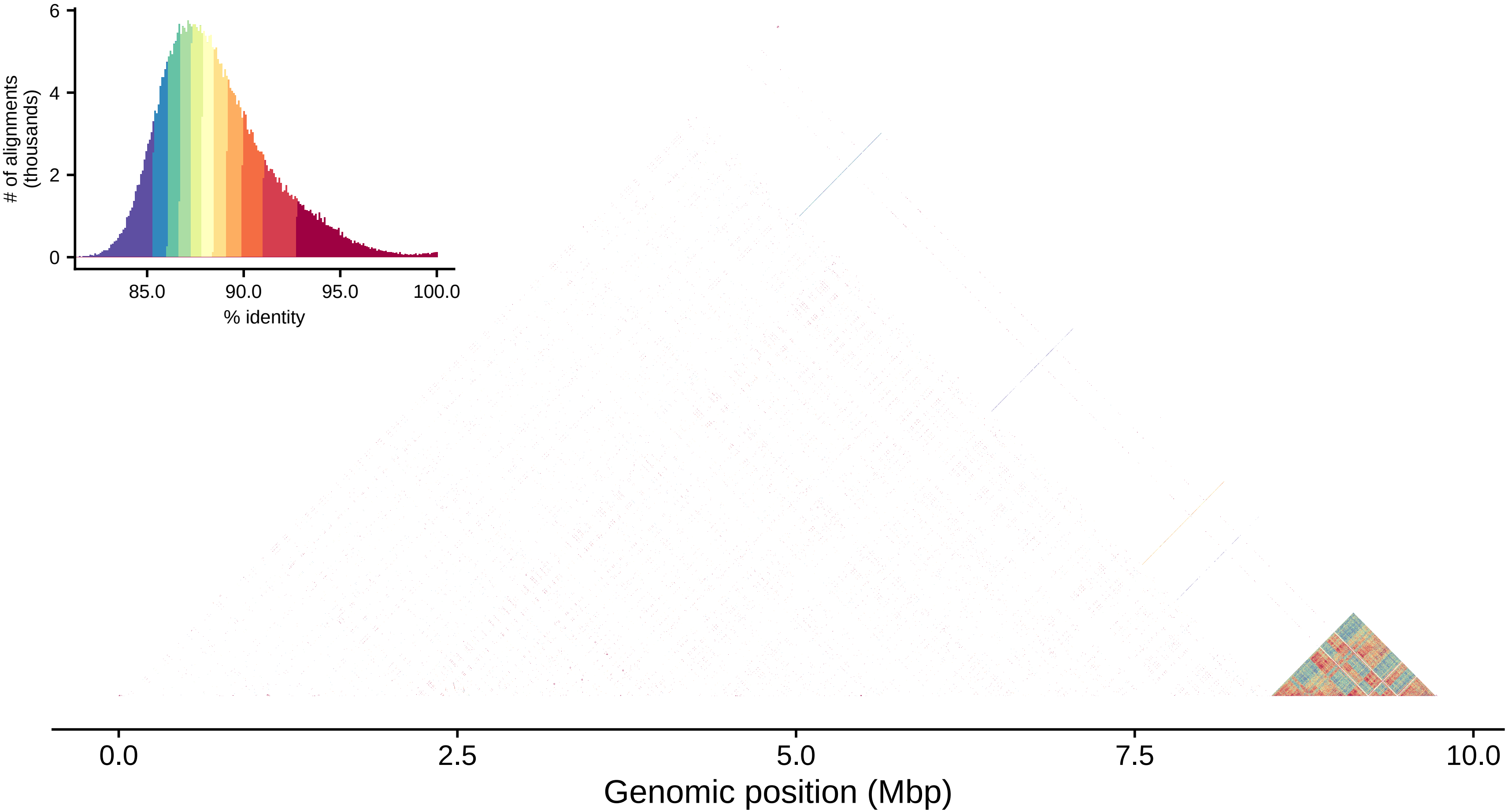

6
4
2
0
### of alignments
(thousands)
85.0
90.0
95.0
100.0
% identity
0.0
2.5
5.0
7.5
10.0
Genomic position (Mbp)

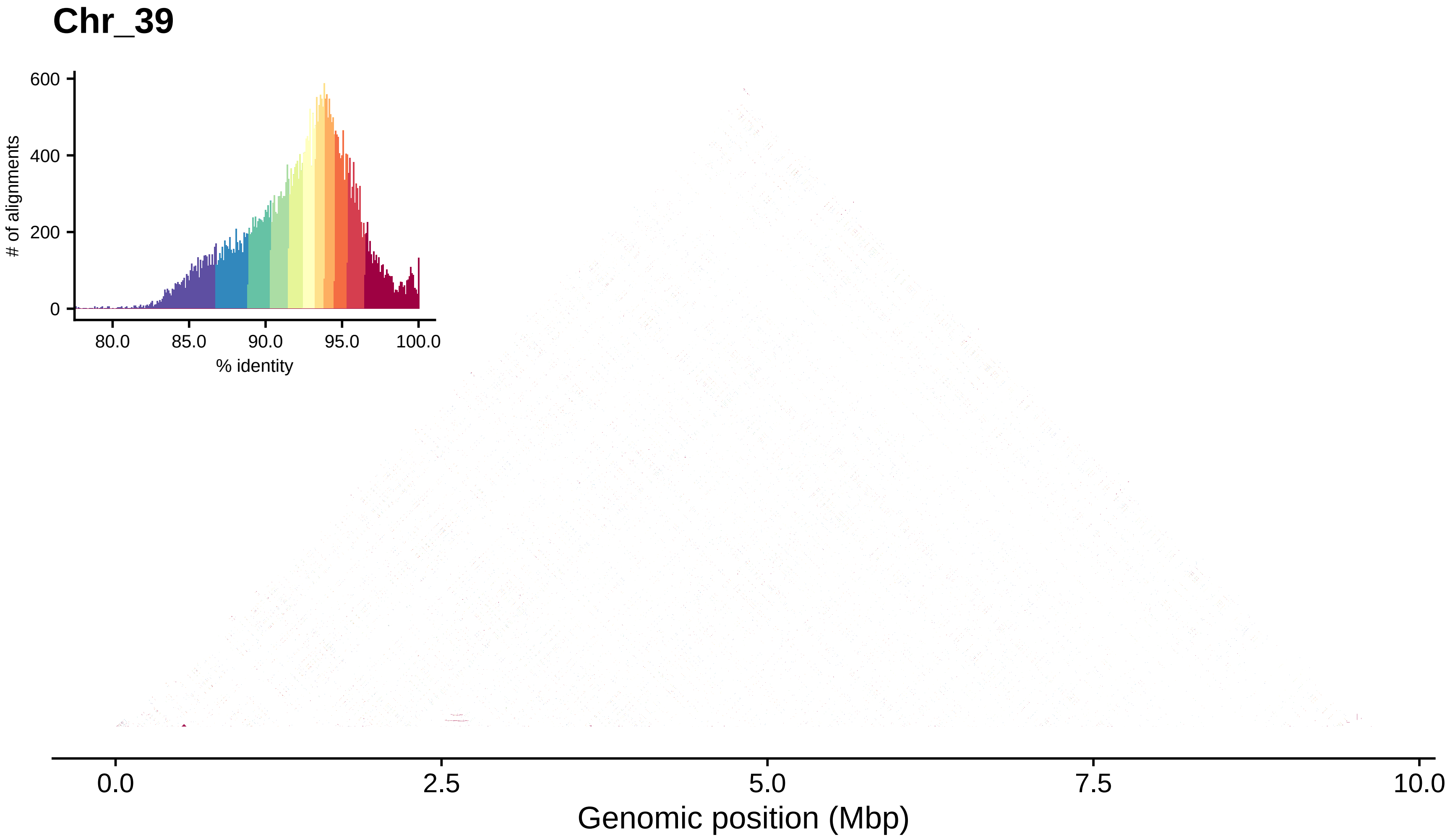
Chr_39
600
400
200
0
### of alignments
80.0
85.0
90.0
95.0
100.0
% identity
0.0
2.5
5.0
7.5
10.0
Genomic position (Mbp)

### Chr_40

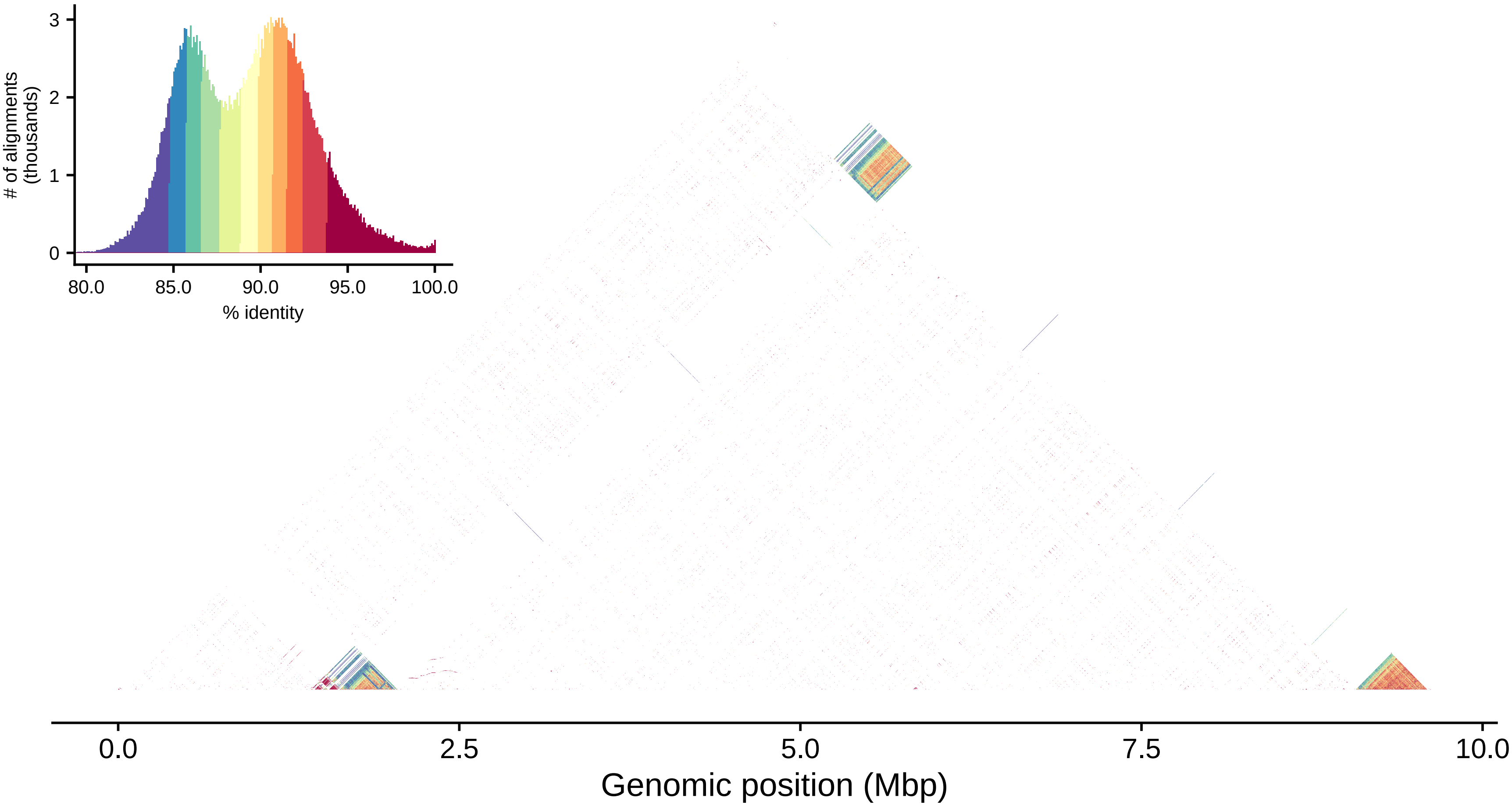

### of alignments
(thousands)
0
1
2
3
80.0
85.0
90.0
95.0
100.0
% identity
0.0
2.5
5.0
7.5
10.0
Genomic position (Mbp)

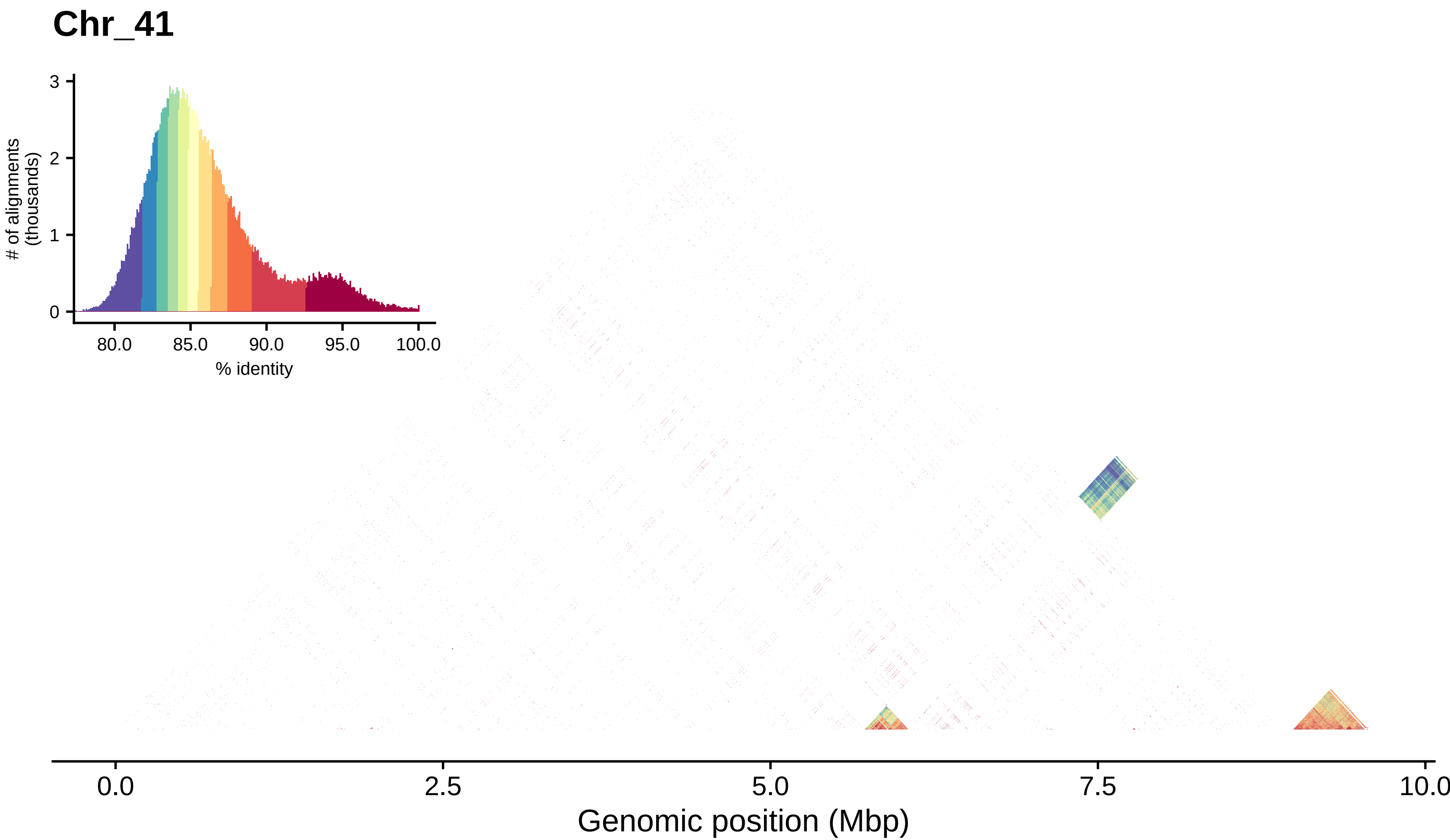
Chr_41
### of alignments
(thousands)
3
2
1
0
80.0
85.0
90.0
95.0
100.0
% identity
0.0
2.5
5.0
7.5
10.0
Genomic position (Mbp)

#### Chr_42

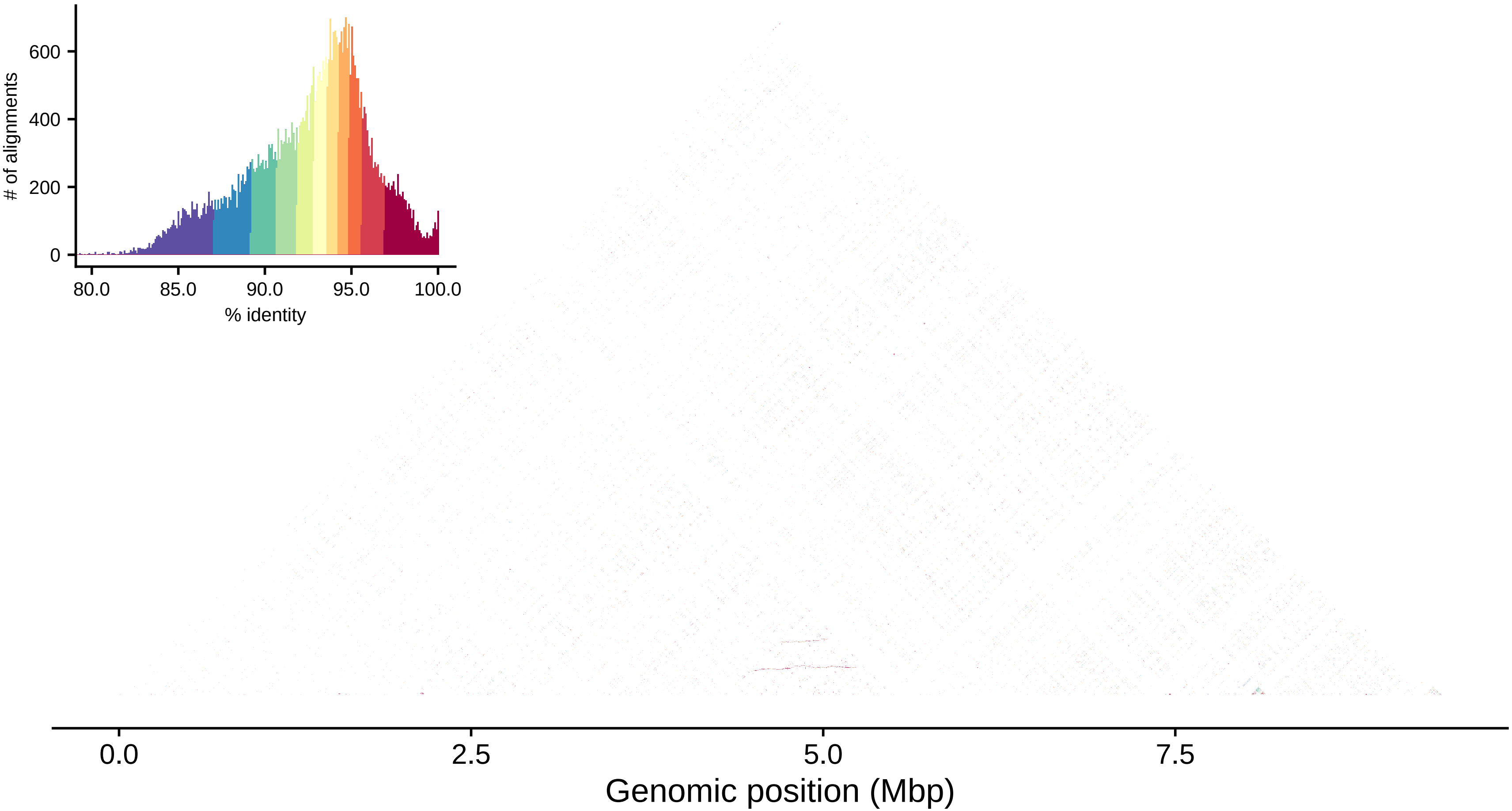

### of alignments
0
200
400
600
80.0
85.0
90.0
95.0
100.0
% identity
0.0
2.5
5.0
7.5
Genomic position (Mbp)

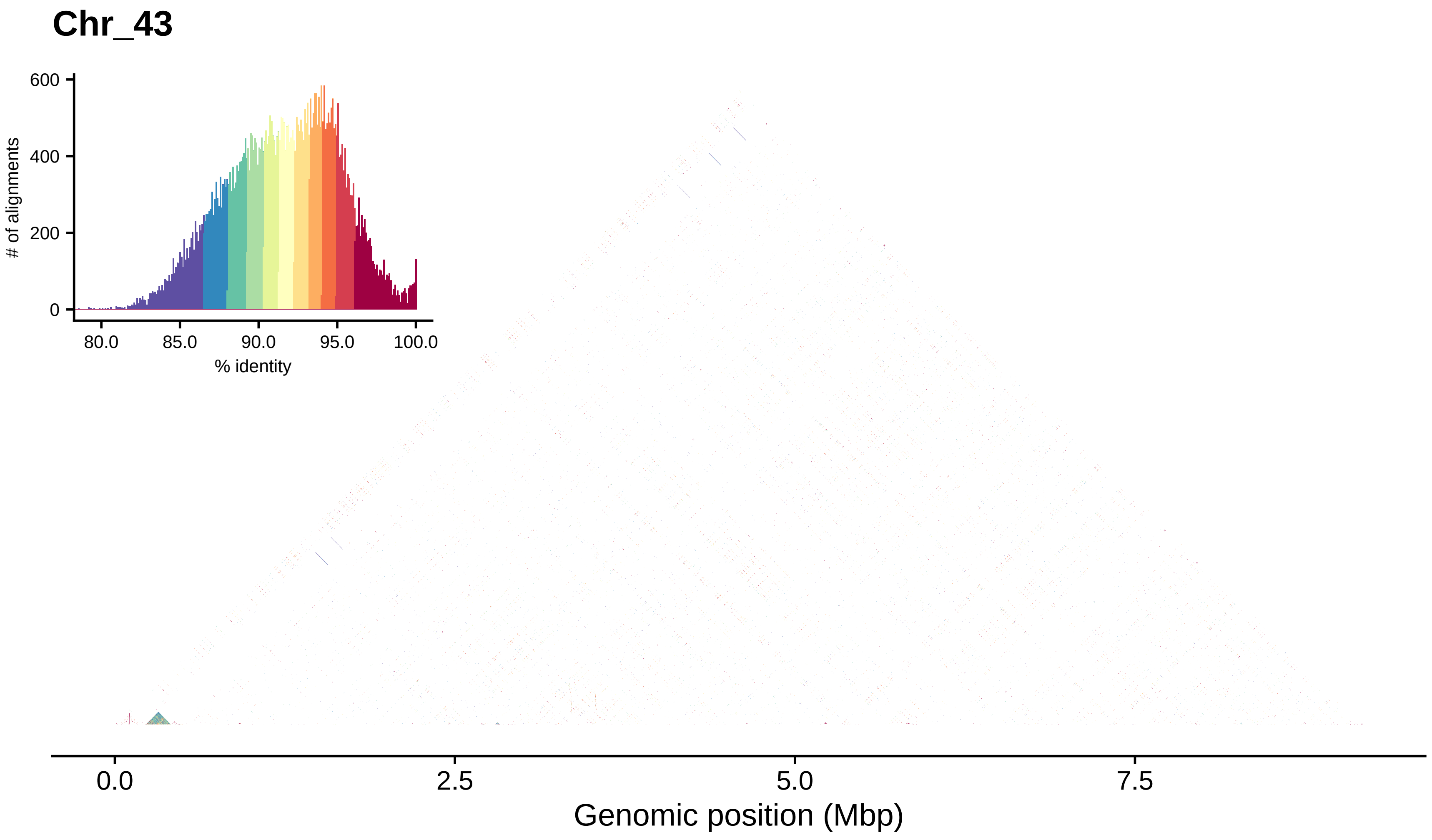
Chr_43
### of alignments
600
400
200
0
80.0
85.0
90.0
95.0
100.0
% identity
0.0
2.5
5.0
7.5
Genomic position (Mbp)

#### Chr_44

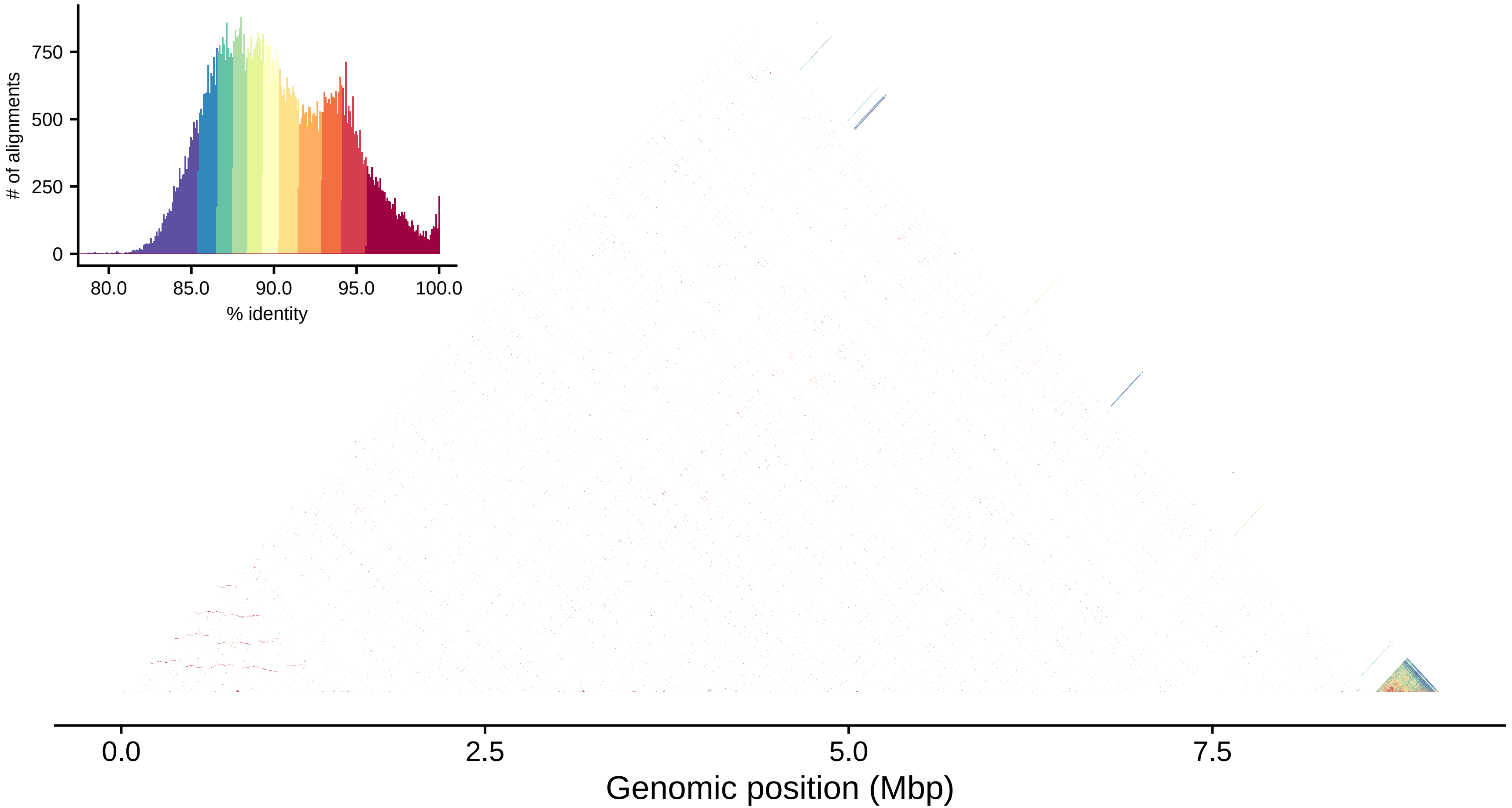

### of alignments
750
500
250
0
80.0
85.0
90.0
95.0
100.0
% identity
0.0
2.5
5.0
7.5
Genomic position (Mbp)

### Chr_45

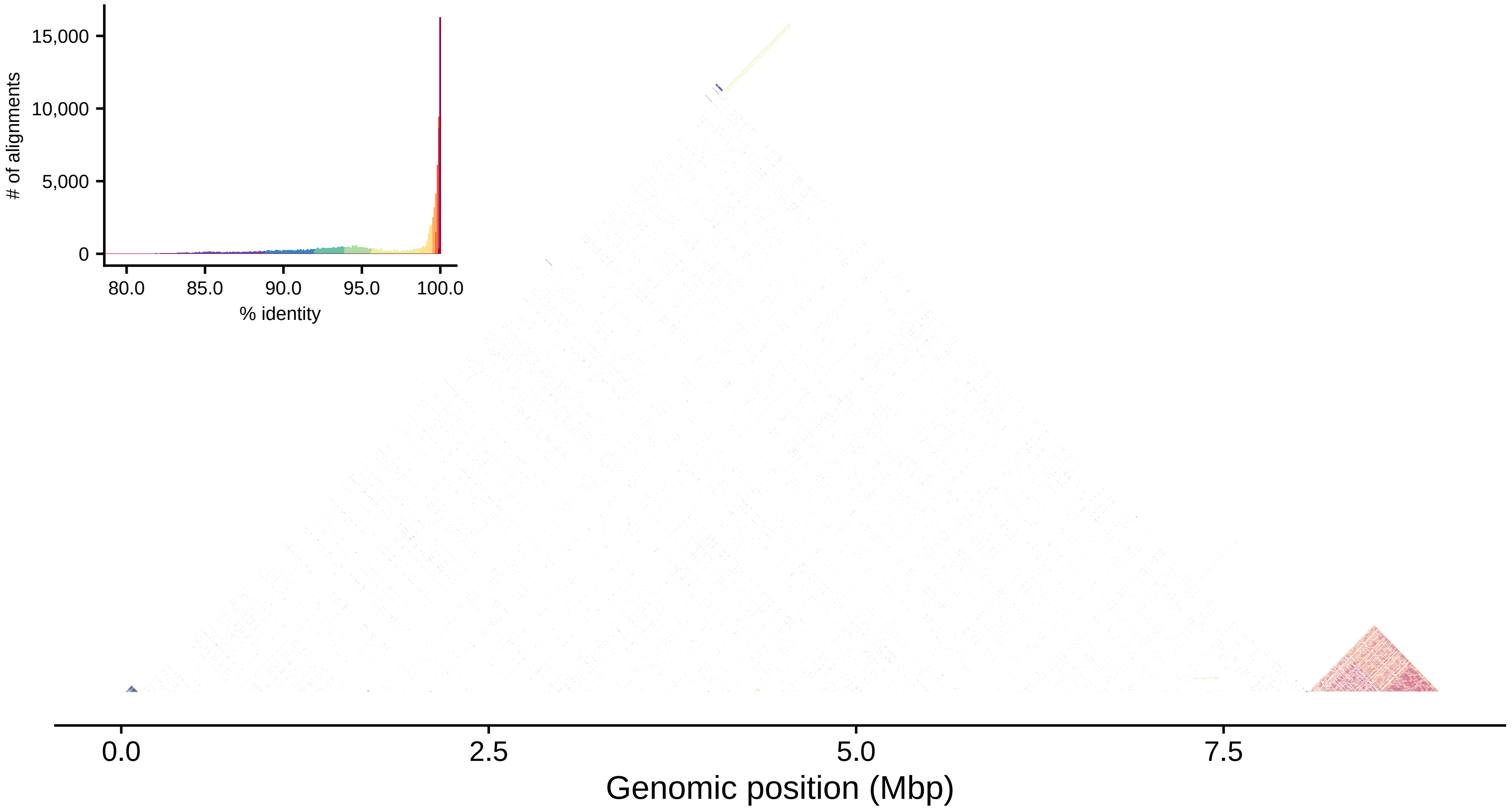
15,000
10,000
5,000
0
### of alignments
80.0
85.0
90.0
95.0
100.0
% identity
0.0
2.5
5.0
7.5
Genomic position (Mbp)

#### Chr_46

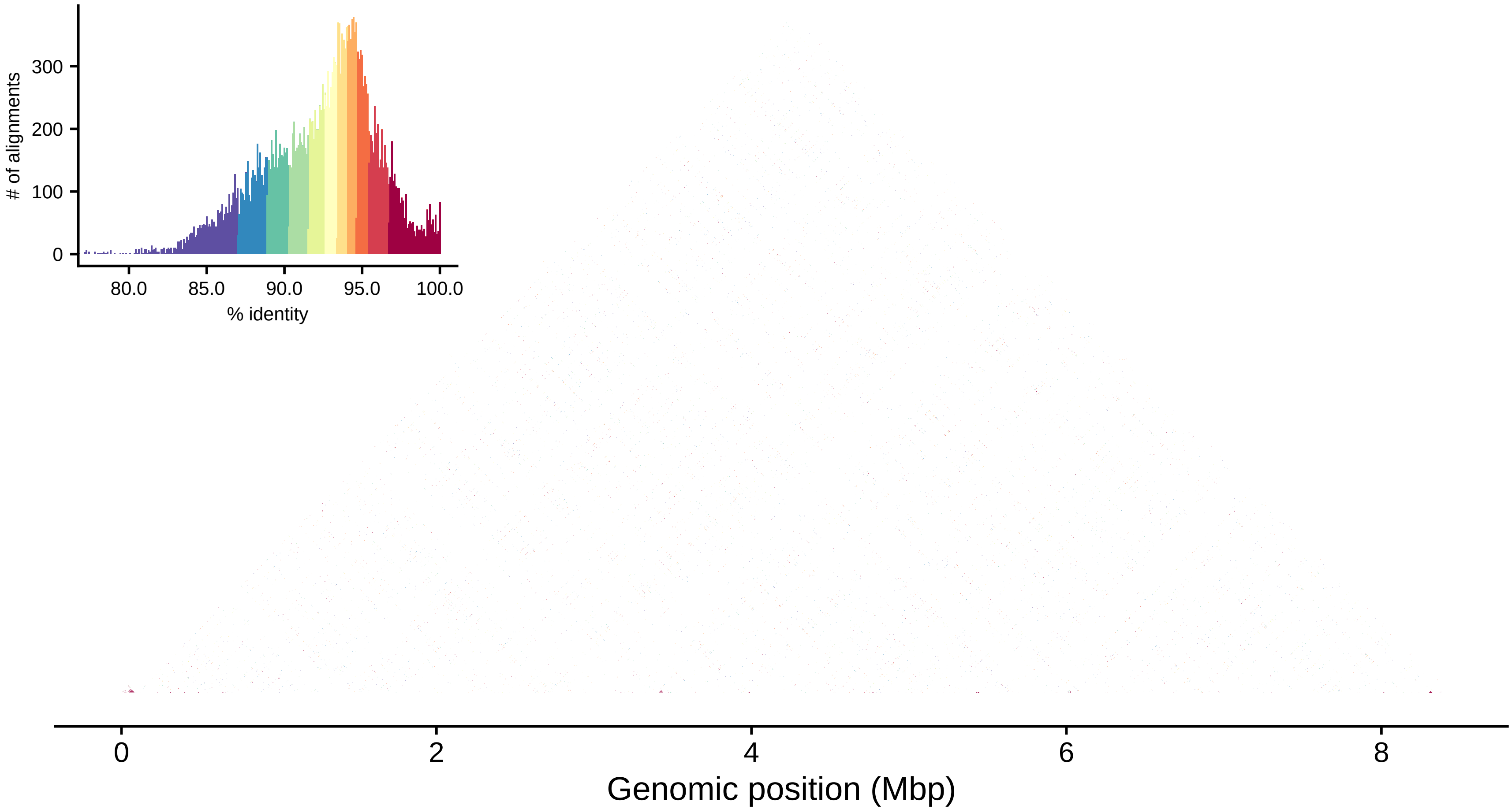

### of alignments
300
200
100
0
80.0
85.0
90.0
95.0
100.0
% identity
0
2
4
6
8
Genomic position (Mbp)

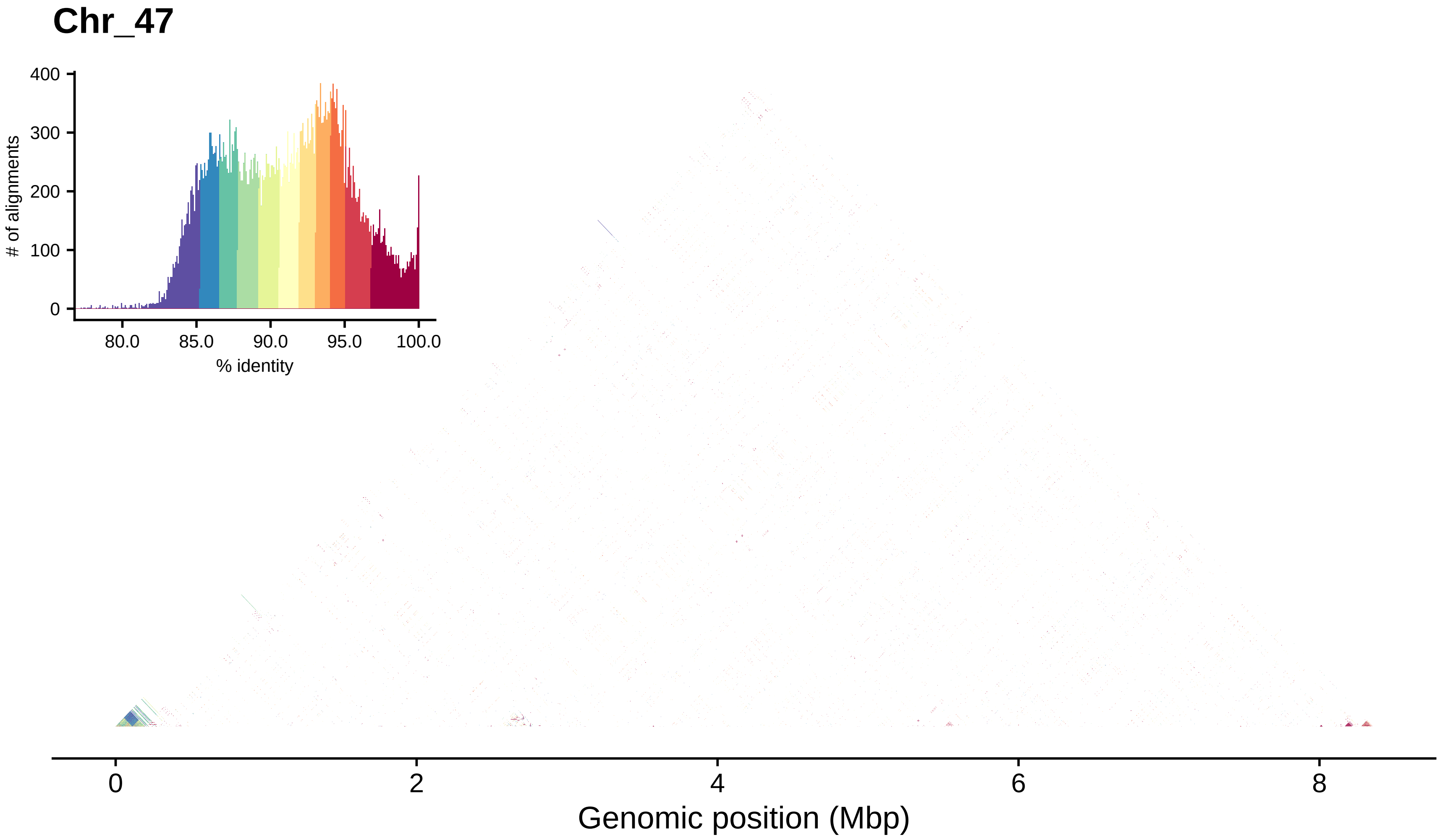

Chr_47
400
300
200
100
0
### of alignments
80.0
85.0
90.0
95.0
100.0
% identity
0
2
4
6
8
Genomic position (Mbp)

### Chr_48

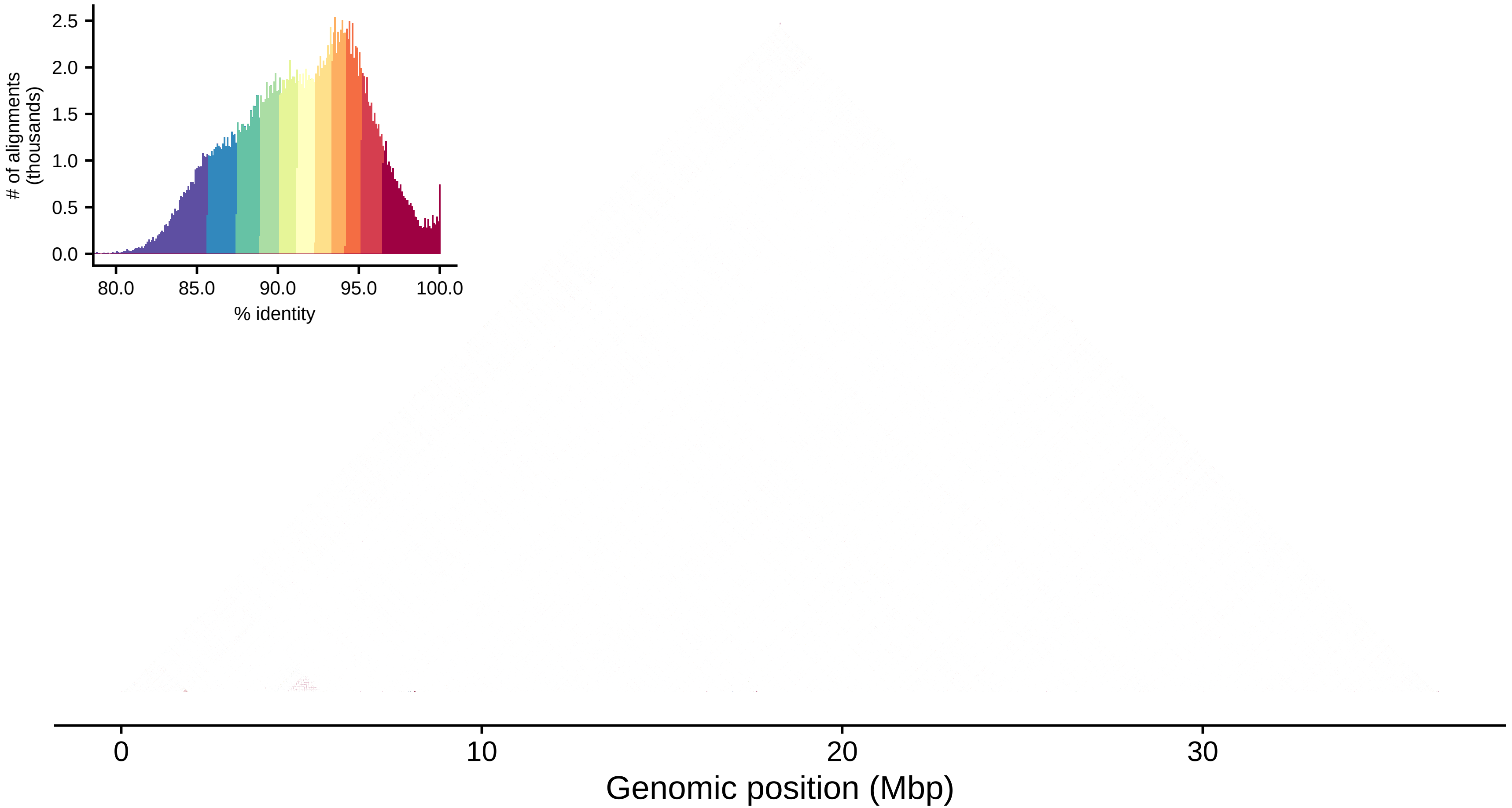

### of alignments
(thousands)
2.5
2.0
1.5
1.0
0.5
0.0
80.0
85.0
90.0
95.0
100.0
% identity
0
10
20
30
Genomic position (Mbp)
