## Additional File 2 for "Determinants of chromosomal rearrangements in holocentric *Leptidea* butterflies"

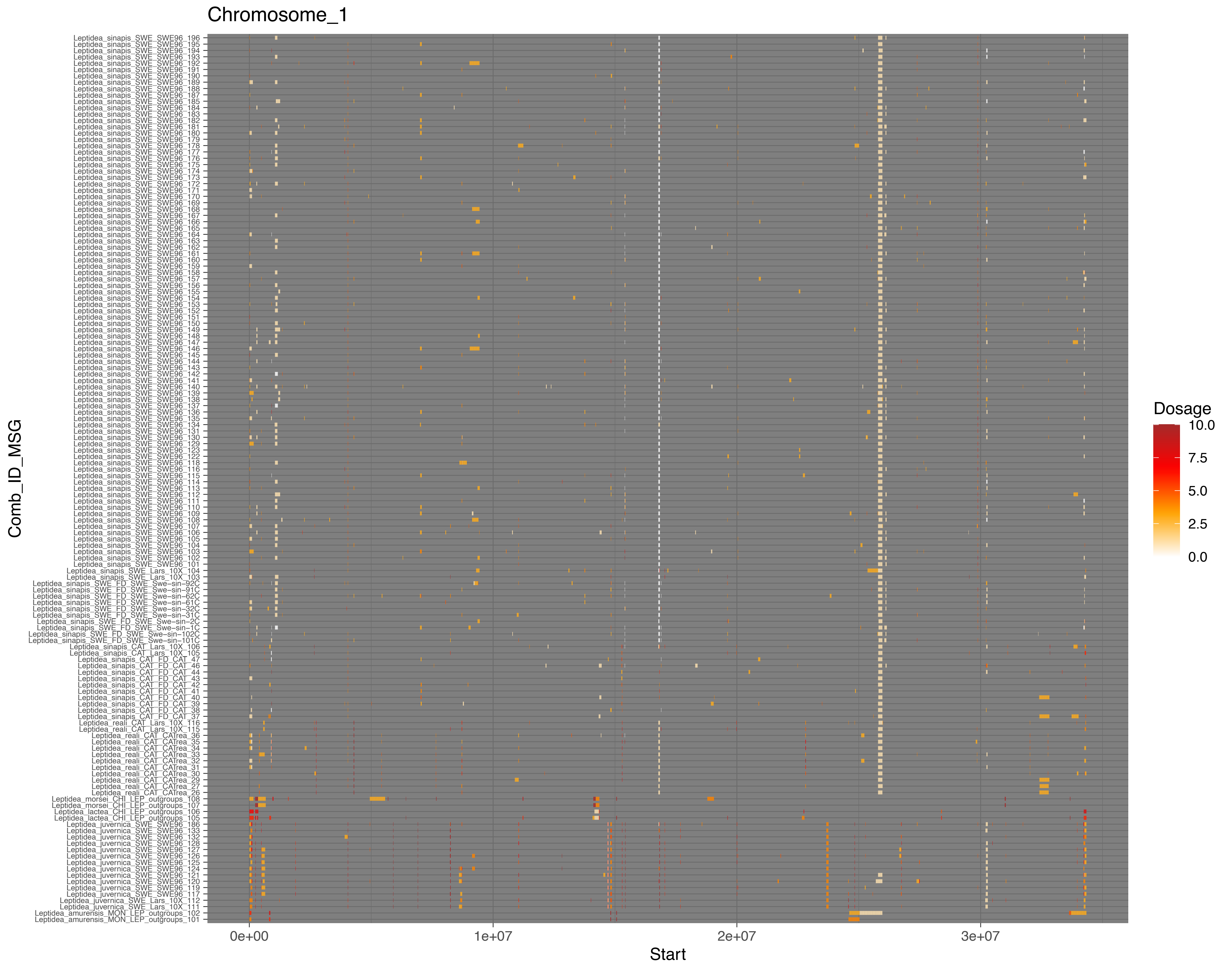

Chromosome_1
Comb_ID_MSG
Leptidea_sinapis_SWE_SWE96_196
Leptidea_sinapis_SWE_SWE96_195
Leptidea_sinapis_SWE_SWE96_194
Leptidea_sinapis_SWE_SWE96_193
Leptidea_sinapis_SWE_SWE96_192
Leptidea_sinapis_SWE_SWE96_191
Leptidea_sinapis_SWE_SWE96_190
Leptidea_sinapis_SWE_SWE96_189
Leptidea_sinapis_SWE_SWE96_188
Leptidea_sinapis_SWE_SWE96_187
Leptidea_sinapis_SWE_SWE96_185
Leptidea_sinapis_SWE_SWE96_184
Leptidea_sinapis_SWE_SWE96_183
Leptidea_sinapis_SWE_SWE96_182
Leptidea_sinapis_SWE_SWE96_181
Leptidea_sinapis_SWE_SWE96_180
Leptidea_sinapis_SWE_SWE96_179
Leptidea_sinapis_SWE_SWE96_178
Leptidea_sinapis_SWE_SWE96_177
Leptidea_sinapis_SWE_SWE96_176
Leptidea_sinapis_SWE_SWE96_175
Leptidea_sinapis_SWE_SWE96_174
Leptidea_sinapis_SWE_SWE96_173
Leptidea_sinapis_SWE_SWE96_172
Leptidea_sinapis_SWE_SWE96_171
Leptidea_sinapis_SWE_SWE96_170
Leptidea_sinapis_SWE_SWE96_169
Leptidea_sinapis_SWE_SWE96_168
Leptidea_sinapis_SWE_SWE96_167
Leptidea_sinapis_SWE_SWE96_166
Leptidea_sinapis_SWE_SWE96_165
Leptidea_sinapis_SWE_SWE96_164
Leptidea_sinapis_SWE_SWE96_163
Leptidea_sinapis_SWE_SWE96_162
Leptidea_sinapis_SWE_SWE96_161
Leptidea_sinapis_SWE_SWE96_160
Leptidea_sinapis_SWE_SWE96_159
Leptidea_sinapis_SWE_SWE96_158
Leptidea_sinapis_SWE_SWE96_157
Leptidea_sinapis_SWE_SWE96_156
Leptidea_sinapis_SWE_SWE96_155
Leptidea_sinapis_SWE_SWE96_154
Leptidea_sinapis_SWE_SWE96_153
Leptidea_sinapis_SWE_SWE96_152
Leptidea_sinapis_SWE_SWE96_151
Leptidea_sinapis_SWE_SWE96_150
Leptidea_sinapis_SWE_SWE96_149
Leptidea_sinapis_SWE_SWE96_148
Leptidea_sinapis_SWE_SWE96_147
Leptidea_sinapis_SWE_SWE96_146
Leptidea_sinapis_SWE_SWE96_145
Leptidea_sinapis_SWE_SWE96_144
Leptidea_sinapis_SWE_SWE96_143
Leptidea_sinapis_SWE_SWE96_142
Leptidea_sinapis_SWE_SWE96_141
Leptidea_sinapis_SWE_SWE96_140
Leptidea_sinapis_SWE_SWE96_139
Leptidea_sinapis_SWE_SWE96_138
Leptidea_sinapis_SWE_SWE96_137
Leptidea_sinapis_SWE_SWE96_136
Leptidea_sinapis_SWE_SWE96_135
Leptidea_sinapis_SWE_SWE96_134
Leptidea_sinapis_SWE_SWE96_131
Leptidea_sinapis_SWE_SWE96_130
Leptidea_sinapis_SWE_SWE96_129
Leptidea_sinapis_SWE_SWE96_123
Leptidea_sinapis_SWE_SWE96_122
Leptidea_sinapis_SWE_SWE96_118
Leptidea_sinapis_SWE_SWE96_116
Leptidea_sinapis_SWE_SWE96_115
Leptidea_sinapis_SWE_SWE96_114
Leptidea_sinapis_SWE_SWE96_113
Leptidea_sinapis_SWE_SWE96_112
Leptidea_sinapis_SWE_SWE96_111
Leptidea_sinapis_SWE_SWE96_110
Leptidea_sinapis_SWE_SWE96_109
Leptidea_sinapis_SWE_SWE96_108
Leptidea_sinapis_SWE_SWE96_107
Leptidea_sinapis_SWE_SWE96_106
Leptidea_sinapis_SWE_SWE96_105
Leptidea_sinapis_SWE_SWE96_104
Leptidea_sinapis_SWE_SWE96_103
Leptidea_sinapis_SWE_SWE96_102
Leptidea_sinapis_SWE_SWE96_101
Leptidea_sinapis_SWE_Lars_10X_104
Leptidea_sinapis_SWE_Lars_10X_103
Leptidea_sinapis_SWE_FD_SWE_Swe-sin-92C
Leptidea_sinapis_SWE_FD_SWE_Swe-sin-91C
Leptidea_sinapis_SWE_FD_SWE_Swe-sin-62C
Leptidea_sinapis_SWE_FD_SWE_Swe-sin-61C
Leptidea_sinapis_SWE_FD_SWE_Swe-sin-32C
Leptidea_sinapis_SWE_FD_SWE_Swe-sin-31C
Leptidea_sinapis_SWE_FD_SWE_Swe-sin-2C
Leptidea_sinapis_SWE_FD_SWE_Swe-sin-1C
Leptidea_sinapis_SWE_FD_SWE_Swe-sin-102C
Leptidea_sinapis_SWE_FD_SWE_Swe-sin-101C
Leptidea_sinapis_CAT_Lars_10X_106
Leptidea_sinapis_CAT_Lars_10X_105
Leptidea_sinapis_CAT_FD_CAT_47
Leptidea_sinapis_CAT_FD_CAT_46
Leptidea_sinapis_CAT_FD_CAT_44
Leptidea_sinapis_CAT_FD_CAT_43
Leptidea_sinapis_CAT_FD_CAT_42
Leptidea_sinapis_CAT_FD_CAT_41
Leptidea_sinapis_CAT_FD_CAT_40
Leptidea_sinapis_CAT_FD_CAT_39
Leptidea_sinapis_CAT_FD_CAT_38
Leptidea_sinapis_CAT_FD_CAT_37
Leptidea_reali_CAT_Lars_10X_116
Leptidea_reali_CAT_Lars_10X_115
Leptidea_reali_CAT_CATrea_36
Leptidea_reali_CAT_CATrea_35
Leptidea_reali_CAT_CATrea_34
Leptidea_reali_CAT_CATrea_33
Leptidea_reali_CAT_CATrea_32
Leptidea_reali_CAT_CATrea_31
Leptidea_reali_CAT_CATrea_30
Leptidea_reali_CAT_CATrea_29
Leptidea_reali_CAT_CATrea_27
Leptidea_reali_CAT_CATrea_26
Leptidea_morsei_CHI_LEP_outgroups_108
Leptidea_morsei_CHI_LEP_outgroups_107
Leptidea_lactea_CHI_LEP_outgroups_106
Leptidea_lactea_CHI_LEP_outgroups_105
Leptidea_juvernica_SWE_SWE96_186
Leptidea_juvernica_SWE_SWE96_133
Leptidea_juvernica_SWE_SWE96_132
Leptidea_juvernica_SWE_SWE96_128
Leptidea_juvernica_SWE_SWE96_127
Leptidea_juvernica_SWE_SWE96_126
Leptidea_juvernica_SWE_SWE96_125
Leptidea_juvernica_SWE_SWE96_124
Leptidea_juvernica_SWE_SWE96_121
Leptidea_juvernica_SWE_SWE96_120
Leptidea_juvernica_SWE_SWE96_119
Leptidea_juvernica_SWE_SWE96_117
Leptidea_juvernica_SWE_Lars_10X_112
Leptidea_juvernica_SWE_Lars_10X_11
Leptidea_amurensis_MON_LEP_outgroups_102
Leptidea_amurensis_MON_LEP_outgroups_101
0e+00
1e+07
2e+07
3e+07
Start
Dosage
10.0
7.5
5.0
2.5
0.0

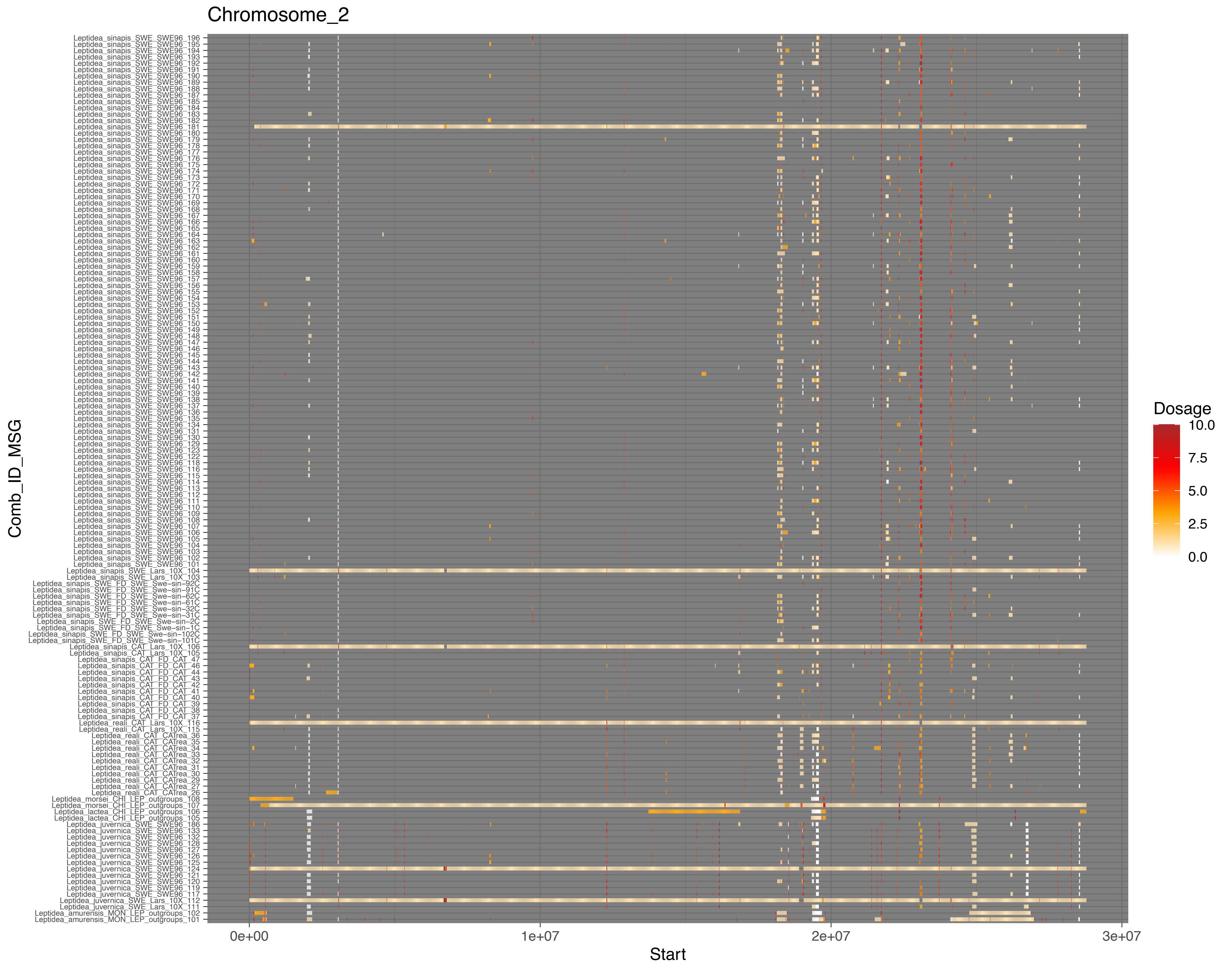
Chromosome_2
Leptidea_sinapis_SWE_SWE96_196
Leptidea_sinapis_SWE_SWE96_195
Leptidea_sinapis_SWE_SWE96_194
Leptidea_sinapis_SWE_SWE96_193
Leptidea_sinapis_SWE_SWE96_192
Leptidea_sinapis_SWE_SWE96_191
Leptidea_sinapis_SWE_SWE96_190
Leptidea_sinapis_SWE_SWE96_189
Leptidea_sinapis_SWE_SWE96_188
Leptidea_sinapis_SWE_SWE96_187
Leptidea_sinapis_SWE_SWE96_185
Leptidea_sinapis_SWE_SWE96_184
Leptidea_sinapis_SWE_SWE96_183
Leptidea_sinapis_SWE_SWE96_182
Leptidea_sinapis_SWE_SWE96_181
Leptidea_sinapis_SWE_SWE96_180
Leptidea_sinapis_SWE_SWE96_179
Leptidea_sinapis_SWE_SWE96_178
Leptidea_sinapis_SWE_SWE96_177
Leptidea_sinapis_SWE_SWE96_176
Leptidea_sinapis_SWE_SWE96_175
Leptidea_sinapis_SWE_SWE96_174
Leptidea_sinapis_SWE_SWE96_173
Leptidea_sinapis_SWE_SWE96_172
Leptidea_sinapis_SWE_SWE96_171
Leptidea_sinapis_SWE_SWE96_170
Leptidea_sinapis_SWE_SWE96_169
Leptidea_sinapis_SWE_SWE96_168
Leptidea_sinapis_SWE_SWE96_167
Leptidea_sinapis_SWE_SWE96_166
Leptidea_sinapis_SWE_SWE96_165
Leptidea_sinapis_SWE_SWE96_164
Leptidea_sinapis_SWE_SWE96_163
Leptidea_sinapis_SWE_SWE96_162
Leptidea_sinapis_SWE_SWE96_161
Leptidea_sinapis_SWE_SWE96_160
Leptidea_sinapis_SWE_SWE96_159
Leptidea_sinapis_SWE_SWE96_158
Leptidea_sinapis_SWE_SWE96_157
Leptidea_sinapis_SWE_SWE96_156
Leptidea_sinapis_SWE_SWE96_155
Leptidea_sinapis_SWE_SWE96_154
Leptidea_sinapis_SWE_SWE96_153
Leptidea_sinapis_SWE_SWE96_152
Leptidea_sinapis_SWE_SWE96_151
Leptidea_sinapis_SWE_SWE96_150
Leptidea_sinapis_SWE_SWE96_149
Leptidea_sinapis_SWE_SWE96_148
Leptidea_sinapis_SWE_SWE96_147
Leptidea_sinapis_SWE_SWE96_146
Leptidea_sinapis_SWE_SWE96_145
Leptidea_sinapis_SWE_SWE96_144
Leptidea_sinapis_SWE_SWE96_143
Leptidea_sinapis_SWE_SWE96_142
Leptidea_sinapis_SWE_SWE96_141
Leptidea_sinapis_SWE_SWE96_140
Leptidea_sinapis_SWE_SWE96_139
Leptidea_sinapis_SWE_SWE96_138
Leptidea_sinapis_SWE_SWE96_137
Leptidea_sinapis_SWE_SWE96_136
Leptidea_sinapis_SWE_SWE96_135
Leptidea_sinapis_SWE_SWE96_134
Leptidea_sinapis_SWE_SWE96_131
Leptidea_sinapis_SWE_SWE96_130
Leptidea_sinapis_SWE_SWE96_129
Leptidea_sinapis_SWE_SWE96_123
Leptidea_sinapis_SWE_SWE96_122
Leptidea_sinapis_SWE_SWE96_118
Leptidea_sinapis_SWE_SWE96_116
Leptidea_sinapis_SWE_SWE96_115
Leptidea_sinapis_SWE_SWE96_114
Leptidea_sinapis_SWE_SWE96_113
Leptidea_sinapis_SWE_SWE96_112
Leptidea_sinapis_SWE_SWE96_111
Leptidea_sinapis_SWE_SWE96_110
Leptidea_sinapis_SWE_SWE96_109
Leptidea_sinapis_SWE_SWE96_108
Leptidea_sinapis_SWE_SWE96_107
Leptidea_sinapis_SWE_SWE96_106
Leptidea_sinapis_SWE_SWE96_105
Leptidea_sinapis_SWE_SWE96_104
Leptidea_sinapis_SWE_SWE96_103
Leptidea_sinapis_SWE_SWE96_102
Leptidea_sinapis_SWE_SWE96_101
Leptidea_sinapis_SWE_Lars_10X_104
Leptidea_sinapis_SWE_Lars_10X_103
Leptidea_sinapis_SWE_FD_SWE_Swe-sin-92C
Leptidea_sinapis_SWE_FD_SWE_Swe-sin-91C
Leptidea_sinapis_SWE_FD_SWE_Swe-sin-62C
Leptidea_sinapis_SWE_FD_SWE_Swe-sin-61C
Leptidea_sinapis_SWE_FD_SWE_Swe-sin-32C
Leptidea_sinapis_SWE_FD_SWE_Swe-sin-31C
Leptidea_sinapis_SWE_FD_SWE_Swe-sin-2C
Leptidea_sinapis_SWE_FD_SWE_Swe-sin-1C
Leptidea_sinapis_SWE_FD_SWE_Swe-sin-102C
Leptidea_sinapis_SWE_FD_SWE_Swe-sin-101C
Leptidea_sinapis_CAT_Lars_10X_106
Leptidea_sinapis_CAT_Lars_10X_105
Leptidea_sinapis_CAT_FD_CAT_47
Leptidea_sinapis_CAT_FD_CAT_46
Leptidea_sinapis_CAT_FD_CAT_44
Leptidea_sinapis_CAT_FD_CAT_43
Leptidea_sinapis_CAT_FD_CAT_42
Leptidea_sinapis_CAT_FD_CAT_41
Leptidea_sinapis_CAT_FD_CAT_40
Leptidea_sinapis_CAT_FD_CAT_39
Leptidea_sinapis_CAT_FD_CAT_38
Leptidea_sinapis_CAT_FD_CAT_37
Leptidea_reali_CAT_Lars_10X_116
Leptidea_reali_CAT_Lars_10X_115
Leptidea_reali_CAT_CATrea_36
Leptidea_reali_CAT_CATrea_35
Leptidea_reali_CAT_CATrea_34
Leptidea_reali_CAT_CATrea_33
Leptidea_reali_CAT_CATrea_32
Leptidea_reali_CAT_CATrea_31
Leptidea_reali_CAT_CATrea_30
Leptidea_reali_CAT_CATrea_29
Leptidea_reali_CAT_CATrea_27
Leptidea_reali_CAT_CATrea_26
Leptidea_morsei_CHI_LEP_outgroups_108
Leptidea_morsei_CHI_LEP_outgroups_107
Leptidea_lactea_CHI_LEP_outgroups_106
Leptidea_lactea_CHI_LEP_outgroups_105
Leptidea_juvernica_SWE_SWE96_186
Leptidea_juvernica_SWE_SWE96_133
Leptidea_juvernica_SWE_SWE96_132
Leptidea_juvernica_SWE_SWE96_128
Leptidea_juvernica_SWE_SWE96_127
Leptidea_juvernica_SWE_SWE96_126
Leptidea_juvernica_SWE_SWE96_125
Leptidea_juvernica_SWE_SWE96_124
Leptidea_juvernica_SWE_SWE96_121
Leptidea_juvernica_SWE_SWE96_120
Leptidea_juvernica_SWE_SWE96_119
Leptidea_juvernica_SWE_SWE96_117
Leptidea_juvernica_SWE_Lars_10X_112
Leptidea_juvernica_SWE_Lars_10X_111
Leptidea_amurensis_MON_LEP_outgroups_102
Leptidea_amurensis_MON_LEP_outgroups_101
Comb_ID_MSG
0e+00
1e+07
2e+07
3e+07
Start
Dosage
10.0
7.5
5.0
2.5
0.0

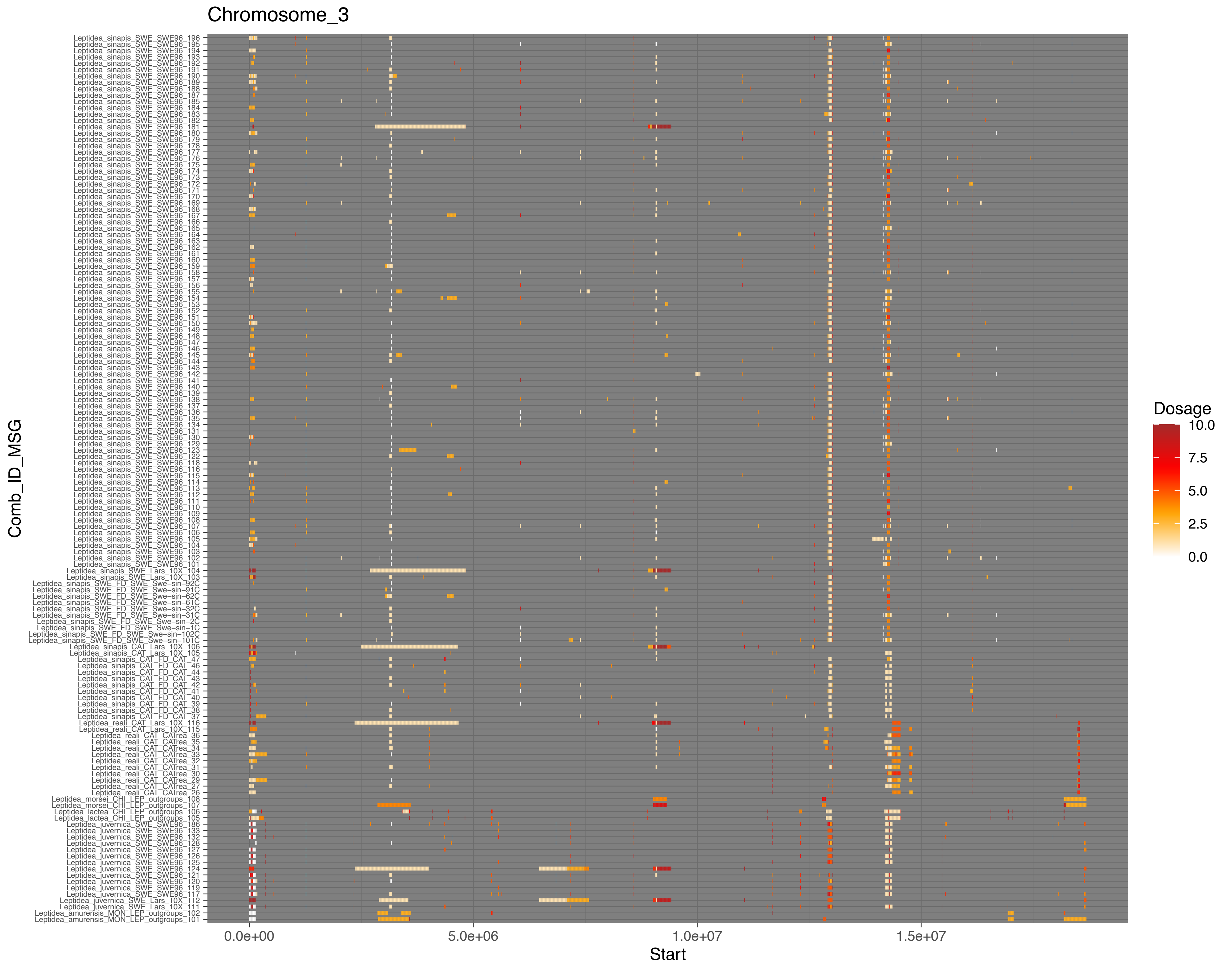
Chromosome_3
Comb_ID_MSG
Start
0.0e+00
5.0e+06
1.0e+07
1.5e+07
Dosage
10.0
7.5
5.0
2.5
0.0

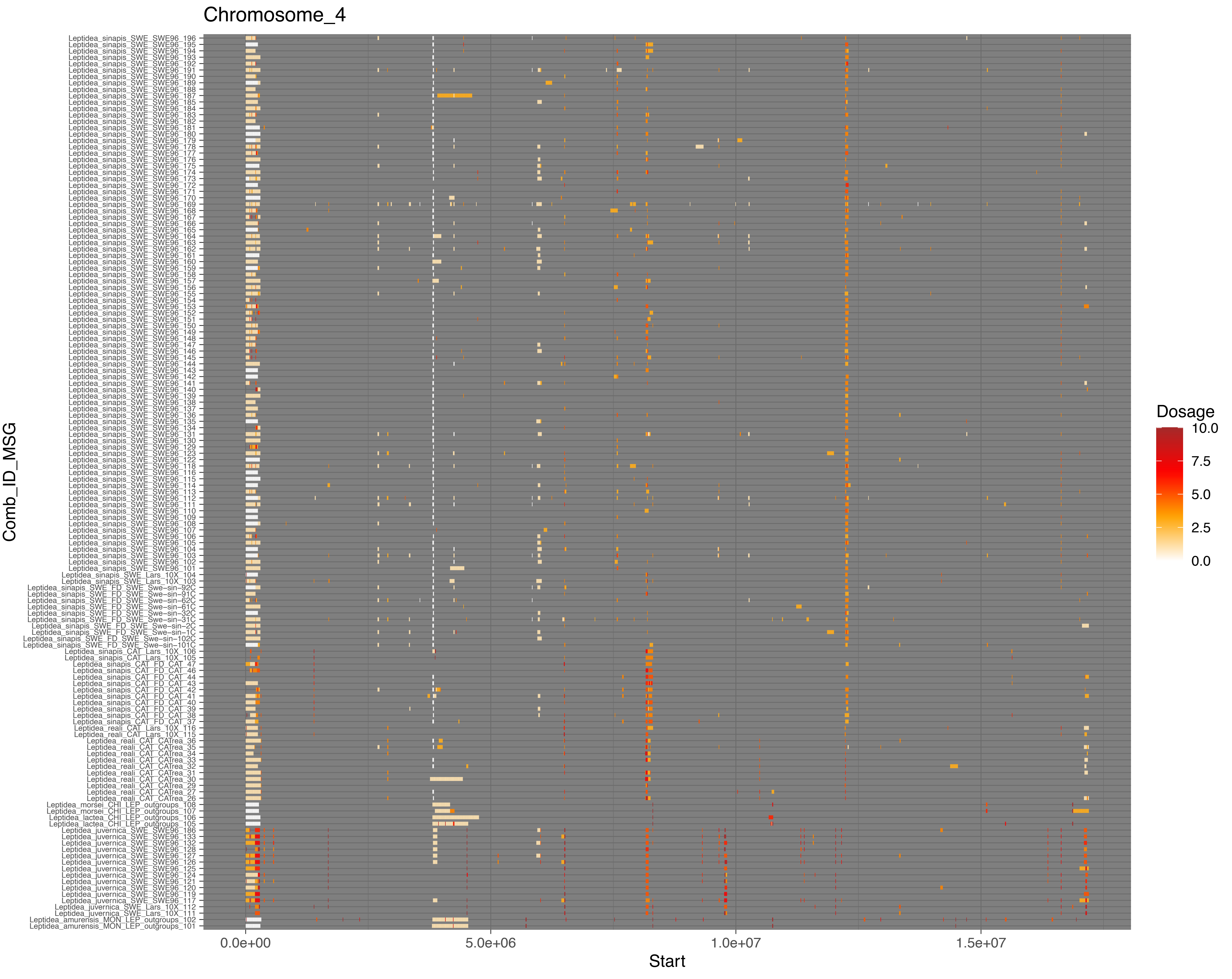
Chromosome_4
Leptidea_sinapis_SWE_SWE96_196
Leptidea_sinapis_SWE_SWE96_195
Leptidea_sinapis_SWE_SWE96_194
Leptidea_sinapis_SWE_SWE96_193
Leptidea_sinapis_SWE_SWE96_192
Leptidea_sinapis_SWE_SWE96_191
Leptidea_sinapis_SWE_SWE96_190
Leptidea_sinapis_SWE_SWE96_189
Leptidea_sinapis_SWE_SWE96_188
Leptidea_sinapis_SWE_SWE96_187
Leptidea_sinapis_SWE_SWE96_185
Leptidea_sinapis_SWE_SWE96_184
Leptidea_sinapis_SWE_SWE96_183
Leptidea_sinapis_SWE_SWE96_182
Leptidea_sinapis_SWE_SWE96_181
Leptidea_sinapis_SWE_SWE96_180
Leptidea_sinapis_SWE_SWE96_179
Leptidea_sinapis_SWE_SWE96_178
Leptidea_sinapis_SWE_SWE96_177
Leptidea_sinapis_SWE_SWE96_176
Leptidea_sinapis_SWE_SWE96_175
Leptidea_sinapis_SWE_SWE96_174
Leptidea_sinapis_SWE_SWE96_173
Leptidea_sinapis_SWE_SWE96_172
Leptidea_sinapis_SWE_SWE96_171
Leptidea_sinapis_SWE_SWE96_170
Leptidea_sinapis_SWE_SWE96_169
Leptidea_sinapis_SWE_SWE96_168
Leptidea_sinapis_SWE_SWE96_167
Leptidea_sinapis_SWE_SWE96_166
Leptidea_sinapis_SWE_SWE96_165
Leptidea_sinapis_SWE_SWE96_164
Leptidea_sinapis_SWE_SWE96_163
Leptidea_sinapis_SWE_SWE96_162
Leptidea_sinapis_SWE_SWE96_161
Leptidea_sinapis_SWE_SWE96_160
Leptidea_sinapis_SWE_SWE96_159
Leptidea_sinapis_SWE_SWE96_158
Leptidea_sinapis_SWE_SWE96_157
Leptidea_sinapis_SWE_SWE96_156
Leptidea_sinapis_SWE_SWE96_155
Leptidea_sinapis_SWE_SWE96_154
Leptidea_sinapis_SWE_SWE96_153
Leptidea_sinapis_SWE_SWE96_152
Leptidea_sinapis_SWE_SWE96_151
Leptidea_sinapis_SWE_SWE96_150
Leptidea_sinapis_SWE_SWE96_149
Leptidea_sinapis_SWE_SWE96_148
Leptidea_sinapis_SWE_SWE96_147
Leptidea_sinapis_SWE_SWE96_146
Leptidea_sinapis_SWE_SWE96_145
Leptidea_sinapis_SWE_SWE96_144
Leptidea_sinapis_SWE_SWE96_143
Leptidea_sinapis_SWE_SWE96_142
Leptidea_sinapis_SWE_SWE96_141
Leptidea_sinapis_SWE_SWE96_140
Leptidea_sinapis_SWE_SWE96_139
Leptidea_sinapis_SWE_SWE96_138
Leptidea_sinapis_SWE_SWE96_137
Leptidea_sinapis_SWE_SWE96_136
Leptidea_sinapis_SWE_SWE96_135
Leptidea_sinapis_SWE_SWE96_134
Leptidea_sinapis_SWE_SWE96_131
Leptidea_sinapis_SWE_SWE96_130
Leptidea_sinapis_SWE_SWE96_129
Leptidea_sinapis_SWE_SWE96_123
Leptidea_sinapis_SWE_SWE96_122
Leptidea_sinapis_SWE_SWE96_118
Leptidea_sinapis_SWE_SWE96_116
Leptidea_sinapis_SWE_SWE96_115
Leptidea_sinapis_SWE_SWE96_114
Leptidea_sinapis_SWE_SWE96_113
Leptidea_sinapis_SWE_SWE96_112
Leptidea_sinapis_SWE_SWE96_111
Leptidea_sinapis_SWE_SWE96_110
Leptidea_sinapis_SWE_SWE96_109
Leptidea_sinapis_SWE_SWE96_108
Leptidea_sinapis_SWE_SWE96_107
Leptidea_sinapis_SWE_SWE96_106
Leptidea_sinapis_SWE_SWE96_105
Leptidea_sinapis_SWE_SWE96_104
Leptidea_sinapis_SWE_SWE96_103
Leptidea_sinapis_SWE_SWE96_102
Leptidea_sinapis_SWE_SWE96_101
Leptidea_sinapis_SWE_Lars_10X_104
Leptidea_sinapis_SWE_Lars_10X_103
Leptidea_sinapis_SWE_FD_SWE_Swe–sin–92C
Leptidea_sinapis_SWE_FD_SWE_Swe–sin–91C
Leptidea_sinapis_SWE_FD_SWE_Swe–sin–62C
Leptidea_sinapis_SWE_FD_SWE_Swe–sin–61C
Leptidea_sinapis_SWE_FD_SWE_Swe–sin–32C
Leptidea_sinapis_SWE_FD_SWE_Swe–sin–31C
Leptidea_sinapis_SWE_FD_SWE_Swe–sin–2C
Leptidea_sinapis_SWE_FD_SWE_Swe–sin–1C
Leptidea_sinapis_SWE_FD_SWE_Swe–sin–102C
Leptidea_sinapis_SWE_FD_SWE_Swe–sin–101C
Leptidea_sinapis_CAT_Lars_10X_106
Leptidea_sinapis_CAT_Lars_10X_105
Leptidea_sinapis_CAT_FD_CAT_47
Leptidea_sinapis_CAT_FD_CAT_46
Leptidea_sinapis_CAT_FD_CAT_44
Leptidea_sinapis_CAT_FD_CAT_43
Leptidea_sinapis_CAT_FD_CAT_42
Leptidea_sinapis_CAT_FD_CAT_41
Leptidea_sinapis_CAT_FD_CAT_40
Leptidea_sinapis_CAT_FD_CAT_39
Leptidea_sinapis_CAT_FD_CAT_38
Leptidea_sinapis_CAT_FD_CAT_37
Leptidea_reali_CAT_Lars_10X_116
Leptidea_reali_CAT_Lars_10X_115
Leptidea_reali_CAT_CATrea_36
Leptidea_reali_CAT_CATrea_35
Leptidea_reali_CAT_CATrea_34
Leptidea_reali_CAT_CATrea_33
Leptidea_reali_CAT_CATrea_32
Leptidea_reali_CAT_CATrea_31
Leptidea_reali_CAT_CATrea_30
Leptidea_reali_CAT_CATrea_29
Leptidea_reali_CAT_CATrea_27
Leptidea_reali_CAT_CATrea_26
Leptidea_morsei_CHI_LEP_outgroups_108
Leptidea_morsei_CHI_LEP_outgroups_107
Leptidea_lactea_CHI_LEP_outgroups_106
Leptidea_lactea_CHI_LEP_outgroups_105
Leptidea_juvernica_SWE_SWE96_186
Leptidea_juvernica_SWE_SWE96_133
Leptidea_juvernica_SWE_SWE96_132
Leptidea_juvernica_SWE_SWE96_128
Leptidea_juvernica_SWE_SWE96_127
Leptidea_juvernica_SWE_SWE96_126
Leptidea_juvernica_SWE_SWE96_125
Leptidea_juvernica_SWE_SWE96_124
Leptidea_juvernica_SWE_SWE96_121
Leptidea_juvernica_SWE_SWE96_120
Leptidea_juvernica_SWE_SWE96_119
Leptidea_juvernica_SWE_SWE96_117
Leptidea_juvernica_SWE_Lars_10X_112
Leptidea_juvernica_SWE_Lars_10X_111
Leptidea_amurensis_MON_LEP_outgroups_102
Leptidea_amurensis_MON_LEP_outgroups_101
Comb_ID_MSG
0.0e+00
5.0e+06
1.0e+07
1.5e+07
Start
Dosage
10.0
7.5
5.0
2.5
0.0

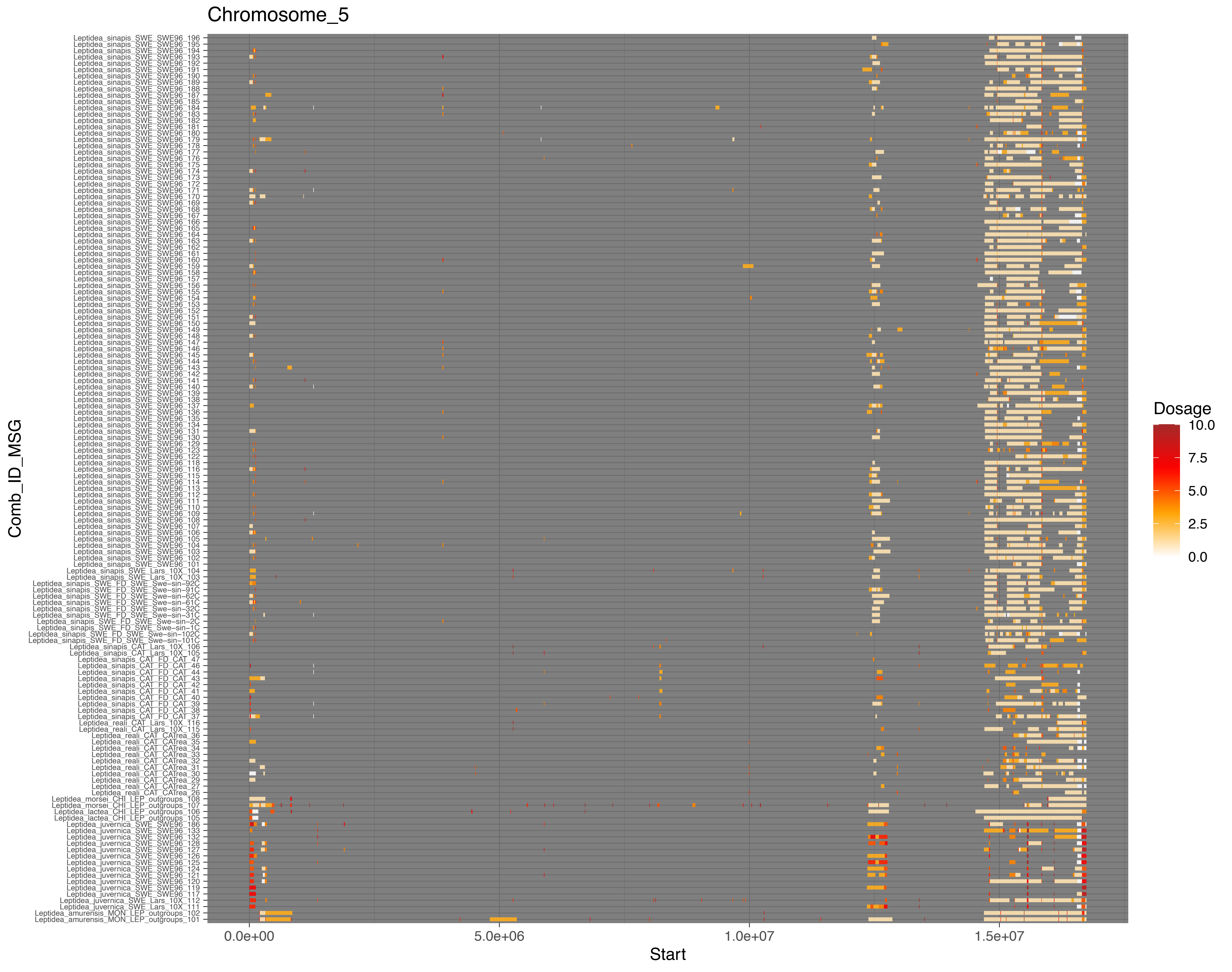

Chromosome_5
Leptidea_sinapis_SWE_SWE96_196
Leptidea_sinapis_SWE_SWE96_195
Leptidea_sinapis_SWE_SWE96_194
Leptidea_sinapis_SWE_SWE96_193
Leptidea_sinapis_SWE_SWE96_192
Leptidea_sinapis_SWE_SWE96_191
Leptidea_sinapis_SWE_SWE96_190
Leptidea_sinapis_SWE_SWE96_189
Leptidea_sinapis_SWE_SWE96_188
Leptidea_sinapis_SWE_SWE96_187
Leptidea_sinapis_SWE_SWE96_185
Leptidea_sinapis_SWE_SWE96_184
Leptidea_sinapis_SWE_SWE96_183
Leptidea_sinapis_SWE_SWE96_182
Leptidea_sinapis_SWE_SWE96_181
Leptidea_sinapis_SWE_SWE96_180
Leptidea_sinapis_SWE_SWE96_179
Leptidea_sinapis_SWE_SWE96_178
Leptidea_sinapis_SWE_SWE96_177
Leptidea_sinapis_SWE_SWE96_176
Leptidea_sinapis_SWE_SWE96_175
Leptidea_sinapis_SWE_SWE96_174
Leptidea_sinapis_SWE_SWE96_173
Leptidea_sinapis_SWE_SWE96_172
Leptidea_sinapis_SWE_SWE96_171
Leptidea_sinapis_SWE_SWE96_170
Leptidea_sinapis_SWE_SWE96_169
Leptidea_sinapis_SWE_SWE96_168
Leptidea_sinapis_SWE_SWE96_167
Leptidea_sinapis_SWE_SWE96_166
Leptidea_sinapis_SWE_SWE96_165
Leptidea_sinapis_SWE_SWE96_164
Leptidea_sinapis_SWE_SWE96_163
Leptidea_sinapis_SWE_SWE96_162
Leptidea_sinapis_SWE_SWE96_161
Leptidea_sinapis_SWE_SWE96_160
Leptidea_sinapis_SWE_SWE96_159
Leptidea_sinapis_SWE_SWE96_158
Leptidea_sinapis_SWE_SWE96_157
Leptidea_sinapis_SWE_SWE96_156
Leptidea_sinapis_SWE_SWE96_155
Leptidea_sinapis_SWE_SWE96_154
Leptidea_sinapis_SWE_SWE96_153
Leptidea_sinapis_SWE_SWE96_152
Leptidea_sinapis_SWE_SWE96_151
Leptidea_sinapis_SWE_SWE96_150
Leptidea_sinapis_SWE_SWE96_149
Leptidea_sinapis_SWE_SWE96_148
Leptidea_sinapis_SWE_SWE96_147
Leptidea_sinapis_SWE_SWE96_146
Leptidea_sinapis_SWE_SWE96_145
Leptidea_sinapis_SWE_SWE96_144
Leptidea_sinapis_SWE_SWE96_143
Leptidea_sinapis_SWE_SWE96_142
Leptidea_sinapis_SWE_SWE96_141
Leptidea_sinapis_SWE_SWE96_140
Leptidea_sinapis_SWE_SWE96_139
Leptidea_sinapis_SWE_SWE96_138
Leptidea_sinapis_SWE_SWE96_137
Leptidea_sinapis_SWE_SWE96_136
Leptidea_sinapis_SWE_SWE96_135
Leptidea_sinapis_SWE_SWE96_134
Leptidea_sinapis_SWE_SWE96_131
Leptidea_sinapis_SWE_SWE96_130
Leptidea_sinapis_SWE_SWE96_129
Leptidea_sinapis_SWE_SWE96_123
Leptidea_sinapis_SWE_SWE96_122
Leptidea_sinapis_SWE_SWE96_118
Leptidea_sinapis_SWE_SWE96_116
Leptidea_sinapis_SWE_SWE96_115
Leptidea_sinapis_SWE_SWE96_114
Leptidea_sinapis_SWE_SWE96_113
Leptidea_sinapis_SWE_SWE96_112
Leptidea_sinapis_SWE_SWE96_111
Leptidea_sinapis_SWE_SWE96_110
Leptidea_sinapis_SWE_SWE96_109
Leptidea_sinapis_SWE_SWE96_108
Leptidea_sinapis_SWE_SWE96_107
Leptidea_sinapis_SWE_SWE96_106
Leptidea_sinapis_SWE_SWE96_105
Leptidea_sinapis_SWE_SWE96_104
Leptidea_sinapis_SWE_SWE96_103
Leptidea_sinapis_SWE_SWE96_102
Leptidea_sinapis_SWE_SWE96_101
Leptidea_sinapis_SWE_Lars_10X_104
Leptidea_sinapis_SWE_Lars_10X_103
Leptidea_sinapis_SWE_FD_SWE_Swe-sin-92C
Leptidea_sinapis_SWE_FD_SWE_Swe-sin-91C
Leptidea_sinapis_SWE_FD_SWE_Swe-sin-62C
Leptidea_sinapis_SWE_FD_SWE_Swe-sin-61C
Leptidea_sinapis_SWE_FD_SWE_Swe-sin-32C
Leptidea_sinapis_SWE_FD_SWE_Swe-sin-31C
Leptidea_sinapis_SWE_FD_SWE_Swe-sin-2C
Leptidea_sinapis_SWE_FD_SWE_Swe-sin-1C
Leptidea_sinapis_SWE_FD_SWE_Swe-sin-102C
Leptidea_sinapis_SWE_FD_SWE_Swe-sin-101C
Leptidea_sinapis_CAT_Lars_10X_106
Leptidea_sinapis_CAT_Lars_10X_105
Leptidea_sinapis_CAT_FD_CAT_47
Leptidea_sinapis_CAT_FD_CAT_46
Leptidea_sinapis_CAT_FD_CAT_44
Leptidea_sinapis_CAT_FD_CAT_43
Leptidea_sinapis_CAT_FD_CAT_42
Leptidea_sinapis_CAT_FD_CAT_41
Leptidea_sinapis_CAT_FD_CAT_40
Leptidea_sinapis_CAT_FD_CAT_39
Leptidea_sinapis_CAT_FD_CAT_38
Leptidea_sinapis_CAT_FD_CAT_37
Leptidea_reali_CAT_Lars_10X_116
Leptidea_reali_CAT_Lars_10X_115
Leptidea_reali_CAT_CATrea_36
Leptidea_reali_CAT_CATrea_35
Leptidea_reali_CAT_CATrea_34
Leptidea_reali_CAT_CATrea_33
Leptidea_reali_CAT_CATrea_32
Leptidea_reali_CAT_CATrea_31
Leptidea_reali_CAT_CATrea_30
Leptidea_reali_CAT_CATrea_29
Leptidea_reali_CAT_CATrea_27
Leptidea_reali_CAT_CATrea_26
Leptidea_morsei_CHI_LEP_outgroups_108
Leptidea_morsei_CHI_LEP_outgroups_107
Leptidea_lactea_CHI_LEP_outgroups_106
Leptidea_lactea_CHI_LEP_outgroups_105
Leptidea_juvernica_SWE_SWE96_186
Leptidea_juvernica_SWE_SWE96_133
Leptidea_juvernica_SWE_SWE96_132
Leptidea_juvernica_SWE_SWE96_128
Leptidea_juvernica_SWE_SWE96_127
Leptidea_juvernica_SWE_SWE96_126
Leptidea_juvernica_SWE_SWE96_125
Leptidea_juvernica_SWE_SWE96_124
Leptidea_juvernica_SWE_SWE96_121
Leptidea_juvernica_SWE_SWE96_120
Leptidea_juvernica_SWE_SWE96_119
Leptidea_juvernica_SWE_SWE96_117
Leptidea_juvernica_SWE_Lars_10X_112
Leptidea_juvernica_SWE_Lars_10X_111
Leptidea_amurensis_MON_LEP_outgroups_102
Leptidea_amurensis_MON_LEP_outgroups_101
Comb_ID_MSG
0.0e+00
5.0e+06
1.0e+07
1.5e+07
Start
Dosage
10.0
7.5
5.0
2.5
0.0

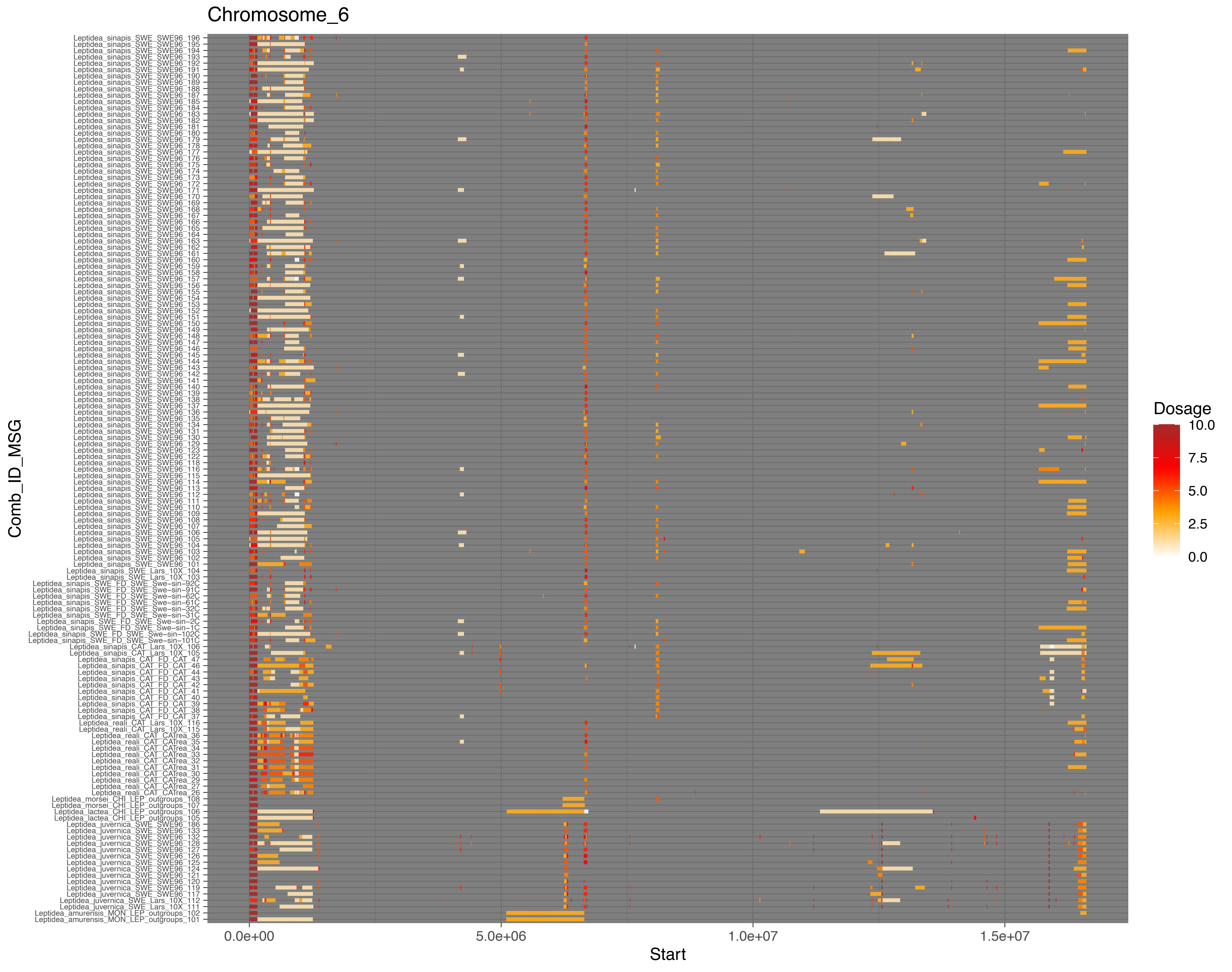
Chromosome_6
Leptidea_sinapis_SWE_SWE96_196
Leptidea_sinapis_SWE_SWE96_195
Leptidea_sinapis_SWE_SWE96_194
Leptidea_sinapis_SWE_SWE96_193
Leptidea_sinapis_SWE_SWE96_192
Leptidea_sinapis_SWE_SWE96_191
Leptidea_sinapis_SWE_SWE96_190
Leptidea_sinapis_SWE_SWE96_189
Leptidea_sinapis_SWE_SWE96_188
Leptidea_sinapis_SWE_SWE96_187
Leptidea_sinapis_SWE_SWE96_185
Leptidea_sinapis_SWE_SWE96_184
Leptidea_sinapis_SWE_SWE96_183
Leptidea_sinapis_SWE_SWE96_182
Leptidea_sinapis_SWE_SWE96_181
Leptidea_sinapis_SWE_SWE96_180
Leptidea_sinapis_SWE_SWE96_179
Leptidea_sinapis_SWE_SWE96_178
Leptidea_sinapis_SWE_SWE96_177
Leptidea_sinapis_SWE_SWE96_176
Leptidea_sinapis_SWE_SWE96_175
Leptidea_sinapis_SWE_SWE96_174
Leptidea_sinapis_SWE_SWE96_173
Leptidea_sinapis_SWE_SWE96_172
Leptidea_sinapis_SWE_SWE96_171
Leptidea_sinapis_SWE_SWE96_170
Leptidea_sinapis_SWE_SWE96_169
Leptidea_sinapis_SWE_SWE96_168
Leptidea_sinapis_SWE_SWE96_167
Leptidea_sinapis_SWE_SWE96_166
Leptidea_sinapis_SWE_SWE96_165
Leptidea_sinapis_SWE_SWE96_164
Leptidea_sinapis_SWE_SWE96_163
Leptidea_sinapis_SWE_SWE96_162
Leptidea_sinapis_SWE_SWE96_161
Leptidea_sinapis_SWE_SWE96_160
Leptidea_sinapis_SWE_SWE96_159
Leptidea_sinapis_SWE_SWE96_158
Leptidea_sinapis_SWE_SWE96_157
Leptidea_sinapis_SWE_SWE96_156
Leptidea_sinapis_SWE_SWE96_155
Leptidea_sinapis_SWE_SWE96_154
Leptidea_sinapis_SWE_SWE96_153
Leptidea_sinapis_SWE_SWE96_152
Leptidea_sinapis_SWE_SWE96_151
Leptidea_sinapis_SWE_SWE96_150
Leptidea_sinapis_SWE_SWE96_149
Leptidea_sinapis_SWE_SWE96_148
Leptidea_sinapis_SWE_SWE96_147
Leptidea_sinapis_SWE_SWE96_146
Leptidea_sinapis_SWE_SWE96_145
Leptidea_sinapis_SWE_SWE96_144
Leptidea_sinapis_SWE_SWE96_143
Leptidea_sinapis_SWE_SWE96_142
Leptidea_sinapis_SWE_SWE96_141
Leptidea_sinapis_SWE_SWE96_140
Leptidea_sinapis_SWE_SWE96_139
Leptidea_sinapis_SWE_SWE96_138
Leptidea_sinapis_SWE_SWE96_137
Leptidea_sinapis_SWE_SWE96_136
Leptidea_sinapis_SWE_SWE96_135
Leptidea_sinapis_SWE_SWE96_134
Leptidea_sinapis_SWE_SWE96_131
Leptidea_sinapis_SWE_SWE96_130
Leptidea_sinapis_SWE_SWE96_129
Leptidea_sinapis_SWE_SWE96_123
Leptidea_sinapis_SWE_SWE96_122
Leptidea_sinapis_SWE_SWE96_118
Leptidea_sinapis_SWE_SWE96_116
Leptidea_sinapis_SWE_SWE96_115
Leptidea_sinapis_SWE_SWE96_114
Leptidea_sinapis_SWE_SWE96_113
Leptidea_sinapis_SWE_SWE96_112
Leptidea_sinapis_SWE_SWE96_111
Leptidea_sinapis_SWE_SWE96_110
Leptidea_sinapis_SWE_SWE96_109
Leptidea_sinapis_SWE_SWE96_108
Leptidea_sinapis_SWE_SWE96_107
Leptidea_sinapis_SWE_SWE96_106
Leptidea_sinapis_SWE_SWE96_105
Leptidea_sinapis_SWE_SWE96_104
Leptidea_sinapis_SWE_SWE96_103
Leptidea_sinapis_SWE_SWE96_102
Leptidea_sinapis_SWE_SWE96_101
Leptidea_sinapis_SWE_Lars_10X_104
Leptidea_sinapis_SWE_Lars_10X_103
Leptidea_sinapis_SWE_FD_SWE_Swe−sin−92C
Leptidea_sinapis_SWE_FD_SWE_Swe−sin−91C
Leptidea_sinapis_SWE_FD_SWE_Swe−sin−62C
Leptidea_sinapis_SWE_FD_SWE_Swe−sin−61C
Leptidea_sinapis_SWE_FD_SWE_Swe−sin−32C
Leptidea_sinapis_SWE_FD_SWE_Swe−sin−31C
Leptidea_sinapis_SWE_FD_SWE_Swe−sin−2C
Leptidea_sinapis_SWE_FD_SWE_Swe−sin−1C
Leptidea_sinapis_SWE_FD_SWE_Swe−sin−102C
Leptidea_sinapis_SWE_FD_SWE_Swe−sin−101C
Leptidea_sinapis_CAT_Lars_10X_106
Leptidea_sinapis_CAT_Lars_10X_105
Leptidea_sinapis_CAT_FD_CAT_47
Leptidea_sinapis_CAT_FD_CAT_46
Leptidea_sinapis_CAT_FD_CAT_44
Leptidea_sinapis_CAT_FD_CAT_43
Leptidea_sinapis_CAT_FD_CAT_42
Leptidea_sinapis_CAT_FD_CAT_41
Leptidea_sinapis_CAT_FD_CAT_40
Leptidea_sinapis_CAT_FD_CAT_39
Leptidea_sinapis_CAT_FD_CAT_38
Leptidea_sinapis_CAT_FD_CAT_37
Leptidea_reali_CAT_Lars_10X_116
Leptidea_reali_CAT_Lars_10X_115
Leptidea_reali_CAT_CATrea_36
Leptidea_reali_CAT_CATrea_35
Leptidea_reali_CAT_CATrea_34
Leptidea_reali_CAT_CATrea_33
Leptidea_reali_CAT_CATrea_32
Leptidea_reali_CAT_CATrea_31
Leptidea_reali_CAT_CATrea_30
Leptidea_reali_CAT_CATrea_29
Leptidea_reali_CAT_CATrea_27
Leptidea_reali_CAT_CATrea_26
Leptidea_morsei_CHI_LEP_outgroups_108
Leptidea_morsei_CHI_LEP_outgroups_107
Leptidea_lactea_CHI_LEP_outgroups_106
Leptidea_lactea_CHI_LEP_outgroups_105
Leptidea_juvernica_SWE_SWE96_186
Leptidea_juvernica_SWE_SWE96_133
Leptidea_juvernica_SWE_SWE96_132
Leptidea_juvernica_SWE_SWE96_128
Leptidea_juvernica_SWE_SWE96_127
Leptidea_juvernica_SWE_SWE96_126
Leptidea_juvernica_SWE_SWE96_125
Leptidea_juvernica_SWE_SWE96_124
Leptidea_juvernica_SWE_SWE96_121
Leptidea_juvernica_SWE_SWE96_120
Leptidea_juvernica_SWE_SWE96_119
Leptidea_juvernica_SWE_SWE96_117
Leptidea_juvernica_SWE_Lars_10X_112
Leptidea_juvernica_SWE_Lars_10X_111
Leptidea_amurensis_MON_LEP_outgroups_102
Leptidea_amurensis_MON_LEP_outgroups_101
Comb_ID_MSG
0.0e+00
5.0e+06
1.0e+07
1.5e+07
Start
Dosage
10.0
7.5
5.0
2.5
0.0

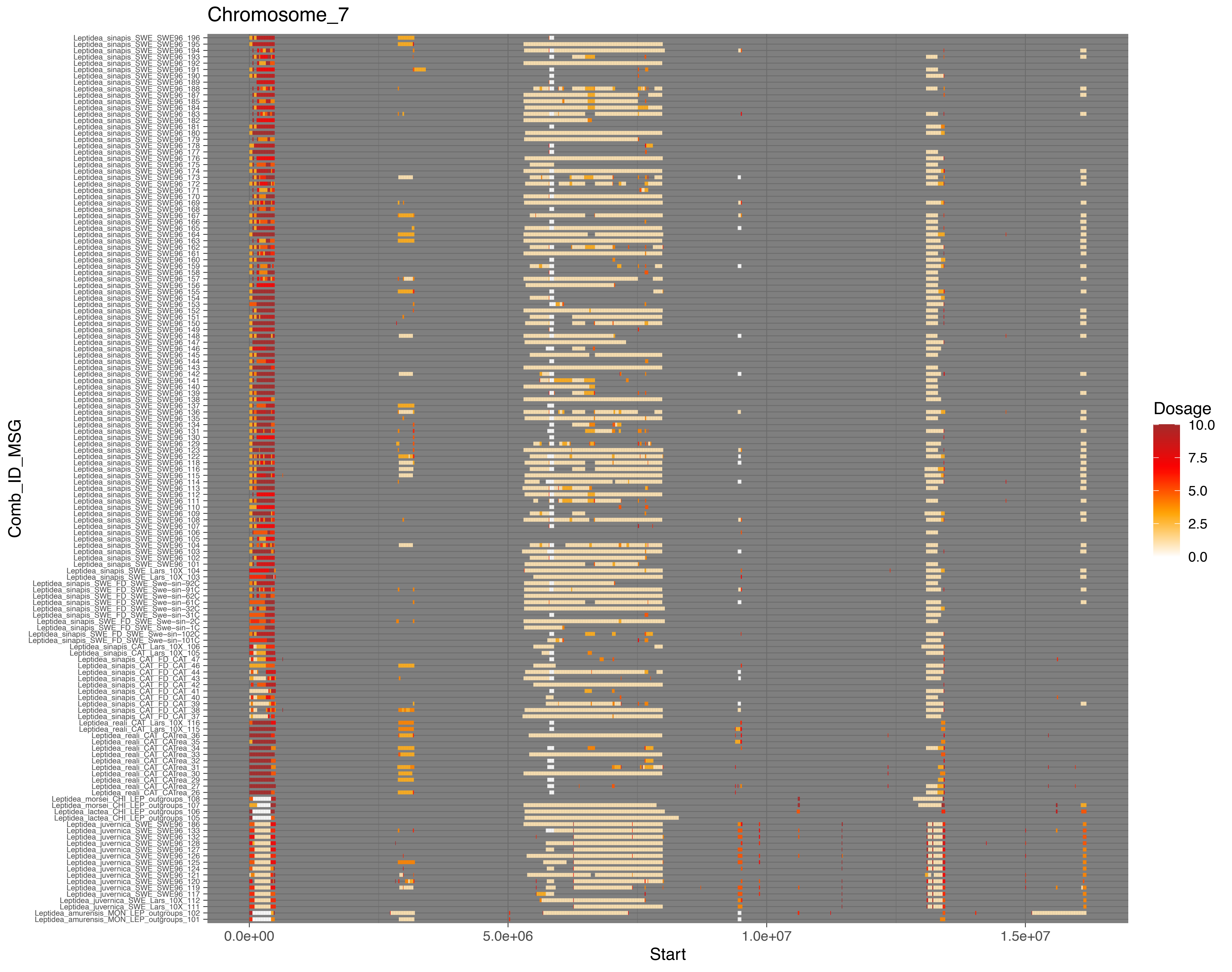
Chromosome_7
Comb_ID_MSG
Start
0.0e+00
5.0e+06
1.0e+07
1.5e+07
Dosage
10.0
7.5
5.0
2.5
0.0

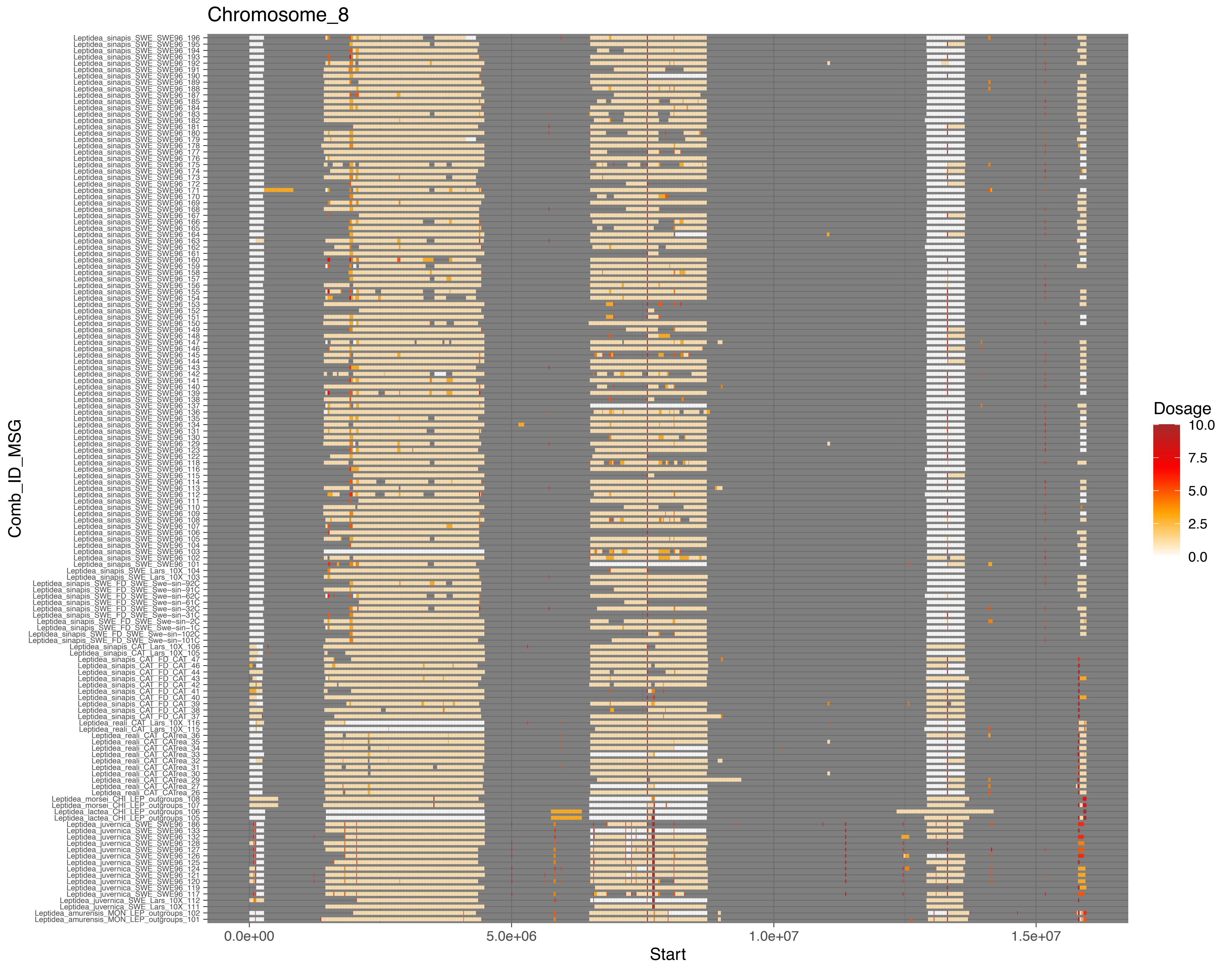
Chromosome_8
Comb_ID_MSG
Start
0.0e+00
5.0e+06
1.0e+07
1.5e+07
Dosage
10.0
7.5
5.0
2.5
0.0

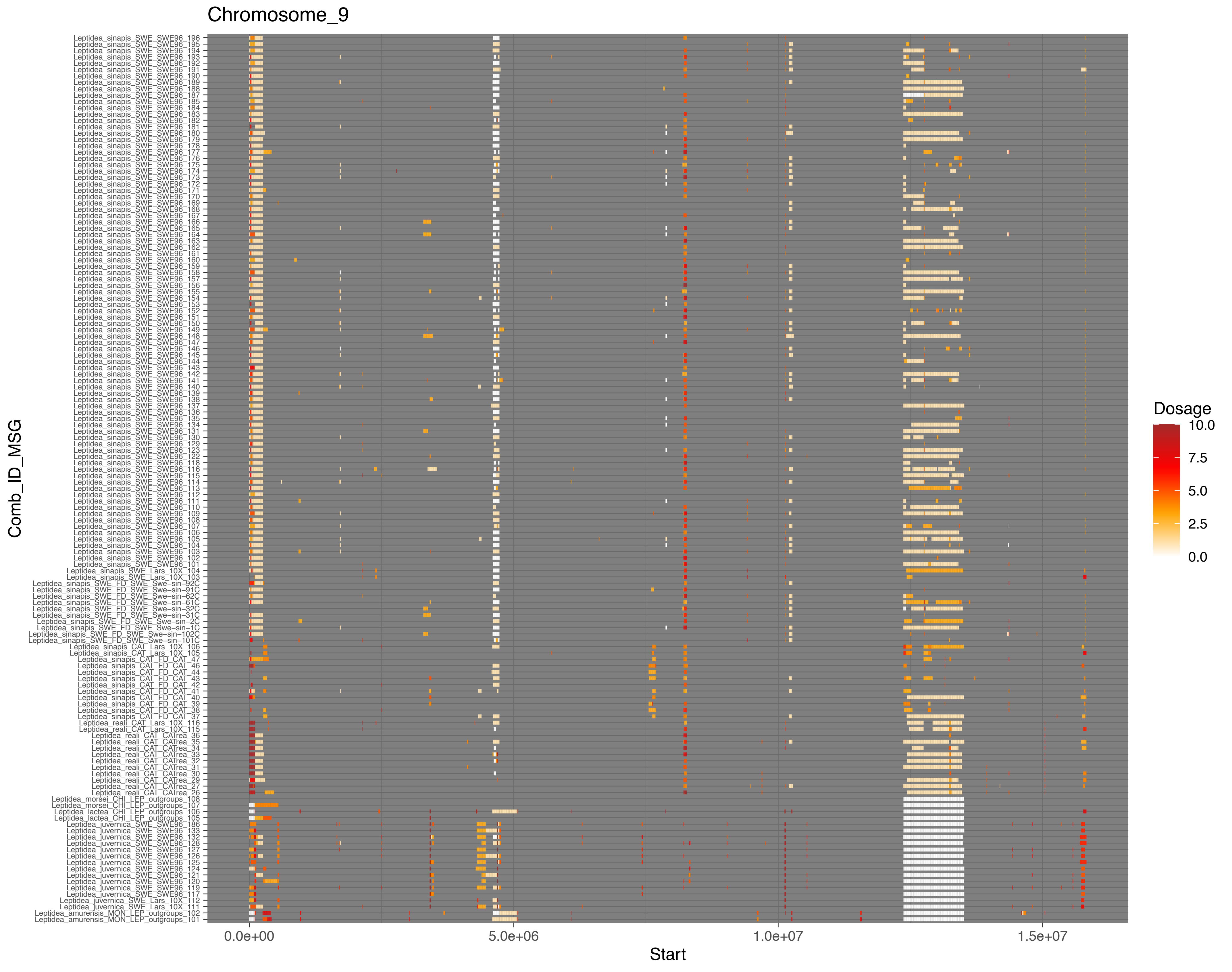
Chromosome_9
Comb_ID_MSG
Start
0.0e+00
5.0e+06
1.0e+07
1.5e+07
Dosage
10.0
7.5
5.0
2.5
0.0

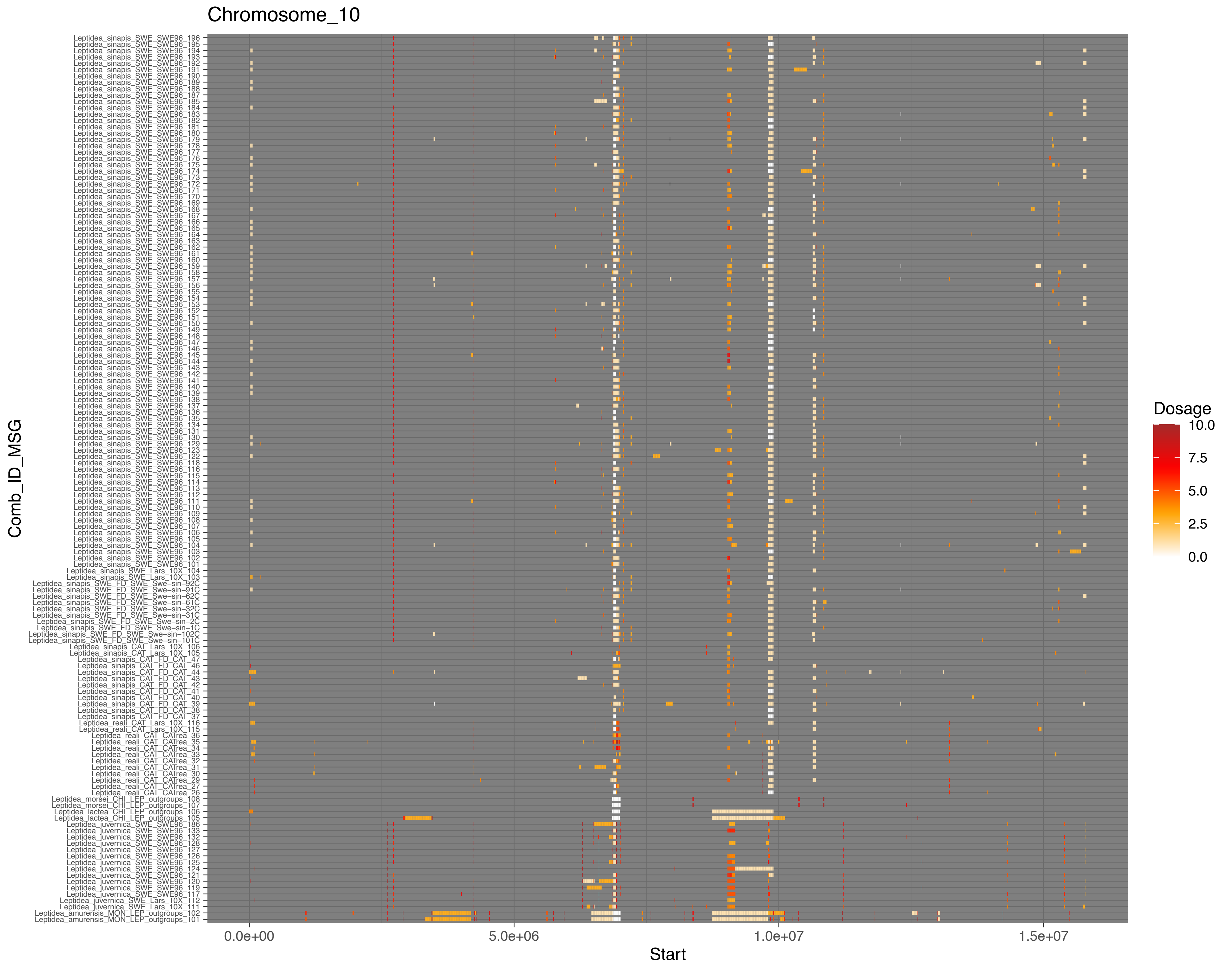

Chromosome_10
Comb_ID_MSG
Leptidea_sinapis_SWE_SWE96_196
Leptidea_sinapis_SWE_SWE96_195
Leptidea_sinapis_SWE_SWE96_194
Leptidea_sinapis_SWE_SWE96_193
Leptidea_sinapis_SWE_SWE96_192
Leptidea_sinapis_SWE_SWE96_191
Leptidea_sinapis_SWE_SWE96_190
Leptidea_sinapis_SWE_SWE96_189
Leptidea_sinapis_SWE_SWE96_188
Leptidea_sinapis_SWE_SWE96_187
Leptidea_sinapis_SWE_SWE96_185
Leptidea_sinapis_SWE_SWE96_184
Leptidea_sinapis_SWE_SWE96_183
Leptidea_sinapis_SWE_SWE96_182
Leptidea_sinapis_SWE_SWE96_181
Leptidea_sinapis_SWE_SWE96_180
Leptidea_sinapis_SWE_SWE96_179
Leptidea_sinapis_SWE_SWE96_178
Leptidea_sinapis_SWE_SWE96_177
Leptidea_sinapis_SWE_SWE96_176
Leptidea_sinapis_SWE_SWE96_175
Leptidea_sinapis_SWE_SWE96_174
Leptidea_sinapis_SWE_SWE96_173
Leptidea_sinapis_SWE_SWE96_172
Leptidea_sinapis_SWE_SWE96_171
Leptidea_sinapis_SWE_SWE96_170
Leptidea_sinapis_SWE_SWE96_169
Leptidea_sinapis_SWE_SWE96_168
Leptidea_sinapis_SWE_SWE96_167
Leptidea_sinapis_SWE_SWE96_166
Leptidea_sinapis_SWE_SWE96_165
Leptidea_sinapis_SWE_SWE96_164
Leptidea_sinapis_SWE_SWE96_163
Leptidea_sinapis_SWE_SWE96_162
Leptidea_sinapis_SWE_SWE96_161
Leptidea_sinapis_SWE_SWE96_160
Leptidea_sinapis_SWE_SWE96_159
Leptidea_sinapis_SWE_SWE96_158
Leptidea_sinapis_SWE_SWE96_157
Leptidea_sinapis_SWE_SWE96_156
Leptidea_sinapis_SWE_SWE96_155
Leptidea_sinapis_SWE_SWE96_154
Leptidea_sinapis_SWE_SWE96_153
Leptidea_sinapis_SWE_SWE96_152
Leptidea_sinapis_SWE_SWE96_151
Leptidea_sinapis_SWE_SWE96_150
Leptidea_sinapis_SWE_SWE96_149
Leptidea_sinapis_SWE_SWE96_148
Leptidea_sinapis_SWE_SWE96_147
Leptidea_sinapis_SWE_SWE96_146
Leptidea_sinapis_SWE_SWE96_145
Leptidea_sinapis_SWE_SWE96_144
Leptidea_sinapis_SWE_SWE96_143
Leptidea_sinapis_SWE_SWE96_142
Leptidea_sinapis_SWE_SWE96_141
Leptidea_sinapis_SWE_SWE96_140
Leptidea_sinapis_SWE_SWE96_139
Leptidea_sinapis_SWE_SWE96_138
Leptidea_sinapis_SWE_SWE96_137
Leptidea_sinapis_SWE_SWE96_136
Leptidea_sinapis_SWE_SWE96_135
Leptidea_sinapis_SWE_SWE96_134
Leptidea_sinapis_SWE_SWE96_131
Leptidea_sinapis_SWE_SWE96_130
Leptidea_sinapis_SWE_SWE96_129
Leptidea_sinapis_SWE_SWE96_123
Leptidea_sinapis_SWE_SWE96_122
Leptidea_sinapis_SWE_SWE96_118
Leptidea_sinapis_SWE_SWE96_116
Leptidea_sinapis_SWE_SWE96_115
Leptidea_sinapis_SWE_SWE96_114
Leptidea_sinapis_SWE_SWE96_113
Leptidea_sinapis_SWE_SWE96_112
Leptidea_sinapis_SWE_SWE96_111
Leptidea_sinapis_SWE_SWE96_110
Leptidea_sinapis_SWE_SWE96_109
Leptidea_sinapis_SWE_SWE96_108
Leptidea_sinapis_SWE_SWE96_107
Leptidea_sinapis_SWE_SWE96_106
Leptidea_sinapis_SWE_SWE96_105
Leptidea_sinapis_SWE_SWE96_104
Leptidea_sinapis_SWE_SWE96_103
Leptidea_sinapis_SWE_SWE96_102
Leptidea_sinapis_SWE_SWE96_101
Leptidea_sinapis_SWE_Lars_10X_104
Leptidea_sinapis_SWE_Lars_10X_103
Leptidea_sinapis_SWE_FD_SWE_Swe-sin-92C
Leptidea_sinapis_SWE_FD_SWE_Swe-sin-91C
Leptidea_sinapis_SWE_FD_SWE_Swe-sin-62C
Leptidea_sinapis_SWE_FD_SWE_Swe-sin-61C
Leptidea_sinapis_SWE_FD_SWE_Swe-sin-32C
Leptidea_sinapis_SWE_FD_SWE_Swe-sin-31C
Leptidea_sinapis_SWE_FD_SWE_Swe-sin-2C
Leptidea_sinapis_SWE_FD_SWE_Swe-sin-1C
Leptidea_sinapis_SWE_FD_SWE_Swe-sin-102C
Leptidea_sinapis_SWE_FD_SWE_Swe-sin-101C
Leptidea_sinapis_CAT_Lars_10X_106
Leptidea_sinapis_CAT_Lars_10X_105
Leptidea_sinapis_CAT_FD_CAT_47
Leptidea_sinapis_CAT_FD_CAT_46
Leptidea_sinapis_CAT_FD_CAT_44
Leptidea_sinapis_CAT_FD_CAT_43
Leptidea_sinapis_CAT_FD_CAT_42
Leptidea_sinapis_CAT_FD_CAT_41
Leptidea_sinapis_CAT_FD_CAT_40
Leptidea_sinapis_CAT_FD_CAT_39
Leptidea_sinapis_CAT_FD_CAT_38
Leptidea_sinapis_CAT_FD_CAT_37
Leptidea_reali_CAT_Lars_10X_116
Leptidea_reali_CAT_Lars_10X_115
Leptidea_reali_CAT_CATrea_36
Leptidea_reali_CAT_CATrea_35
Leptidea_reali_CAT_CATrea_34
Leptidea_reali_CAT_CATrea_33
Leptidea_reali_CAT_CATrea_32
Leptidea_reali_CAT_CATrea_31
Leptidea_reali_CAT_CATrea_30
Leptidea_reali_CAT_CATrea_29
Leptidea_reali_CAT_CATrea_27
Leptidea_reali_CAT_CATrea_26
Leptidea_morsei_CHI_LEP_outgroups_108
Leptidea_morsei_CHI_LEP_outgroups_107
Leptidea_lactea_CHI_LEP_outgroups_106
Leptidea_lactea_CHI_LEP_outgroups_105
Leptidea_juvernica_SWE_SWE96_186
Leptidea_juvernica_SWE_SWE96_133
Leptidea_juvernica_SWE_SWE96_132
Leptidea_juvernica_SWE_SWE96_128
Leptidea_juvernica_SWE_SWE96_127
Leptidea_juvernica_SWE_SWE96_126
Leptidea_juvernica_SWE_SWE96_125
Leptidea_juvernica_SWE_SWE96_124
Leptidea_juvernica_SWE_SWE96_121
Leptidea_juvernica_SWE_SWE96_120
Leptidea_juvernica_SWE_SWE96_119
Leptidea_juvernica_SWE_SWE96_117
Leptidea_juvernica_SWE_Lars_10X_112
Leptidea_juvernica_SWE_Lars_10X_111
Leptidea_amurensis_MON_LEP_outgroups_102
Leptidea_amurensis_MON_LEP_outgroups_101
0.0e+00
5.0e+06
1.0e+07
1.5e+07
Start
Dosage
10.0
7.5
5.0
2.5
0.0

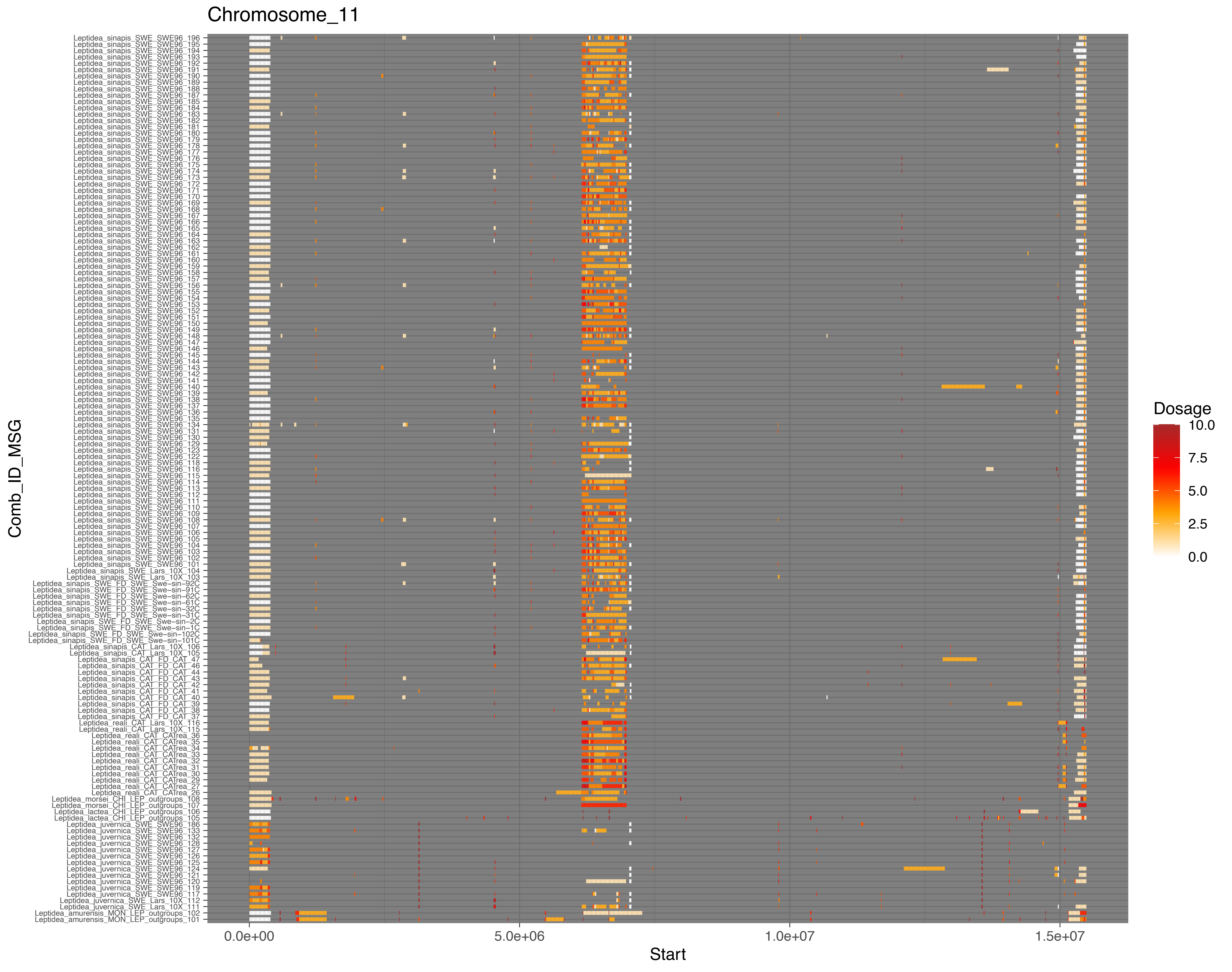

Chromosome_11
Comb_ID_MSG
Start
0.0e+00
5.0e+06
1.0e+07
1.5e+07
Dosage
10.0
7.5
5.0
2.5
0.0

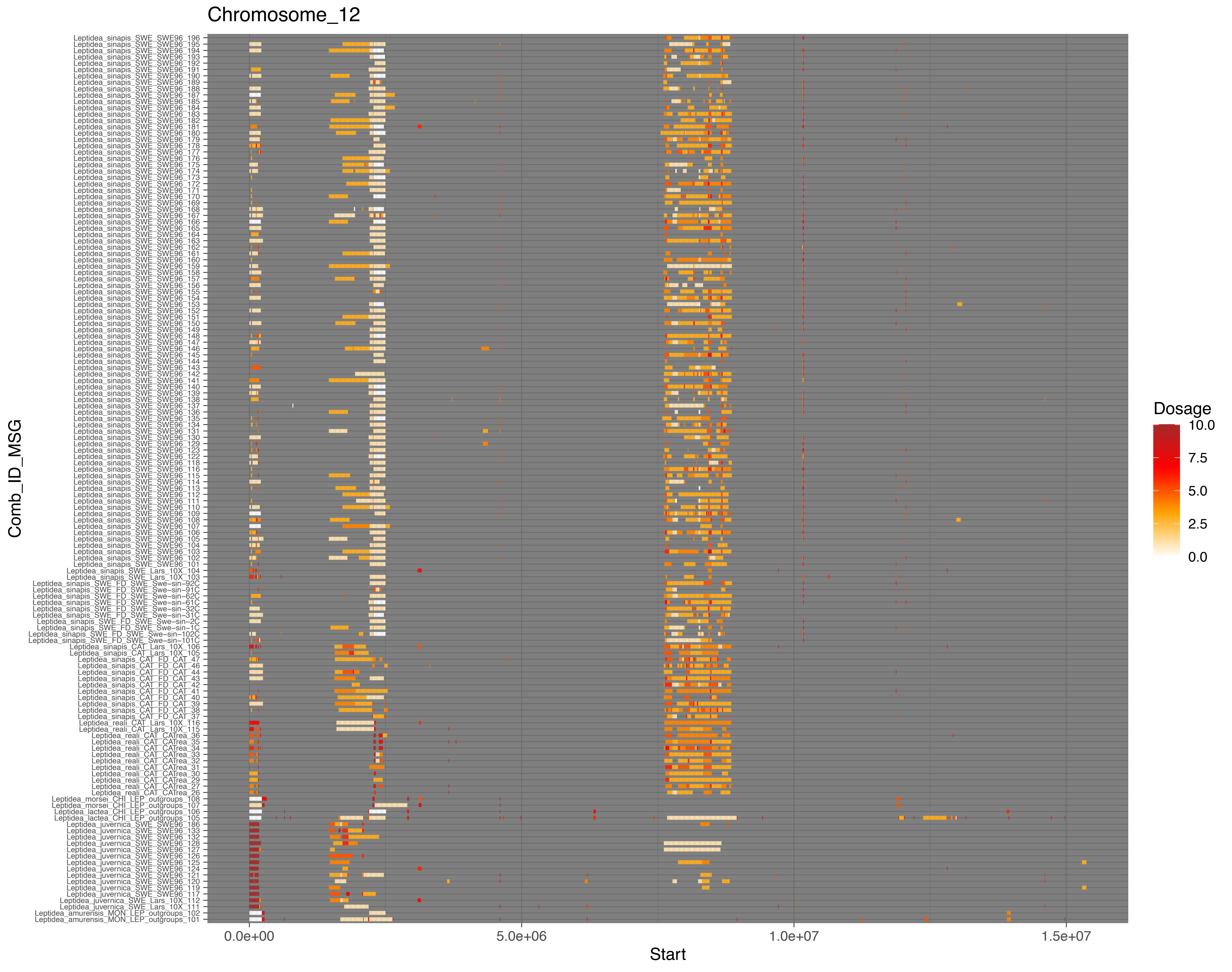
Chromosome_12
Comb_ID_MSG
Start
0.0e+00
5.0e+06
1.0e+07
1.5e+07
Dosage
10.0
7.5
5.0
2.5
0.0

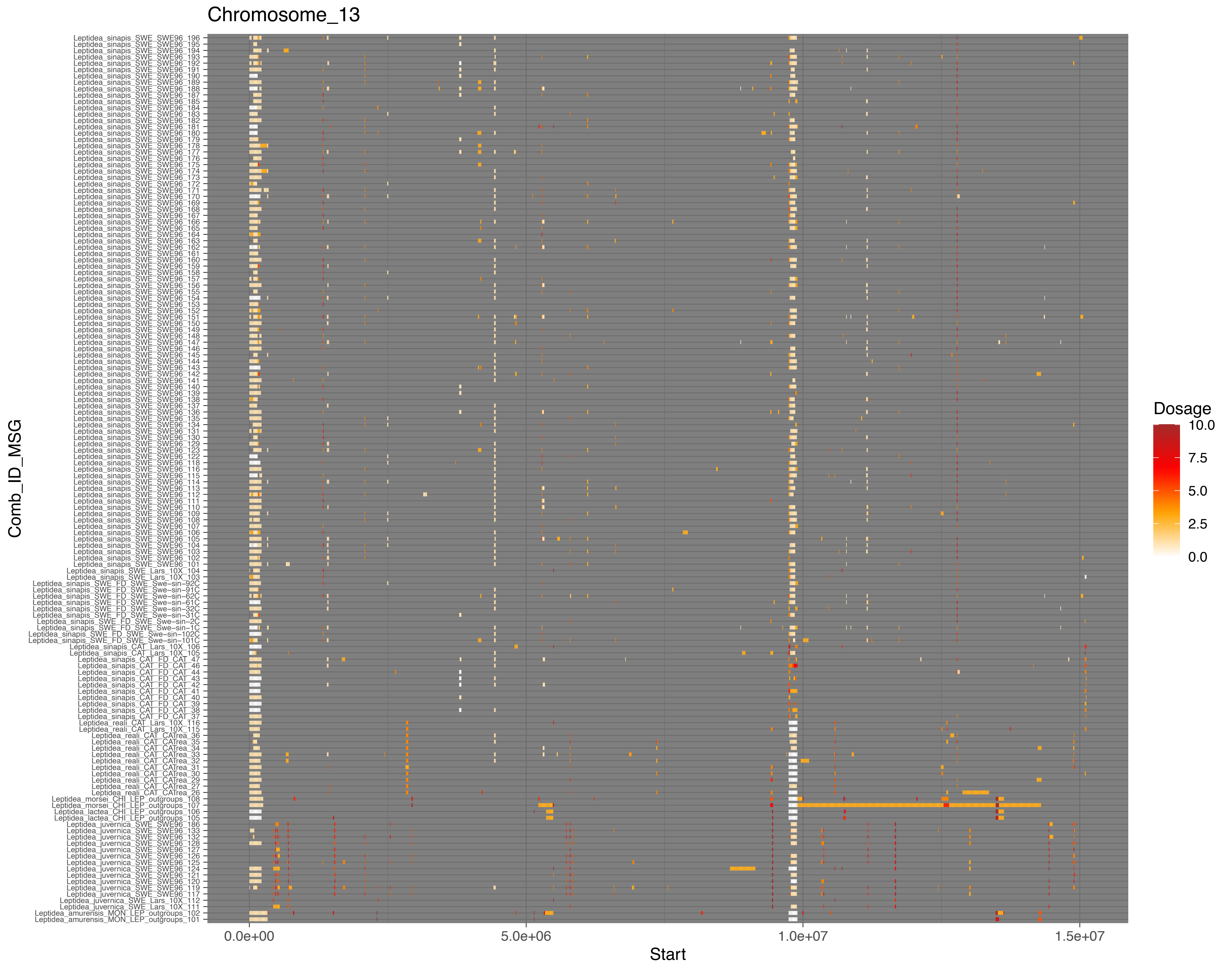
Chromosome_13
Comb_ID_MSG
Leptidea_sinapis_SWE_SWE96_196
Leptidea_sinapis_SWE_SWE96_195
Leptidea_sinapis_SWE_SWE96_194
Leptidea_sinapis_SWE_SWE96_193
Leptidea_sinapis_SWE_SWE96_192
Leptidea_sinapis_SWE_SWE96_191
Leptidea_sinapis_SWE_SWE96_190
Leptidea_sinapis_SWE_SWE96_189
Leptidea_sinapis_SWE_SWE96_188
Leptidea_sinapis_SWE_SWE96_187
Leptidea_sinapis_SWE_SWE96_185
Leptidea_sinapis_SWE_SWE96_184
Leptidea_sinapis_SWE_SWE96_183
Leptidea_sinapis_SWE_SWE96_182
Leptidea_sinapis_SWE_SWE96_181
Leptidea_sinapis_SWE_SWE96_180
Leptidea_sinapis_SWE_SWE96_179
Leptidea_sinapis_SWE_SWE96_178
Leptidea_sinapis_SWE_SWE96_177
Leptidea_sinapis_SWE_SWE96_176
Leptidea_sinapis_SWE_SWE96_175
Leptidea_sinapis_SWE_SWE96_174
Leptidea_sinapis_SWE_SWE96_173
Leptidea_sinapis_SWE_SWE96_172
Leptidea_sinapis_SWE_SWE96_171
Leptidea_sinapis_SWE_SWE96_170
Leptidea_sinapis_SWE_SWE96_169
Leptidea_sinapis_SWE_SWE96_168
Leptidea_sinapis_SWE_SWE96_167
Leptidea_sinapis_SWE_SWE96_166
Leptidea_sinapis_SWE_SWE96_165
Leptidea_sinapis_SWE_SWE96_164
Leptidea_sinapis_SWE_SWE96_163
Leptidea_sinapis_SWE_SWE96_162
Leptidea_sinapis_SWE_SWE96_161
Leptidea_sinapis_SWE_SWE96_160
Leptidea_sinapis_SWE_SWE96_159
Leptidea_sinapis_SWE_SWE96_158
Leptidea_sinapis_SWE_SWE96_157
Leptidea_sinapis_SWE_SWE96_156
Leptidea_sinapis_SWE_SWE96_155
Leptidea_sinapis_SWE_SWE96_154
Leptidea_sinapis_SWE_SWE96_153
Leptidea_sinapis_SWE_SWE96_152
Leptidea_sinapis_SWE_SWE96_151
Leptidea_sinapis_SWE_SWE96_150
Leptidea_sinapis_SWE_SWE96_149
Leptidea_sinapis_SWE_SWE96_148
Leptidea_sinapis_SWE_SWE96_147
Leptidea_sinapis_SWE_SWE96_146
Leptidea_sinapis_SWE_SWE96_145
Leptidea_sinapis_SWE_SWE96_144
Leptidea_sinapis_SWE_SWE96_143
Leptidea_sinapis_SWE_SWE96_142
Leptidea_sinapis_SWE_SWE96_141
Leptidea_sinapis_SWE_SWE96_140
Leptidea_sinapis_SWE_SWE96_139
Leptidea_sinapis_SWE_SWE96_138
Leptidea_sinapis_SWE_SWE96_137
Leptidea_sinapis_SWE_SWE96_136
Leptidea_sinapis_SWE_SWE96_135
Leptidea_sinapis_SWE_SWE96_134
Leptidea_sinapis_SWE_SWE96_131
Leptidea_sinapis_SWE_SWE96_130
Leptidea_sinapis_SWE_SWE96_129
Leptidea_sinapis_SWE_SWE96_123
Leptidea_sinapis_SWE_SWE96_122
Leptidea_sinapis_SWE_SWE96_118
Leptidea_sinapis_SWE_SWE96_116
Leptidea_sinapis_SWE_SWE96_115
Leptidea_sinapis_SWE_SWE96_114
Leptidea_sinapis_SWE_SWE96_113
Leptidea_sinapis_SWE_SWE96_112
Leptidea_sinapis_SWE_SWE96_111
Leptidea_sinapis_SWE_SWE96_110
Leptidea_sinapis_SWE_SWE96_109
Leptidea_sinapis_SWE_SWE96_108
Leptidea_sinapis_SWE_SWE96_107
Leptidea_sinapis_SWE_SWE96_106
Leptidea_sinapis_SWE_SWE96_105
Leptidea_sinapis_SWE_SWE96_104
Leptidea_sinapis_SWE_SWE96_103
Leptidea_sinapis_SWE_SWE96_102
Leptidea_sinapis_SWE_SWE96_101
Leptidea_sinapis_SWE_Lars_10X_104
Leptidea_sinapis_SWE_Lars_10X_103
Leptidea_sinapis_SWE_FD_SWE_Swe-sin-92C
Leptidea_sinapis_SWE_FD_SWE_Swe-sin-91C
Leptidea_sinapis_SWE_FD_SWE_Swe-sin-62C
Leptidea_sinapis_SWE_FD_SWE_Swe-sin-61C
Leptidea_sinapis_SWE_FD_SWE_Swe-sin-32C
Leptidea_sinapis_SWE_FD_SWE_Swe-sin-31C
Leptidea_sinapis_SWE_FD_SWE_Swe-sin-2C
Leptidea_sinapis_SWE_FD_SWE_Swe-sin-1C
Leptidea_sinapis_SWE_FD_SWE_Swe-sin-102C
Leptidea_sinapis_SWE_FD_SWE_Swe-sin-101C
Leptidea_sinapis_CAT_Lars_10X_106
Leptidea_sinapis_CAT_Lars_10X_105
Leptidea_sinapis_CAT_FD_CAT_47
Leptidea_sinapis_CAT_FD_CAT_46
Leptidea_sinapis_CAT_FD_CAT_44
Leptidea_sinapis_CAT_FD_CAT_43
Leptidea_sinapis_CAT_FD_CAT_42
Leptidea_sinapis_CAT_FD_CAT_41
Leptidea_sinapis_CAT_FD_CAT_40
Leptidea_sinapis_CAT_FD_CAT_39
Leptidea_sinapis_CAT_FD_CAT_38
Leptidea_sinapis_CAT_FD_CAT_37
Leptidea_reali_CAT_Lars_10X_116
Leptidea_reali_CAT_Lars_10X_115
Leptidea_reali_CAT_CATrea_36
Leptidea_reali_CAT_CATrea_35
Leptidea_reali_CAT_CATrea_34
Leptidea_reali_CAT_CATrea_33
Leptidea_reali_CAT_CATrea_32
Leptidea_reali_CAT_CATrea_31
Leptidea_reali_CAT_CATrea_30
Leptidea_reali_CAT_CATrea_29
Leptidea_reali_CAT_CATrea_27
Leptidea_reali_CAT_CATrea_26
Leptidea_morsei_CHI_LEP_outgroups_108
Leptidea_morsei_CHI_LEP_outgroups_107
Leptidea_lactea_CHI_LEP_outgroups_106
Leptidea_lactea_CHI_LEP_outgroups_105
Leptidea_juvernica_SWE_SWE96_186
Leptidea_juvernica_SWE_SWE96_133
Leptidea_juvernica_SWE_SWE96_132
Leptidea_juvernica_SWE_SWE96_128
Leptidea_juvernica_SWE_SWE96_127
Leptidea_juvernica_SWE_SWE96_126
Leptidea_juvernica_SWE_SWE96_125
Leptidea_juvernica_SWE_SWE96_124
Leptidea_juvernica_SWE_SWE96_121
Leptidea_juvernica_SWE_SWE96_120
Leptidea_juvernica_SWE_SWE96_119
Leptidea_juvernica_SWE_SWE96_117
Leptidea_juvernica_SWE_Lars_10X_112
Leptidea_juvernica_SWE_Lars_10X_111
Leptidea_amurensis_MON_LEP_outgroups_102
Leptidea_amurensis_MON_LEP_outgroups_101
0.0e+00
5.0e+06
1.0e+07
1.5e+07
Start
Dosage
10.0
7.5
5.0
2.5
0.0

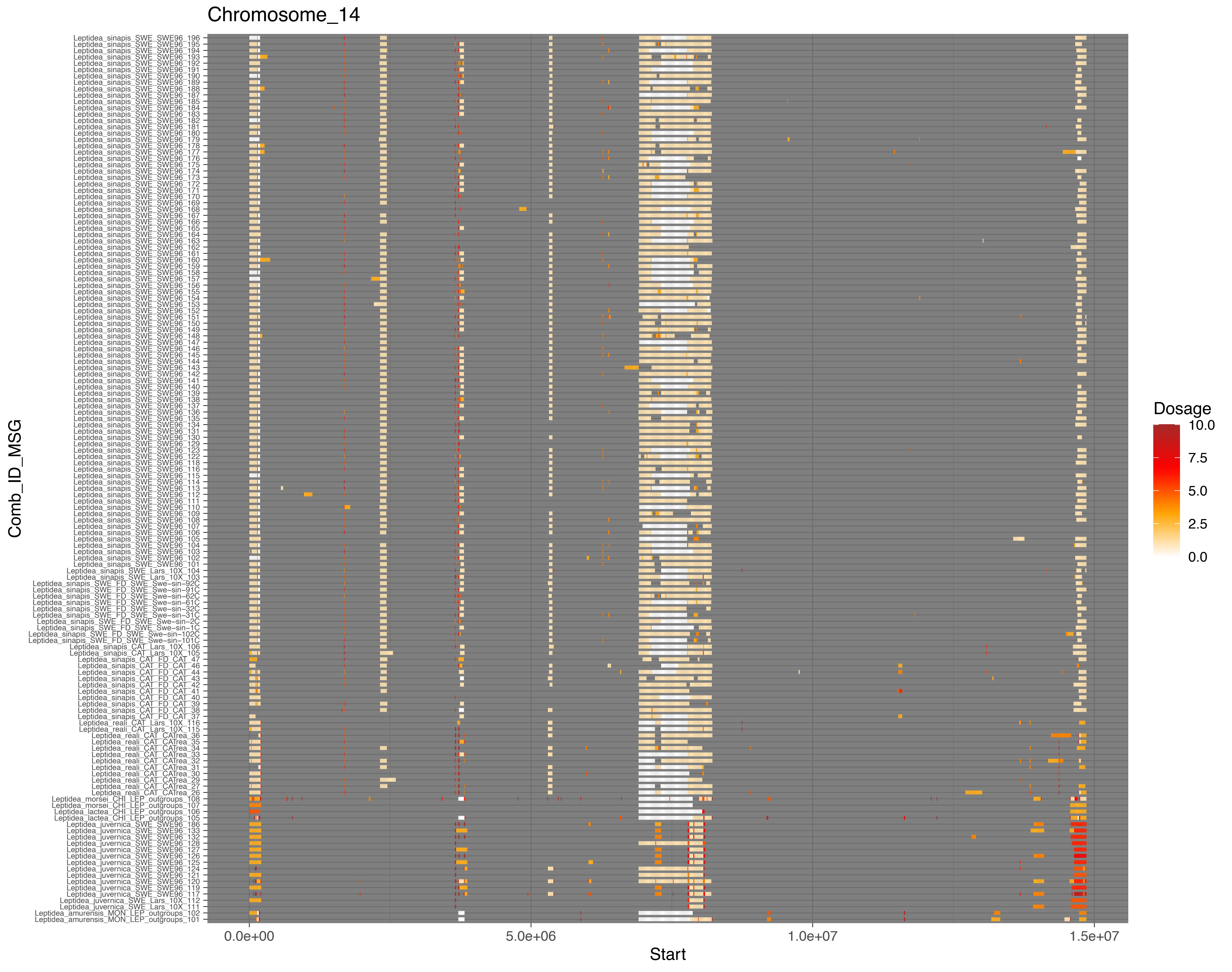
Chromosome_14
Comb_ID_MSG
Leptidea_sinapis_SWE_SWE96_196
Leptidea_sinapis_SWE_SWE96_195
Leptidea_sinapis_SWE_SWE96_194
Leptidea_sinapis_SWE_SWE96_193
Leptidea_sinapis_SWE_SWE96_192
Leptidea_sinapis_SWE_SWE96_191
Leptidea_sinapis_SWE_SWE96_190
Leptidea_sinapis_SWE_SWE96_189
Leptidea_sinapis_SWE_SWE96_188
Leptidea_sinapis_SWE_SWE96_187
Leptidea_sinapis_SWE_SWE96_185
Leptidea_sinapis_SWE_SWE96_184
Leptidea_sinapis_SWE_SWE96_183
Leptidea_sinapis_SWE_SWE96_182
Leptidea_sinapis_SWE_SWE96_181
Leptidea_sinapis_SWE_SWE96_180
Leptidea_sinapis_SWE_SWE96_179
Leptidea_sinapis_SWE_SWE96_178
Leptidea_sinapis_SWE_SWE96_177
Leptidea_sinapis_SWE_SWE96_176
Leptidea_sinapis_SWE_SWE96_175
Leptidea_sinapis_SWE_SWE96_174
Leptidea_sinapis_SWE_SWE96_173
Leptidea_sinapis_SWE_SWE96_172
Leptidea_sinapis_SWE_SWE96_171
Leptidea_sinapis_SWE_SWE96_170
Leptidea_sinapis_SWE_SWE96_169
Leptidea_sinapis_SWE_SWE96_168
Leptidea_sinapis_SWE_SWE96_167
Leptidea_sinapis_SWE_SWE96_166
Leptidea_sinapis_SWE_SWE96_165
Leptidea_sinapis_SWE_SWE96_164
Leptidea_sinapis_SWE_SWE96_163
Leptidea_sinapis_SWE_SWE96_162
Leptidea_sinapis_SWE_SWE96_161
Leptidea_sinapis_SWE_SWE96_160
Leptidea_sinapis_SWE_SWE96_159
Leptidea_sinapis_SWE_SWE96_158
Leptidea_sinapis_SWE_SWE96_157
Leptidea_sinapis_SWE_SWE96_156
Leptidea_sinapis_SWE_SWE96_155
Leptidea_sinapis_SWE_SWE96_154
Leptidea_sinapis_SWE_SWE96_153
Leptidea_sinapis_SWE_SWE96_152
Leptidea_sinapis_SWE_SWE96_151
Leptidea_sinapis_SWE_SWE96_150
Leptidea_sinapis_SWE_SWE96_149
Leptidea_sinapis_SWE_SWE96_148
Leptidea_sinapis_SWE_SWE96_147
Leptidea_sinapis_SWE_SWE96_146
Leptidea_sinapis_SWE_SWE96_145
Leptidea_sinapis_SWE_SWE96_144
Leptidea_sinapis_SWE_SWE96_143
Leptidea_sinapis_SWE_SWE96_142
Leptidea_sinapis_SWE_SWE96_141
Leptidea_sinapis_SWE_SWE96_140
Leptidea_sinapis_SWE_SWE96_139
Leptidea_sinapis_SWE_SWE96_138
Leptidea_sinapis_SWE_SWE96_137
Leptidea_sinapis_SWE_SWE96_136
Leptidea_sinapis_SWE_SWE96_135
Leptidea_sinapis_SWE_SWE96_134
Leptidea_sinapis_SWE_SWE96_131
Leptidea_sinapis_SWE_SWE96_130
Leptidea_sinapis_SWE_SWE96_129
Leptidea_sinapis_SWE_SWE96_123
Leptidea_sinapis_SWE_SWE96_122
Leptidea_sinapis_SWE_SWE96_118
Leptidea_sinapis_SWE_SWE96_116
Leptidea_sinapis_SWE_SWE96_115
Leptidea_sinapis_SWE_SWE96_114
Leptidea_sinapis_SWE_SWE96_113
Leptidea_sinapis_SWE_SWE96_112
Leptidea_sinapis_SWE_SWE96_110
Leptidea_sinapis_SWE_SWE96_111
Leptidea_sinapis_SWE_SWE96_109
Leptidea_sinapis_SWE_SWE96_108
Leptidea_sinapis_SWE_SWE96_107
Leptidea_sinapis_SWE_SWE96_106
Leptidea_sinapis_SWE_SWE96_105
Leptidea_sinapis_SWE_SWE96_104
Leptidea_sinapis_SWE_SWE96_103
Leptidea_sinapis_SWE_SWE96_102
Leptidea_sinapis_SWE_SWE96_101
Leptidea_sinapis_SWE_Lars_10X_104
Leptidea_sinapis_SWE_Lars_10X_103
Leptidea_sinapis_SWE_FD_SWE_Swe-sin-92C
Leptidea_sinapis_SWE_FD_SWE_Swe-sin-91C
Leptidea_sinapis_SWE_FD_SWE_Swe-sin-62C
Leptidea_sinapis_SWE_FD_SWE_Swe-sin-61C
Leptidea_sinapis_SWE_FD_SWE_Swe-sin-32C
Leptidea_sinapis_SWE_FD_SWE_Swe-sin-31C
Leptidea_sinapis_SWE_FD_SWE_Swe-sin-2C
Leptidea_sinapis_SWE_FD_SWE_Swe-sin-1C
Leptidea_sinapis_SWE_FD_SWE_Swe-sin-102C
Leptidea_sinapis_SWE_FD_SWE_Swe-sin-101C
Leptidea_sinapis_CAT_Lars_10X_106
Leptidea_sinapis_CAT_Lars_10X_105
Leptidea_sinapis_CAT_FD_CAT_47
Leptidea_sinapis_CAT_FD_CAT_46
Leptidea_sinapis_CAT_FD_CAT_44
Leptidea_sinapis_CAT_FD_CAT_43
Leptidea_sinapis_CAT_FD_CAT_42
Leptidea_sinapis_CAT_FD_CAT_41
Leptidea_sinapis_CAT_FD_CAT_40
Leptidea_sinapis_CAT_FD_CAT_39
Leptidea_sinapis_CAT_FD_CAT_38
Leptidea_sinapis_CAT_FD_CAT_37
Leptidea_reali_CAT_Lars_10X_116
Leptidea_reali_CAT_Lars_10X_115
Leptidea_reali_CAT_CAtrea_36
Leptidea_reali_CAT_CAtrea_35
Leptidea_reali_CAT_CAtrea_34
Leptidea_reali_CAT_CAtrea_33
Leptidea_reali_CAT_CAtrea_32
Leptidea_reali_CAT_CAtrea_31
Leptidea_reali_CAT_CAtrea_30
Leptidea_reali_CAT_CAtrea_29
Leptidea_reali_CAT_CAtrea_27
Leptidea_reali_CAT_CAtrea_26
Leptidea_morsei_CHI_LEP_outgroups_108
Leptidea_morsei_CHI_LEP_outgroups_107
Leptidea_lactea_CHI_LEP_outgroups_106
Leptidea_lactea_CHI_LEP_outgroups_105
Leptidea_juvernica_SWE_SWE96_186
Leptidea_juvernica_SWE_SWE96_133
Leptidea_juvernica_SWE_SWE96_132
Leptidea_juvernica_SWE_SWE96_128
Leptidea_juvernica_SWE_SWE96_127
Leptidea_juvernica_SWE_SWE96_126
Leptidea_juvernica_SWE_SWE96_125
Leptidea_juvernica_SWE_SWE96_124
Leptidea_juvernica_SWE_SWE96_121
Leptidea_juvernica_SWE_SWE96_120
Leptidea_juvernica_SWE_SWE96_119
Leptidea_juvernica_SWE_SWE96_117
Leptidea_juvernica_SWE_Lars_10X_112
Leptidea_juvernica_SWE_Lars_10X_111
Leptidea_amurensis_MON_LEP_outgroups_102
Leptidea_amurensis_MON_LEP_outgroups_101
0.0e+00
5.0e+06
1.0e+07
1.5e+07
Start
Dosage
10.0
7.5
5.0
2.5
0.0

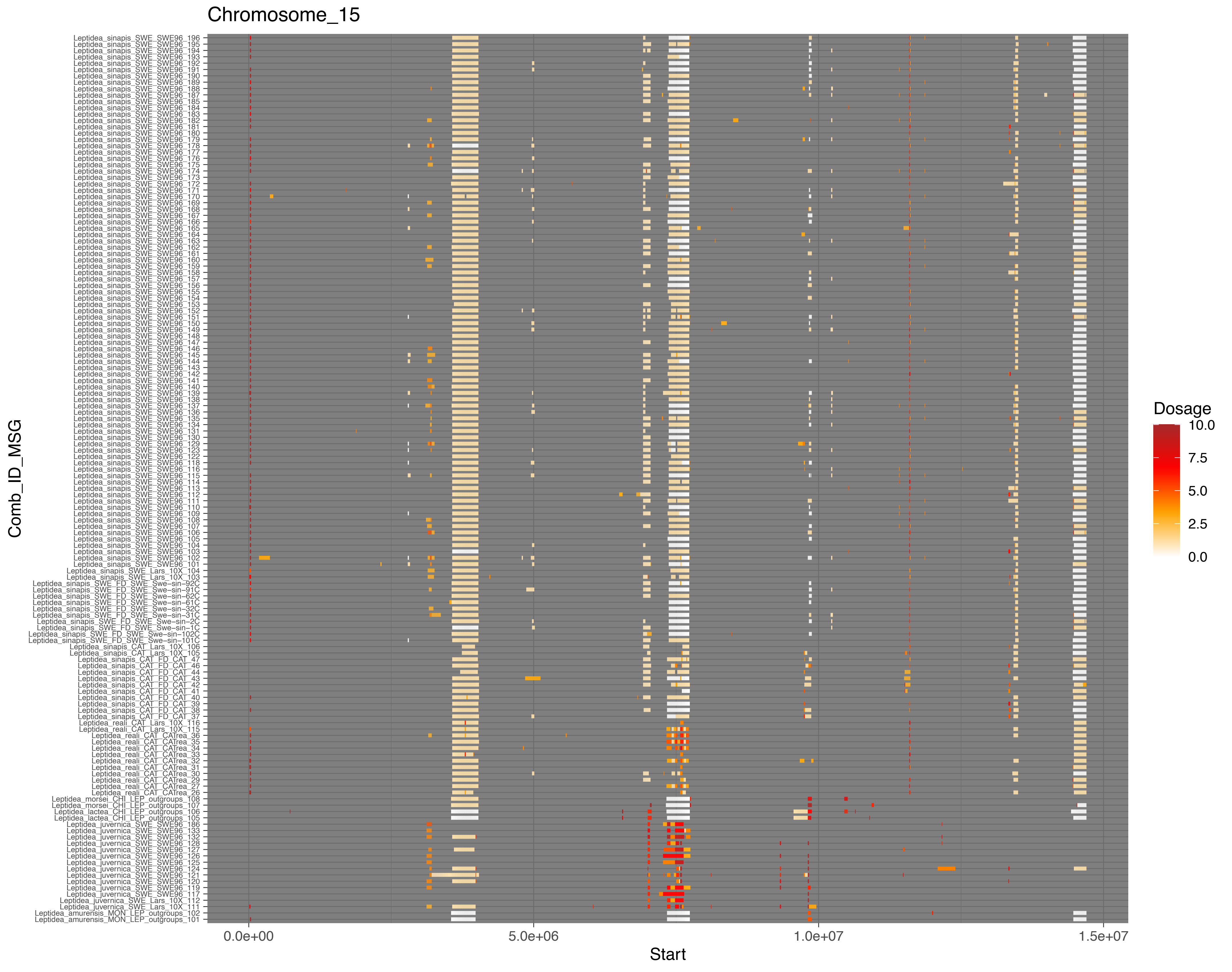
Chromosome_15
Comb_ID_MSG
Leptidea_sinapis_SWE_SWE96_196
Leptidea_sinapis_SWE_SWE96_195
Leptidea_sinapis_SWE_SWE96_194
Leptidea_sinapis_SWE_SWE96_193
Leptidea_sinapis_SWE_SWE96_192
Leptidea_sinapis_SWE_SWE96_191
Leptidea_sinapis_SWE_SWE96_190
Leptidea_sinapis_SWE_SWE96_189
Leptidea_sinapis_SWE_SWE96_188
Leptidea_sinapis_SWE_SWE96_187
Leptidea_sinapis_SWE_SWE96_185
Leptidea_sinapis_SWE_SWE96_184
Leptidea_sinapis_SWE_SWE96_183
Leptidea_sinapis_SWE_SWE96_182
Leptidea_sinapis_SWE_SWE96_181
Leptidea_sinapis_SWE_SWE96_180
Leptidea_sinapis_SWE_SWE96_179
Leptidea_sinapis_SWE_SWE96_178
Leptidea_sinapis_SWE_SWE96_177
Leptidea_sinapis_SWE_SWE96_176
Leptidea_sinapis_SWE_SWE96_175
Leptidea_sinapis_SWE_SWE96_174
Leptidea_sinapis_SWE_SWE96_173
Leptidea_sinapis_SWE_SWE96_172
Leptidea_sinapis_SWE_SWE96_171
Leptidea_sinapis_SWE_SWE96_170
Leptidea_sinapis_SWE_SWE96_169
Leptidea_sinapis_SWE_SWE96_168
Leptidea_sinapis_SWE_SWE96_167
Leptidea_sinapis_SWE_SWE96_166
Leptidea_sinapis_SWE_SWE96_165
Leptidea_sinapis_SWE_SWE96_164
Leptidea_sinapis_SWE_SWE96_163
Leptidea_sinapis_SWE_SWE96_162
Leptidea_sinapis_SWE_SWE96_161
Leptidea_sinapis_SWE_SWE96_160
Leptidea_sinapis_SWE_SWE96_159
Leptidea_sinapis_SWE_SWE96_158
Leptidea_sinapis_SWE_SWE96_157
Leptidea_sinapis_SWE_SWE96_156
Leptidea_sinapis_SWE_SWE96_155
Leptidea_sinapis_SWE_SWE96_154
Leptidea_sinapis_SWE_SWE96_153
Leptidea_sinapis_SWE_SWE96_152
Leptidea_sinapis_SWE_SWE96_151
Leptidea_sinapis_SWE_SWE96_150
Leptidea_sinapis_SWE_SWE96_149
Leptidea_sinapis_SWE_SWE96_148
Leptidea_sinapis_SWE_SWE96_147
Leptidea_sinapis_SWE_SWE96_146
Leptidea_sinapis_SWE_SWE96_145
Leptidea_sinapis_SWE_SWE96_144
Leptidea_sinapis_SWE_SWE96_143
Leptidea_sinapis_SWE_SWE96_142
Leptidea_sinapis_SWE_SWE96_141
Leptidea_sinapis_SWE_SWE96_140
Leptidea_sinapis_SWE_SWE96_139
Leptidea_sinapis_SWE_SWE96_138
Leptidea_sinapis_SWE_SWE96_137
Leptidea_sinapis_SWE_SWE96_136
Leptidea_sinapis_SWE_SWE96_135
Leptidea_sinapis_SWE_SWE96_134
Leptidea_sinapis_SWE_SWE96_131
Leptidea_sinapis_SWE_SWE96_130
Leptidea_sinapis_SWE_SWE96_129
Leptidea_sinapis_SWE_SWE96_123
Leptidea_sinapis_SWE_SWE96_122
Leptidea_sinapis_SWE_SWE96_118
Leptidea_sinapis_SWE_SWE96_116
Leptidea_sinapis_SWE_SWE96_115
Leptidea_sinapis_SWE_SWE96_114
Leptidea_sinapis_SWE_SWE96_113
Leptidea_sinapis_SWE_SWE96_112
Leptidea_sinapis_SWE_SWE96_111
Leptidea_sinapis_SWE_SWE96_110
Leptidea_sinapis_SWE_SWE96_109
Leptidea_sinapis_SWE_SWE96_108
Leptidea_sinapis_SWE_SWE96_107
Leptidea_sinapis_SWE_SWE96_106
Leptidea_sinapis_SWE_SWE96_105
Leptidea_sinapis_SWE_SWE96_104
Leptidea_sinapis_SWE_SWE96_103
Leptidea_sinapis_SWE_SWE96_102
Leptidea_sinapis_SWE_SWE96_101
Leptidea_sinapis_SWE_Lars_10X_104
Leptidea_sinapis_SWE_Lars_10X_103
Leptidea_sinapis_SWE_FD_SWE_Swe−sin−92C
Leptidea_sinapis_SWE_FD_SWE_Swe−sin−91C
Leptidea_sinapis_SWE_FD_SWE_Swe−sin−62C
Leptidea_sinapis_SWE_FD_SWE_Swe−sin−61C
Leptidea_sinapis_SWE_FD_SWE_Swe−sin−32C
Leptidea_sinapis_SWE_FD_SWE_Swe−sin−31C
Leptidea_sinapis_SWE_FD_SWE_Swe−sin−2C
Leptidea_sinapis_SWE_FD_SWE_Swe−sin−1C
Leptidea_sinapis_SWE_FD_SWE_Swe−sin−102C
Leptidea_sinapis_SWE_FD_SWE_Swe−sin−101C
Leptidea_sinapis_CAT_Lars_10X_106
Leptidea_sinapis_CAT_Lars_10X_105
Leptidea_sinapis_CAT_FD_CAT_47
Leptidea_sinapis_CAT_FD_CAT_46
Leptidea_sinapis_CAT_FD_CAT_44
Leptidea_sinapis_CAT_FD_CAT_43
Leptidea_sinapis_CAT_FD_CAT_42
Leptidea_sinapis_CAT_FD_CAT_41
Leptidea_sinapis_CAT_FD_CAT_40
Leptidea_sinapis_CAT_FD_CAT_39
Leptidea_sinapis_CAT_FD_CAT_38
Leptidea_sinapis_CAT_FD_CAT_37
Leptidea_reali_CAT_Lars_10X_116
Leptidea_reali_CAT_Lars_10X_115
Leptidea_reali_CAT_CATrea_36
Leptidea_reali_CAT_CATrea_35
Leptidea_reali_CAT_CATrea_34
Leptidea_reali_CAT_CATrea_33
Leptidea_reali_CAT_CATrea_32
Leptidea_reali_CAT_CATrea_31
Leptidea_reali_CAT_CATrea_30
Leptidea_reali_CAT_CATrea_29
Leptidea_reali_CAT_CATrea_27
Leptidea_reali_CAT_CATrea_26
Leptidea_morsei_CHI_LEP_outgroups_108
Leptidea_morsei_CHI_LEP_outgroups_107
Leptidea_lactea_CHI_LEP_outgroups_106
Leptidea_lactea_CHI_LEP_outgroups_105
Leptidea_juvernica_SWE_SWE96_186
Leptidea_juvernica_SWE_SWE96_133
Leptidea_juvernica_SWE_SWE96_132
Leptidea_juvernica_SWE_SWE96_128
Leptidea_juvernica_SWE_SWE96_127
Leptidea_juvernica_SWE_SWE96_126
Leptidea_juvernica_SWE_SWE96_125
Leptidea_juvernica_SWE_SWE96_124
Leptidea_juvernica_SWE_SWE96_121
Leptidea_juvernica_SWE_SWE96_120
Leptidea_juvernica_SWE_SWE96_119
Leptidea_juvernica_SWE_SWE96_117
Leptidea_juvernica_SWE_Lars_10X_112
Leptidea_juvernica_SWE_Lars_10X_111
Leptidea_amurensis_MON_LEP_outgroups_102
Leptidea_amurensis_MON_LEP_outgroups_101
0.0e+00
5.0e+06
1.0e+07
1.5e+07
Start
Dosage
10.0
7.5
5.0
2.5
0.0

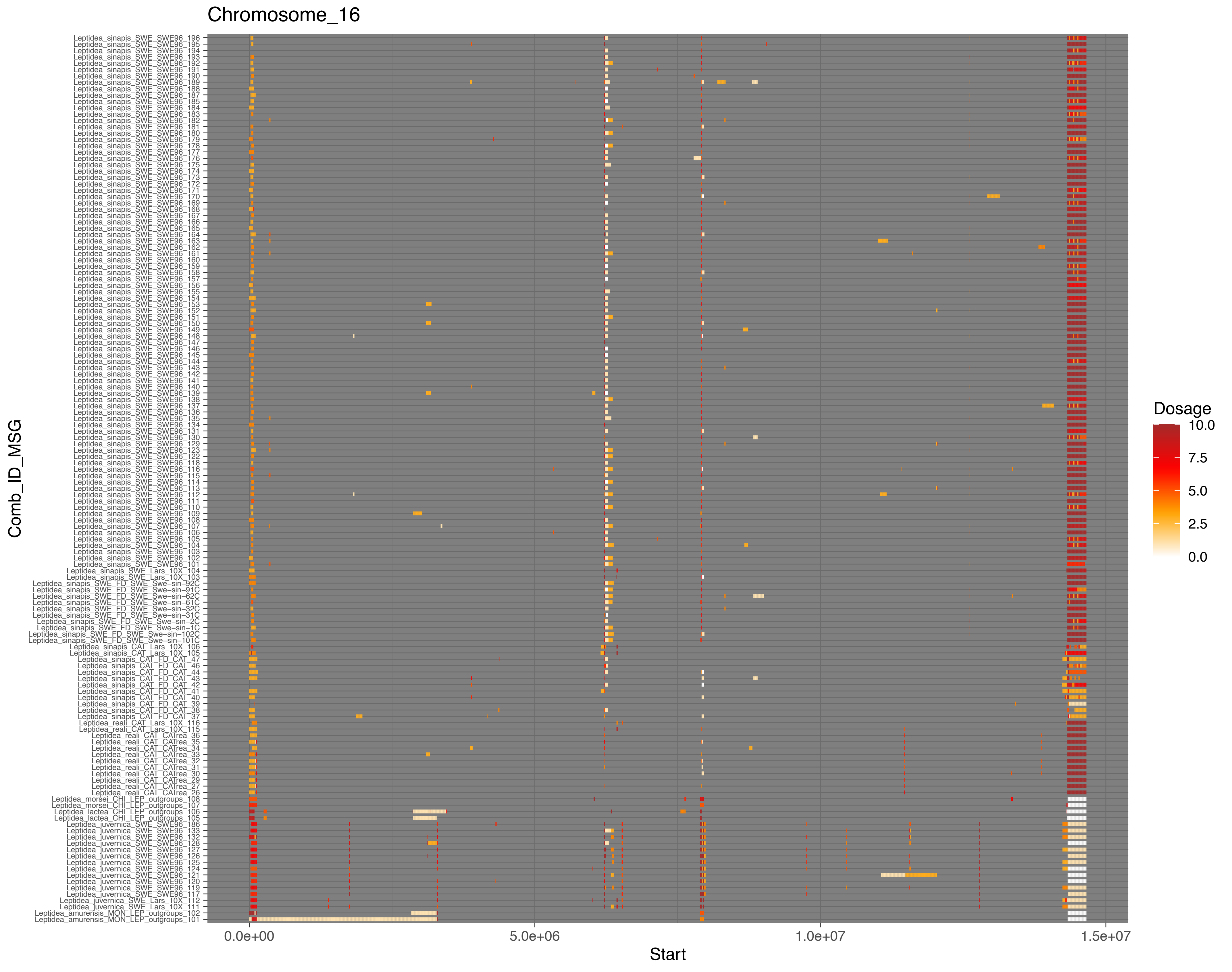

Chromosome_16
Leptidea_sinapis_SWE_SWE96_196
Leptidea_sinapis_SWE_SWE96_195
Leptidea_sinapis_SWE_SWE96_194
Leptidea_sinapis_SWE_SWE96_193
Leptidea_sinapis_SWE_SWE96_192
Leptidea_sinapis_SWE_SWE96_191
Leptidea_sinapis_SWE_SWE96_190
Leptidea_sinapis_SWE_SWE96_189
Leptidea_sinapis_SWE_SWE96_188
Leptidea_sinapis_SWE_SWE96_187
Leptidea_sinapis_SWE_SWE96_185
Leptidea_sinapis_SWE_SWE96_184
Leptidea_sinapis_SWE_SWE96_183
Leptidea_sinapis_SWE_SWE96_182
Leptidea_sinapis_SWE_SWE96_181
Leptidea_sinapis_SWE_SWE96_180
Leptidea_sinapis_SWE_SWE96_179
Leptidea_sinapis_SWE_SWE96_178
Leptidea_sinapis_SWE_SWE96_177
Leptidea_sinapis_SWE_SWE96_176
Leptidea_sinapis_SWE_SWE96_175
Leptidea_sinapis_SWE_SWE96_174
Leptidea_sinapis_SWE_SWE96_173
Leptidea_sinapis_SWE_SWE96_172
Leptidea_sinapis_SWE_SWE96_171
Leptidea_sinapis_SWE_SWE96_170
Leptidea_sinapis_SWE_SWE96_169
Leptidea_sinapis_SWE_SWE96_168
Leptidea_sinapis_SWE_SWE96_167
Leptidea_sinapis_SWE_SWE96_166
Leptidea_sinapis_SWE_SWE96_165
Leptidea_sinapis_SWE_SWE96_164
Leptidea_sinapis_SWE_SWE96_163
Leptidea_sinapis_SWE_SWE96_162
Leptidea_sinapis_SWE_SWE96_161
Leptidea_sinapis_SWE_SWE96_160
Leptidea_sinapis_SWE_SWE96_159
Leptidea_sinapis_SWE_SWE96_158
Leptidea_sinapis_SWE_SWE96_157
Leptidea_sinapis_SWE_SWE96_156
Leptidea_sinapis_SWE_SWE96_155
Leptidea_sinapis_SWE_SWE96_154
Leptidea_sinapis_SWE_SWE96_153
Leptidea_sinapis_SWE_SWE96_152
Leptidea_sinapis_SWE_SWE96_151
Leptidea_sinapis_SWE_SWE96_150
Leptidea_sinapis_SWE_SWE96_149
Leptidea_sinapis_SWE_SWE96_148
Leptidea_sinapis_SWE_SWE96_147
Leptidea_sinapis_SWE_SWE96_146
Leptidea_sinapis_SWE_SWE96_145
Leptidea_sinapis_SWE_SWE96_144
Leptidea_sinapis_SWE_SWE96_143
Leptidea_sinapis_SWE_SWE96_142
Leptidea_sinapis_SWE_SWE96_141
Leptidea_sinapis_SWE_SWE96_140
Leptidea_sinapis_SWE_SWE96_139
Leptidea_sinapis_SWE_SWE96_138
Leptidea_sinapis_SWE_SWE96_137
Leptidea_sinapis_SWE_SWE96_136
Leptidea_sinapis_SWE_SWE96_135
Leptidea_sinapis_SWE_SWE96_134
Leptidea_sinapis_SWE_SWE96_131
Leptidea_sinapis_SWE_SWE96_130
Leptidea_sinapis_SWE_SWE96_129
Leptidea_sinapis_SWE_SWE96_123
Leptidea_sinapis_SWE_SWE96_122
Leptidea_sinapis_SWE_SWE96_118
Leptidea_sinapis_SWE_SWE96_116
Leptidea_sinapis_SWE_SWE96_115
Leptidea_sinapis_SWE_SWE96_114
Leptidea_sinapis_SWE_SWE96_113
Leptidea_sinapis_SWE_SWE96_112
Leptidea_sinapis_SWE_SWE96_111
Leptidea_sinapis_SWE_SWE96_110
Leptidea_sinapis_SWE_SWE96_109
Leptidea_sinapis_SWE_SWE96_108
Leptidea_sinapis_SWE_SWE96_107
Leptidea_sinapis_SWE_SWE96_106
Leptidea_sinapis_SWE_SWE96_105
Leptidea_sinapis_SWE_SWE96_104
Leptidea_sinapis_SWE_SWE96_103
Leptidea_sinapis_SWE_SWE96_102
Leptidea_sinapis_SWE_SWE96_101
Leptidea_sinapis_SWE_Lars_10X_104
Leptidea_sinapis_SWE_Lars_10X_103
Leptidea_sinapis_SWE_FD_SWE_Swe-sin-92C
Leptidea_sinapis_SWE_FD_SWE_Swe-sin-91C
Leptidea_sinapis_SWE_FD_SWE_Swe-sin-62C
Leptidea_sinapis_SWE_FD_SWE_Swe-sin-61C
Leptidea_sinapis_SWE_FD_SWE_Swe-sin-32C
Leptidea_sinapis_SWE_FD_SWE_Swe-sin-31C
Leptidea_sinapis_SWE_FD_SWE_Swe-sin-2C
Leptidea_sinapis_SWE_FD_SWE_Swe-sin-1C
Leptidea_sinapis_SWE_FD_SWE_Swe-sin-102C
Leptidea_sinapis_SWE_FD_SWE_Swe-sin-101C
Leptidea_sinapis_CAT_Lars_10X_106
Leptidea_sinapis_CAT_Lars_10X_105
Leptidea_sinapis_CAT_FD_CAT_47
Leptidea_sinapis_CAT_FD_CAT_46
Leptidea_sinapis_CAT_FD_CAT_44
Leptidea_sinapis_CAT_FD_CAT_43
Leptidea_sinapis_CAT_FD_CAT_42
Leptidea_sinapis_CAT_FD_CAT_41
Leptidea_sinapis_CAT_FD_CAT_40
Leptidea_sinapis_CAT_FD_CAT_39
Leptidea_sinapis_CAT_FD_CAT_38
Leptidea_sinapis_CAT_FD_CAT_37
Leptidea_reali_CAT_Lars_10X_116
Leptidea_reali_CAT_Lars_10X_115
Leptidea_reali_CAT_CATrea_36
Leptidea_reali_CAT_CATrea_35
Leptidea_reali_CAT_CATrea_34
Leptidea_reali_CAT_CATrea_33
Leptidea_reali_CAT_CATrea_32
Leptidea_reali_CAT_CATrea_31
Leptidea_reali_CAT_CATrea_30
Leptidea_reali_CAT_CATrea_29
Leptidea_reali_CAT_CATrea_27
Leptidea_reali_CAT_CATrea_26
Leptidea_morsei_CHI_LEP_outgroups_108
Leptidea_morsei_CHI_LEP_outgroups_107
Leptidea_lactea_CHI_LEP_outgroups_106
Leptidea_lactea_CHI_LEP_outgroups_105
Leptidea_juvernica_SWE_SWE96_186
Leptidea_juvernica_SWE_SWE96_133
Leptidea_juvernica_SWE_SWE96_132
Leptidea_juvernica_SWE_SWE96_128
Leptidea_juvernica_SWE_SWE96_127
Leptidea_juvernica_SWE_SWE96_126
Leptidea_juvernica_SWE_SWE96_125
Leptidea_juvernica_SWE_SWE96_124
Leptidea_juvernica_SWE_SWE96_121
Leptidea_juvernica_SWE_SWE96_120
Leptidea_juvernica_SWE_SWE96_119
Leptidea_juvernica_SWE_SWE96_117
Leptidea_juvernica_SWE_Lars_10X_112
Leptidea_juvernica_SWE_Lars_10X_111
Leptidea_amurensis_MON_LEP_outgroups_102
Leptidea_amurensis_MON_LEP_outgroups_101
Comb_ID_MSG
0.0e+00
5.0e+06
1.0e+07
1.5e+07
Start
Dosage
10.0
7.5
5.0
2.5
0.0

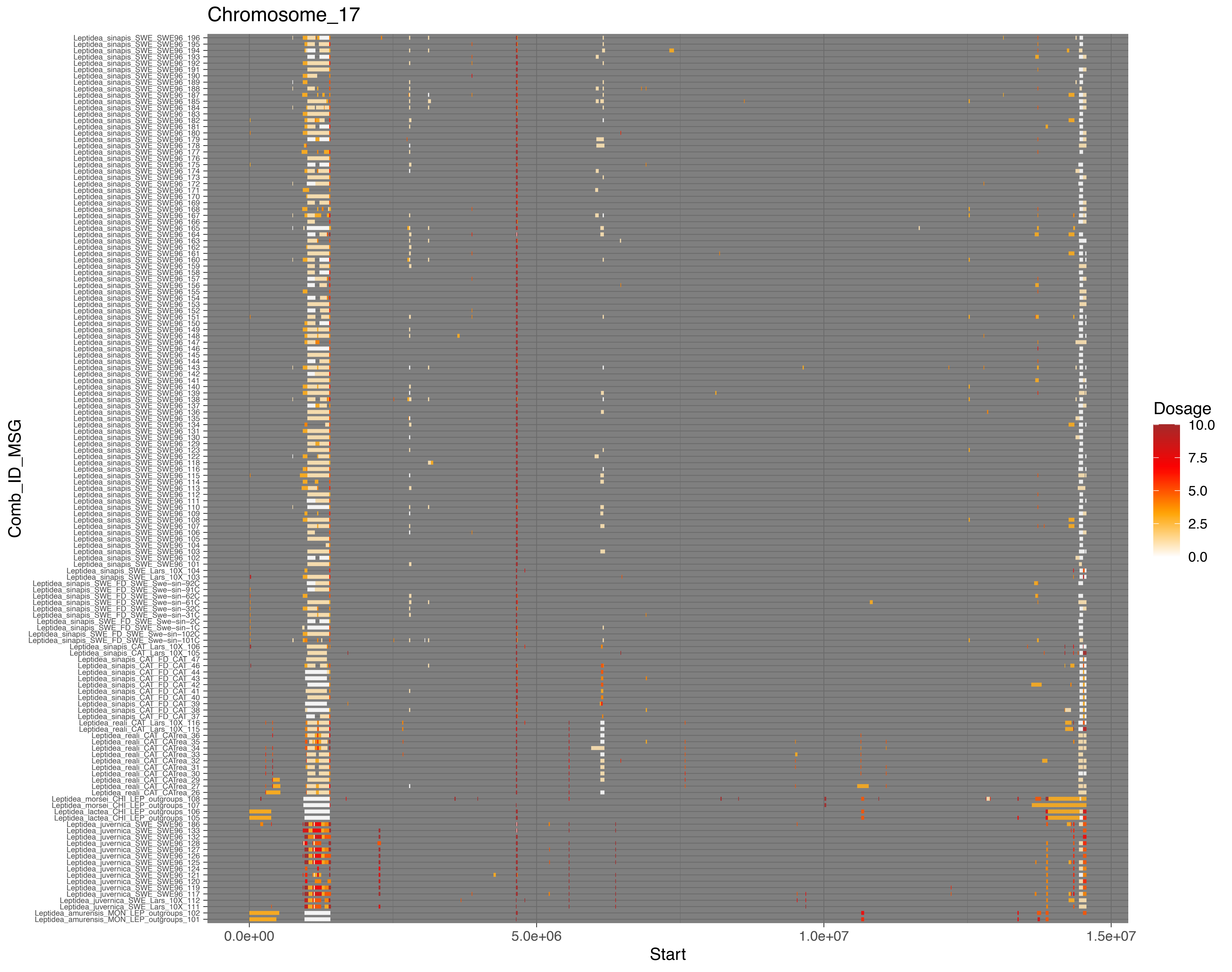

Chromosome_17
Comb_ID_MSG
Leptidea_sinapis_SWE_SWE96_196
Leptidea_sinapis_SWE_SWE96_195
Leptidea_sinapis_SWE_SWE96_194
Leptidea_sinapis_SWE_SWE96_193
Leptidea_sinapis_SWE_SWE96_192
Leptidea_sinapis_SWE_SWE96_191
Leptidea_sinapis_SWE_SWE96_190
Leptidea_sinapis_SWE_SWE96_189
Leptidea_sinapis_SWE_SWE96_188
Leptidea_sinapis_SWE_SWE96_187
Leptidea_sinapis_SWE_SWE96_185
Leptidea_sinapis_SWE_SWE96_184
Leptidea_sinapis_SWE_SWE96_183
Leptidea_sinapis_SWE_SWE96_182
Leptidea_sinapis_SWE_SWE96_181
Leptidea_sinapis_SWE_SWE96_180
Leptidea_sinapis_SWE_SWE96_179
Leptidea_sinapis_SWE_SWE96_178
Leptidea_sinapis_SWE_SWE96_177
Leptidea_sinapis_SWE_SWE96_176
Leptidea_sinapis_SWE_SWE96_175
Leptidea_sinapis_SWE_SWE96_174
Leptidea_sinapis_SWE_SWE96_173
Leptidea_sinapis_SWE_SWE96_172
Leptidea_sinapis_SWE_SWE96_171
Leptidea_sinapis_SWE_SWE96_170
Leptidea_sinapis_SWE_SWE96_169
Leptidea_sinapis_SWE_SWE96_168
Leptidea_sinapis_SWE_SWE96_167
Leptidea_sinapis_SWE_SWE96_166
Leptidea_sinapis_SWE_SWE96_165
Leptidea_sinapis_SWE_SWE96_164
Leptidea_sinapis_SWE_SWE96_163
Leptidea_sinapis_SWE_SWE96_162
Leptidea_sinapis_SWE_SWE96_161
Leptidea_sinapis_SWE_SWE96_160
Leptidea_sinapis_SWE_SWE96_159
Leptidea_sinapis_SWE_SWE96_158
Leptidea_sinapis_SWE_SWE96_157
Leptidea_sinapis_SWE_SWE96_156
Leptidea_sinapis_SWE_SWE96_155
Leptidea_sinapis_SWE_SWE96_154
Leptidea_sinapis_SWE_SWE96_153
Leptidea_sinapis_SWE_SWE96_152
Leptidea_sinapis_SWE_SWE96_151
Leptidea_sinapis_SWE_SWE96_150
Leptidea_sinapis_SWE_SWE96_149
Leptidea_sinapis_SWE_SWE96_148
Leptidea_sinapis_SWE_SWE96_147
Leptidea_sinapis_SWE_SWE96_146
Leptidea_sinapis_SWE_SWE96_145
Leptidea_sinapis_SWE_SWE96_144
Leptidea_sinapis_SWE_SWE96_143
Leptidea_sinapis_SWE_SWE96_142
Leptidea_sinapis_SWE_SWE96_141
Leptidea_sinapis_SWE_SWE96_140
Leptidea_sinapis_SWE_SWE96_139
Leptidea_sinapis_SWE_SWE96_138
Leptidea_sinapis_SWE_SWE96_137
Leptidea_sinapis_SWE_SWE96_136
Leptidea_sinapis_SWE_SWE96_135
Leptidea_sinapis_SWE_SWE96_134
Leptidea_sinapis_SWE_SWE96_131
Leptidea_sinapis_SWE_SWE96_130
Leptidea_sinapis_SWE_SWE96_129
Leptidea_sinapis_SWE_SWE96_123
Leptidea_sinapis_SWE_SWE96_122
Leptidea_sinapis_SWE_SWE96_118
Leptidea_sinapis_SWE_SWE96_116
Leptidea_sinapis_SWE_SWE96_115
Leptidea_sinapis_SWE_SWE96_114
Leptidea_sinapis_SWE_SWE96_113
Leptidea_sinapis_SWE_SWE96_112
Leptidea_sinapis_SWE_SWE96_111
Leptidea_sinapis_SWE_SWE96_110
Leptidea_sinapis_SWE_SWE96_109
Leptidea_sinapis_SWE_SWE96_108
Leptidea_sinapis_SWE_SWE96_107
Leptidea_sinapis_SWE_SWE96_106
Leptidea_sinapis_SWE_SWE96_105
Leptidea_sinapis_SWE_SWE96_104
Leptidea_sinapis_SWE_SWE96_103
Leptidea_sinapis_SWE_SWE96_102
Leptidea_sinapis_SWE_SWE96_101
Leptidea_sinapis_SWE_Lars_10X_104
Leptidea_sinapis_SWE_Lars_10X_103
Leptidea_sinapis_SWE_FD_SWE_Swe−sin−92C
Leptidea_sinapis_SWE_FD_SWE_Swe−sin−91C
Leptidea_sinapis_SWE_FD_SWE_Swe−sin−62C
Leptidea_sinapis_SWE_FD_SWE_Swe−sin−61C
Leptidea_sinapis_SWE_FD_SWE_Swe−sin−32C
Leptidea_sinapis_SWE_FD_SWE_Swe−sin−31C
Leptidea_sinapis_SWE_FD_SWE_Swe−sin−2C
Leptidea_sinapis_SWE_FD_SWE_Swe−sin−1C
Leptidea_sinapis_SWE_FD_SWE_Swe−sin−102C
Leptidea_sinapis_SWE_FD_SWE_Swe−sin−101C
Leptidea_sinapis_CAT_Lars_10X_106
Leptidea_sinapis_CAT_Lars_10X_105
Leptidea_sinapis_CAT_FD_CAT_47
Leptidea_sinapis_CAT_FD_CAT_46
Leptidea_sinapis_CAT_FD_CAT_44
Leptidea_sinapis_CAT_FD_CAT_43
Leptidea_sinapis_CAT_FD_CAT_42
Leptidea_sinapis_CAT_FD_CAT_40
Leptidea_sinapis_CAT_FD_CAT_41
Leptidea_sinapis_CAT_FD_CAT_39
Leptidea_sinapis_CAT_FD_CAT_38
Leptidea_sinapis_CAT_FD_CAT_37
Leptidea_reali_CAT_Lars_10X_116
Leptidea_reali_CAT_Lars_10X_115
Leptidea_reali_CAT_CAtrea_36
Leptidea_reali_CAT_CAtrea_35
Leptidea_reali_CAT_CAtrea_34
Leptidea_reali_CAT_CAtrea_33
Leptidea_reali_CAT_CAtrea_32
Leptidea_reali_CAT_CAtrea_31
Leptidea_reali_CAT_CAtrea_30
Leptidea_reali_CAT_CAtrea_29
Leptidea_reali_CAT_CAtrea_27
Leptidea_reali_CAT_CAtrea_26
Leptidea_morsei_CHI_LEP_outgroups_108
Leptidea_morsei_CHI_LEP_outgroups_107
Leptidea_lactea_CHI_LEP_outgroups_106
Leptidea_lactea_CHI_LEP_outgroups_105
Leptidea_juvernica_SWE_SWE96_186
Leptidea_juvernica_SWE_SWE96_133
Leptidea_juvernica_SWE_SWE96_132
Leptidea_juvernica_SWE_SWE96_128
Leptidea_juvernica_SWE_SWE96_127
Leptidea_juvernica_SWE_SWE96_126
Leptidea_juvernica_SWE_SWE96_125
Leptidea_juvernica_SWE_SWE96_124
Leptidea_juvernica_SWE_SWE96_121
Leptidea_juvernica_SWE_SWE96_120
Leptidea_juvernica_SWE_SWE96_119
Leptidea_juvernica_SWE_SWE96_117
Leptidea_juvernica_SWE_Lars_10X_112
Leptidea_juvernica_SWE_Lars_10X_111
Leptidea_amurensis_MON_LEP_outgroups_102
Leptidea_amurensis_MON_LEP_outgroups_101
0.0e+00
5.0e+06
1.0e+07
1.5e+07
Start
Dosage
10.0
7.5
5.0
2.5
0.0

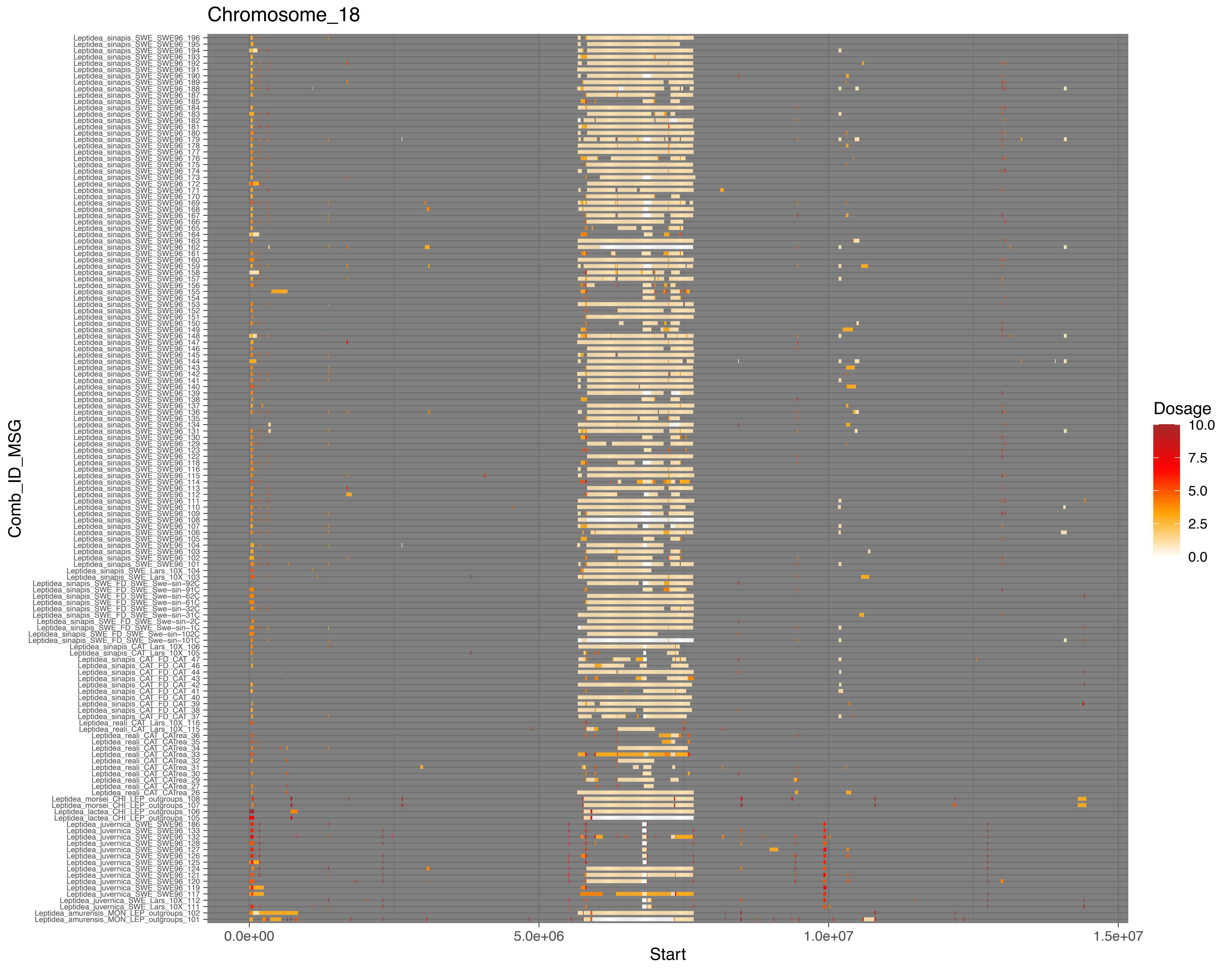
Chromosome_18
Comb_ID_MSG
Start
0.0e+00
5.0e+06
1.0e+07
1.5e+07
Dosage
10.0
7.5
5.0
2.5
0.0

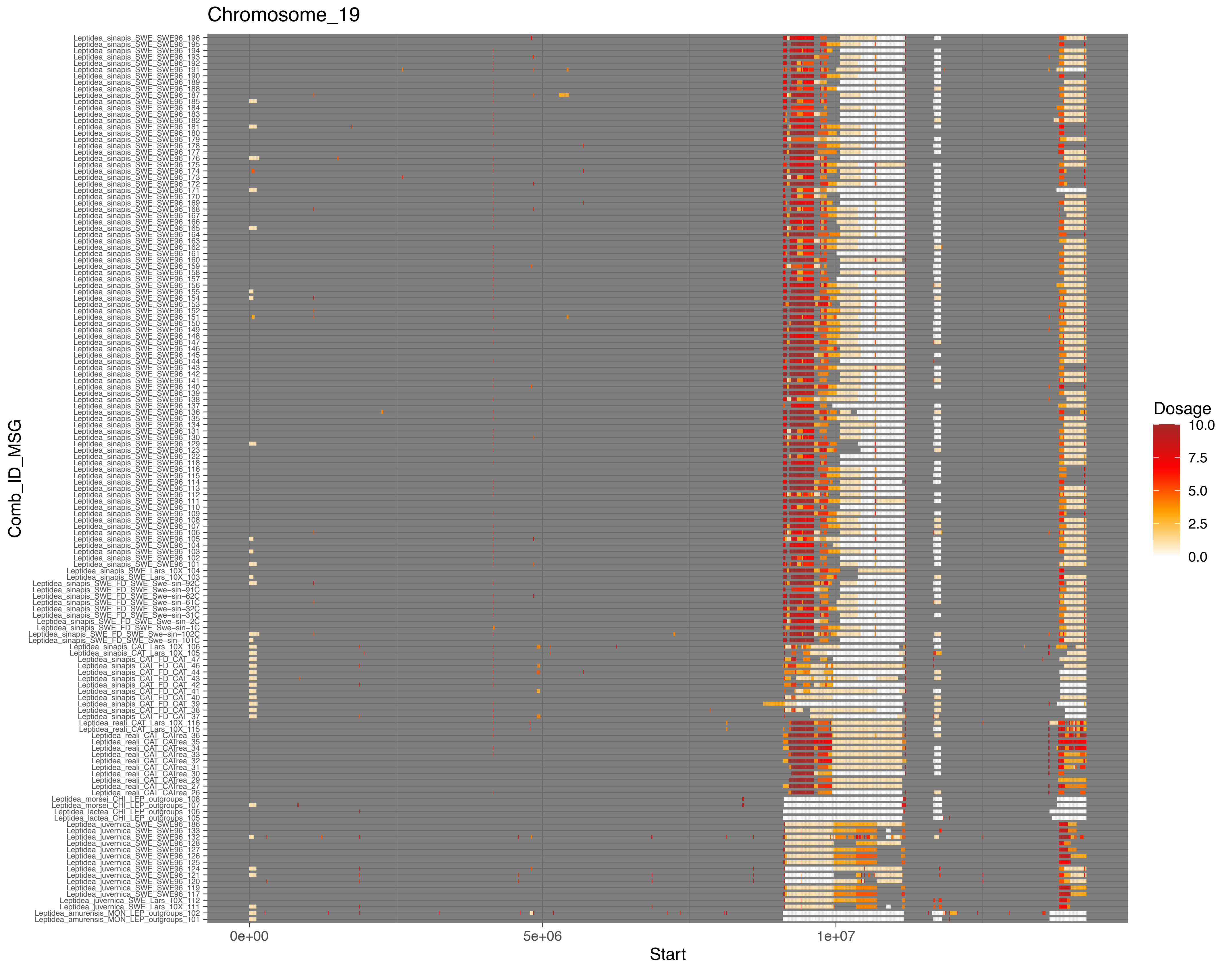
Chromosome_19
Comb_ID_MSG
Leptidea_sinapis_SWE_SWE96_196
Leptidea_sinapis_SWE_SWE96_195
Leptidea_sinapis_SWE_SWE96_194
Leptidea_sinapis_SWE_SWE96_193
Leptidea_sinapis_SWE_SWE96_192
Leptidea_sinapis_SWE_SWE96_191
Leptidea_sinapis_SWE_SWE96_190
Leptidea_sinapis_SWE_SWE96_189
Leptidea_sinapis_SWE_SWE96_188
Leptidea_sinapis_SWE_SWE96_187
Leptidea_sinapis_SWE_SWE96_185
Leptidea_sinapis_SWE_SWE96_184
Leptidea_sinapis_SWE_SWE96_183
Leptidea_sinapis_SWE_SWE96_182
Leptidea_sinapis_SWE_SWE96_181
Leptidea_sinapis_SWE_SWE96_180
Leptidea_sinapis_SWE_SWE96_179
Leptidea_sinapis_SWE_SWE96_178
Leptidea_sinapis_SWE_SWE96_177
Leptidea_sinapis_SWE_SWE96_176
Leptidea_sinapis_SWE_SWE96_175
Leptidea_sinapis_SWE_SWE96_174
Leptidea_sinapis_SWE_SWE96_173
Leptidea_sinapis_SWE_SWE96_172
Leptidea_sinapis_SWE_SWE96_171
Leptidea_sinapis_SWE_SWE96_170
Leptidea_sinapis_SWE_SWE96_169
Leptidea_sinapis_SWE_SWE96_168
Leptidea_sinapis_SWE_SWE96_167
Leptidea_sinapis_SWE_SWE96_166
Leptidea_sinapis_SWE_SWE96_165
Leptidea_sinapis_SWE_SWE96_164
Leptidea_sinapis_SWE_SWE96_163
Leptidea_sinapis_SWE_SWE96_162
Leptidea_sinapis_SWE_SWE96_161
Leptidea_sinapis_SWE_SWE96_160
Leptidea_sinapis_SWE_SWE96_159
Leptidea_sinapis_SWE_SWE96_158
Leptidea_sinapis_SWE_SWE96_157
Leptidea_sinapis_SWE_SWE96_156
Leptidea_sinapis_SWE_SWE96_155
Leptidea_sinapis_SWE_SWE96_154
Leptidea_sinapis_SWE_SWE96_153
Leptidea_sinapis_SWE_SWE96_152
Leptidea_sinapis_SWE_SWE96_151
Leptidea_sinapis_SWE_SWE96_150
Leptidea_sinapis_SWE_SWE96_149
Leptidea_sinapis_SWE_SWE96_148
Leptidea_sinapis_SWE_SWE96_147
Leptidea_sinapis_SWE_SWE96_146
Leptidea_sinapis_SWE_SWE96_145
Leptidea_sinapis_SWE_SWE96_144
Leptidea_sinapis_SWE_SWE96_143
Leptidea_sinapis_SWE_SWE96_142
Leptidea_sinapis_SWE_SWE96_141
Leptidea_sinapis_SWE_SWE96_140
Leptidea_sinapis_SWE_SWE96_139
Leptidea_sinapis_SWE_SWE96_138
Leptidea_sinapis_SWE_SWE96_137
Leptidea_sinapis_SWE_SWE96_136
Leptidea_sinapis_SWE_SWE96_135
Leptidea_sinapis_SWE_SWE96_134
Leptidea_sinapis_SWE_SWE96_131
Leptidea_sinapis_SWE_SWE96_130
Leptidea_sinapis_SWE_SWE96_129
Leptidea_sinapis_SWE_SWE96_123
Leptidea_sinapis_SWE_SWE96_122
Leptidea_sinapis_SWE_SWE96_118
Leptidea_sinapis_SWE_SWE96_116
Leptidea_sinapis_SWE_SWE96_115
Leptidea_sinapis_SWE_SWE96_114
Leptidea_sinapis_SWE_SWE96_113
Leptidea_sinapis_SWE_SWE96_112
Leptidea_sinapis_SWE_SWE96_111
Leptidea_sinapis_SWE_SWE96_110
Leptidea_sinapis_SWE_SWE96_109
Leptidea_sinapis_SWE_SWE96_108
Leptidea_sinapis_SWE_SWE96_107
Leptidea_sinapis_SWE_SWE96_106
Leptidea_sinapis_SWE_SWE96_105
Leptidea_sinapis_SWE_SWE96_104
Leptidea_sinapis_SWE_SWE96_103
Leptidea_sinapis_SWE_SWE96_102
Leptidea_sinapis_SWE_SWE96_101
Leptidea_sinapis_SWE_Lars_10X_104
Leptidea_sinapis_SWE_Lars_10X_103
Leptidea_sinapis_SWE_FD_SWE_Swe−sin−92C
Leptidea_sinapis_SWE_FD_SWE_Swe−sin−91C
Leptidea_sinapis_SWE_FD_SWE_Swe−sin−62C
Leptidea_sinapis_SWE_FD_SWE_Swe−sin−61C
Leptidea_sinapis_SWE_FD_SWE_Swe−sin−32C
Leptidea_sinapis_SWE_FD_SWE_Swe−sin−31C
Leptidea_sinapis_SWE_FD_SWE_Swe−sin−2C
Leptidea_sinapis_SWE_FD_SWE_Swe−sin−1C
Leptidea_sinapis_SWE_FD_SWE_Swe−sin−102C
Leptidea_sinapis_SWE_FD_SWE_Swe−sin−101C
Leptidea_sinapis_CAT_Lars_10X_106
Leptidea_sinapis_CAT_Lars_10X_105
Leptidea_sinapis_CAT_FD_CAT_47
Leptidea_sinapis_CAT_FD_CAT_46
Leptidea_sinapis_CAT_FD_CAT_44
Leptidea_sinapis_CAT_FD_CAT_43
Leptidea_sinapis_CAT_FD_CAT_42
Leptidea_sinapis_CAT_FD_CAT_41
Leptidea_sinapis_CAT_FD_CAT_40
Leptidea_sinapis_CAT_FD_CAT_39
Leptidea_sinapis_CAT_FD_CAT_38
Leptidea_sinapis_CAT_FD_CAT_37
Leptidea_reali_CAT_Lars_10X_116
Leptidea_reali_CAT_Lars_10X_115
Leptidea_reali_CAT_CATrea_36
Leptidea_reali_CAT_CATrea_35
Leptidea_reali_CAT_CATrea_34
Leptidea_reali_CAT_CATrea_33
Leptidea_reali_CAT_CATrea_32
Leptidea_reali_CAT_CATrea_31
Leptidea_reali_CAT_CATrea_30
Leptidea_reali_CAT_CATrea_29
Leptidea_reali_CAT_CATrea_27
Leptidea_reali_CAT_CATrea_26
Leptidea_morsei_CHI_LEP_outgroups_108
Leptidea_morsei_CHI_LEP_outgroups_107
Leptidea_lactea_CHI_LEP_outgroups_106
Leptidea_lactea_CHI_LEP_outgroups_105
Leptidea_juvernica_SWE_SWE96_186
Leptidea_juvernica_SWE_SWE96_133
Leptidea_juvernica_SWE_SWE96_132
Leptidea_juvernica_SWE_SWE96_128
Leptidea_juvernica_SWE_SWE96_127
Leptidea_juvernica_SWE_SWE96_126
Leptidea_juvernica_SWE_SWE96_125
Leptidea_juvernica_SWE_SWE96_124
Leptidea_juvernica_SWE_SWE96_121
Leptidea_juvernica_SWE_SWE96_120
Leptidea_juvernica_SWE_SWE96_119
Leptidea_juvernica_SWE_SWE96_117
Leptidea_juvernica_SWE_Lars_10X_112
Leptidea_juvernica_SWE_Lars_10X_111
Leptidea_amurensis_MON_LEP_outgroups_102
Leptidea_amurensis_MON_LEP_outgroups_101
0e+00
5e+06
1e+07
Start
Dosage
10.0
7.5
5.0
2.5
0.0

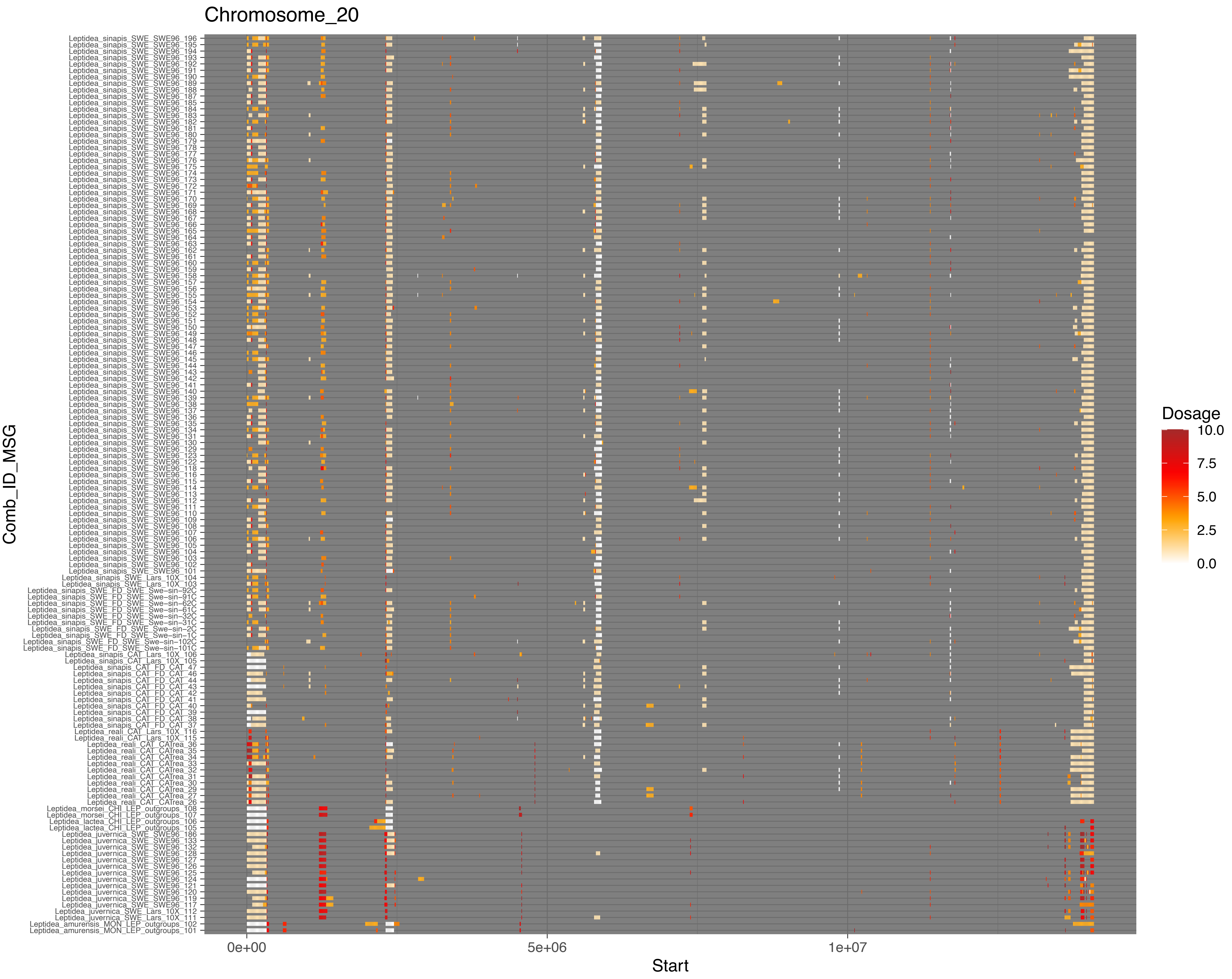
Chromosome_20
Comb_ID_MSG
Leptidea_sinapis_SWE_SWE96_196
Leptidea_sinapis_SWE_SWE96_195
Leptidea_sinapis_SWE_SWE96_194
Leptidea_sinapis_SWE_SWE96_193
Leptidea_sinapis_SWE_SWE96_192
Leptidea_sinapis_SWE_SWE96_191
Leptidea_sinapis_SWE_SWE96_190
Leptidea_sinapis_SWE_SWE96_189
Leptidea_sinapis_SWE_SWE96_188
Leptidea_sinapis_SWE_SWE96_187
Leptidea_sinapis_SWE_SWE96_185
Leptidea_sinapis_SWE_SWE96_184
Leptidea_sinapis_SWE_SWE96_183
Leptidea_sinapis_SWE_SWE96_182
Leptidea_sinapis_SWE_SWE96_181
Leptidea_sinapis_SWE_SWE96_180
Leptidea_sinapis_SWE_SWE96_179
Leptidea_sinapis_SWE_SWE96_178
Leptidea_sinapis_SWE_SWE96_177
Leptidea_sinapis_SWE_SWE96_176
Leptidea_sinapis_SWE_SWE96_175
Leptidea_sinapis_SWE_SWE96_174
Leptidea_sinapis_SWE_SWE96_173
Leptidea_sinapis_SWE_SWE96_172
Leptidea_sinapis_SWE_SWE96_171
Leptidea_sinapis_SWE_SWE96_170
Leptidea_sinapis_SWE_SWE96_169
Leptidea_sinapis_SWE_SWE96_168
Leptidea_sinapis_SWE_SWE96_167
Leptidea_sinapis_SWE_SWE96_166
Leptidea_sinapis_SWE_SWE96_165
Leptidea_sinapis_SWE_SWE96_164
Leptidea_sinapis_SWE_SWE96_163
Leptidea_sinapis_SWE_SWE96_162
Leptidea_sinapis_SWE_SWE96_161
Leptidea_sinapis_SWE_SWE96_160
Leptidea_sinapis_SWE_SWE96_159
Leptidea_sinapis_SWE_SWE96_158
Leptidea_sinapis_SWE_SWE96_157
Leptidea_sinapis_SWE_SWE96_156
Leptidea_sinapis_SWE_SWE96_155
Leptidea_sinapis_SWE_SWE96_154
Leptidea_sinapis_SWE_SWE96_153
Leptidea_sinapis_SWE_SWE96_152
Leptidea_sinapis_SWE_SWE96_151
Leptidea_sinapis_SWE_SWE96_150
Leptidea_sinapis_SWE_SWE96_149
Leptidea_sinapis_SWE_SWE96_148
Leptidea_sinapis_SWE_SWE96_147
Leptidea_sinapis_SWE_SWE96_146
Leptidea_sinapis_SWE_SWE96_145
Leptidea_sinapis_SWE_SWE96_144
Leptidea_sinapis_SWE_SWE96_143
Leptidea_sinapis_SWE_SWE96_142
Leptidea_sinapis_SWE_SWE96_141
Leptidea_sinapis_SWE_SWE96_140
Leptidea_sinapis_SWE_SWE96_139
Leptidea_sinapis_SWE_SWE96_138
Leptidea_sinapis_SWE_SWE96_137
Leptidea_sinapis_SWE_SWE96_136
Leptidea_sinapis_SWE_SWE96_135
Leptidea_sinapis_SWE_SWE96_134
Leptidea_sinapis_SWE_SWE96_131
Leptidea_sinapis_SWE_SWE96_130
Leptidea_sinapis_SWE_SWE96_129
Leptidea_sinapis_SWE_SWE96_123
Leptidea_sinapis_SWE_SWE96_122
Leptidea_sinapis_SWE_SWE96_118
Leptidea_sinapis_SWE_SWE96_116
Leptidea_sinapis_SWE_SWE96_115
Leptidea_sinapis_SWE_SWE96_114
Leptidea_sinapis_SWE_SWE96_113
Leptidea_sinapis_SWE_SWE96_112
Leptidea_sinapis_SWE_SWE96_111
Leptidea_sinapis_SWE_SWE96_110
Leptidea_sinapis_SWE_SWE96_109
Leptidea_sinapis_SWE_SWE96_108
Leptidea_sinapis_SWE_SWE96_107
Leptidea_sinapis_SWE_SWE96_106
Leptidea_sinapis_SWE_SWE96_105
Leptidea_sinapis_SWE_SWE96_104
Leptidea_sinapis_SWE_SWE96_103
Leptidea_sinapis_SWE_SWE96_102
Leptidea_sinapis_SWE_SWE96_101
Leptidea_sinapis_SWE_Lars_10X_104
Leptidea_sinapis_SWE_Lars_10X_103
Leptidea_sinapis_SWE_FD_SWE_Swe-sin-92C
Leptidea_sinapis_SWE_FD_SWE_Swe-sin-91C
Leptidea_sinapis_SWE_FD_SWE_Swe-sin-62C
Leptidea_sinapis_SWE_FD_SWE_Swe-sin-61C
Leptidea_sinapis_SWE_FD_SWE_Swe-sin-32C
Leptidea_sinapis_SWE_FD_SWE_Swe-sin-31C
Leptidea_sinapis_SWE_FD_SWE_Swe-sin-2C
Leptidea_sinapis_SWE_FD_SWE_Swe-sin-1C
Leptidea_sinapis_SWE_FD_SWE_Swe-sin-102C
Leptidea_sinapis_SWE_FD_SWE_Swe-sin-101C
Leptidea_sinapis_CAT_Lars_10X_106
Leptidea_sinapis_CAT_Lars_10X_105
Leptidea_sinapis_CAT_FD_CAT_47
Leptidea_sinapis_CAT_FD_CAT_46
Leptidea_sinapis_CAT_FD_CAT_44
Leptidea_sinapis_CAT_FD_CAT_43
Leptidea_sinapis_CAT_FD_CAT_42
Leptidea_sinapis_CAT_FD_CAT_41
Leptidea_sinapis_CAT_FD_CAT_40
Leptidea_sinapis_CAT_FD_CAT_39
Leptidea_sinapis_CAT_FD_CAT_38
Leptidea_sinapis_CAT_FD_CAT_37
Leptidea_reali_CAT_Lars_10X_116
Leptidea_reali_CAT_Lars_10X_115
Leptidea_reali_CAT_CATrea_36
Leptidea_reali_CAT_CATrea_35
Leptidea_reali_CAT_CATrea_34
Leptidea_reali_CAT_CATrea_33
Leptidea_reali_CAT_CATrea_32
Leptidea_reali_CAT_CATrea_31
Leptidea_reali_CAT_CATrea_30
Leptidea_reali_CAT_CATrea_29
Leptidea_reali_CAT_CATrea_27
Leptidea_reali_CAT_CATrea_26
Leptidea_morsei_CHI_LEP_outgroups_108
Leptidea_morsei_CHI_LEP_outgroups_107
Leptidea_lactea_CHI_LEP_outgroups_106
Leptidea_lactea_CHI_LEP_outgroups_105
Leptidea_juvernica_SWE_SWE96_186
Leptidea_juvernica_SWE_SWE96_133
Leptidea_juvernica_SWE_SWE96_132
Leptidea_juvernica_SWE_SWE96_128
Leptidea_juvernica_SWE_SWE96_127
Leptidea_juvernica_SWE_SWE96_126
Leptidea_juvernica_SWE_SWE96_125
Leptidea_juvernica_SWE_SWE96_124
Leptidea_juvernica_SWE_SWE96_121
Leptidea_juvernica_SWE_SWE96_120
Leptidea_juvernica_SWE_SWE96_119
Leptidea_juvernica_SWE_SWE96_117
Leptidea_juvernica_SWE_Lars_10X_112
Leptidea_juvernica_SWE_Lars_10X_111
Leptidea_amurensis_MON_LEP_outgroups_102
Leptidea_amurensis_MON_LEP_outgroups_101
0e+00
5e+06
1e+07
Start
Dosage
10.0
7.5
5.0
2.5
0.0

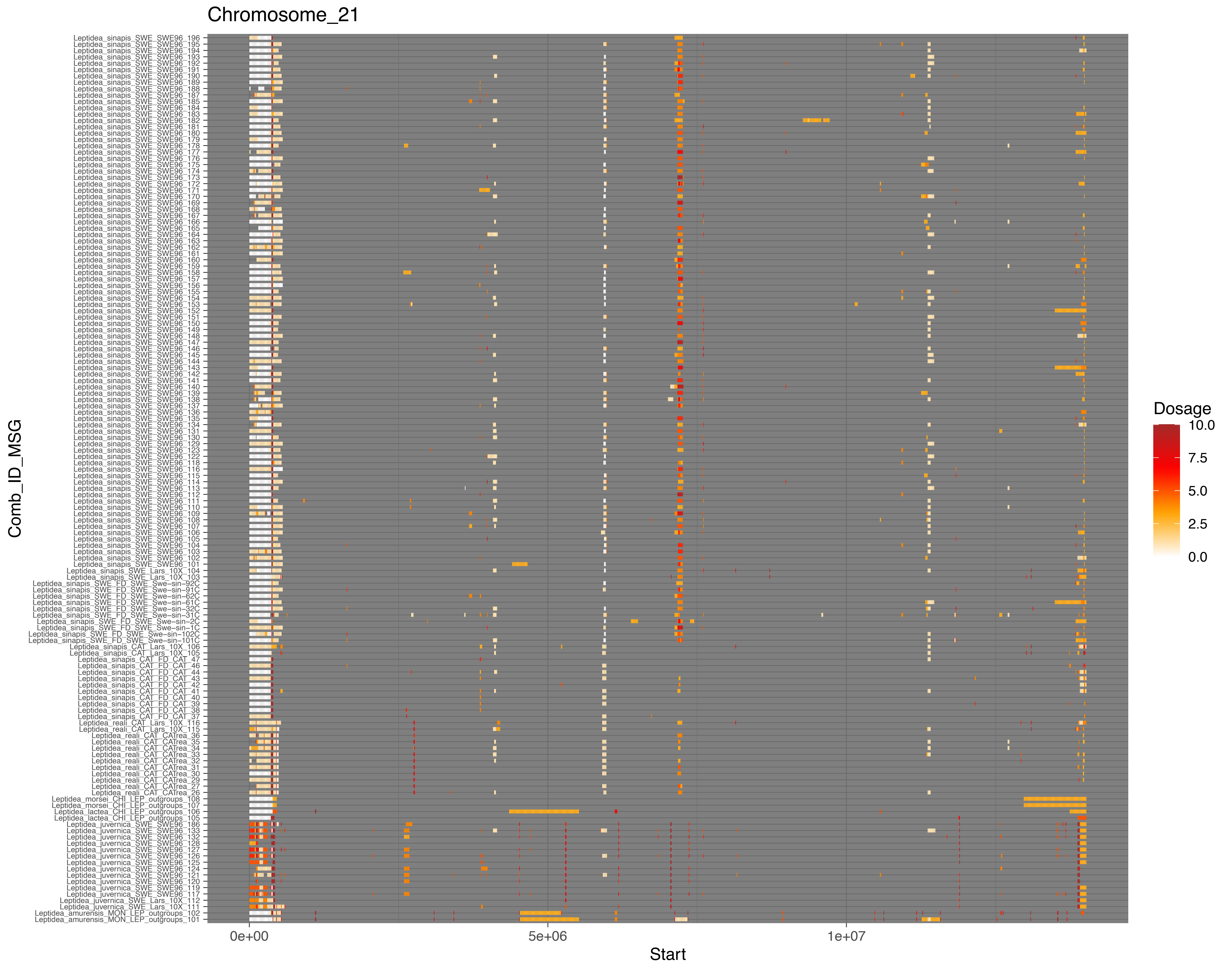
Chromosome_21
Comb_ID_MSG
Start
0e+00
5e+06
1e+07
Dosage
10.0
7.5
5.0
2.5
0.0

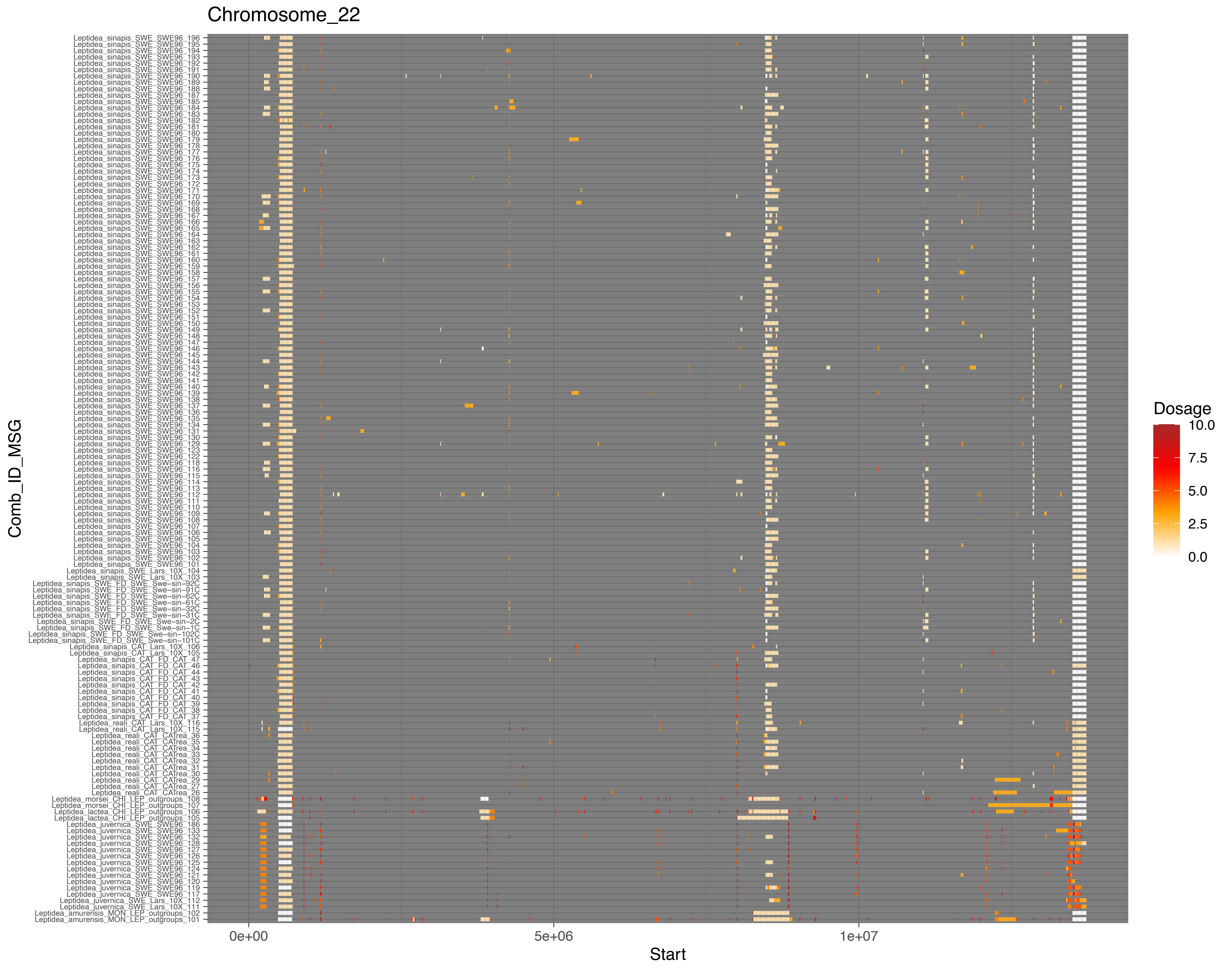
Chromosome_22
Comb_ID_MSG
Start
0e+00
5e+06
1e+07
Dosage
10.0
7.5
5.0
2.5
0.0

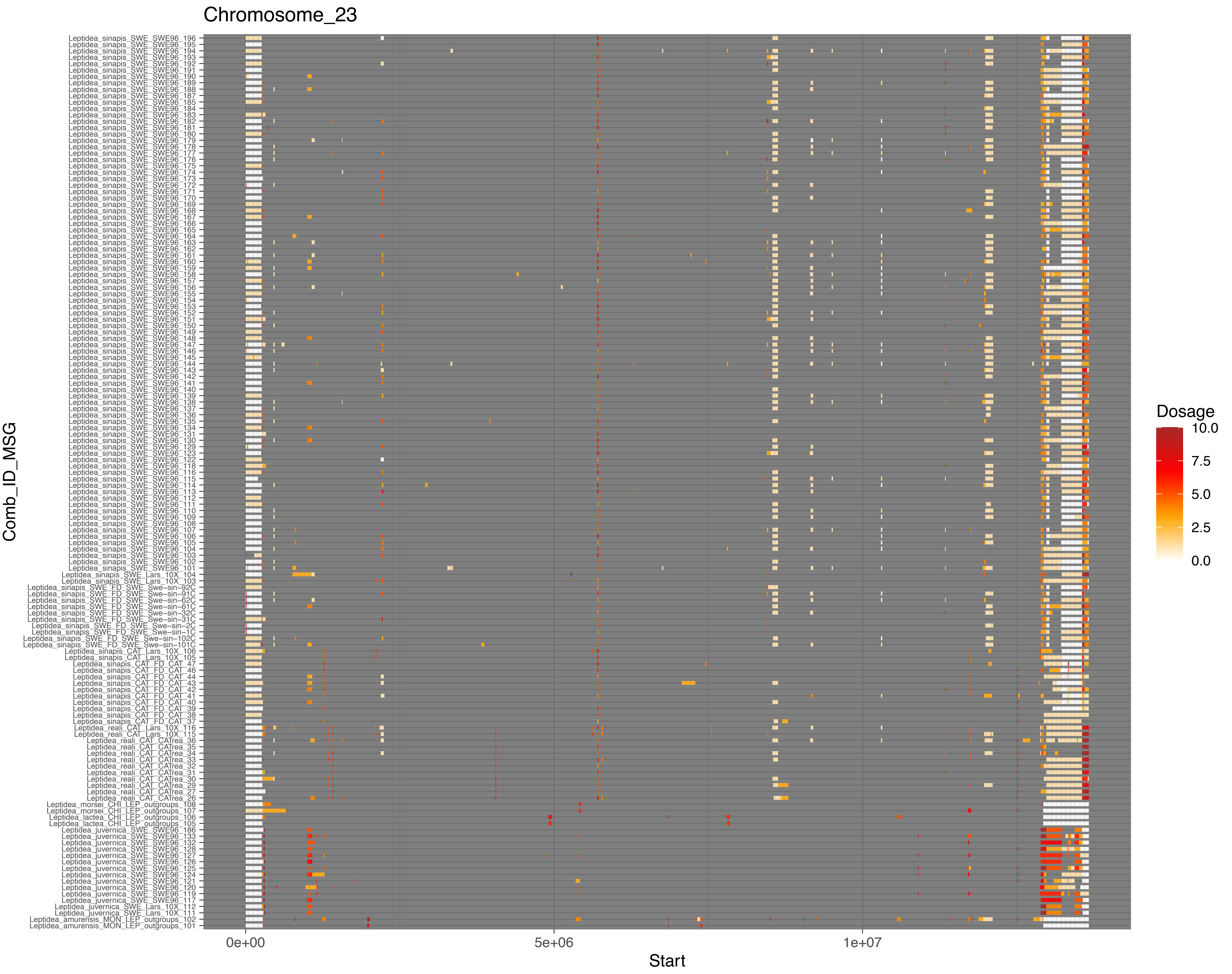

Chromosome_23
Comb_ID_MSG
Leptidea_sinapis_SWE_SWE96_196
Leptidea_sinapis_SWE_SWE96_195
Leptidea_sinapis_SWE_SWE96_194
Leptidea_sinapis_SWE_SWE96_193
Leptidea_sinapis_SWE_SWE96_192
Leptidea_sinapis_SWE_SWE96_191
Leptidea_sinapis_SWE_SWE96_190
Leptidea_sinapis_SWE_SWE96_189
Leptidea_sinapis_SWE_SWE96_188
Leptidea_sinapis_SWE_SWE96_187
Leptidea_sinapis_SWE_SWE96_185
Leptidea_sinapis_SWE_SWE96_184
Leptidea_sinapis_SWE_SWE96_183
Leptidea_sinapis_SWE_SWE96_182
Leptidea_sinapis_SWE_SWE96_181
Leptidea_sinapis_SWE_SWE96_180
Leptidea_sinapis_SWE_SWE96_179
Leptidea_sinapis_SWE_SWE96_178
Leptidea_sinapis_SWE_SWE96_177
Leptidea_sinapis_SWE_SWE96_176
Leptidea_sinapis_SWE_SWE96_175
Leptidea_sinapis_SWE_SWE96_174
Leptidea_sinapis_SWE_SWE96_173
Leptidea_sinapis_SWE_SWE96_172
Leptidea_sinapis_SWE_SWE96_171
Leptidea_sinapis_SWE_SWE96_170
Leptidea_sinapis_SWE_SWE96_169
Leptidea_sinapis_SWE_SWE96_168
Leptidea_sinapis_SWE_SWE96_167
Leptidea_sinapis_SWE_SWE96_166
Leptidea_sinapis_SWE_SWE96_165
Leptidea_sinapis_SWE_SWE96_164
Leptidea_sinapis_SWE_SWE96_163
Leptidea_sinapis_SWE_SWE96_162
Leptidea_sinapis_SWE_SWE96_161
Leptidea_sinapis_SWE_SWE96_160
Leptidea_sinapis_SWE_SWE96_159
Leptidea_sinapis_SWE_SWE96_158
Leptidea_sinapis_SWE_SWE96_157
Leptidea_sinapis_SWE_SWE96_156
Leptidea_sinapis_SWE_SWE96_155
Leptidea_sinapis_SWE_SWE96_154
Leptidea_sinapis_SWE_SWE96_153
Leptidea_sinapis_SWE_SWE96_152
Leptidea_sinapis_SWE_SWE96_151
Leptidea_sinapis_SWE_SWE96_150
Leptidea_sinapis_SWE_SWE96_149
Leptidea_sinapis_SWE_SWE96_148
Leptidea_sinapis_SWE_SWE96_147
Leptidea_sinapis_SWE_SWE96_146
Leptidea_sinapis_SWE_SWE96_145
Leptidea_sinapis_SWE_SWE96_144
Leptidea_sinapis_SWE_SWE96_143
Leptidea_sinapis_SWE_SWE96_142
Leptidea_sinapis_SWE_SWE96_141
Leptidea_sinapis_SWE_SWE96_140
Leptidea_sinapis_SWE_SWE96_139
Leptidea_sinapis_SWE_SWE96_138
Leptidea_sinapis_SWE_SWE96_137
Leptidea_sinapis_SWE_SWE96_136
Leptidea_sinapis_SWE_SWE96_135
Leptidea_sinapis_SWE_SWE96_134
Leptidea_sinapis_SWE_SWE96_131
Leptidea_sinapis_SWE_SWE96_130
Leptidea_sinapis_SWE_SWE96_129
Leptidea_sinapis_SWE_SWE96_123
Leptidea_sinapis_SWE_SWE96_122
Leptidea_sinapis_SWE_SWE96_118
Leptidea_sinapis_SWE_SWE96_116
Leptidea_sinapis_SWE_SWE96_115
Leptidea_sinapis_SWE_SWE96_114
Leptidea_sinapis_SWE_SWE96_113
Leptidea_sinapis_SWE_SWE96_112
Leptidea_sinapis_SWE_SWE96_111
Leptidea_sinapis_SWE_SWE96_110
Leptidea_sinapis_SWE_SWE96_109
Leptidea_sinapis_SWE_SWE96_108
Leptidea_sinapis_SWE_SWE96_107
Leptidea_sinapis_SWE_SWE96_106
Leptidea_sinapis_SWE_SWE96_105
Leptidea_sinapis_SWE_SWE96_104
Leptidea_sinapis_SWE_SWE96_103
Leptidea_sinapis_SWE_SWE96_102
Leptidea_sinapis_SWE_SWE96_101
Leptidea_sinapis_SWE_Lars_10X_104
Leptidea_sinapis_SWE_Lars_10X_103
Leptidea_sinapis_SWE_FD_SWE_Swe−sin−92C
Leptidea_sinapis_SWE_FD_SWE_Swe−sin−91C
Leptidea_sinapis_SWE_FD_SWE_Swe−sin−62C
Leptidea_sinapis_SWE_FD_SWE_Swe−sin−61C
Leptidea_sinapis_SWE_FD_SWE_Swe−sin−32C
Leptidea_sinapis_SWE_FD_SWE_Swe−sin−31C
Leptidea_sinapis_SWE_FD_SWE_Swe−sin−2C
Leptidea_sinapis_SWE_FD_SWE_Swe−sin−1C
Leptidea_sinapis_SWE_FD_SWE_Swe−sin−102C
Leptidea_sinapis_SWE_FD_SWE_Swe−sin−101C
Leptidea_sinapis_CAT_Lars_10X_106
Leptidea_sinapis_CAT_Lars_10X_105
Leptidea_sinapis_CAT_FD_CAT_47
Leptidea_sinapis_CAT_FD_CAT_46
Leptidea_sinapis_CAT_FD_CAT_44
Leptidea_sinapis_CAT_FD_CAT_43
Leptidea_sinapis_CAT_FD_CAT_42
Leptidea_sinapis_CAT_FD_CAT_41
Leptidea_sinapis_CAT_FD_CAT_40
Leptidea_sinapis_CAT_FD_CAT_39
Leptidea_sinapis_CAT_FD_CAT_38
Leptidea_sinapis_CAT_FD_CAT_37
Leptidea_reali_CAT_Lars_10X_116
Leptidea_reali_CAT_Lars_10X_115
Leptidea_reali_CAT_CATrea_36
Leptidea_reali_CAT_CATrea_35
Leptidea_reali_CAT_CATrea_34
Leptidea_reali_CAT_CATrea_33
Leptidea_reali_CAT_CATrea_32
Leptidea_reali_CAT_CATrea_31
Leptidea_reali_CAT_CATrea_30
Leptidea_reali_CAT_CATrea_29
Leptidea_reali_CAT_CATrea_27
Leptidea_reali_CAT_CATrea_26
Leptidea_morsei_CHI_LEP_outgroups_108
Leptidea_morsei_CHI_LEP_outgroups_107
Leptidea_lactea_CHI_LEP_outgroups_106
Leptidea_lactea_CHI_LEP_outgroups_105
Leptidea_juvernica_SWE_SWE96_186
Leptidea_juvernica_SWE_SWE96_133
Leptidea_juvernica_SWE_SWE96_132
Leptidea_juvernica_SWE_SWE96_128
Leptidea_juvernica_SWE_SWE96_127
Leptidea_juvernica_SWE_SWE96_126
Leptidea_juvernica_SWE_SWE96_125
Leptidea_juvernica_SWE_SWE96_124
Leptidea_juvernica_SWE_SWE96_121
Leptidea_juvernica_SWE_SWE96_120
Leptidea_juvernica_SWE_SWE96_119
Leptidea_juvernica_SWE_SWE96_117
Leptidea_juvernica_SWE_Lars_10X_112
Leptidea_juvernica_SWE_Lars_10X_111
Leptidea_amurensis_MON_LEP_outgroups_102
Leptidea_amurensis_MON_LEP_outgroups_101
0e+00
5e+06
1e+07
Start
Dosage
10.0
7.5
5.0
2.5
0.0

Chromosome_24

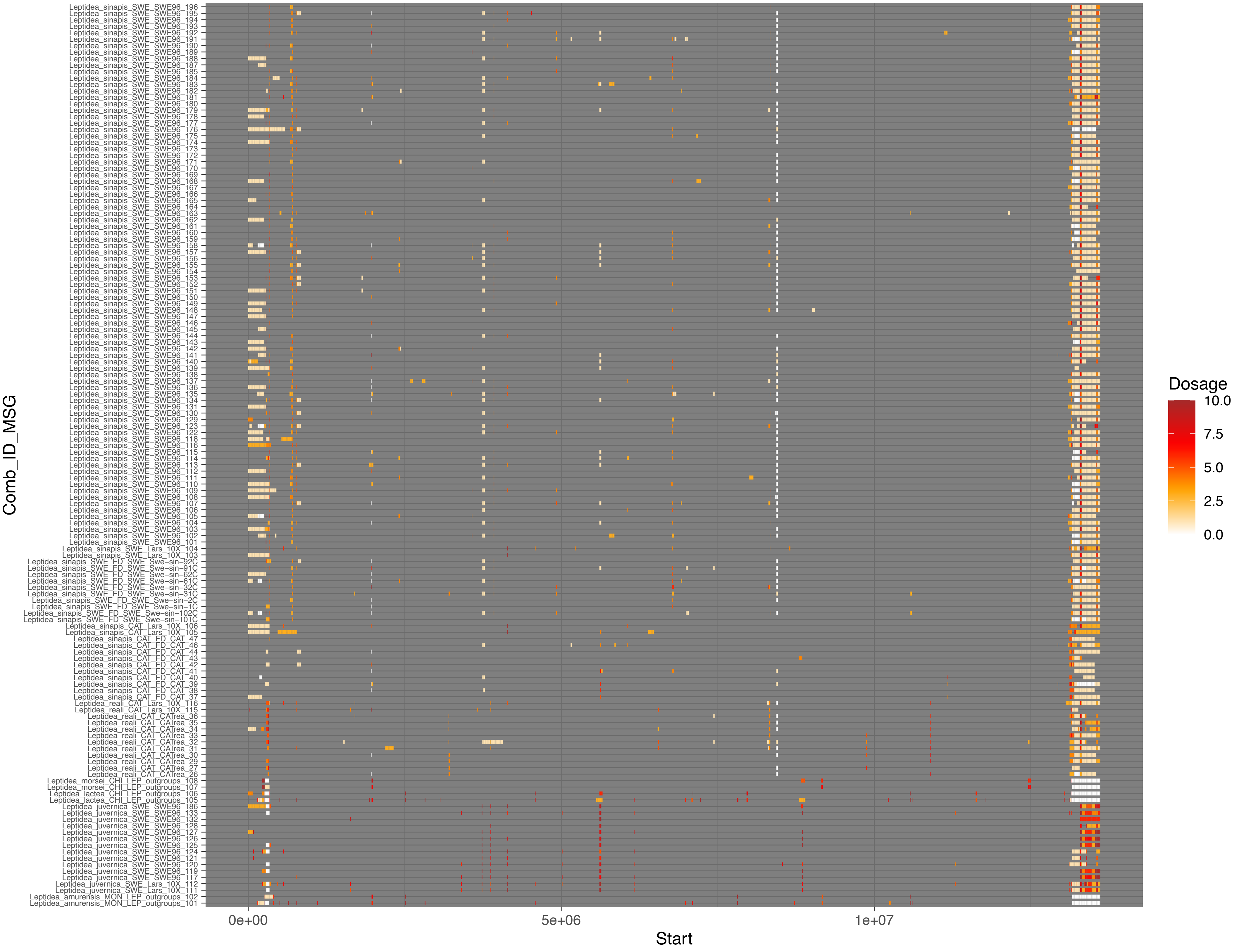

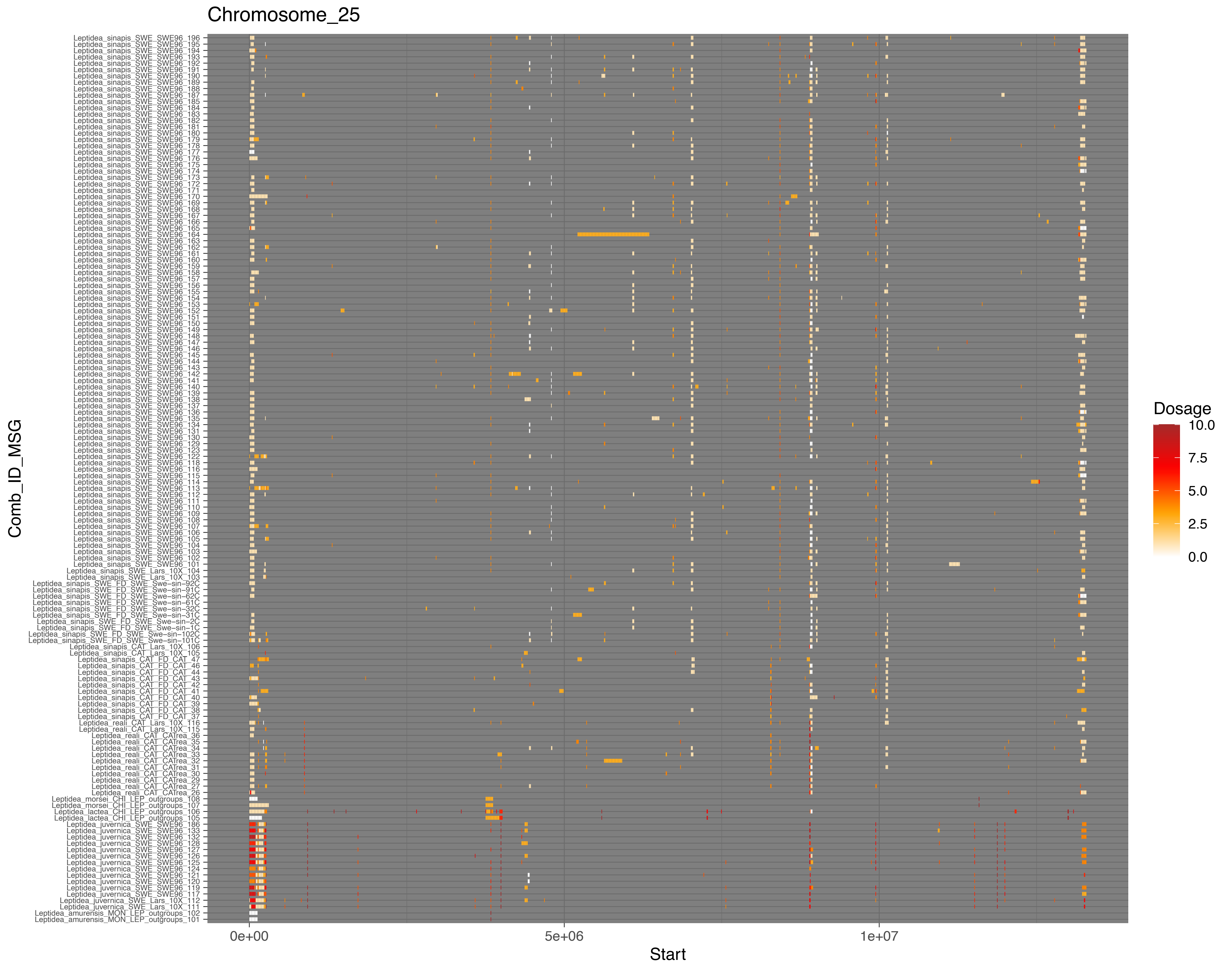
Chromosome_25
Comb_ID_MSG
Start
0e+00
5e+06
1e+07
Dosage
10.0
7.5
5.0
2.5
0.0

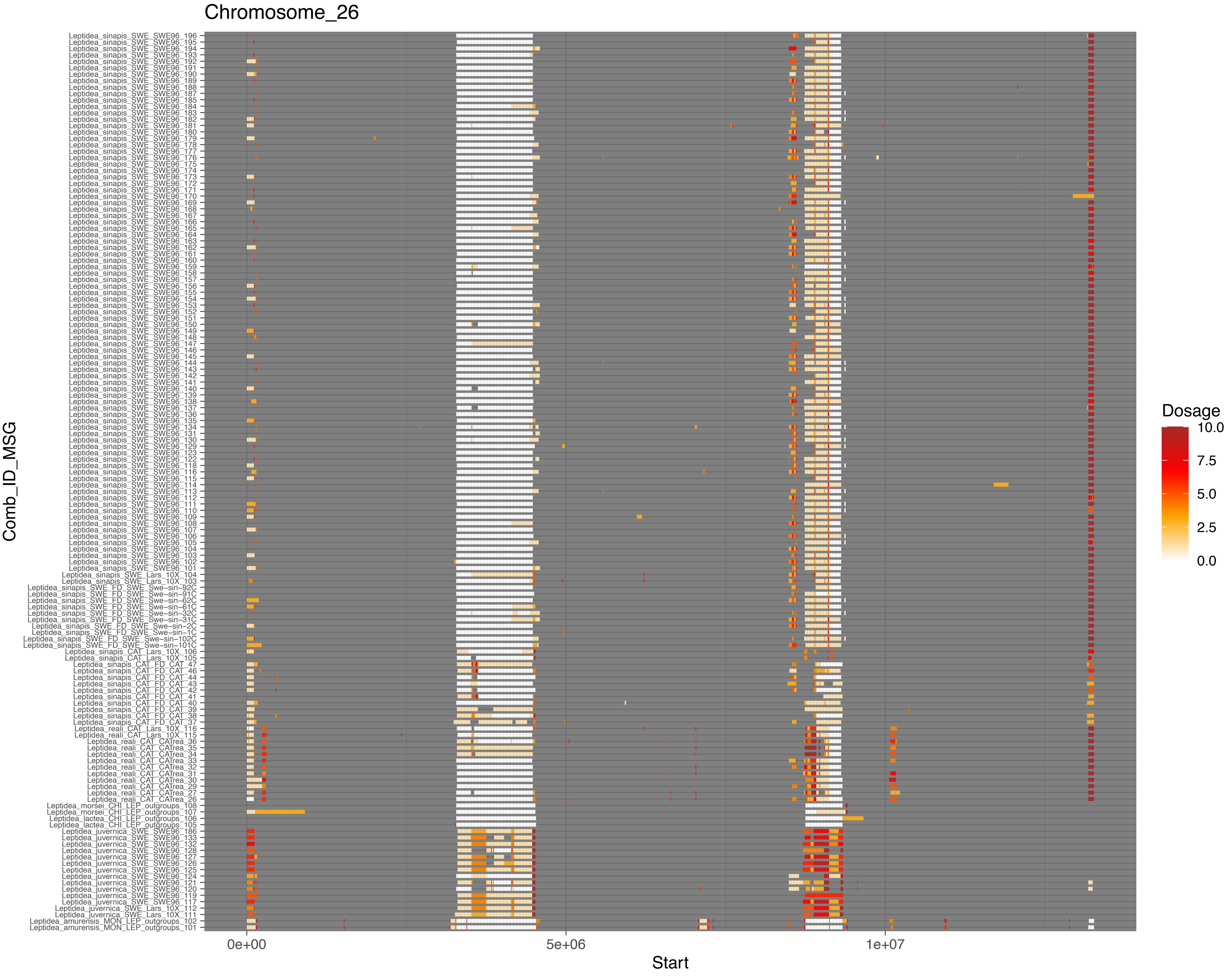

Chromosome_26
Comb_ID_MSG
Leptidea_sinapis_SWE_SWE96_196
Leptidea_sinapis_SWE_SWE96_195
Leptidea_sinapis_SWE_SWE96_194
Leptidea_sinapis_SWE_SWE96_193
Leptidea_sinapis_SWE_SWE96_192
Leptidea_sinapis_SWE_SWE96_191
Leptidea_sinapis_SWE_SWE96_190
Leptidea_sinapis_SWE_SWE96_189
Leptidea_sinapis_SWE_SWE96_188
Leptidea_sinapis_SWE_SWE96_187
Leptidea_sinapis_SWE_SWE96_185
Leptidea_sinapis_SWE_SWE96_184
Leptidea_sinapis_SWE_SWE96_183
Leptidea_sinapis_SWE_SWE96_182
Leptidea_sinapis_SWE_SWE96_181
Leptidea_sinapis_SWE_SWE96_180
Leptidea_sinapis_SWE_SWE96_179
Leptidea_sinapis_SWE_SWE96_178
Leptidea_sinapis_SWE_SWE96_177
Leptidea_sinapis_SWE_SWE96_176
Leptidea_sinapis_SWE_SWE96_175
Leptidea_sinapis_SWE_SWE96_174
Leptidea_sinapis_SWE_SWE96_173
Leptidea_sinapis_SWE_SWE96_172
Leptidea_sinapis_SWE_SWE96_171
Leptidea_sinapis_SWE_SWE96_170
Leptidea_sinapis_SWE_SWE96_169
Leptidea_sinapis_SWE_SWE96_168
Leptidea_sinapis_SWE_SWE96_167
Leptidea_sinapis_SWE_SWE96_166
Leptidea_sinapis_SWE_SWE96_165
Leptidea_sinapis_SWE_SWE96_164
Leptidea_sinapis_SWE_SWE96_163
Leptidea_sinapis_SWE_SWE96_162
Leptidea_sinapis_SWE_SWE96_161
Leptidea_sinapis_SWE_SWE96_160
Leptidea_sinapis_SWE_SWE96_159
Leptidea_sinapis_SWE_SWE96_158
Leptidea_sinapis_SWE_SWE96_157
Leptidea_sinapis_SWE_SWE96_156
Leptidea_sinapis_SWE_SWE96_155
Leptidea_sinapis_SWE_SWE96_154
Leptidea_sinapis_SWE_SWE96_153
Leptidea_sinapis_SWE_SWE96_152
Leptidea_sinapis_SWE_SWE96_151
Leptidea_sinapis_SWE_SWE96_150
Leptidea_sinapis_SWE_SWE96_149
Leptidea_sinapis_SWE_SWE96_148
Leptidea_sinapis_SWE_SWE96_147
Leptidea_sinapis_SWE_SWE96_146
Leptidea_sinapis_SWE_SWE96_145
Leptidea_sinapis_SWE_SWE96_144
Leptidea_sinapis_SWE_SWE96_143
Leptidea_sinapis_SWE_SWE96_142
Leptidea_sinapis_SWE_SWE96_141
Leptidea_sinapis_SWE_SWE96_140
Leptidea_sinapis_SWE_SWE96_139
Leptidea_sinapis_SWE_SWE96_138
Leptidea_sinapis_SWE_SWE96_137
Leptidea_sinapis_SWE_SWE96_136
Leptidea_sinapis_SWE_SWE96_135
Leptidea_sinapis_SWE_SWE96_134
Leptidea_sinapis_SWE_SWE96_131
Leptidea_sinapis_SWE_SWE96_130
Leptidea_sinapis_SWE_SWE96_129
Leptidea_sinapis_SWE_SWE96_123
Leptidea_sinapis_SWE_SWE96_122
Leptidea_sinapis_SWE_SWE96_118
Leptidea_sinapis_SWE_SWE96_116
Leptidea_sinapis_SWE_SWE96_115
Leptidea_sinapis_SWE_SWE96_114
Leptidea_sinapis_SWE_SWE96_113
Leptidea_sinapis_SWE_SWE96_112
Leptidea_sinapis_SWE_SWE96_111
Leptidea_sinapis_SWE_SWE96_110
Leptidea_sinapis_SWE_SWE96_109
Leptidea_sinapis_SWE_SWE96_108
Leptidea_sinapis_SWE_SWE96_107
Leptidea_sinapis_SWE_SWE96_106
Leptidea_sinapis_SWE_SWE96_105
Leptidea_sinapis_SWE_SWE96_104
Leptidea_sinapis_SWE_SWE96_103
Leptidea_sinapis_SWE_SWE96_102
Leptidea_sinapis_SWE_SWE96_101
Leptidea_sinapis_SWE_Lars_10X_104
Leptidea_sinapis_SWE_Lars_10X_103
Leptidea_sinapis_SWE_FD_SWE_Swe-sin-92C
Leptidea_sinapis_SWE_FD_SWE_Swe-sin-91C
Leptidea_sinapis_SWE_FD_SWE_Swe-sin-62C
Leptidea_sinapis_SWE_FD_SWE_Swe-sin-61C
Leptidea_sinapis_SWE_FD_SWE_Swe-sin-32C
Leptidea_sinapis_SWE_FD_SWE_Swe-sin-31C
Leptidea_sinapis_SWE_FD_SWE_Swe-sin-2C
Leptidea_sinapis_SWE_FD_SWE_Swe-sin-1C
Leptidea_sinapis_SWE_FD_SWE_Swe-sin-102C
Leptidea_sinapis_SWE_FD_SWE_Swe-sin-101C
Leptidea_sinapis_CAT_Lars_10X_106
Leptidea_sinapis_CAT_Lars_10X_105
Leptidea_sinapis_CAT_FD_CAT_47
Leptidea_sinapis_CAT_FD_CAT_46
Leptidea_sinapis_CAT_FD_CAT_44
Leptidea_sinapis_CAT_FD_CAT_43
Leptidea_sinapis_CAT_FD_CAT_42
Leptidea_sinapis_CAT_FD_CAT_41
Leptidea_sinapis_CAT_FD_CAT_40
Leptidea_sinapis_CAT_FD_CAT_39
Leptidea_sinapis_CAT_FD_CAT_38
Leptidea_sinapis_CAT_FD_CAT_37
Leptidea_reali_CAT_Lars_10X_116
Leptidea_reali_CAT_Lars_10X_115
Leptidea_reali_CAT_CATrea_36
Leptidea_reali_CAT_CATrea_35
Leptidea_reali_CAT_CATrea_34
Leptidea_reali_CAT_CATrea_33
Leptidea_reali_CAT_CATrea_32
Leptidea_reali_CAT_CATrea_31
Leptidea_reali_CAT_CATrea_30
Leptidea_reali_CAT_CATrea_29
Leptidea_reali_CAT_CATrea_27
Leptidea_reali_CAT_CATrea_26
Leptidea_morsei_CHI_LEP_outgroups_108
Leptidea_morsei_CHI_LEP_outgroups_107
Leptidea_lactea_CHI_LEP_outgroups_106
Leptidea_lactea_CHI_LEP_outgroups_105
Leptidea_juvernica_SWE_SWE96_186
Leptidea_juvernica_SWE_SWE96_133
Leptidea_juvernica_SWE_SWE96_132
Leptidea_juvernica_SWE_SWE96_128
Leptidea_juvernica_SWE_SWE96_127
Leptidea_juvernica_SWE_SWE96_126
Leptidea_juvernica_SWE_SWE96_125
Leptidea_juvernica_SWE_SWE96_124
Leptidea_juvernica_SWE_SWE96_121
Leptidea_juvernica_SWE_SWE96_120
Leptidea_juvernica_SWE_SWE96_119
Leptidea_juvernica_SWE_SWE96_117
Leptidea_juvernica_SWE_Lars_10X_112
Leptidea_juvernica_SWE_Lars_10X_111
Leptidea_amurensis_MON_LEP_outgroups_102
Leptidea_amurensis_MON_LEP_outgroups_101
0e+00
5e+06
1e+07
Start
Dosage
10.0
7.5
5.0
2.5
0.0

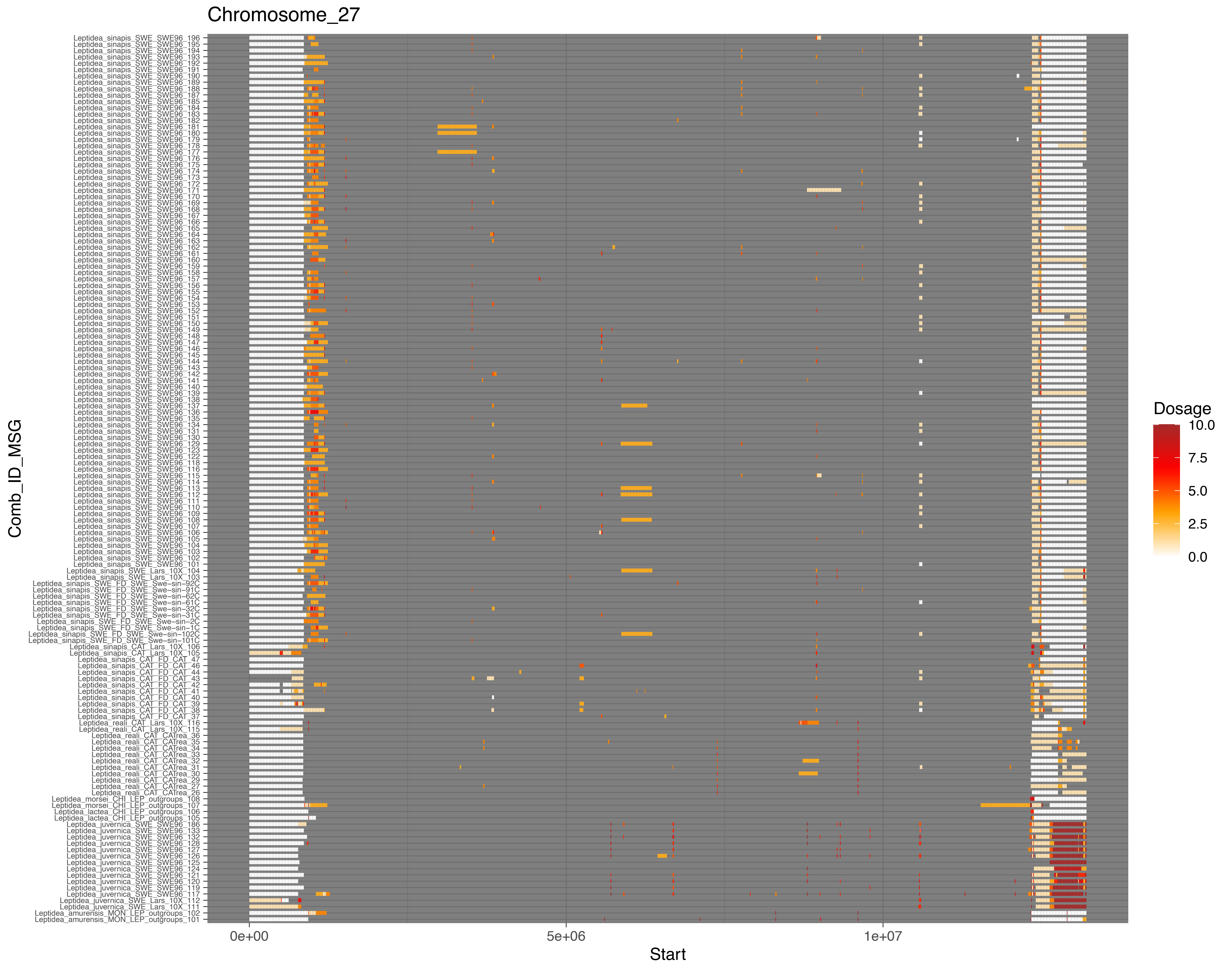

Chromosome_27
Comb_ID_MSG
Start
0e+00
5e+06
1e+07
Dosage
10.0
7.5
5.0
2.5
0.0

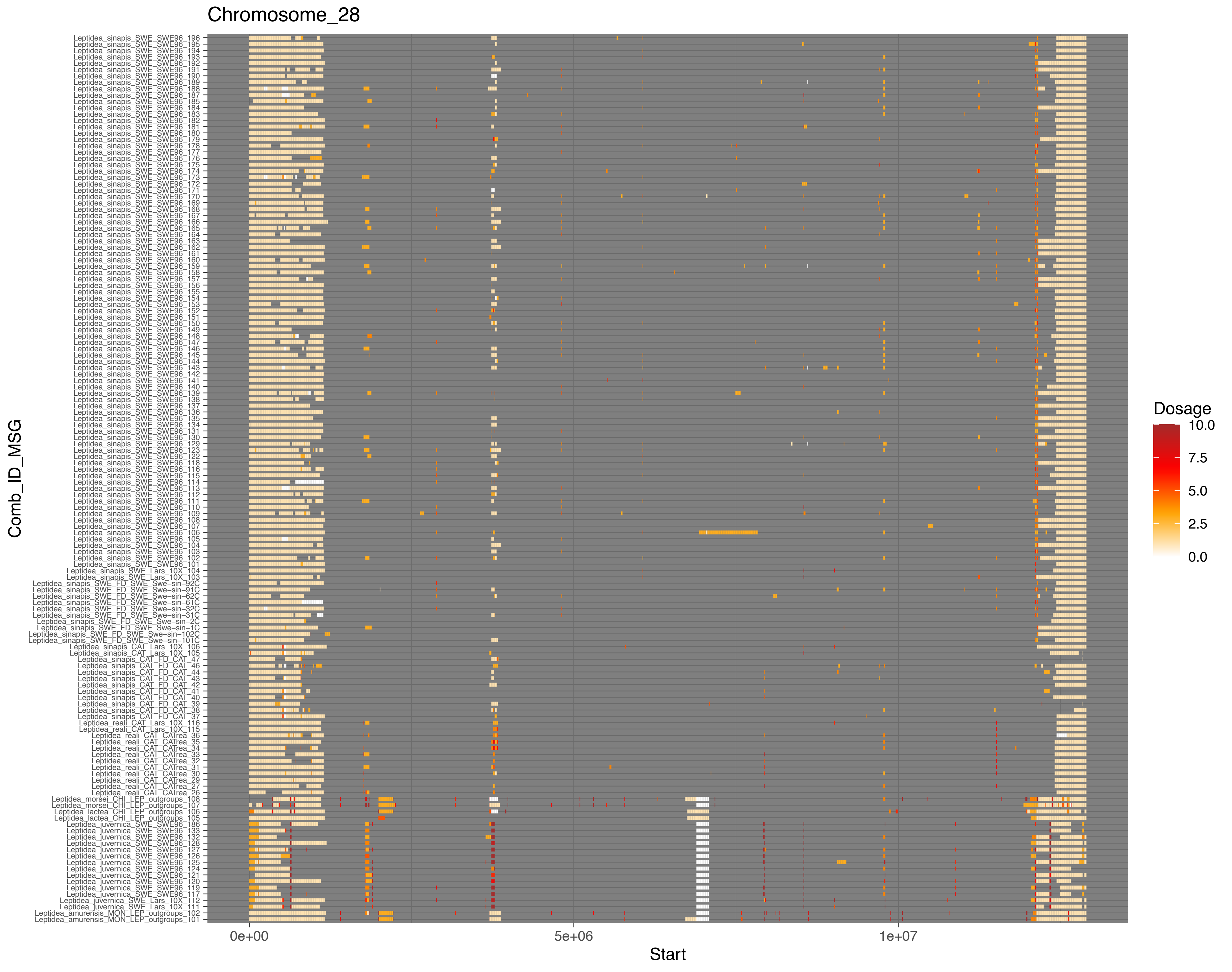
Chromosome_28
Leptidea_sinapis_SWE_SWE96_196
Leptidea_sinapis_SWE_SWE96_195
Leptidea_sinapis_SWE_SWE96_194
Leptidea_sinapis_SWE_SWE96_193
Leptidea_sinapis_SWE_SWE96_192
Leptidea_sinapis_SWE_SWE96_191
Leptidea_sinapis_SWE_SWE96_190
Leptidea_sinapis_SWE_SWE96_189
Leptidea_sinapis_SWE_SWE96_188
Leptidea_sinapis_SWE_SWE96_187
Leptidea_sinapis_SWE_SWE96_185
Leptidea_sinapis_SWE_SWE96_184
Leptidea_sinapis_SWE_SWE96_183
Leptidea_sinapis_SWE_SWE96_182
Leptidea_sinapis_SWE_SWE96_181
Leptidea_sinapis_SWE_SWE96_180
Leptidea_sinapis_SWE_SWE96_179
Leptidea_sinapis_SWE_SWE96_178
Leptidea_sinapis_SWE_SWE96_177
Leptidea_sinapis_SWE_SWE96_176
Leptidea_sinapis_SWE_SWE96_175
Leptidea_sinapis_SWE_SWE96_174
Leptidea_sinapis_SWE_SWE96_173
Leptidea_sinapis_SWE_SWE96_172
Leptidea_sinapis_SWE_SWE96_171
Leptidea_sinapis_SWE_SWE96_170
Leptidea_sinapis_SWE_SWE96_169
Leptidea_sinapis_SWE_SWE96_168
Leptidea_sinapis_SWE_SWE96_167
Leptidea_sinapis_SWE_SWE96_166
Leptidea_sinapis_SWE_SWE96_165
Leptidea_sinapis_SWE_SWE96_164
Leptidea_sinapis_SWE_SWE96_163
Leptidea_sinapis_SWE_SWE96_162
Leptidea_sinapis_SWE_SWE96_161
Leptidea_sinapis_SWE_SWE96_160
Leptidea_sinapis_SWE_SWE96_159
Leptidea_sinapis_SWE_SWE96_158
Leptidea_sinapis_SWE_SWE96_157
Leptidea_sinapis_SWE_SWE96_156
Leptidea_sinapis_SWE_SWE96_155
Leptidea_sinapis_SWE_SWE96_154
Leptidea_sinapis_SWE_SWE96_153
Leptidea_sinapis_SWE_SWE96_152
Leptidea_sinapis_SWE_SWE96_151
Leptidea_sinapis_SWE_SWE96_150
Leptidea_sinapis_SWE_SWE96_149
Leptidea_sinapis_SWE_SWE96_148
Leptidea_sinapis_SWE_SWE96_147
Leptidea_sinapis_SWE_SWE96_146
Leptidea_sinapis_SWE_SWE96_145
Leptidea_sinapis_SWE_SWE96_144
Leptidea_sinapis_SWE_SWE96_143
Leptidea_sinapis_SWE_SWE96_142
Leptidea_sinapis_SWE_SWE96_141
Leptidea_sinapis_SWE_SWE96_140
Leptidea_sinapis_SWE_SWE96_139
Leptidea_sinapis_SWE_SWE96_138
Leptidea_sinapis_SWE_SWE96_137
Leptidea_sinapis_SWE_SWE96_136
Leptidea_sinapis_SWE_SWE96_135
Leptidea_sinapis_SWE_SWE96_134
Leptidea_sinapis_SWE_SWE96_131
Leptidea_sinapis_SWE_SWE96_130
Leptidea_sinapis_SWE_SWE96_129
Leptidea_sinapis_SWE_SWE96_123
Leptidea_sinapis_SWE_SWE96_122
Leptidea_sinapis_SWE_SWE96_118
Leptidea_sinapis_SWE_SWE96_116
Leptidea_sinapis_SWE_SWE96_115
Leptidea_sinapis_SWE_SWE96_114
Leptidea_sinapis_SWE_SWE96_113
Leptidea_sinapis_SWE_SWE96_112
Leptidea_sinapis_SWE_SWE96_111
Leptidea_sinapis_SWE_SWE96_110
Leptidea_sinapis_SWE_SWE96_109
Leptidea_sinapis_SWE_SWE96_108
Leptidea_sinapis_SWE_SWE96_107
Leptidea_sinapis_SWE_SWE96_106
Leptidea_sinapis_SWE_SWE96_105
Leptidea_sinapis_SWE_SWE96_104
Leptidea_sinapis_SWE_SWE96_103
Leptidea_sinapis_SWE_SWE96_102
Leptidea_sinapis_SWE_SWE96_101
Leptidea_sinapis_SWE_Lars_10X_104
Leptidea_sinapis_SWE_Lars_10X_103
Leptidea_sinapis_SWE_FD_SWE_Swe−sin−92C
Leptidea_sinapis_SWE_FD_SWE_Swe−sin−91C
Leptidea_sinapis_SWE_FD_SWE_Swe−sin−62C
Leptidea_sinapis_SWE_FD_SWE_Swe−sin−61C
Leptidea_sinapis_SWE_FD_SWE_Swe−sin−32C
Leptidea_sinapis_SWE_FD_SWE_Swe−sin−31C
Leptidea_sinapis_SWE_FD_SWE_Swe−sin−2C
Leptidea_sinapis_SWE_FD_SWE_Swe−sin−1C
Leptidea_sinapis_SWE_FD_SWE_Swe−sin−102C
Leptidea_sinapis_SWE_FD_SWE_Swe−sin−101C
Leptidea_sinapis_CAT_Lars_10X_106
Leptidea_sinapis_CAT_Lars_10X_105
Leptidea_sinapis_CAT_FD_CAT_47
Leptidea_sinapis_CAT_FD_CAT_46
Leptidea_sinapis_CAT_FD_CAT_44
Leptidea_sinapis_CAT_FD_CAT_43
Leptidea_sinapis_CAT_FD_CAT_42
Leptidea_sinapis_CAT_FD_CAT_41
Leptidea_sinapis_CAT_FD_CAT_40
Leptidea_sinapis_CAT_FD_CAT_39
Leptidea_sinapis_CAT_FD_CAT_38
Leptidea_sinapis_CAT_FD_CAT_37
Leptidea_reali_CAT_Lars_10X_116
Leptidea_reali_CAT_Lars_10X_115
Leptidea_reali_CAT_CATrea_36
Leptidea_reali_CAT_CATrea_35
Leptidea_reali_CAT_CATrea_34
Leptidea_reali_CAT_CATrea_33
Leptidea_reali_CAT_CATrea_32
Leptidea_reali_CAT_CATrea_31
Leptidea_reali_CAT_CATrea_30
Leptidea_reali_CAT_CATrea_29
Leptidea_reali_CAT_CATrea_27
Leptidea_reali_CAT_CATrea_26
Leptidea_morsei_CHI_LEP_outgroups_108
Leptidea_morsei_CHI_LEP_outgroups_107
Leptidea_lactea_CHI_LEP_outgroups_106
Leptidea_lactea_CHI_LEP_outgroups_105
Leptidea_juvernica_SWE_SWE96_186
Leptidea_juvernica_SWE_SWE96_133
Leptidea_juvernica_SWE_SWE96_132
Leptidea_juvernica_SWE_SWE96_128
Leptidea_juvernica_SWE_SWE96_127
Leptidea_juvernica_SWE_SWE96_126
Leptidea_juvernica_SWE_SWE96_125
Leptidea_juvernica_SWE_SWE96_124
Leptidea_juvernica_SWE_SWE96_121
Leptidea_juvernica_SWE_SWE96_120
Leptidea_juvernica_SWE_SWE96_119
Leptidea_juvernica_SWE_SWE96_117
Leptidea_juvernica_SWE_Lars_10X_112
Leptidea_juvernica_SWE_Lars_10X_111
Leptidea_amurensis_MON_LEP_outgroups_102
Leptidea_amurensis_MON_LEP_outgroups_101
Comb_ID_MSG
0e+00
5e+06
1e+07
Start
Dosage
10.0
7.5
5.0
2.5
0.0

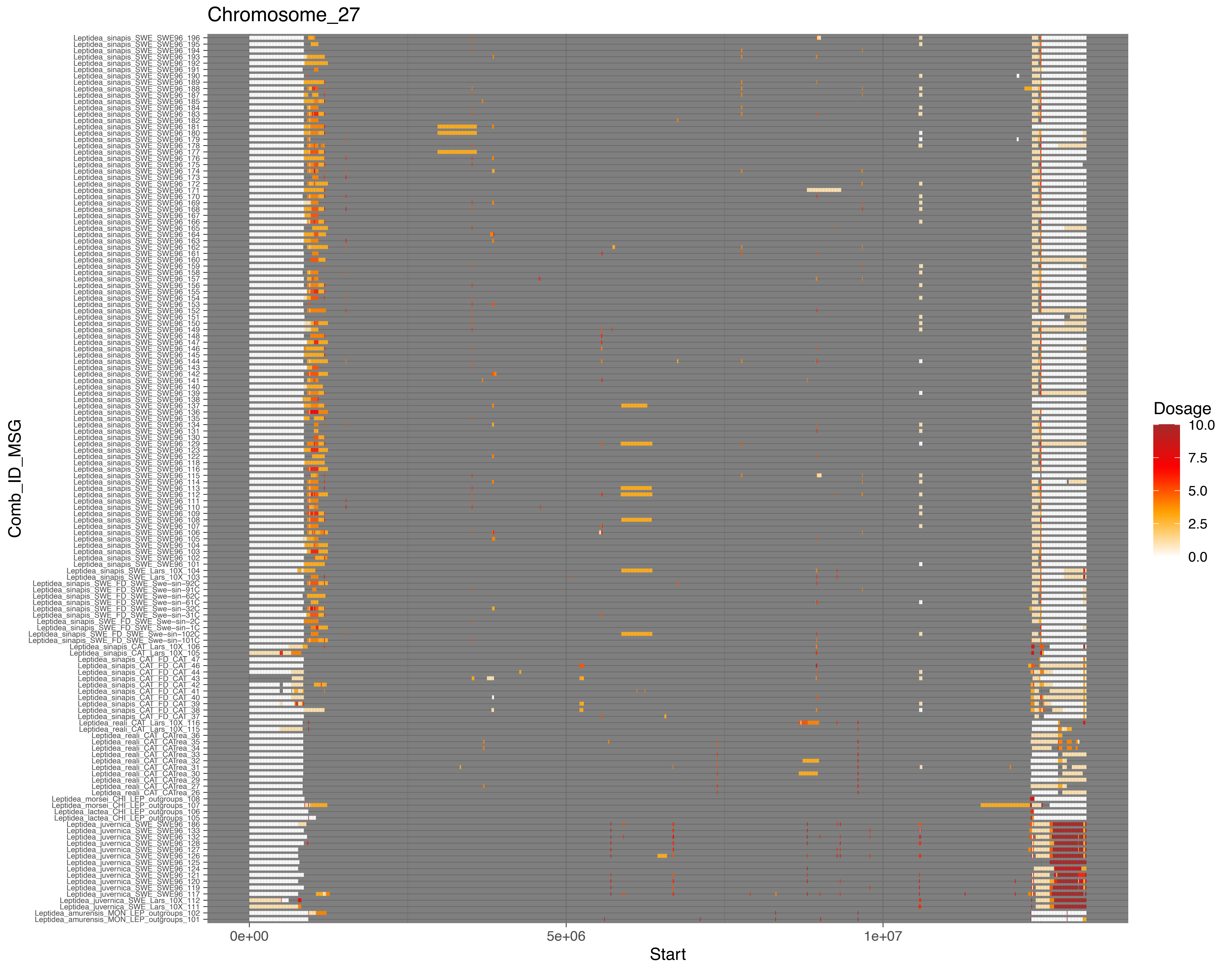
Chromosome_27
Comb_ID_MSG
Start
0e+00
5e+06
1e+07
Dosage
10.0
7.5
5.0
2.5
0.0

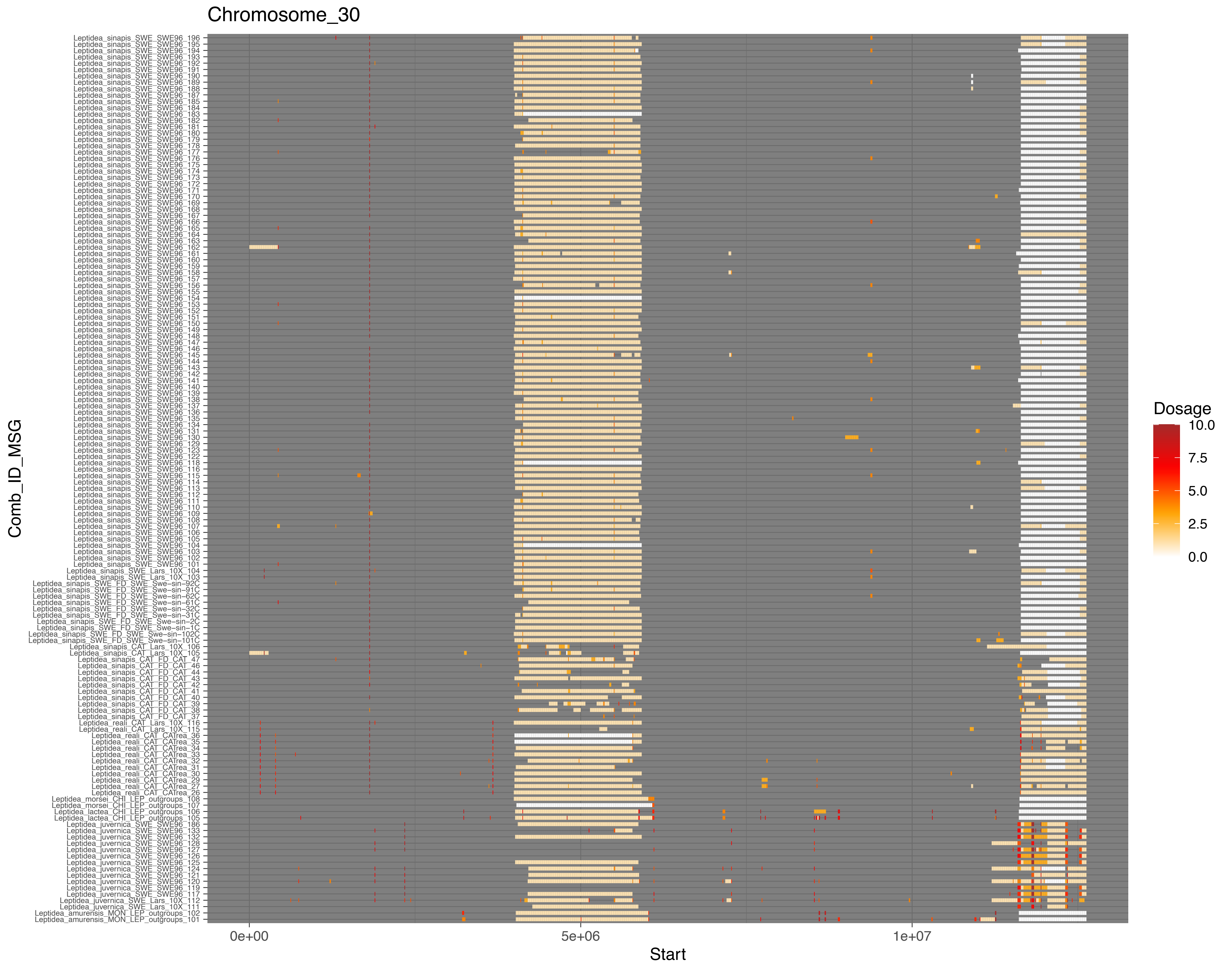
Chromosome_30
Comb_ID_MSG
Start
0e+00
5e+06
1e+07
Dosage
10.0
7.5
5.0
2.5
0.0

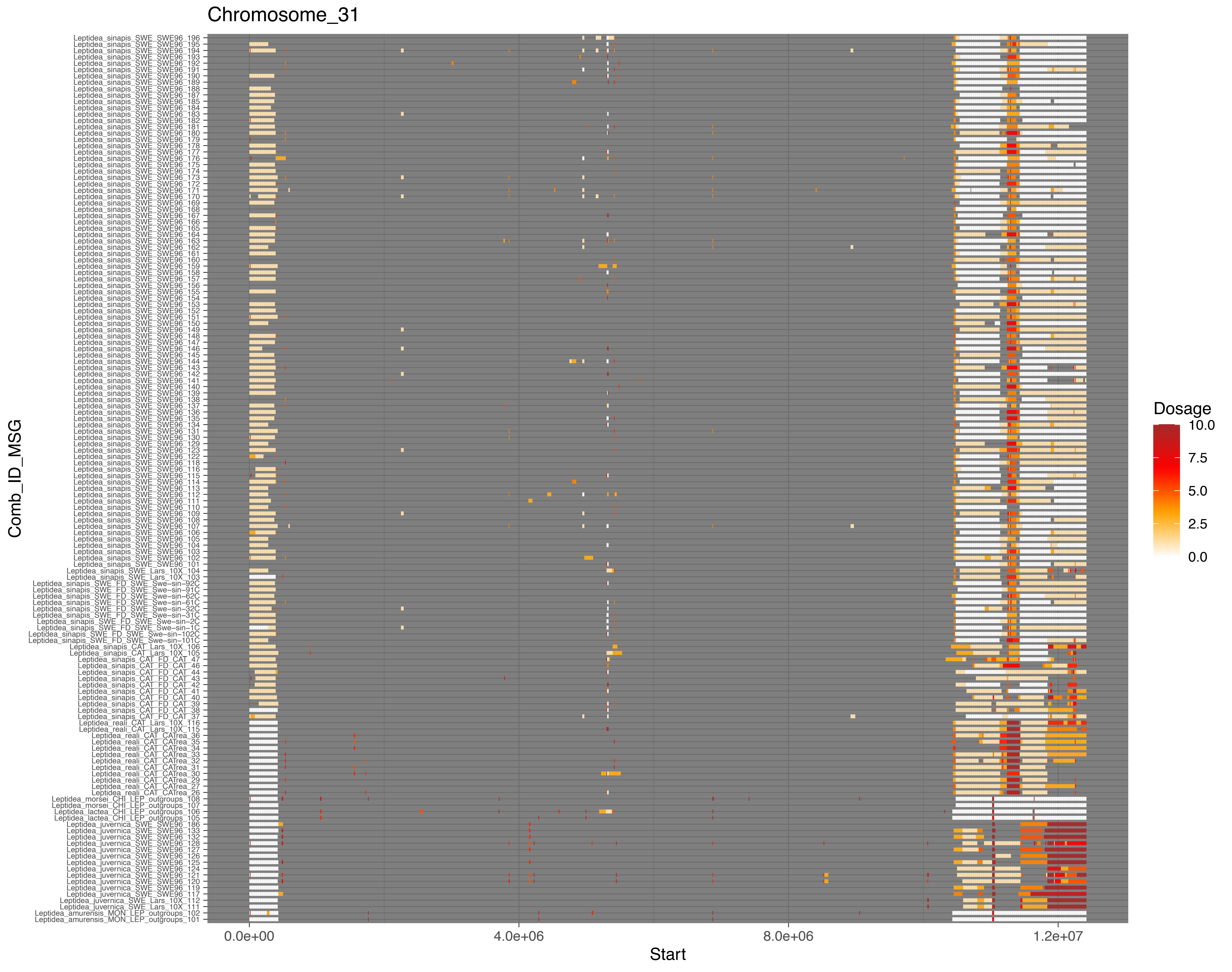

Chromosome_31
Leptidea_sinapis_SWE_SWE96_196
Leptidea_sinapis_SWE_SWE96_195
Leptidea_sinapis_SWE_SWE96_194
Leptidea_sinapis_SWE_SWE96_193
Leptidea_sinapis_SWE_SWE96_192
Leptidea_sinapis_SWE_SWE96_191
Leptidea_sinapis_SWE_SWE96_190
Leptidea_sinapis_SWE_SWE96_189
Leptidea_sinapis_SWE_SWE96_188
Leptidea_sinapis_SWE_SWE96_187
Leptidea_sinapis_SWE_SWE96_185
Leptidea_sinapis_SWE_SWE96_184
Leptidea_sinapis_SWE_SWE96_183
Leptidea_sinapis_SWE_SWE96_182
Leptidea_sinapis_SWE_SWE96_181
Leptidea_sinapis_SWE_SWE96_180
Leptidea_sinapis_SWE_SWE96_179
Leptidea_sinapis_SWE_SWE96_178
Leptidea_sinapis_SWE_SWE96_177
Leptidea_sinapis_SWE_SWE96_176
Leptidea_sinapis_SWE_SWE96_175
Leptidea_sinapis_SWE_SWE96_174
Leptidea_sinapis_SWE_SWE96_173
Leptidea_sinapis_SWE_SWE96_172
Leptidea_sinapis_SWE_SWE96_171
Leptidea_sinapis_SWE_SWE96_170
Leptidea_sinapis_SWE_SWE96_169
Leptidea_sinapis_SWE_SWE96_168
Leptidea_sinapis_SWE_SWE96_167
Leptidea_sinapis_SWE_SWE96_166
Leptidea_sinapis_SWE_SWE96_165
Leptidea_sinapis_SWE_SWE96_164
Leptidea_sinapis_SWE_SWE96_163
Leptidea_sinapis_SWE_SWE96_162
Leptidea_sinapis_SWE_SWE96_161
Leptidea_sinapis_SWE_SWE96_160
Leptidea_sinapis_SWE_SWE96_159
Leptidea_sinapis_SWE_SWE96_158
Leptidea_sinapis_SWE_SWE96_157
Leptidea_sinapis_SWE_SWE96_156
Leptidea_sinapis_SWE_SWE96_155
Leptidea_sinapis_SWE_SWE96_154
Leptidea_sinapis_SWE_SWE96_153
Leptidea_sinapis_SWE_SWE96_152
Leptidea_sinapis_SWE_SWE96_151
Leptidea_sinapis_SWE_SWE96_150
Leptidea_sinapis_SWE_SWE96_149
Leptidea_sinapis_SWE_SWE96_148
Leptidea_sinapis_SWE_SWE96_147
Leptidea_sinapis_SWE_SWE96_146
Leptidea_sinapis_SWE_SWE96_145
Leptidea_sinapis_SWE_SWE96_144
Leptidea_sinapis_SWE_SWE96_143
Leptidea_sinapis_SWE_SWE96_142
Leptidea_sinapis_SWE_SWE96_141
Leptidea_sinapis_SWE_SWE96_140
Leptidea_sinapis_SWE_SWE96_139
Leptidea_sinapis_SWE_SWE96_138
Leptidea_sinapis_SWE_SWE96_137
Leptidea_sinapis_SWE_SWE96_136
Leptidea_sinapis_SWE_SWE96_135
Leptidea_sinapis_SWE_SWE96_134
Leptidea_sinapis_SWE_SWE96_131
Leptidea_sinapis_SWE_SWE96_130
Leptidea_sinapis_SWE_SWE96_129
Leptidea_sinapis_SWE_SWE96_123
Leptidea_sinapis_SWE_SWE96_122
Leptidea_sinapis_SWE_SWE96_118
Leptidea_sinapis_SWE_SWE96_116
Leptidea_sinapis_SWE_SWE96_115
Leptidea_sinapis_SWE_SWE96_114
Leptidea_sinapis_SWE_SWE96_113
Leptidea_sinapis_SWE_SWE96_112
Leptidea_sinapis_SWE_SWE96_111
Leptidea_sinapis_SWE_SWE96_110
Leptidea_sinapis_SWE_SWE96_109
Leptidea_sinapis_SWE_SWE96_108
Leptidea_sinapis_SWE_SWE96_107
Leptidea_sinapis_SWE_SWE96_106
Leptidea_sinapis_SWE_SWE96_105
Leptidea_sinapis_SWE_SWE96_104
Leptidea_sinapis_SWE_SWE96_103
Leptidea_sinapis_SWE_SWE96_102
Leptidea_sinapis_SWE_SWE96_101
Leptidea_sinapis_SWE_Lars_10X_104
Leptidea_sinapis_SWE_Lars_10X_103
Leptidea_sinapis_SWE_FD_SWE_Swe-sin-92C
Leptidea_sinapis_SWE_FD_SWE_Swe-sin-91C
Leptidea_sinapis_SWE_FD_SWE_Swe-sin-62C
Leptidea_sinapis_SWE_FD_SWE_Swe-sin-61C
Leptidea_sinapis_SWE_FD_SWE_Swe-sin-32C
Leptidea_sinapis_SWE_FD_SWE_Swe-sin-31C
Leptidea_sinapis_SWE_FD_SWE_Swe-sin-2C
Leptidea_sinapis_SWE_FD_SWE_Swe-sin-1C
Leptidea_sinapis_SWE_FD_SWE_Swe-sin-102C
Leptidea_sinapis_SWE_FD_SWE_Swe-sin-101C
Leptidea_sinapis_CAT_Lars_10X_106
Leptidea_sinapis_CAT_Lars_10X_105
Leptidea_sinapis_CAT_FD_CAT_47
Leptidea_sinapis_CAT_FD_CAT_46
Leptidea_sinapis_CAT_FD_CAT_44
Leptidea_sinapis_CAT_FD_CAT_43
Leptidea_sinapis_CAT_FD_CAT_42
Leptidea_sinapis_CAT_FD_CAT_41
Leptidea_sinapis_CAT_FD_CAT_40
Leptidea_sinapis_CAT_FD_CAT_39
Leptidea_sinapis_CAT_FD_CAT_38
Leptidea_sinapis_CAT_FD_CAT_37
Leptidea_reali_CAT_Lars_10X_116
Leptidea_reali_CAT_Lars_10X_115
Leptidea_reali_CAT_CATrea_36
Leptidea_reali_CAT_CATrea_35
Leptidea_reali_CAT_CATrea_34
Leptidea_reali_CAT_CATrea_33
Leptidea_reali_CAT_CATrea_32
Leptidea_reali_CAT_CATrea_31
Leptidea_reali_CAT_CATrea_30
Leptidea_reali_CAT_CATrea_29
Leptidea_reali_CAT_CATrea_27
Leptidea_reali_CAT_CATrea_26
Leptidea_morsei_CHI_LEP_outgroups_108
Leptidea_morsei_CHI_LEP_outgroups_107
Leptidea_lactea_CHI_LEP_outgroups_106
Leptidea_lactea_CHI_LEP_outgroups_105
Leptidea_juvernica_SWE_SWE96_186
Leptidea_juvernica_SWE_SWE96_133
Leptidea_juvernica_SWE_SWE96_132
Leptidea_juvernica_SWE_SWE96_128
Leptidea_juvernica_SWE_SWE96_127
Leptidea_juvernica_SWE_SWE96_126
Leptidea_juvernica_SWE_SWE96_125
Leptidea_juvernica_SWE_SWE96_124
Leptidea_juvernica_SWE_SWE96_121
Leptidea_juvernica_SWE_SWE96_120
Leptidea_juvernica_SWE_SWE96_119
Leptidea_juvernica_SWE_SWE96_117
Leptidea_juvernica_SWE_Lars_10X_112
Leptidea_juvernica_SWE_Lars_10X_111
Leptidea_amurensis_MON_LEP_outgroups_102
Leptidea_amurensis_MON_LEP_outgroups_101
Comb_ID_MSG
0.0e+00
4.0e+06
8.0e+06
1.2e+07
Start
Dosage
10.0
7.5
5.0
2.5
0.0

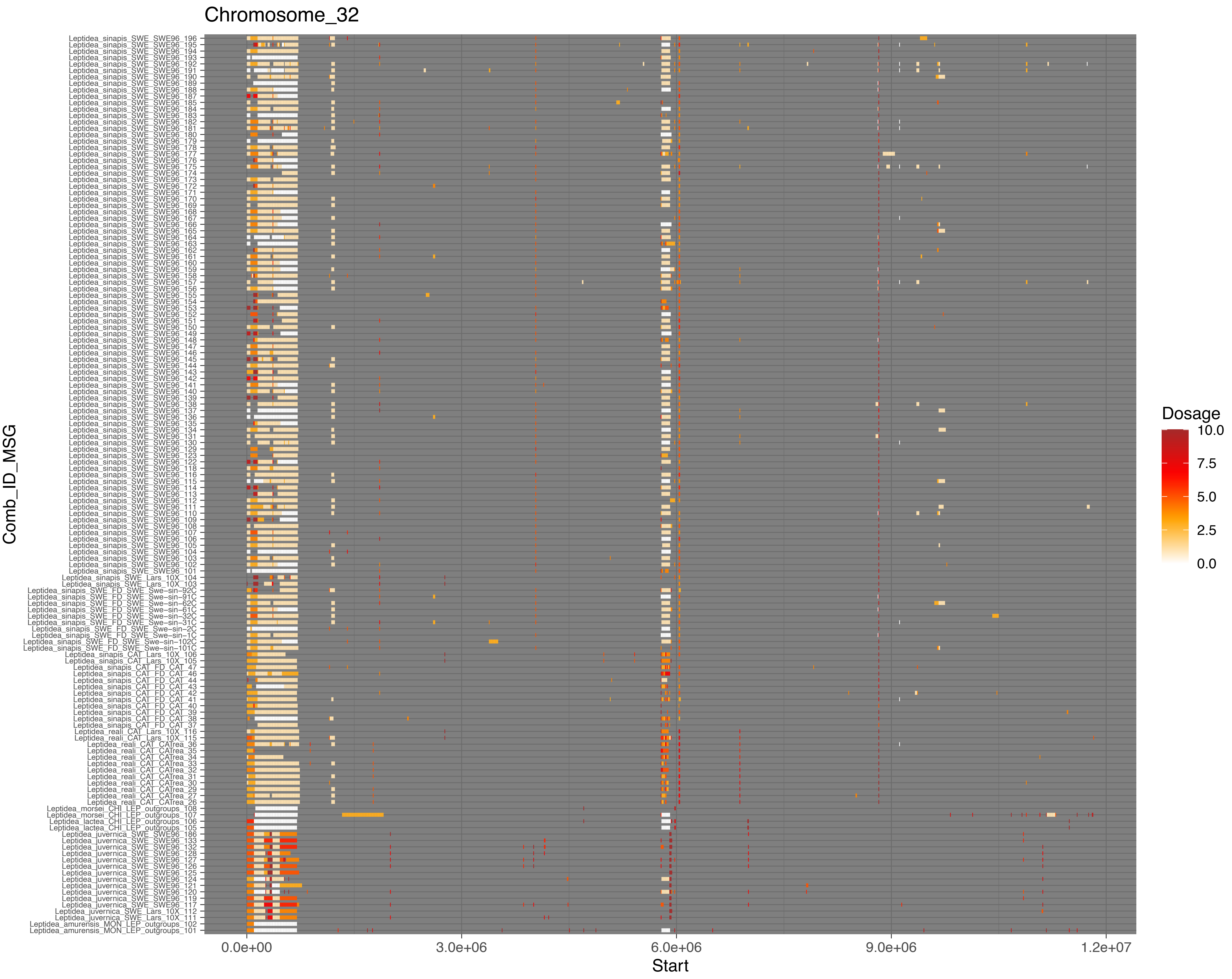

Chromosome_32
Leptidea_sinapis_SWE_SWE96_196
Leptidea_sinapis_SWE_SWE96_195
Leptidea_sinapis_SWE_SWE96_194
Leptidea_sinapis_SWE_SWE96_193
Leptidea_sinapis_SWE_SWE96_192
Leptidea_sinapis_SWE_SWE96_191
Leptidea_sinapis_SWE_SWE96_190
Leptidea_sinapis_SWE_SWE96_189
Leptidea_sinapis_SWE_SWE96_188
Leptidea_sinapis_SWE_SWE96_187
Leptidea_sinapis_SWE_SWE96_185
Leptidea_sinapis_SWE_SWE96_184
Leptidea_sinapis_SWE_SWE96_183
Leptidea_sinapis_SWE_SWE96_182
Leptidea_sinapis_SWE_SWE96_181
Leptidea_sinapis_SWE_SWE96_180
Leptidea_sinapis_SWE_SWE96_179
Leptidea_sinapis_SWE_SWE96_178
Leptidea_sinapis_SWE_SWE96_177
Leptidea_sinapis_SWE_SWE96_176
Leptidea_sinapis_SWE_SWE96_175
Leptidea_sinapis_SWE_SWE96_174
Leptidea_sinapis_SWE_SWE96_173
Leptidea_sinapis_SWE_SWE96_172
Leptidea_sinapis_SWE_SWE96_171
Leptidea_sinapis_SWE_SWE96_170
Leptidea_sinapis_SWE_SWE96_169
Leptidea_sinapis_SWE_SWE96_168
Leptidea_sinapis_SWE_SWE96_167
Leptidea_sinapis_SWE_SWE96_166
Leptidea_sinapis_SWE_SWE96_165
Leptidea_sinapis_SWE_SWE96_164
Leptidea_sinapis_SWE_SWE96_163
Leptidea_sinapis_SWE_SWE96_162
Leptidea_sinapis_SWE_SWE96_161
Leptidea_sinapis_SWE_SWE96_160
Leptidea_sinapis_SWE_SWE96_159
Leptidea_sinapis_SWE_SWE96_158
Leptidea_sinapis_SWE_SWE96_157
Leptidea_sinapis_SWE_SWE96_156
Leptidea_sinapis_SWE_SWE96_155
Leptidea_sinapis_SWE_SWE96_154
Leptidea_sinapis_SWE_SWE96_153
Leptidea_sinapis_SWE_SWE96_152
Leptidea_sinapis_SWE_SWE96_151
Leptidea_sinapis_SWE_SWE96_150
Leptidea_sinapis_SWE_SWE96_149
Leptidea_sinapis_SWE_SWE96_148
Leptidea_sinapis_SWE_SWE96_147
Leptidea_sinapis_SWE_SWE96_146
Leptidea_sinapis_SWE_SWE96_145
Leptidea_sinapis_SWE_SWE96_144
Leptidea_sinapis_SWE_SWE96_143
Leptidea_sinapis_SWE_SWE96_142
Leptidea_sinapis_SWE_SWE96_141
Leptidea_sinapis_SWE_SWE96_140
Leptidea_sinapis_SWE_SWE96_139
Leptidea_sinapis_SWE_SWE96_138
Leptidea_sinapis_SWE_SWE96_137
Leptidea_sinapis_SWE_SWE96_136
Leptidea_sinapis_SWE_SWE96_135
Leptidea_sinapis_SWE_SWE96_134
Leptidea_sinapis_SWE_SWE96_131
Leptidea_sinapis_SWE_SWE96_130
Leptidea_sinapis_SWE_SWE96_129
Leptidea_sinapis_SWE_SWE96_123
Leptidea_sinapis_SWE_SWE96_122
Leptidea_sinapis_SWE_SWE96_118
Leptidea_sinapis_SWE_SWE96_116
Leptidea_sinapis_SWE_SWE96_115
Leptidea_sinapis_SWE_SWE96_114
Leptidea_sinapis_SWE_SWE96_113
Leptidea_sinapis_SWE_SWE96_112
Leptidea_sinapis_SWE_SWE96_111
Leptidea_sinapis_SWE_SWE96_110
Leptidea_sinapis_SWE_SWE96_109
Leptidea_sinapis_SWE_SWE96_108
Leptidea_sinapis_SWE_SWE96_107
Leptidea_sinapis_SWE_SWE96_106
Leptidea_sinapis_SWE_SWE96_105
Leptidea_sinapis_SWE_SWE96_104
Leptidea_sinapis_SWE_SWE96_103
Leptidea_sinapis_SWE_SWE96_102
Leptidea_sinapis_SWE_SWE96_101
Leptidea_sinapis_SWE_Lars_10X_104
Leptidea_sinapis_SWE_Lars_10X_103
Leptidea_sinapis_SWE_FD_SWE_Swe-sin-92C
Leptidea_sinapis_SWE_FD_SWE_Swe-sin-91C
Leptidea_sinapis_SWE_FD_SWE_Swe-sin-62C
Leptidea_sinapis_SWE_FD_SWE_Swe-sin-61C
Leptidea_sinapis_SWE_FD_SWE_Swe-sin-32C
Leptidea_sinapis_SWE_FD_SWE_Swe-sin-31C
Leptidea_sinapis_SWE_FD_SWE_Swe-sin-2C
Leptidea_sinapis_SWE_FD_SWE_Swe-sin-1C
Leptidea_sinapis_SWE_FD_SWE_Swe-sin-102C
Leptidea_sinapis_SWE_FD_SWE_Swe-sin-101C
Leptidea_sinapis_CAT_Lars_10X_106
Leptidea_sinapis_CAT_Lars_10X_105
Leptidea_sinapis_CAT_FD_CAT_47
Leptidea_sinapis_CAT_FD_CAT_46
Leptidea_sinapis_CAT_FD_CAT_44
Leptidea_sinapis_CAT_FD_CAT_43
Leptidea_sinapis_CAT_FD_CAT_42
Leptidea_sinapis_CAT_FD_CAT_41
Leptidea_sinapis_CAT_FD_CAT_40
Leptidea_sinapis_CAT_FD_CAT_39
Leptidea_sinapis_CAT_FD_CAT_38
Leptidea_sinapis_CAT_FD_CAT_37
Leptidea_reali_CAT_Lars_10X_116
Leptidea_reali_CAT_Lars_10X_115
Leptidea_reali_CAT_CATrea_36
Leptidea_reali_CAT_CATrea_35
Leptidea_reali_CAT_CATrea_34
Leptidea_reali_CAT_CATrea_33
Leptidea_reali_CAT_CATrea_32
Leptidea_reali_CAT_CATrea_31
Leptidea_reali_CAT_CATrea_30
Leptidea_reali_CAT_CATrea_29
Leptidea_reali_CAT_CATrea_27
Leptidea_reali_CAT_CATrea_26
Leptidea_morsei_CHI_LEP_outgroups_108
Leptidea_morsei_CHI_LEP_outgroups_107
Leptidea_lactea_CHI_LEP_outgroups_106
Leptidea_lactea_CHI_LEP_outgroups_105
Leptidea_juvernica_SWE_SWE96_186
Leptidea_juvernica_SWE_SWE96_133
Leptidea_juvernica_SWE_SWE96_132
Leptidea_juvernica_SWE_SWE96_128
Leptidea_juvernica_SWE_SWE96_127
Leptidea_juvernica_SWE_SWE96_126
Leptidea_juvernica_SWE_SWE96_125
Leptidea_juvernica_SWE_SWE96_124
Leptidea_juvernica_SWE_SWE96_121
Leptidea_juvernica_SWE_SWE96_120
Leptidea_juvernica_SWE_SWE96_119
Leptidea_juvernica_SWE_SWE96_117
Leptidea_juvernica_SWE_Lars_10X_112
Leptidea_juvernica_SWE_Lars_10X_111
Leptidea_amurensis_MON_LEP_outgroups_102
Leptidea_amurensis_MON_LEP_outgroups_101
Comb_ID_MSG
0.0e+00
3.0e+06
6.0e+06
9.0e+06
1.2e+07
Start
Dosage
10.0
7.5
5.0
2.5
0.0

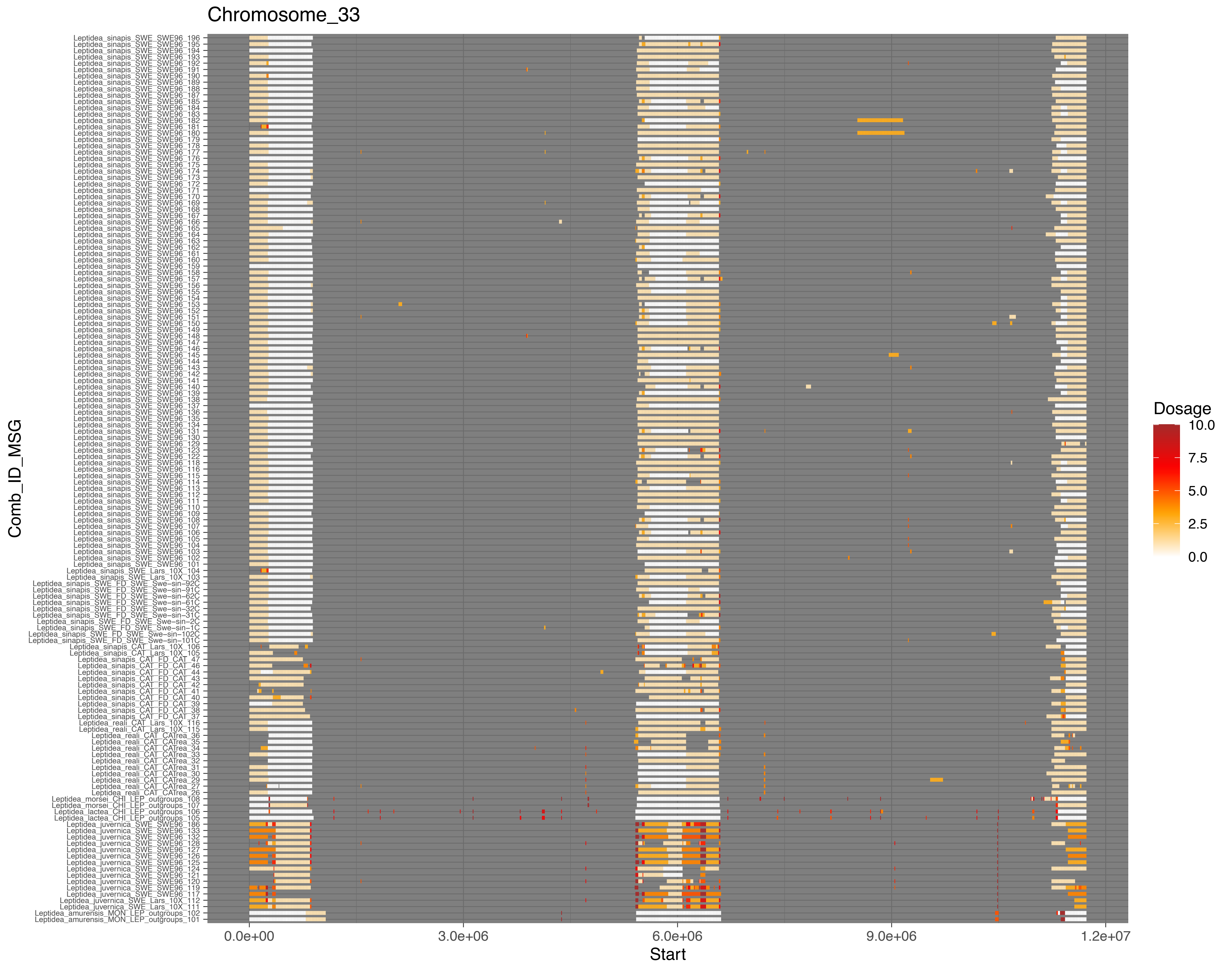
Chromosome_33
Comb_ID_MSG
Leptidea_sinapis_SWE_SWE96_196
Leptidea_sinapis_SWE_SWE96_195
Leptidea_sinapis_SWE_SWE96_194
Leptidea_sinapis_SWE_SWE96_193
Leptidea_sinapis_SWE_SWE96_192
Leptidea_sinapis_SWE_SWE96_191
Leptidea_sinapis_SWE_SWE96_190
Leptidea_sinapis_SWE_SWE96_189
Leptidea_sinapis_SWE_SWE96_188
Leptidea_sinapis_SWE_SWE96_187
Leptidea_sinapis_SWE_SWE96_185
Leptidea_sinapis_SWE_SWE96_184
Leptidea_sinapis_SWE_SWE96_183
Leptidea_sinapis_SWE_SWE96_182
Leptidea_sinapis_SWE_SWE96_181
Leptidea_sinapis_SWE_SWE96_180
Leptidea_sinapis_SWE_SWE96_179
Leptidea_sinapis_SWE_SWE96_178
Leptidea_sinapis_SWE_SWE96_177
Leptidea_sinapis_SWE_SWE96_176
Leptidea_sinapis_SWE_SWE96_175
Leptidea_sinapis_SWE_SWE96_174
Leptidea_sinapis_SWE_SWE96_173
Leptidea_sinapis_SWE_SWE96_172
Leptidea_sinapis_SWE_SWE96_171
Leptidea_sinapis_SWE_SWE96_170
Leptidea_sinapis_SWE_SWE96_169
Leptidea_sinapis_SWE_SWE96_168
Leptidea_sinapis_SWE_SWE96_167
Leptidea_sinapis_SWE_SWE96_166
Leptidea_sinapis_SWE_SWE96_165
Leptidea_sinapis_SWE_SWE96_164
Leptidea_sinapis_SWE_SWE96_163
Leptidea_sinapis_SWE_SWE96_162
Leptidea_sinapis_SWE_SWE96_161
Leptidea_sinapis_SWE_SWE96_160
Leptidea_sinapis_SWE_SWE96_159
Leptidea_sinapis_SWE_SWE96_158
Leptidea_sinapis_SWE_SWE96_157
Leptidea_sinapis_SWE_SWE96_156
Leptidea_sinapis_SWE_SWE96_155
Leptidea_sinapis_SWE_SWE96_154
Leptidea_sinapis_SWE_SWE96_153
Leptidea_sinapis_SWE_SWE96_152
Leptidea_sinapis_SWE_SWE96_151
Leptidea_sinapis_SWE_SWE96_150
Leptidea_sinapis_SWE_SWE96_149
Leptidea_sinapis_SWE_SWE96_148
Leptidea_sinapis_SWE_SWE96_147
Leptidea_sinapis_SWE_SWE96_146
Leptidea_sinapis_SWE_SWE96_145
Leptidea_sinapis_SWE_SWE96_144
Leptidea_sinapis_SWE_SWE96_143
Leptidea_sinapis_SWE_SWE96_142
Leptidea_sinapis_SWE_SWE96_141
Leptidea_sinapis_SWE_SWE96_140
Leptidea_sinapis_SWE_SWE96_139
Leptidea_sinapis_SWE_SWE96_138
Leptidea_sinapis_SWE_SWE96_137
Leptidea_sinapis_SWE_SWE96_136
Leptidea_sinapis_SWE_SWE96_135
Leptidea_sinapis_SWE_SWE96_134
Leptidea_sinapis_SWE_SWE96_131
Leptidea_sinapis_SWE_SWE96_130
Leptidea_sinapis_SWE_SWE96_129
Leptidea_sinapis_SWE_SWE96_123
Leptidea_sinapis_SWE_SWE96_122
Leptidea_sinapis_SWE_SWE96_118
Leptidea_sinapis_SWE_SWE96_116
Leptidea_sinapis_SWE_SWE96_115
Leptidea_sinapis_SWE_SWE96_114
Leptidea_sinapis_SWE_SWE96_113
Leptidea_sinapis_SWE_SWE96_112
Leptidea_sinapis_SWE_SWE96_111
Leptidea_sinapis_SWE_SWE96_110
Leptidea_sinapis_SWE_SWE96_109
Leptidea_sinapis_SWE_SWE96_108
Leptidea_sinapis_SWE_SWE96_107
Leptidea_sinapis_SWE_SWE96_106
Leptidea_sinapis_SWE_SWE96_105
Leptidea_sinapis_SWE_SWE96_104
Leptidea_sinapis_SWE_SWE96_103
Leptidea_sinapis_SWE_SWE96_102
Leptidea_sinapis_SWE_SWE96_101
Leptidea_sinapis_SWE_Lars_10X_104
Leptidea_sinapis_SWE_Lars_10X_103
Leptidea_sinapis_SWE_FD_SWE_Swe-sin-92C
Leptidea_sinapis_SWE_FD_SWE_Swe-sin-91C
Leptidea_sinapis_SWE_FD_SWE_Swe-sin-62C
Leptidea_sinapis_SWE_FD_SWE_Swe-sin-61C
Leptidea_sinapis_SWE_FD_SWE_Swe-sin-32C
Leptidea_sinapis_SWE_FD_SWE_Swe-sin-31C
Leptidea_sinapis_SWE_FD_SWE_Swe-sin-2C
Leptidea_sinapis_SWE_FD_SWE_Swe-sin-1C
Leptidea_sinapis_SWE_FD_SWE_Swe-sin-102C
Leptidea_sinapis_SWE_FD_SWE_Swe-sin-101C
Leptidea_sinapis_CAT_Lars_10X_106
Leptidea_sinapis_CAT_Lars_10X_105
Leptidea_sinapis_CAT_FD_CAT_47
Leptidea_sinapis_CAT_FD_CAT_46
Leptidea_sinapis_CAT_FD_CAT_44
Leptidea_sinapis_CAT_FD_CAT_43
Leptidea_sinapis_CAT_FD_CAT_42
Leptidea_sinapis_CAT_FD_CAT_41
Leptidea_sinapis_CAT_FD_CAT_40
Leptidea_sinapis_CAT_FD_CAT_39
Leptidea_sinapis_CAT_FD_CAT_38
Leptidea_sinapis_CAT_FD_CAT_37
Leptidea_reali_CAT_Lars_10X_116
Leptidea_reali_CAT_Lars_10X_115
Leptidea_reali_CAT_CAtrea_36
Leptidea_reali_CAT_CAtrea_35
Leptidea_reali_CAT_CAtrea_34
Leptidea_reali_CAT_CAtrea_33
Leptidea_reali_CAT_CAtrea_32
Leptidea_reali_CAT_CAtrea_31
Leptidea_reali_CAT_CAtrea_30
Leptidea_reali_CAT_CAtrea_29
Leptidea_reali_CAT_CAtrea_27
Leptidea_reali_CAT_CAtrea_26
Leptidea_morsei_CHI_LEP_outgroups_108
Leptidea_morsei_CHI_LEP_outgroups_107
Leptidea_lactea_CHI_LEP_outgroups_106
Leptidea_lactea_CHI_LEP_outgroups_105
Leptidea_juvernica_SWE_SWE96_186
Leptidea_juvernica_SWE_SWE96_133
Leptidea_juvernica_SWE_SWE96_132
Leptidea_juvernica_SWE_SWE96_128
Leptidea_juvernica_SWE_SWE96_127
Leptidea_juvernica_SWE_SWE96_126
Leptidea_juvernica_SWE_SWE96_125
Leptidea_juvernica_SWE_SWE96_124
Leptidea_juvernica_SWE_SWE96_121
Leptidea_juvernica_SWE_SWE96_120
Leptidea_juvernica_SWE_SWE96_119
Leptidea_juvernica_SWE_SWE96_117
Leptidea_juvernica_SWE_Lars_10X_112
Leptidea_juvernica_SWE_Lars_10X_111
Leptidea_amurensis_MON_LEP_outgroups_102
Leptidea_amurensis_MON_LEP_outgroups_101
0.0e+00
3.0e+06
6.0e+06
9.0e+06
1.2e+07
Start
Dosage
10.0
7.5
5.0
2.5
0.0

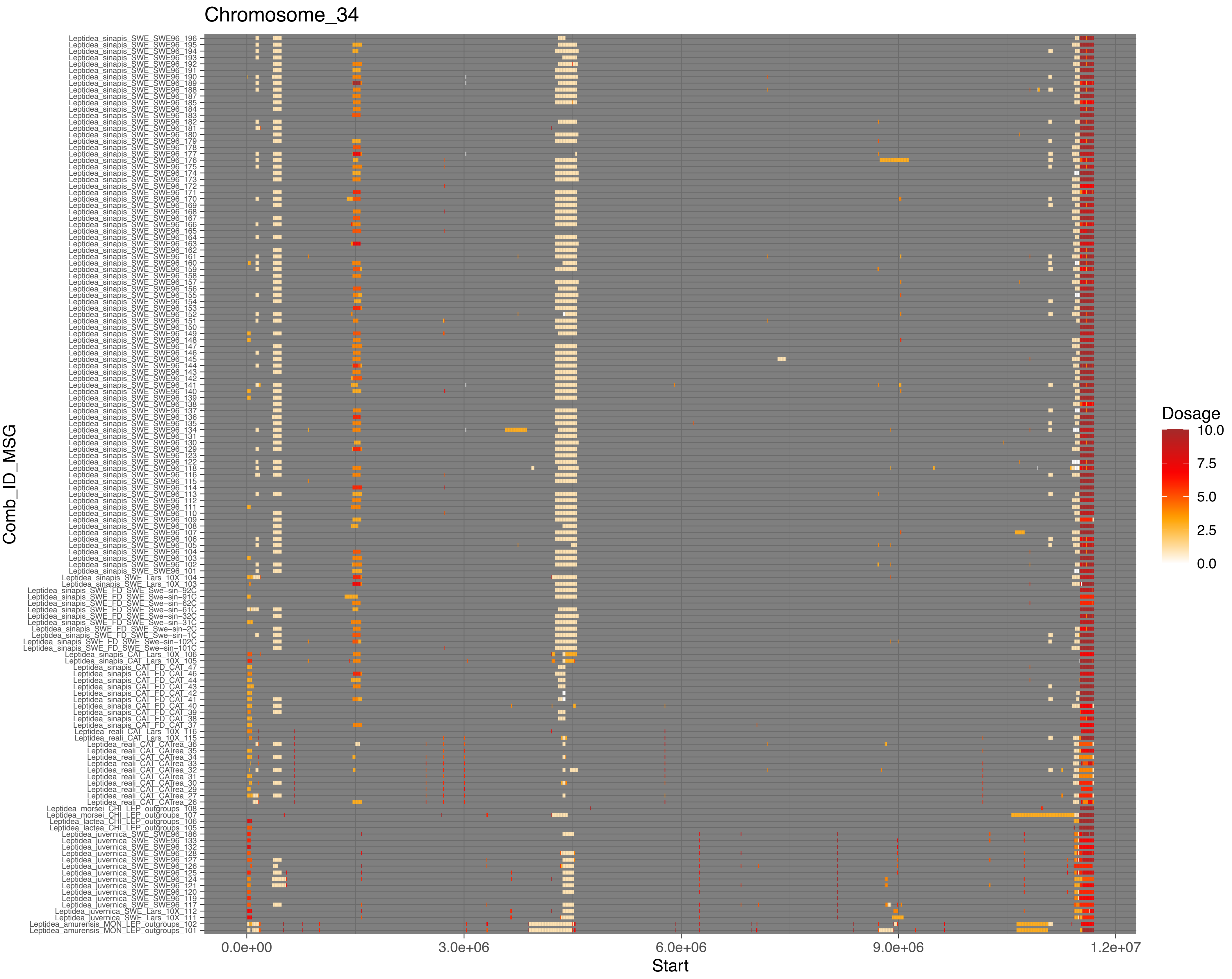

Chromosome_34
Comb_ID_MSG
Start
0.0e+00
3.0e+06
6.0e+06
9.0e+06
1.2e+07
Dosage
10.0
7.5
5.0
2.5
0.0

Chromosome_35

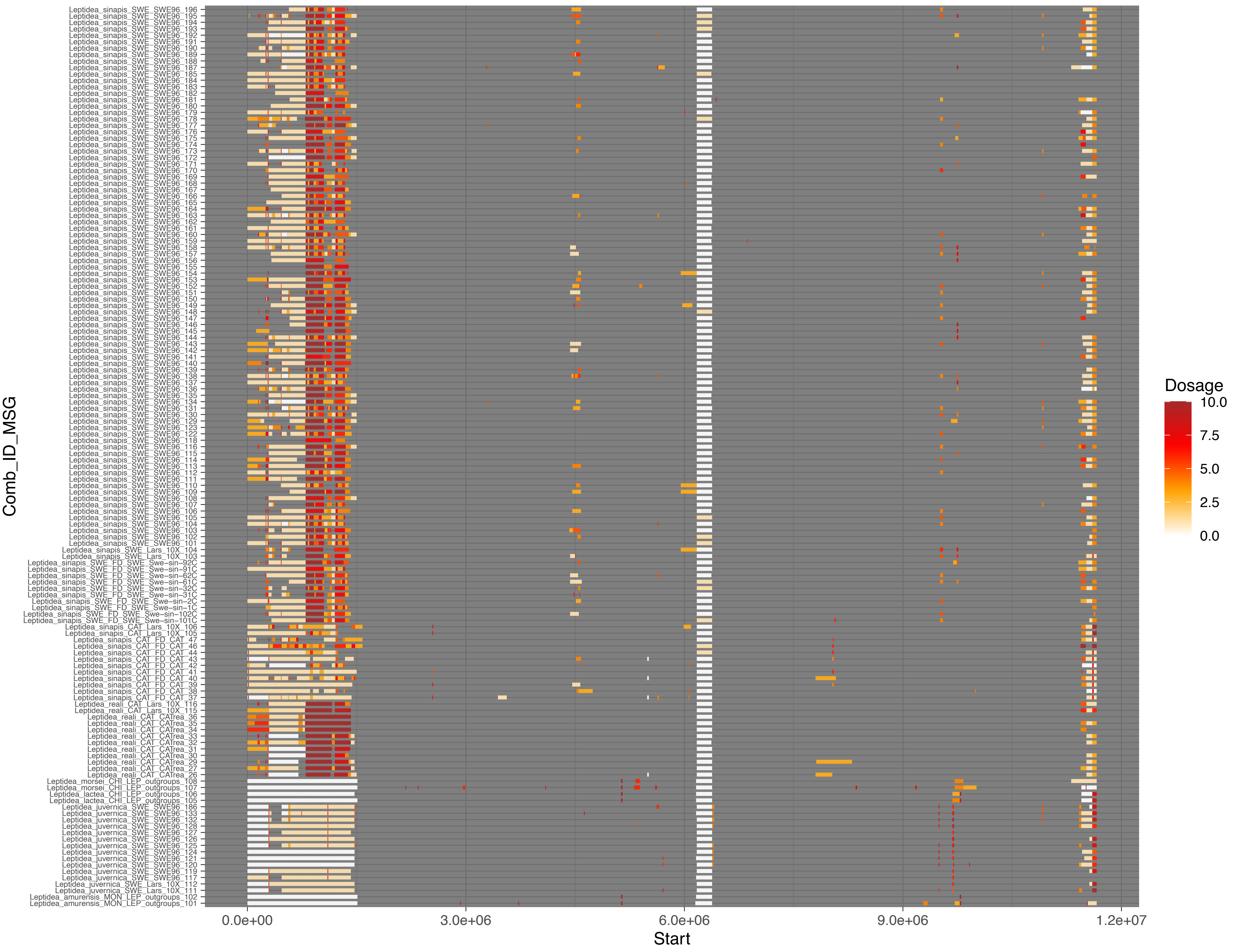

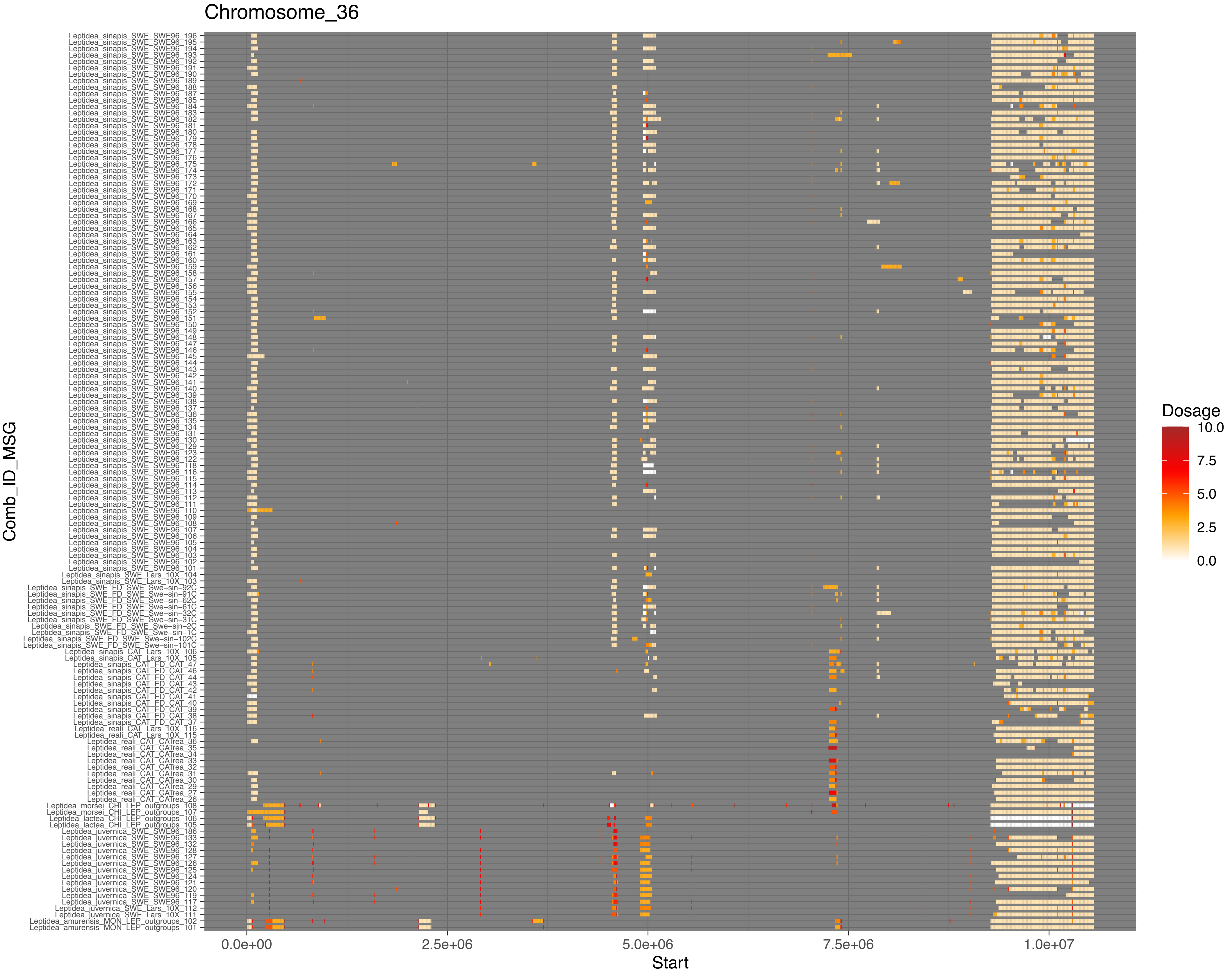
Chromosome_36
Comb_ID_MSG
Start
0.0e+00
2.5e+06
5.0e+06
7.5e+06
1.0e+07
Dosage
10.0
7.5
5.0
2.5
0.0
Leptidea_sinapis_SWE_SWE96_196
Leptidea_sinapis_SWE_SWE96_195
Leptidea_sinapis_SWE_SWE96_194
Leptidea_sinapis_SWE_SWE96_193
Leptidea_sinapis_SWE_SWE96_192
Leptidea_sinapis_SWE_SWE96_191
Leptidea_sinapis_SWE_SWE96_190
Leptidea_sinapis_SWE_SWE96_189
Leptidea_sinapis_SWE_SWE96_188
Leptidea_sinapis_SWE_SWE96_187
Leptidea_sinapis_SWE_SWE96_185
Leptidea_sinapis_SWE_SWE96_184
Leptidea_sinapis_SWE_SWE96_183
Leptidea_sinapis_SWE_SWE96_182
Leptidea_sinapis_SWE_SWE96_181
Leptidea_sinapis_SWE_SWE96_180
Leptidea_sinapis_SWE_SWE96_179
Leptidea_sinapis_SWE_SWE96_178
Leptidea_sinapis_SWE_SWE96_177
Leptidea_sinapis_SWE_SWE96_176
Leptidea_sinapis_SWE_SWE96_175
Leptidea_sinapis_SWE_SWE96_174
Leptidea_sinapis_SWE_SWE96_173
Leptidea_sinapis_SWE_SWE96_172
Leptidea_sinapis_SWE_SWE96_171
Leptidea_sinapis_SWE_SWE96_170
Leptidea_sinapis_SWE_SWE96_169
Leptidea_sinapis_SWE_SWE96_168
Leptidea_sinapis_SWE_SWE96_167
Leptidea_sinapis_SWE_SWE96_166
Leptidea_sinapis_SWE_SWE96_165
Leptidea_sinapis_SWE_SWE96_164
Leptidea_sinapis_SWE_SWE96_163
Leptidea_sinapis_SWE_SWE96_162
Leptidea_sinapis_SWE_SWE96_161
Leptidea_sinapis_SWE_SWE96_160
Leptidea_sinapis_SWE_SWE96_159
Leptidea_sinapis_SWE_SWE96_158
Leptidea_sinapis_SWE_SWE96_157
Leptidea_sinapis_SWE_SWE96_156
Leptidea_sinapis_SWE_SWE96_155
Leptidea_sinapis_SWE_SWE96_154
Leptidea_sinapis_SWE_SWE96_153
Leptidea_sinapis_SWE_SWE96_152
Leptidea_sinapis_SWE_SWE96_151
Leptidea_sinapis_SWE_SWE96_150
Leptidea_sinapis_SWE_SWE96_149
Leptidea_sinapis_SWE_SWE96_148
Leptidea_sinapis_SWE_SWE96_147
Leptidea_sinapis_SWE_SWE96_146
Leptidea_sinapis_SWE_SWE96_145
Leptidea_sinapis_SWE_SWE96_144
Leptidea_sinapis_SWE_SWE96_143
Leptidea_sinapis_SWE_SWE96_142
Leptidea_sinapis_SWE_SWE96_141
Leptidea_sinapis_SWE_SWE96_140
Leptidea_sinapis_SWE_SWE96_139
Leptidea_sinapis_SWE_SWE96_138
Leptidea_sinapis_SWE_SWE96_137
Leptidea_sinapis_SWE_SWE96_136
Leptidea_sinapis_SWE_SWE96_135
Leptidea_sinapis_SWE_SWE96_134
Leptidea_sinapis_SWE_SWE96_131
Leptidea_sinapis_SWE_SWE96_130
Leptidea_sinapis_SWE_SWE96_129
Leptidea_sinapis_SWE_SWE96_123
Leptidea_sinapis_SWE_SWE96_122
Leptidea_sinapis_SWE_SWE96_118
Leptidea_sinapis_SWE_SWE96_116
Leptidea_sinapis_SWE_SWE96_115
Leptidea_sinapis_SWE_SWE96_114
Leptidea_sinapis_SWE_SWE96_113
Leptidea_sinapis_SWE_SWE96_112
Leptidea_sinapis_SWE_SWE96_111
Leptidea_sinapis_SWE_SWE96_110
Leptidea_sinapis_SWE_SWE96_109
Leptidea_sinapis_SWE_SWE96_108
Leptidea_sinapis_SWE_SWE96_107
Leptidea_sinapis_SWE_SWE96_106
Leptidea_sinapis_SWE_SWE96_105
Leptidea_sinapis_SWE_SWE96_104
Leptidea_sinapis_SWE_SWE96_103
Leptidea_sinapis_SWE_SWE96_102
Leptidea_sinapis_SWE_SWE96_101
Leptidea_sinapis_SWE_Lars_10X_104
Leptidea_sinapis_SWE_Lars_10X_103
Leptidea_sinapis_SWE_FD_SWE_Swe-sin-92C
Leptidea_sinapis_SWE_FD_SWE_Swe-sin-91C
Leptidea_sinapis_SWE_FD_SWE_Swe-sin-62C
Leptidea_sinapis_SWE_FD_SWE_Swe-sin-61C
Leptidea_sinapis_SWE_FD_SWE_Swe-sin-32C
Leptidea_sinapis_SWE_FD_SWE_Swe-sin-31C
Leptidea_sinapis_SWE_FD_SWE_Swe-sin-2C
Leptidea_sinapis_SWE_FD_SWE_Swe-sin-1C
Leptidea_sinapis_SWE_FD_SWE_Swe-sin-102C
Leptidea_sinapis_SWE_FD_SWE_Swe-sin-101C
Leptidea_sinapis_CAT_Lars_10X_106
Leptidea_sinapis_CAT_Lars_10X_105
Leptidea_sinapis_CAT_FD_CAT_47
Leptidea_sinapis_CAT_FD_CAT_46
Leptidea_sinapis_CAT_FD_CAT_44
Leptidea_sinapis_CAT_FD_CAT_43
Leptidea_sinapis_CAT_FD_CAT_42
Leptidea_sinapis_CAT_FD_CAT_41
Leptidea_sinapis_CAT_FD_CAT_40
Leptidea_sinapis_CAT_FD_CAT_39
Leptidea_sinapis_CAT_FD_CAT_38
Leptidea_sinapis_CAT_FD_CAT_37
Leptidea_reali_CAT_Lars_10X_116
Leptidea_reali_CAT_Lars_10X_115
Leptidea_reali_CAT_CAtrea_36
Leptidea_reali_CAT_CAtrea_35
Leptidea_reali_CAT_CAtrea_34
Leptidea_reali_CAT_CAtrea_33
Leptidea_reali_CAT_CAtrea_32
Leptidea_reali_CAT_CAtrea_31
Leptidea_reali_CAT_CAtrea_30
Leptidea_reali_CAT_CAtrea_29
Leptidea_reali_CAT_CAtrea_27
Leptidea_reali_CAT_CAtrea_26
Leptidea_morsei_CHI_LEP_outgroups_108
Leptidea_morsei_CHI_LEP_outgroups_107
Leptidea_lactea_CHI_LEP_outgroups_106
Leptidea_lactea_CHI_LEP_outgroups_105
Leptidea_juvernica_SWE_SWE96_186
Leptidea_juvernica_SWE_SWE96_133
Leptidea_juvernica_SWE_SWE96_132
Leptidea_juvernica_SWE_SWE96_128
Leptidea_juvernica_SWE_SWE96_127
Leptidea_juvernica_SWE_SWE96_126
Leptidea_juvernica_SWE_SWE96_125
Leptidea_juvernica_SWE_SWE96_124
Leptidea_juvernica_SWE_SWE96_121
Leptidea_juvernica_SWE_SWE96_120
Leptidea_juvernica_SWE_SWE96_119
Leptidea_juvernica_SWE_SWE96_117
Leptidea_juvernica_SWE_Lars_10X_112
Leptidea_juvernica_SWE_Lars_10X_111
Leptidea_amurensis_MON_LEP_outgroups_102
Leptidea_amurensis_MON_LEP_outgroups_101

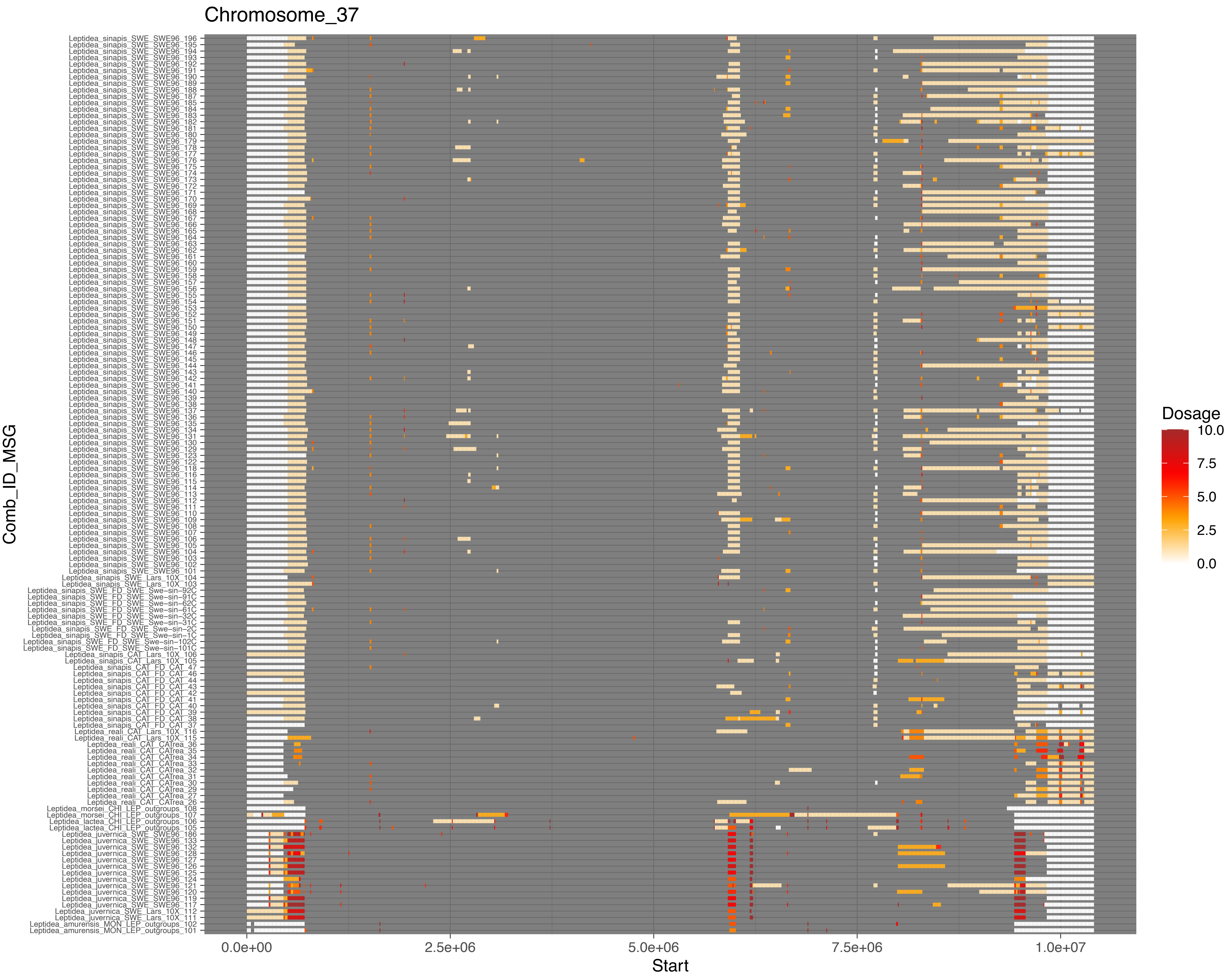

Chromosome_37
Comb_ID_MSG
Start
0.0e+00
2.5e+06
5.0e+06
7.5e+06
1.0e+07
Dosage
10.0
7.5
5.0
2.5
0.0

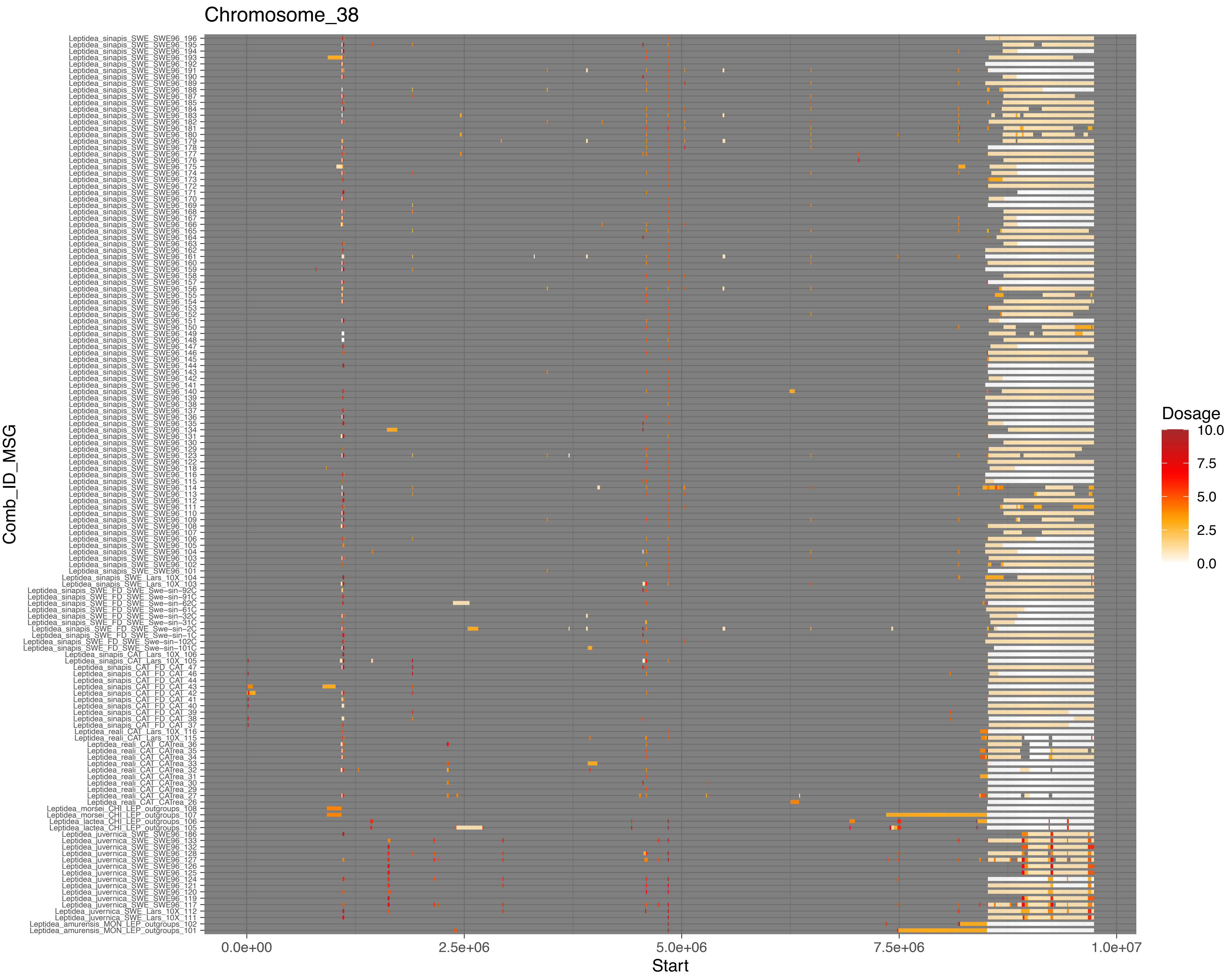

Chromosome_38
Comb_ID_MSG
Leptidea_sinapis_SWE_SWE96_196
Leptidea_sinapis_SWE_SWE96_195
Leptidea_sinapis_SWE_SWE96_194
Leptidea_sinapis_SWE_SWE96_193
Leptidea_sinapis_SWE_SWE96_192
Leptidea_sinapis_SWE_SWE96_191
Leptidea_sinapis_SWE_SWE96_190
Leptidea_sinapis_SWE_SWE96_189
Leptidea_sinapis_SWE_SWE96_188
Leptidea_sinapis_SWE_SWE96_187
Leptidea_sinapis_SWE_SWE96_185
Leptidea_sinapis_SWE_SWE96_184
Leptidea_sinapis_SWE_SWE96_183
Leptidea_sinapis_SWE_SWE96_182
Leptidea_sinapis_SWE_SWE96_181
Leptidea_sinapis_SWE_SWE96_180
Leptidea_sinapis_SWE_SWE96_179
Leptidea_sinapis_SWE_SWE96_178
Leptidea_sinapis_SWE_SWE96_177
Leptidea_sinapis_SWE_SWE96_176
Leptidea_sinapis_SWE_SWE96_175
Leptidea_sinapis_SWE_SWE96_174
Leptidea_sinapis_SWE_SWE96_173
Leptidea_sinapis_SWE_SWE96_172
Leptidea_sinapis_SWE_SWE96_171
Leptidea_sinapis_SWE_SWE96_170
Leptidea_sinapis_SWE_SWE96_169
Leptidea_sinapis_SWE_SWE96_168
Leptidea_sinapis_SWE_SWE96_167
Leptidea_sinapis_SWE_SWE96_166
Leptidea_sinapis_SWE_SWE96_165
Leptidea_sinapis_SWE_SWE96_164
Leptidea_sinapis_SWE_SWE96_163
Leptidea_sinapis_SWE_SWE96_162
Leptidea_sinapis_SWE_SWE96_161
Leptidea_sinapis_SWE_SWE96_160
Leptidea_sinapis_SWE_SWE96_159
Leptidea_sinapis_SWE_SWE96_158
Leptidea_sinapis_SWE_SWE96_157
Leptidea_sinapis_SWE_SWE96_156
Leptidea_sinapis_SWE_SWE96_155
Leptidea_sinapis_SWE_SWE96_154
Leptidea_sinapis_SWE_SWE96_153
Leptidea_sinapis_SWE_SWE96_152
Leptidea_sinapis_SWE_SWE96_151
Leptidea_sinapis_SWE_SWE96_150
Leptidea_sinapis_SWE_SWE96_149
Leptidea_sinapis_SWE_SWE96_148
Leptidea_sinapis_SWE_SWE96_147
Leptidea_sinapis_SWE_SWE96_146
Leptidea_sinapis_SWE_SWE96_145
Leptidea_sinapis_SWE_SWE96_144
Leptidea_sinapis_SWE_SWE96_143
Leptidea_sinapis_SWE_SWE96_142
Leptidea_sinapis_SWE_SWE96_141
Leptidea_sinapis_SWE_SWE96_140
Leptidea_sinapis_SWE_SWE96_139
Leptidea_sinapis_SWE_SWE96_138
Leptidea_sinapis_SWE_SWE96_137
Leptidea_sinapis_SWE_SWE96_136
Leptidea_sinapis_SWE_SWE96_135
Leptidea_sinapis_SWE_SWE96_134
Leptidea_sinapis_SWE_SWE96_131
Leptidea_sinapis_SWE_SWE96_130
Leptidea_sinapis_SWE_SWE96_129
Leptidea_sinapis_SWE_SWE96_123
Leptidea_sinapis_SWE_SWE96_122
Leptidea_sinapis_SWE_SWE96_118
Leptidea_sinapis_SWE_SWE96_116
Leptidea_sinapis_SWE_SWE96_115
Leptidea_sinapis_SWE_SWE96_114
Leptidea_sinapis_SWE_SWE96_113
Leptidea_sinapis_SWE_SWE96_112
Leptidea_sinapis_SWE_SWE96_111
Leptidea_sinapis_SWE_SWE96_110
Leptidea_sinapis_SWE_SWE96_109
Leptidea_sinapis_SWE_SWE96_108
Leptidea_sinapis_SWE_SWE96_107
Leptidea_sinapis_SWE_SWE96_106
Leptidea_sinapis_SWE_SWE96_105
Leptidea_sinapis_SWE_SWE96_104
Leptidea_sinapis_SWE_SWE96_103
Leptidea_sinapis_SWE_SWE96_102
Leptidea_sinapis_SWE_SWE96_101
Leptidea_sinapis_SWE_Lars_10X_104
Leptidea_sinapis_SWE_Lars_10X_103
Leptidea_sinapis_SWE_FD_SWE_Swe−sin−92C
Leptidea_sinapis_SWE_FD_SWE_Swe−sin−91C
Leptidea_sinapis_SWE_FD_SWE_Swe−sin−62C
Leptidea_sinapis_SWE_FD_SWE_Swe−sin−61C
Leptidea_sinapis_SWE_FD_SWE_Swe−sin−32C
Leptidea_sinapis_SWE_FD_SWE_Swe−sin−31C
Leptidea_sinapis_SWE_FD_SWE_Swe−sin−2C
Leptidea_sinapis_SWE_FD_SWE_Swe−sin−1C
Leptidea_sinapis_SWE_FD_SWE_Swe−sin−102C
Leptidea_sinapis_SWE_FD_SWE_Swe−sin−101C
Leptidea_sinapis_CAT_Lars_10X_106
Leptidea_sinapis_CAT_Lars_10X_105
Leptidea_sinapis_CAT_FD_CAT_47
Leptidea_sinapis_CAT_FD_CAT_46
Leptidea_sinapis_CAT_FD_CAT_44
Leptidea_sinapis_CAT_FD_CAT_43
Leptidea_sinapis_CAT_FD_CAT_42
Leptidea_sinapis_CAT_FD_CAT_41
Leptidea_sinapis_CAT_FD_CAT_40
Leptidea_sinapis_CAT_FD_CAT_39
Leptidea_sinapis_CAT_FD_CAT_38
Leptidea_sinapis_CAT_FD_CAT_37
Leptidea_reali_CAT_Lars_10X_116
Leptidea_reali_CAT_Lars_10X_115
Leptidea_reali_CAT_CATrea_36
Leptidea_reali_CAT_CATrea_35
Leptidea_reali_CAT_CATrea_34
Leptidea_reali_CAT_CATrea_33
Leptidea_reali_CAT_CATrea_32
Leptidea_reali_CAT_CATrea_31
Leptidea_reali_CAT_CATrea_30
Leptidea_reali_CAT_CATrea_29
Leptidea_reali_CAT_CATrea_27
Leptidea_reali_CAT_CATrea_26
Leptidea_morsei_CHI_LEP_outgroups_108
Leptidea_morsei_CHI_LEP_outgroups_107
Leptidea_lactea_CHI_LEP_outgroups_106
Leptidea_lactea_CHI_LEP_outgroups_105
Leptidea_juvernica_SWE_SWE96_186
Leptidea_juvernica_SWE_SWE96_133
Leptidea_juvernica_SWE_SWE96_132
Leptidea_juvernica_SWE_SWE96_128
Leptidea_juvernica_SWE_SWE96_127
Leptidea_juvernica_SWE_SWE96_126
Leptidea_juvernica_SWE_SWE96_125
Leptidea_juvernica_SWE_SWE96_124
Leptidea_juvernica_SWE_SWE96_121
Leptidea_juvernica_SWE_SWE96_120
Leptidea_juvernica_SWE_SWE96_119
Leptidea_juvernica_SWE_SWE96_117
Leptidea_juvernica_SWE_Lars_10X_112
Leptidea_juvernica_SWE_Lars_10X_111
Leptidea_amurensis_MON_LEP_outgroups_102
Leptidea_amurensis_MON_LEP_outgroups_101
0.0e+00
2.5e+06
5.0e+06
7.5e+06
1.0e+07
Start
Dosage
10.0
7.5
5.0
2.5
0.0

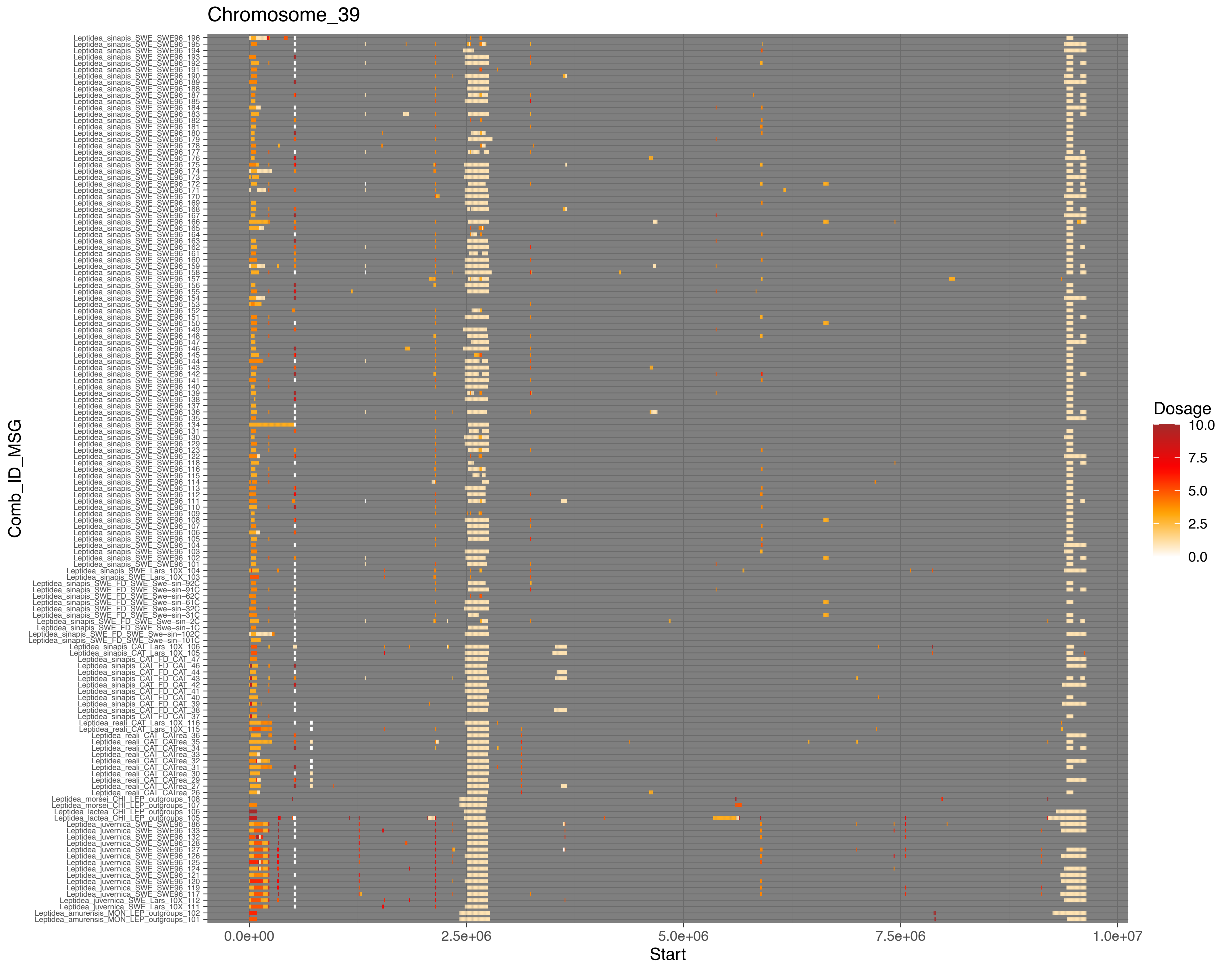

Chromosome_39
Leptidea_sinapis_SWE_SWE96_196
Leptidea_sinapis_SWE_SWE96_195
Leptidea_sinapis_SWE_SWE96_194
Leptidea_sinapis_SWE_SWE96_193
Leptidea_sinapis_SWE_SWE96_192
Leptidea_sinapis_SWE_SWE96_191
Leptidea_sinapis_SWE_SWE96_190
Leptidea_sinapis_SWE_SWE96_189
Leptidea_sinapis_SWE_SWE96_188
Leptidea_sinapis_SWE_SWE96_187
Leptidea_sinapis_SWE_SWE96_185
Leptidea_sinapis_SWE_SWE96_184
Leptidea_sinapis_SWE_SWE96_183
Leptidea_sinapis_SWE_SWE96_182
Leptidea_sinapis_SWE_SWE96_181
Leptidea_sinapis_SWE_SWE96_180
Leptidea_sinapis_SWE_SWE96_179
Leptidea_sinapis_SWE_SWE96_178
Leptidea_sinapis_SWE_SWE96_177
Leptidea_sinapis_SWE_SWE96_176
Leptidea_sinapis_SWE_SWE96_175
Leptidea_sinapis_SWE_SWE96_174
Leptidea_sinapis_SWE_SWE96_173
Leptidea_sinapis_SWE_SWE96_172
Leptidea_sinapis_SWE_SWE96_171
Leptidea_sinapis_SWE_SWE96_170
Leptidea_sinapis_SWE_SWE96_169
Leptidea_sinapis_SWE_SWE96_168
Leptidea_sinapis_SWE_SWE96_167
Leptidea_sinapis_SWE_SWE96_166
Leptidea_sinapis_SWE_SWE96_165
Leptidea_sinapis_SWE_SWE96_164
Leptidea_sinapis_SWE_SWE96_163
Leptidea_sinapis_SWE_SWE96_162
Leptidea_sinapis_SWE_SWE96_161
Leptidea_sinapis_SWE_SWE96_160
Leptidea_sinapis_SWE_SWE96_159
Leptidea_sinapis_SWE_SWE96_158
Leptidea_sinapis_SWE_SWE96_157
Leptidea_sinapis_SWE_SWE96_156
Leptidea_sinapis_SWE_SWE96_155
Leptidea_sinapis_SWE_SWE96_154
Leptidea_sinapis_SWE_SWE96_153
Leptidea_sinapis_SWE_SWE96_152
Leptidea_sinapis_SWE_SWE96_151
Leptidea_sinapis_SWE_SWE96_150
Leptidea_sinapis_SWE_SWE96_149
Leptidea_sinapis_SWE_SWE96_148
Leptidea_sinapis_SWE_SWE96_147
Leptidea_sinapis_SWE_SWE96_146
Leptidea_sinapis_SWE_SWE96_145
Leptidea_sinapis_SWE_SWE96_144
Leptidea_sinapis_SWE_SWE96_143
Leptidea_sinapis_SWE_SWE96_142
Leptidea_sinapis_SWE_SWE96_141
Leptidea_sinapis_SWE_SWE96_140
Leptidea_sinapis_SWE_SWE96_139
Leptidea_sinapis_SWE_SWE96_138
Leptidea_sinapis_SWE_SWE96_137
Leptidea_sinapis_SWE_SWE96_136
Leptidea_sinapis_SWE_SWE96_135
Leptidea_sinapis_SWE_SWE96_134
Leptidea_sinapis_SWE_SWE96_131
Leptidea_sinapis_SWE_SWE96_130
Leptidea_sinapis_SWE_SWE96_129
Leptidea_sinapis_SWE_SWE96_123
Leptidea_sinapis_SWE_SWE96_122
Leptidea_sinapis_SWE_SWE96_118
Leptidea_sinapis_SWE_SWE96_116
Leptidea_sinapis_SWE_SWE96_115
Leptidea_sinapis_SWE_SWE96_114
Leptidea_sinapis_SWE_SWE96_113
Leptidea_sinapis_SWE_SWE96_112
Leptidea_sinapis_SWE_SWE96_111
Leptidea_sinapis_SWE_SWE96_110
Leptidea_sinapis_SWE_SWE96_109
Leptidea_sinapis_SWE_SWE96_108
Leptidea_sinapis_SWE_SWE96_107
Leptidea_sinapis_SWE_SWE96_106
Leptidea_sinapis_SWE_SWE96_105
Leptidea_sinapis_SWE_SWE96_104
Leptidea_sinapis_SWE_SWE96_103
Leptidea_sinapis_SWE_SWE96_102
Leptidea_sinapis_SWE_SWE96_101
Leptidea_sinapis_SWE_Lars_10X_104
Leptidea_sinapis_SWE_Lars_10X_103
Leptidea_sinapis_SWE_FD_SWE_Swe-sin-92C
Leptidea_sinapis_SWE_FD_SWE_Swe-sin-91C
Leptidea_sinapis_SWE_FD_SWE_Swe-sin-62C
Leptidea_sinapis_SWE_FD_SWE_Swe-sin-61C
Leptidea_sinapis_SWE_FD_SWE_Swe-sin-32C
Leptidea_sinapis_SWE_FD_SWE_Swe-sin-31C
Leptidea_sinapis_SWE_FD_SWE_Swe-sin-2C
Leptidea_sinapis_SWE_FD_SWE_Swe-sin-1C
Leptidea_sinapis_SWE_FD_SWE_Swe-sin-102C
Leptidea_sinapis_SWE_FD_SWE_Swe-sin-101C
Leptidea_sinapis_CAT_Lars_10X_106
Leptidea_sinapis_CAT_Lars_10X_105
Leptidea_sinapis_CAT_FD_CAT_47
Leptidea_sinapis_CAT_FD_CAT_46
Leptidea_sinapis_CAT_FD_CAT_44
Leptidea_sinapis_CAT_FD_CAT_43
Leptidea_sinapis_CAT_FD_CAT_42
Leptidea_sinapis_CAT_FD_CAT_41
Leptidea_sinapis_CAT_FD_CAT_40
Leptidea_sinapis_CAT_FD_CAT_39
Leptidea_sinapis_CAT_FD_CAT_38
Leptidea_sinapis_CAT_FD_CAT_37
Leptidea_reali_CAT_Lars_10X_116
Leptidea_reali_CAT_Lars_10X_115
Leptidea_reali_CAT_CATrea_36
Leptidea_reali_CAT_CATrea_35
Leptidea_reali_CAT_CATrea_34
Leptidea_reali_CAT_CATrea_33
Leptidea_reali_CAT_CATrea_32
Leptidea_reali_CAT_CATrea_31
Leptidea_reali_CAT_CATrea_30
Leptidea_reali_CAT_CATrea_29
Leptidea_reali_CAT_CATrea_27
Leptidea_reali_CAT_CATrea_26
Leptidea_morsei_CHI_LEP_outgroups_108
Leptidea_morsei_CHI_LEP_outgroups_107
Leptidea_lactea_CHI_LEP_outgroups_106
Leptidea_lactea_CHI_LEP_outgroups_105
Leptidea_juvernica_SWE_SWE96_186
Leptidea_juvernica_SWE_SWE96_133
Leptidea_juvernica_SWE_SWE96_132
Leptidea_juvernica_SWE_SWE96_128
Leptidea_juvernica_SWE_SWE96_127
Leptidea_juvernica_SWE_SWE96_126
Leptidea_juvernica_SWE_SWE96_125
Leptidea_juvernica_SWE_SWE96_124
Leptidea_juvernica_SWE_SWE96_121
Leptidea_juvernica_SWE_SWE96_120
Leptidea_juvernica_SWE_SWE96_119
Leptidea_juvernica_SWE_SWE96_117
Leptidea_juvernica_SWE_Lars_10X_112
Leptidea_juvernica_SWE_Lars_10X_111
Leptidea_amurensis_MON_LEP_outgroups_102
Leptidea_amurensis_MON_LEP_outgroups_101
Comb_ID_MSG
0.0e+00
2.5e+06
5.0e+06
7.5e+06
1.0e+07
Start
Dosage
10.0
7.5
5.0
2.5
0.0

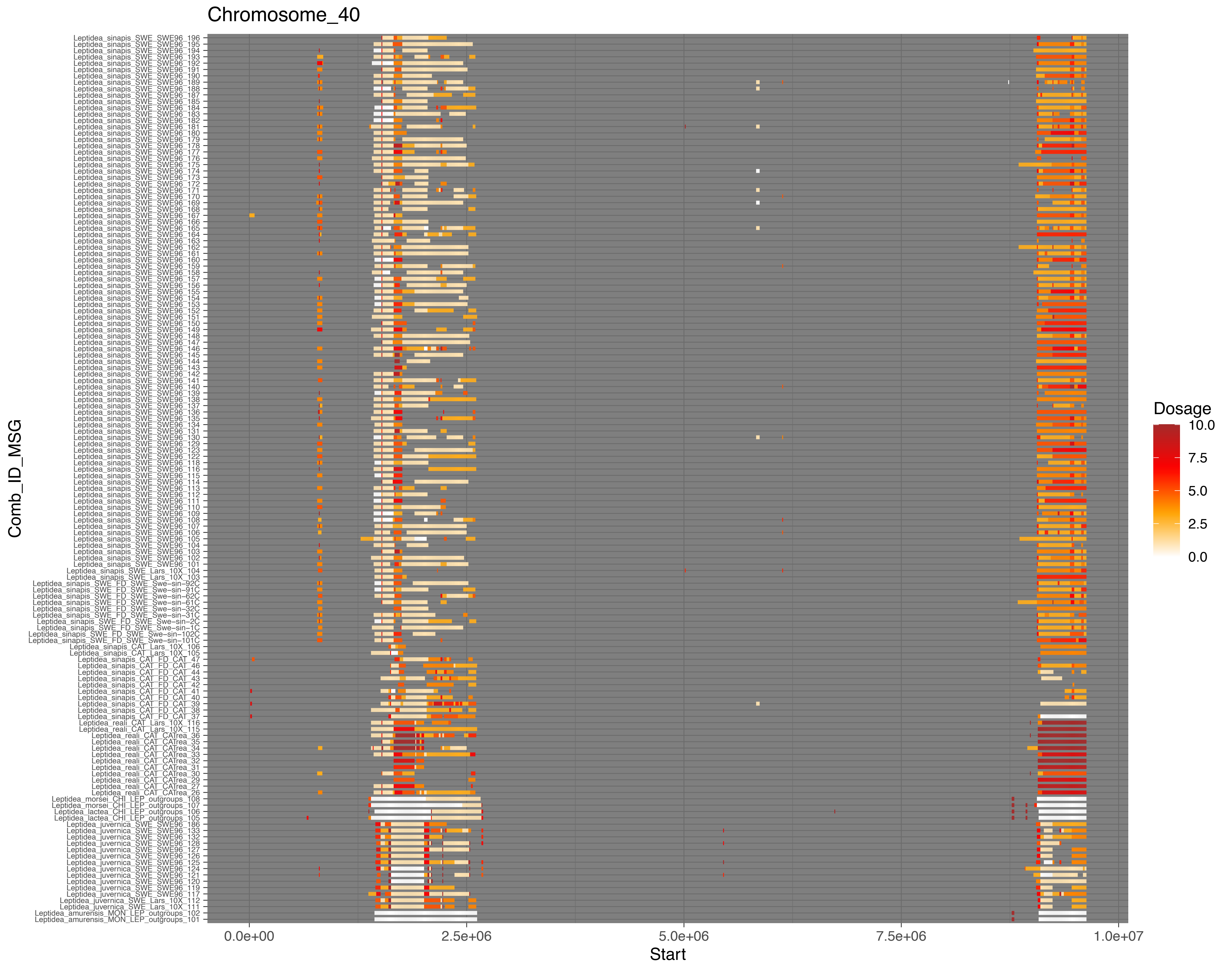
Chromosome_40
Comb_ID_MSG
Start
0.0e+00
2.5e+06
5.0e+06
7.5e+06
1.0e+07
Dosage
10.0
7.5
5.0
2.5
0.0

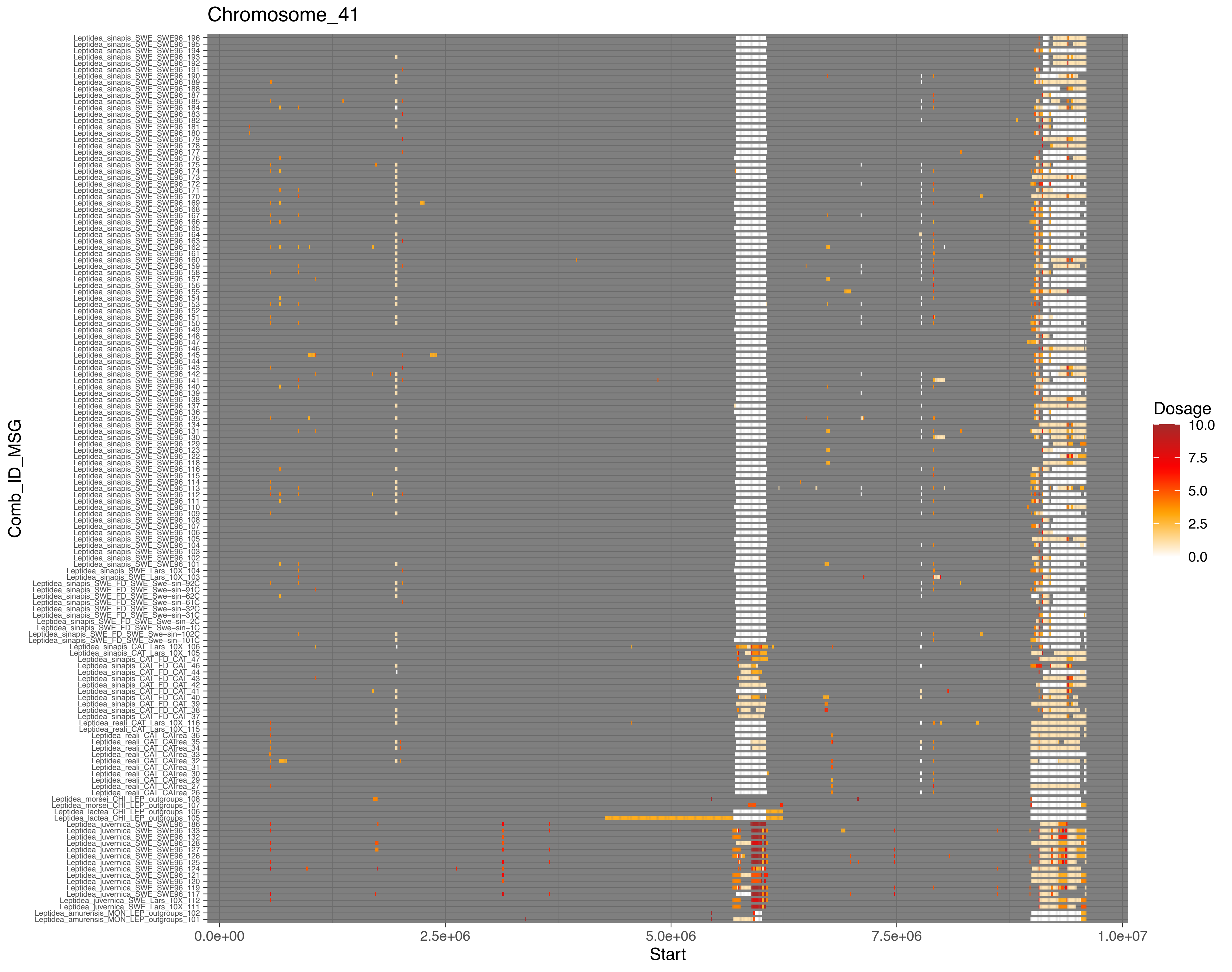

Chromosome_41
Comb_ID_MSG
Start
0.0e+00
2.5e+06
5.0e+06
7.5e+06
1.0e+07
Dosage
10.0
7.5
5.0
2.5
0.0

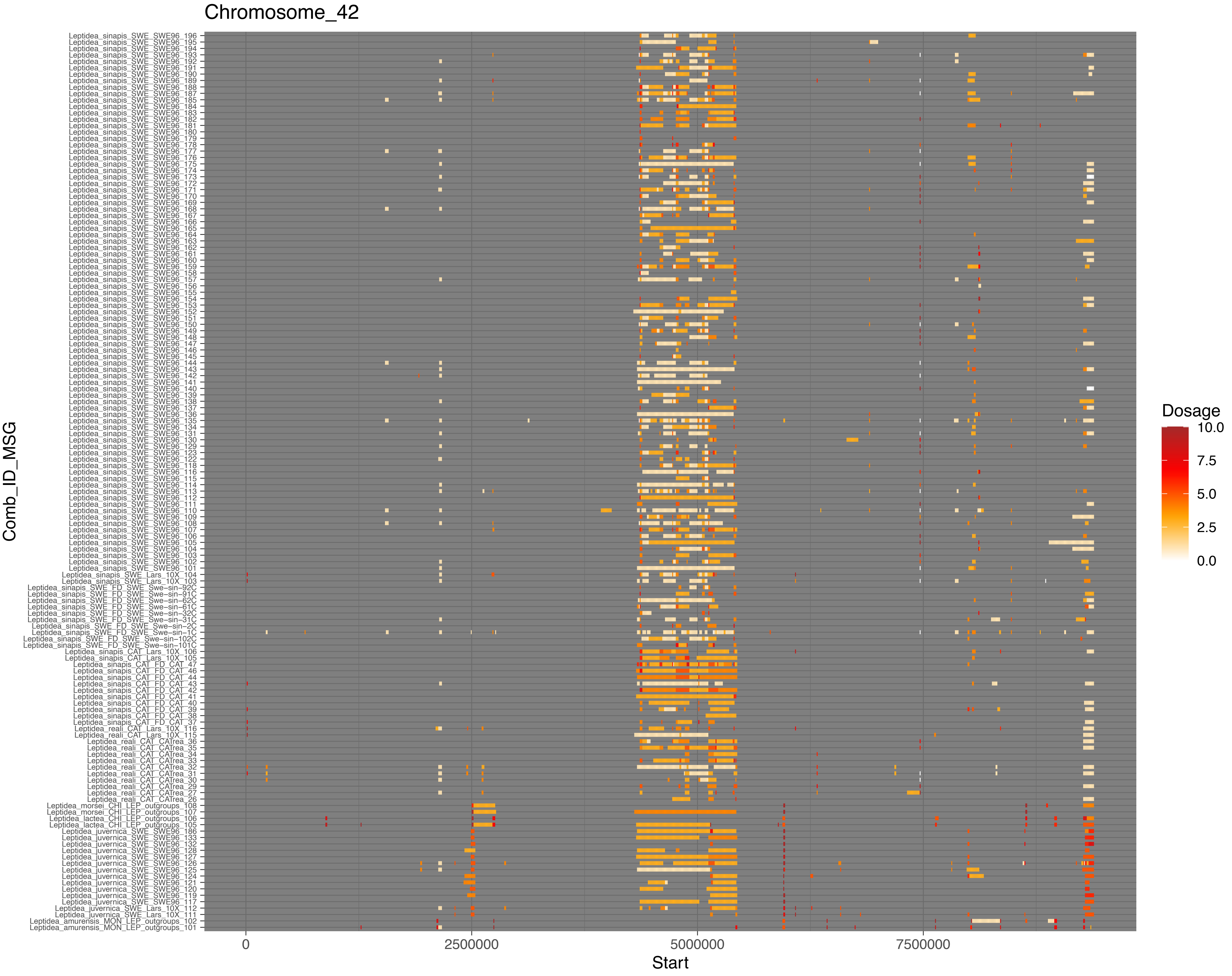

Chromosome_42
Comb_ID_MSG
Start
0
2500000
5000000
7500000
Dosage
10.0
7.5
5.0
2.5
0.0
Leptidea_sinapis_SWE_SWE96_196
Leptidea_sinapis_SWE_SWE96_195
Leptidea_sinapis_SWE_SWE96_194
Leptidea_sinapis_SWE_SWE96_193
Leptidea_sinapis_SWE_SWE96_192
Leptidea_sinapis_SWE_SWE96_191
Leptidea_sinapis_SWE_SWE96_190
Leptidea_sinapis_SWE_SWE96_189
Leptidea_sinapis_SWE_SWE96_188
Leptidea_sinapis_SWE_SWE96_187
Leptidea_sinapis_SWE_SWE96_185
Leptidea_sinapis_SWE_SWE96_184
Leptidea_sinapis_SWE_SWE96_183
Leptidea_sinapis_SWE_SWE96_182
Leptidea_sinapis_SWE_SWE96_181
Leptidea_sinapis_SWE_SWE96_180
Leptidea_sinapis_SWE_SWE96_179
Leptidea_sinapis_SWE_SWE96_178
Leptidea_sinapis_SWE_SWE96_177
Leptidea_sinapis_SWE_SWE96_176
Leptidea_sinapis_SWE_SWE96_175
Leptidea_sinapis_SWE_SWE96_174
Leptidea_sinapis_SWE_SWE96_173
Leptidea_sinapis_SWE_SWE96_172
Leptidea_sinapis_SWE_SWE96_171
Leptidea_sinapis_SWE_SWE96_170
Leptidea_sinapis_SWE_SWE96_169
Leptidea_sinapis_SWE_SWE96_168
Leptidea_sinapis_SWE_SWE96_167
Leptidea_sinapis_SWE_SWE96_166
Leptidea_sinapis_SWE_SWE96_165
Leptidea_sinapis_SWE_SWE96_164
Leptidea_sinapis_SWE_SWE96_163
Leptidea_sinapis_SWE_SWE96_162
Leptidea_sinapis_SWE_SWE96_161
Leptidea_sinapis_SWE_SWE96_160
Leptidea_sinapis_SWE_SWE96_159
Leptidea_sinapis_SWE_SWE96_158
Leptidea_sinapis_SWE_SWE96_157
Leptidea_sinapis_SWE_SWE96_156
Leptidea_sinapis_SWE_SWE96_155
Leptidea_sinapis_SWE_SWE96_154
Leptidea_sinapis_SWE_SWE96_153
Leptidea_sinapis_SWE_SWE96_152
Leptidea_sinapis_SWE_SWE96_151
Leptidea_sinapis_SWE_SWE96_150
Leptidea_sinapis_SWE_SWE96_149
Leptidea_sinapis_SWE_SWE96_148
Leptidea_sinapis_SWE_SWE96_147
Leptidea_sinapis_SWE_SWE96_146
Leptidea_sinapis_SWE_SWE96_145
Leptidea_sinapis_SWE_SWE96_144
Leptidea_sinapis_SWE_SWE96_143
Leptidea_sinapis_SWE_SWE96_142
Leptidea_sinapis_SWE_SWE96_141
Leptidea_sinapis_SWE_SWE96_140
Leptidea_sinapis_SWE_SWE96_139
Leptidea_sinapis_SWE_SWE96_138
Leptidea_sinapis_SWE_SWE96_137
Leptidea_sinapis_SWE_SWE96_136
Leptidea_sinapis_SWE_SWE96_135
Leptidea_sinapis_SWE_SWE96_134
Leptidea_sinapis_SWE_SWE96_131
Leptidea_sinapis_SWE_SWE96_130
Leptidea_sinapis_SWE_SWE96_129
Leptidea_sinapis_SWE_SWE96_123
Leptidea_sinapis_SWE_SWE96_122
Leptidea_sinapis_SWE_SWE96_118
Leptidea_sinapis_SWE_SWE96_116
Leptidea_sinapis_SWE_SWE96_115
Leptidea_sinapis_SWE_SWE96_114
Leptidea_sinapis_SWE_SWE96_113
Leptidea_sinapis_SWE_SWE96_112
Leptidea_sinapis_SWE_SWE96_111
Leptidea_sinapis_SWE_SWE96_110
Leptidea_sinapis_SWE_SWE96_109
Leptidea_sinapis_SWE_SWE96_108
Leptidea_sinapis_SWE_SWE96_107
Leptidea_sinapis_SWE_SWE96_106
Leptidea_sinapis_SWE_SWE96_105
Leptidea_sinapis_SWE_SWE96_104
Leptidea_sinapis_SWE_SWE96_103
Leptidea_sinapis_SWE_SWE96_102
Leptidea_sinapis_SWE_SWE96_101
Leptidea_sinapis_SWE_Lars_10X_104
Leptidea_sinapis_SWE_Lars_10X_103
Leptidea_sinapis_SWE_FD_SWE_Swe-sin-92C
Leptidea_sinapis_SWE_FD_SWE_Swe-sin-91C
Leptidea_sinapis_SWE_FD_SWE_Swe-sin-62C
Leptidea_sinapis_SWE_FD_SWE_Swe-sin-61C
Leptidea_sinapis_SWE_FD_SWE_Swe-sin-32C
Leptidea_sinapis_SWE_FD_SWE_Swe-sin-31C
Leptidea_sinapis_SWE_FD_SWE_Swe-sin-2C
Leptidea_sinapis_SWE_FD_SWE_Swe-sin-1C
Leptidea_sinapis_SWE_FD_SWE_Swe-sin-102C
Leptidea_sinapis_SWE_FD_SWE_Swe-sin-101C
Leptidea_sinapis_CAT_Lars_10X_106
Leptidea_sinapis_CAT_Lars_10X_105
Leptidea_sinapis_CAT_FD_CAT_47
Leptidea_sinapis_CAT_FD_CAT_46
Leptidea_sinapis_CAT_FD_CAT_44
Leptidea_sinapis_CAT_FD_CAT_43
Leptidea_sinapis_CAT_FD_CAT_42
Leptidea_sinapis_CAT_FD_CAT_41
Leptidea_sinapis_CAT_FD_CAT_40
Leptidea_sinapis_CAT_FD_CAT_39
Leptidea_sinapis_CAT_FD_CAT_38
Leptidea_sinapis_CAT_FD_CAT_37
Leptidea_reali_CAT_Lars_10X_116
Leptidea_reali_CAT_Lars_10X_115
Leptidea_reali_CAT_CATrea_36
Leptidea_reali_CAT_CATrea_35
Leptidea_reali_CAT_CATrea_34
Leptidea_reali_CAT_CATrea_33
Leptidea_reali_CAT_CATrea_32
Leptidea_reali_CAT_CATrea_31
Leptidea_reali_CAT_CATrea_30
Leptidea_reali_CAT_CATrea_29
Leptidea_reali_CAT_CATrea_27
Leptidea_reali_CAT_CATrea_26
Leptidea_morsei_CHI_LEP_outgroups_108
Leptidea_morsei_CHI_LEP_outgroups_107
Leptidea_lactea_CHI_LEP_outgroups_106
Leptidea_lactea_CHI_LEP_outgroups_105
Leptidea_juvernica_SWE_SWE96_186
Leptidea_juvernica_SWE_SWE96_133
Leptidea_juvernica_SWE_SWE96_132
Leptidea_juvernica_SWE_SWE96_128
Leptidea_juvernica_SWE_SWE96_127
Leptidea_juvernica_SWE_SWE96_126
Leptidea_juvernica_SWE_SWE96_125
Leptidea_juvernica_SWE_SWE96_124
Leptidea_juvernica_SWE_SWE96_121
Leptidea_juvernica_SWE_SWE96_120
Leptidea_juvernica_SWE_SWE96_119
Leptidea_juvernica_SWE_SWE96_117
Leptidea_juvernica_SWE_Lars_10X_112
Leptidea_juvernica_SWE_Lars_10X_111
Leptidea_amurensis_MON_LEP_outgroups_102
Leptidea_amurensis_MON_LEP_outgroups_101

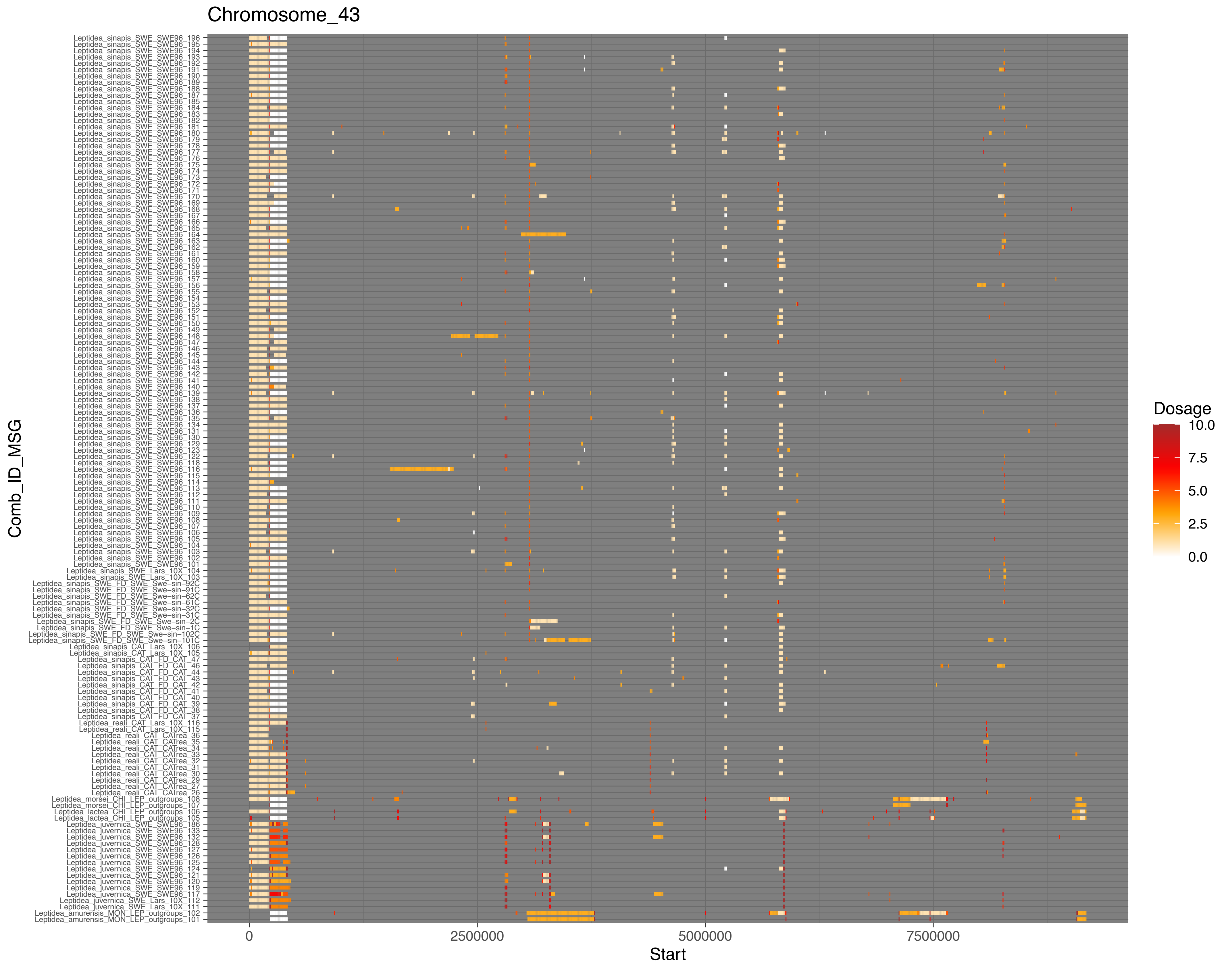
Chromosome_43
Leptidea_sinapis_SWE_SWE96_196
Leptidea_sinapis_SWE_SWE96_195
Leptidea_sinapis_SWE_SWE96_194
Leptidea_sinapis_SWE_SWE96_193
Leptidea_sinapis_SWE_SWE96_192
Leptidea_sinapis_SWE_SWE96_191
Leptidea_sinapis_SWE_SWE96_190
Leptidea_sinapis_SWE_SWE96_189
Leptidea_sinapis_SWE_SWE96_188
Leptidea_sinapis_SWE_SWE96_187
Leptidea_sinapis_SWE_SWE96_185
Leptidea_sinapis_SWE_SWE96_184
Leptidea_sinapis_SWE_SWE96_183
Leptidea_sinapis_SWE_SWE96_182
Leptidea_sinapis_SWE_SWE96_181
Leptidea_sinapis_SWE_SWE96_180
Leptidea_sinapis_SWE_SWE96_179
Leptidea_sinapis_SWE_SWE96_178
Leptidea_sinapis_SWE_SWE96_177
Leptidea_sinapis_SWE_SWE96_176
Leptidea_sinapis_SWE_SWE96_175
Leptidea_sinapis_SWE_SWE96_174
Leptidea_sinapis_SWE_SWE96_173
Leptidea_sinapis_SWE_SWE96_172
Leptidea_sinapis_SWE_SWE96_171
Leptidea_sinapis_SWE_SWE96_170
Leptidea_sinapis_SWE_SWE96_169
Leptidea_sinapis_SWE_SWE96_168
Leptidea_sinapis_SWE_SWE96_167
Leptidea_sinapis_SWE_SWE96_166
Leptidea_sinapis_SWE_SWE96_165
Leptidea_sinapis_SWE_SWE96_164
Leptidea_sinapis_SWE_SWE96_163
Leptidea_sinapis_SWE_SWE96_162
Leptidea_sinapis_SWE_SWE96_161
Leptidea_sinapis_SWE_SWE96_160
Leptidea_sinapis_SWE_SWE96_159
Leptidea_sinapis_SWE_SWE96_158
Leptidea_sinapis_SWE_SWE96_157
Leptidea_sinapis_SWE_SWE96_156
Leptidea_sinapis_SWE_SWE96_155
Leptidea_sinapis_SWE_SWE96_154
Leptidea_sinapis_SWE_SWE96_153
Leptidea_sinapis_SWE_SWE96_152
Leptidea_sinapis_SWE_SWE96_151
Leptidea_sinapis_SWE_SWE96_150
Leptidea_sinapis_SWE_SWE96_149
Leptidea_sinapis_SWE_SWE96_148
Leptidea_sinapis_SWE_SWE96_147
Leptidea_sinapis_SWE_SWE96_146
Leptidea_sinapis_SWE_SWE96_145
Leptidea_sinapis_SWE_SWE96_144
Leptidea_sinapis_SWE_SWE96_143
Leptidea_sinapis_SWE_SWE96_142
Leptidea_sinapis_SWE_SWE96_141
Leptidea_sinapis_SWE_SWE96_140
Leptidea_sinapis_SWE_SWE96_139
Leptidea_sinapis_SWE_SWE96_138
Leptidea_sinapis_SWE_SWE96_137
Leptidea_sinapis_SWE_SWE96_136
Leptidea_sinapis_SWE_SWE96_135
Leptidea_sinapis_SWE_SWE96_134
Leptidea_sinapis_SWE_SWE96_131
Leptidea_sinapis_SWE_SWE96_130
Leptidea_sinapis_SWE_SWE96_129
Leptidea_sinapis_SWE_SWE96_123
Leptidea_sinapis_SWE_SWE96_122
Leptidea_sinapis_SWE_SWE96_118
Leptidea_sinapis_SWE_SWE96_116
Leptidea_sinapis_SWE_SWE96_115
Leptidea_sinapis_SWE_SWE96_114
Leptidea_sinapis_SWE_SWE96_113
Leptidea_sinapis_SWE_SWE96_112
Leptidea_sinapis_SWE_SWE96_111
Leptidea_sinapis_SWE_SWE96_110
Leptidea_sinapis_SWE_SWE96_109
Leptidea_sinapis_SWE_SWE96_108
Leptidea_sinapis_SWE_SWE96_107
Leptidea_sinapis_SWE_SWE96_106
Leptidea_sinapis_SWE_SWE96_105
Leptidea_sinapis_SWE_SWE96_104
Leptidea_sinapis_SWE_SWE96_103
Leptidea_sinapis_SWE_SWE96_102
Leptidea_sinapis_SWE_SWE96_101
Leptidea_sinapis_SWE_Lars_10X_104
Leptidea_sinapis_SWE_Lars_10X_103
Leptidea_sinapis_SWE_FD_SWE_Swe-sin-92C
Leptidea_sinapis_SWE_FD_SWE_Swe-sin-91C
Leptidea_sinapis_SWE_FD_SWE_Swe-sin-62C
Leptidea_sinapis_SWE_FD_SWE_Swe-sin-61C
Leptidea_sinapis_SWE_FD_SWE_Swe-sin-32C
Leptidea_sinapis_SWE_FD_SWE_Swe-sin-31C
Leptidea_sinapis_SWE_FD_SWE_Swe-sin-2C
Leptidea_sinapis_SWE_FD_SWE_Swe-sin-1C
Leptidea_sinapis_SWE_FD_SWE_Swe-sin-102C
Leptidea_sinapis_SWE_FD_SWE_Swe-sin-101C
Leptidea_sinapis_CAT_Lars_10X_106
Leptidea_sinapis_CAT_Lars_10X_105
Leptidea_sinapis_CAT_FD_CAT_47
Leptidea_sinapis_CAT_FD_CAT_46
Leptidea_sinapis_CAT_FD_CAT_44
Leptidea_sinapis_CAT_FD_CAT_43
Leptidea_sinapis_CAT_FD_CAT_42
Leptidea_sinapis_CAT_FD_CAT_41
Leptidea_sinapis_CAT_FD_CAT_40
Leptidea_sinapis_CAT_FD_CAT_39
Leptidea_sinapis_CAT_FD_CAT_38
Leptidea_sinapis_CAT_FD_CAT_37
Leptidea_reali_CAT_Lars_10X_116
Leptidea_reali_CAT_Lars_10X_115
Leptidea_reali_CAT_CATrea_36
Leptidea_reali_CAT_CATrea_35
Leptidea_reali_CAT_CATrea_34
Leptidea_reali_CAT_CATrea_33
Leptidea_reali_CAT_CATrea_32
Leptidea_reali_CAT_CATrea_31
Leptidea_reali_CAT_CATrea_30
Leptidea_reali_CAT_CATrea_29
Leptidea_reali_CAT_CATrea_27
Leptidea_reali_CAT_CATrea_26
Leptidea_morsei_CHI_LEP_outgroups_108
Leptidea_morsei_CHI_LEP_outgroups_107
Leptidea_lactea_CHI_LEP_outgroups_106
Leptidea_lactea_CHI_LEP_outgroups_105
Leptidea_juvernica_SWE_SWE96_186
Leptidea_juvernica_SWE_SWE96_133
Leptidea_juvernica_SWE_SWE96_132
Leptidea_juvernica_SWE_SWE96_128
Leptidea_juvernica_SWE_SWE96_127
Leptidea_juvernica_SWE_SWE96_126
Leptidea_juvernica_SWE_SWE96_125
Leptidea_juvernica_SWE_SWE96_124
Leptidea_juvernica_SWE_SWE96_121
Leptidea_juvernica_SWE_SWE96_120
Leptidea_juvernica_SWE_SWE96_119
Leptidea_juvernica_SWE_SWE96_117
Leptidea_juvernica_SWE_Lars_10X_112
Leptidea_juvernica_SWE_Lars_10X_111
Leptidea_amurensis_MON_LEP_outgroups_102
Leptidea_amurensis_MON_LEP_outgroups_101
Comb_ID_MSG
0
2500000
5000000
7500000
Start
Dosage
10.0
7.5
5.0
2.5
0.0

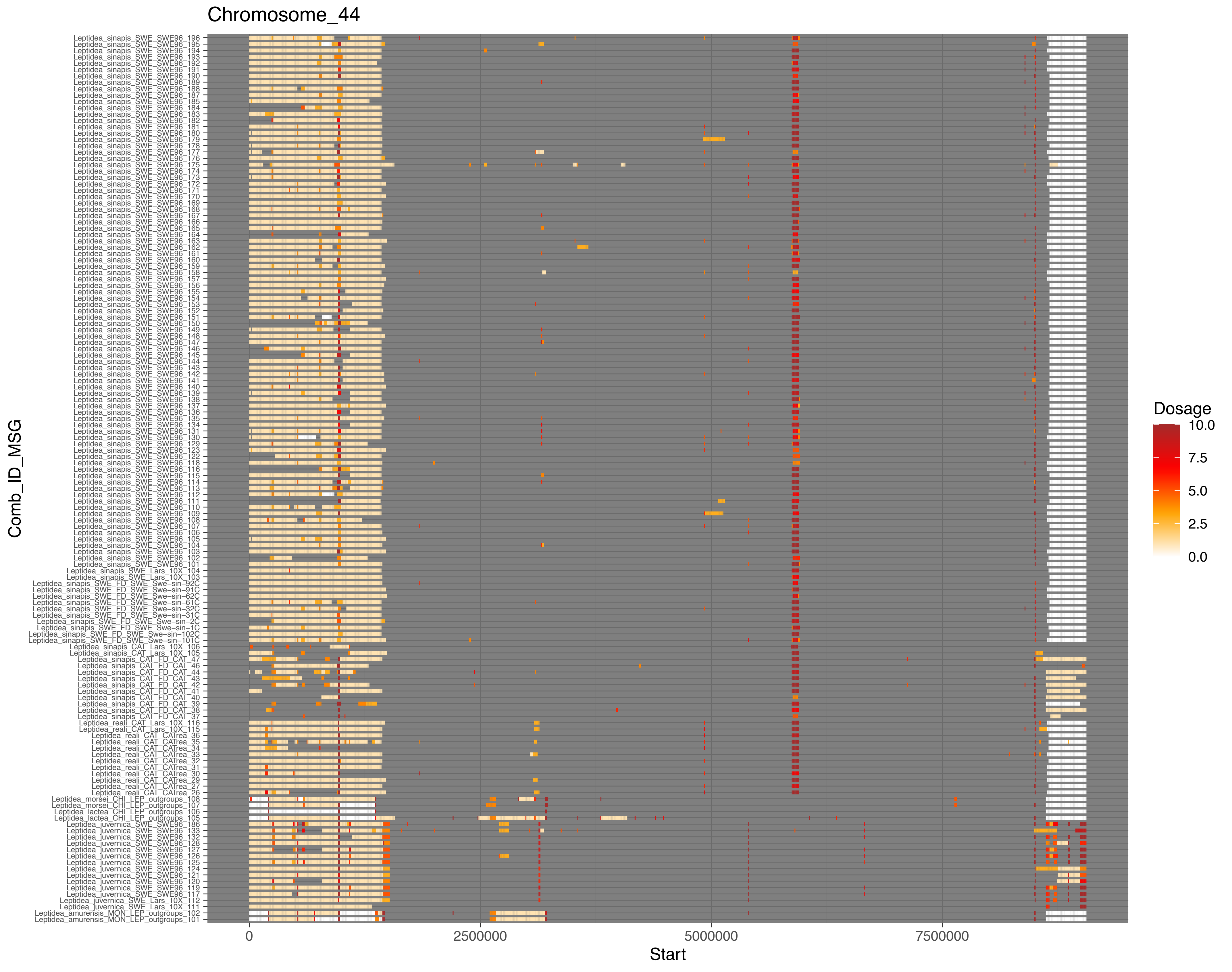
Chromosome_44
Leptidea_sinapis_SWE_SWE96_196
Leptidea_sinapis_SWE_SWE96_195
Leptidea_sinapis_SWE_SWE96_194
Leptidea_sinapis_SWE_SWE96_193
Leptidea_sinapis_SWE_SWE96_192
Leptidea_sinapis_SWE_SWE96_191
Leptidea_sinapis_SWE_SWE96_190
Leptidea_sinapis_SWE_SWE96_189
Leptidea_sinapis_SWE_SWE96_188
Leptidea_sinapis_SWE_SWE96_187
Leptidea_sinapis_SWE_SWE96_185
Leptidea_sinapis_SWE_SWE96_184
Leptidea_sinapis_SWE_SWE96_183
Leptidea_sinapis_SWE_SWE96_182
Leptidea_sinapis_SWE_SWE96_181
Leptidea_sinapis_SWE_SWE96_180
Leptidea_sinapis_SWE_SWE96_179
Leptidea_sinapis_SWE_SWE96_178
Leptidea_sinapis_SWE_SWE96_177
Leptidea_sinapis_SWE_SWE96_176
Leptidea_sinapis_SWE_SWE96_175
Leptidea_sinapis_SWE_SWE96_174
Leptidea_sinapis_SWE_SWE96_173
Leptidea_sinapis_SWE_SWE96_172
Leptidea_sinapis_SWE_SWE96_171
Leptidea_sinapis_SWE_SWE96_170
Leptidea_sinapis_SWE_SWE96_169
Leptidea_sinapis_SWE_SWE96_168
Leptidea_sinapis_SWE_SWE96_167
Leptidea_sinapis_SWE_SWE96_166
Leptidea_sinapis_SWE_SWE96_165
Leptidea_sinapis_SWE_SWE96_164
Leptidea_sinapis_SWE_SWE96_163
Leptidea_sinapis_SWE_SWE96_162
Leptidea_sinapis_SWE_SWE96_161
Leptidea_sinapis_SWE_SWE96_160
Leptidea_sinapis_SWE_SWE96_159
Leptidea_sinapis_SWE_SWE96_158
Leptidea_sinapis_SWE_SWE96_157
Leptidea_sinapis_SWE_SWE96_156
Leptidea_sinapis_SWE_SWE96_155
Leptidea_sinapis_SWE_SWE96_154
Leptidea_sinapis_SWE_SWE96_153
Leptidea_sinapis_SWE_SWE96_152
Leptidea_sinapis_SWE_SWE96_151
Leptidea_sinapis_SWE_SWE96_150
Leptidea_sinapis_SWE_SWE96_149
Leptidea_sinapis_SWE_SWE96_148
Leptidea_sinapis_SWE_SWE96_147
Leptidea_sinapis_SWE_SWE96_146
Leptidea_sinapis_SWE_SWE96_145
Leptidea_sinapis_SWE_SWE96_144
Leptidea_sinapis_SWE_SWE96_143
Leptidea_sinapis_SWE_SWE96_142
Leptidea_sinapis_SWE_SWE96_141
Leptidea_sinapis_SWE_SWE96_140
Leptidea_sinapis_SWE_SWE96_139
Leptidea_sinapis_SWE_SWE96_138
Leptidea_sinapis_SWE_SWE96_137
Leptidea_sinapis_SWE_SWE96_136
Leptidea_sinapis_SWE_SWE96_135
Leptidea_sinapis_SWE_SWE96_134
Leptidea_sinapis_SWE_SWE96_131
Leptidea_sinapis_SWE_SWE96_130
Leptidea_sinapis_SWE_SWE96_129
Leptidea_sinapis_SWE_SWE96_123
Leptidea_sinapis_SWE_SWE96_122
Leptidea_sinapis_SWE_SWE96_118
Leptidea_sinapis_SWE_SWE96_116
Leptidea_sinapis_SWE_SWE96_115
Leptidea_sinapis_SWE_SWE96_114
Leptidea_sinapis_SWE_SWE96_113
Leptidea_sinapis_SWE_SWE96_112
Leptidea_sinapis_SWE_SWE96_111
Leptidea_sinapis_SWE_SWE96_110
Leptidea_sinapis_SWE_SWE96_109
Leptidea_sinapis_SWE_SWE96_108
Leptidea_sinapis_SWE_SWE96_107
Leptidea_sinapis_SWE_SWE96_106
Leptidea_sinapis_SWE_SWE96_105
Leptidea_sinapis_SWE_SWE96_104
Leptidea_sinapis_SWE_SWE96_103
Leptidea_sinapis_SWE_SWE96_102
Leptidea_sinapis_SWE_SWE96_101
Leptidea_sinapis_SWE_Lars_10X_104
Leptidea_sinapis_SWE_Lars_10X_103
Leptidea_sinapis_SWE_FD_SWE_Swe−sin−92C
Leptidea_sinapis_SWE_FD_SWE_Swe−sin−91C
Leptidea_sinapis_SWE_FD_SWE_Swe−sin−62C
Leptidea_sinapis_SWE_FD_SWE_Swe−sin−61C
Leptidea_sinapis_SWE_FD_SWE_Swe−sin−32C
Leptidea_sinapis_SWE_FD_SWE_Swe−sin−31C
Leptidea_sinapis_SWE_FD_SWE_Swe−sin−2C
Leptidea_sinapis_SWE_FD_SWE_Swe−sin−1C
Leptidea_sinapis_SWE_FD_SWE_Swe−sin−102C
Leptidea_sinapis_SWE_FD_SWE_Swe−sin−101C
Leptidea_sinapis_CAT_Lars_10X_106
Leptidea_sinapis_CAT_Lars_10X_105
Leptidea_sinapis_CAT_FD_CAT_47
Leptidea_sinapis_CAT_FD_CAT_46
Leptidea_sinapis_CAT_FD_CAT_44
Leptidea_sinapis_CAT_FD_CAT_43
Leptidea_sinapis_CAT_FD_CAT_42
Leptidea_sinapis_CAT_FD_CAT_41
Leptidea_sinapis_CAT_FD_CAT_40
Leptidea_sinapis_CAT_FD_CAT_39
Leptidea_sinapis_CAT_FD_CAT_38
Leptidea_sinapis_CAT_FD_CAT_37
Leptidea_reali_CAT_Lars_10X_116
Leptidea_reali_CAT_Lars_10X_115
Leptidea_reali_CAT_CATrea_36
Leptidea_reali_CAT_CATrea_35
Leptidea_reali_CAT_CATrea_34
Leptidea_reali_CAT_CATrea_33
Leptidea_reali_CAT_CATrea_32
Leptidea_reali_CAT_CATrea_31
Leptidea_reali_CAT_CATrea_30
Leptidea_reali_CAT_CATrea_29
Leptidea_reali_CAT_CATrea_27
Leptidea_reali_CAT_CATrea_26
Leptidea_morsei_CHI_LEP_outgroups_108
Leptidea_morsei_CHI_LEP_outgroups_107
Leptidea_lactea_CHI_LEP_outgroups_106
Leptidea_lactea_CHI_LEP_outgroups_105
Leptidea_juvernica_SWE_SWE96_186
Leptidea_juvernica_SWE_SWE96_133
Leptidea_juvernica_SWE_SWE96_132
Leptidea_juvernica_SWE_SWE96_128
Leptidea_juvernica_SWE_SWE96_127
Leptidea_juvernica_SWE_SWE96_126
Leptidea_juvernica_SWE_SWE96_125
Leptidea_juvernica_SWE_SWE96_124
Leptidea_juvernica_SWE_SWE96_121
Leptidea_juvernica_SWE_SWE96_120
Leptidea_juvernica_SWE_SWE96_119
Leptidea_juvernica_SWE_SWE96_117
Leptidea_juvernica_SWE_Lars_10X_112
Leptidea_juvernica_SWE_Lars_10X_111
Leptidea_amurensis_MON_LEP_outgroups_102
Leptidea_amurensis_MON_LEP_outgroups_101
Comb_ID_MSG
0
2500000
5000000
7500000
Start
Dosage
10.0
7.5
5.0
2.5
0.0

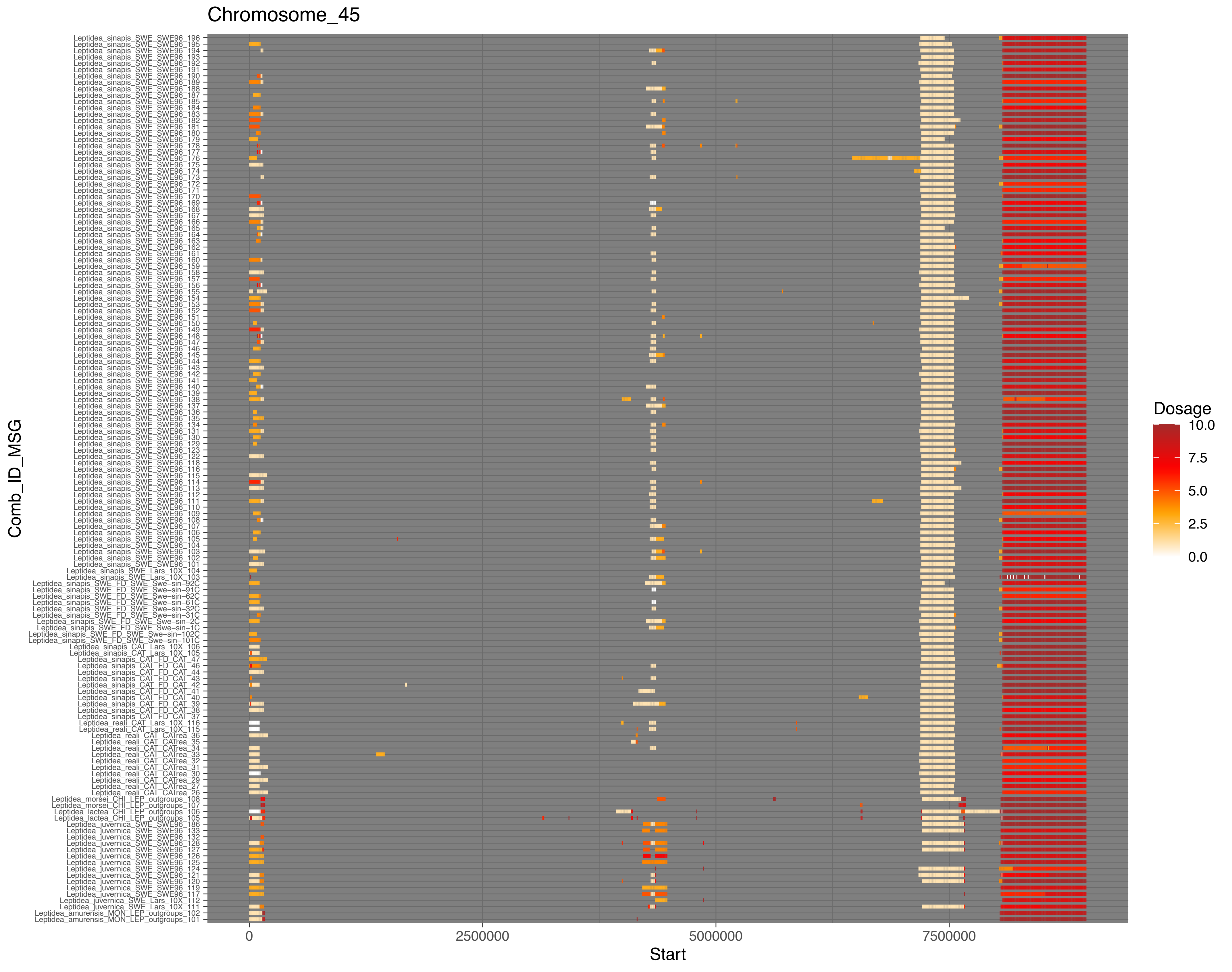
Chromosome_45
Leptidea_sinapis_SWE_SWE96_196
Leptidea_sinapis_SWE_SWE96_195
Leptidea_sinapis_SWE_SWE96_194
Leptidea_sinapis_SWE_SWE96_193
Leptidea_sinapis_SWE_SWE96_192
Leptidea_sinapis_SWE_SWE96_191
Leptidea_sinapis_SWE_SWE96_190
Leptidea_sinapis_SWE_SWE96_189
Leptidea_sinapis_SWE_SWE96_188
Leptidea_sinapis_SWE_SWE96_187
Leptidea_sinapis_SWE_SWE96_185
Leptidea_sinapis_SWE_SWE96_184
Leptidea_sinapis_SWE_SWE96_183
Leptidea_sinapis_SWE_SWE96_182
Leptidea_sinapis_SWE_SWE96_181
Leptidea_sinapis_SWE_SWE96_180
Leptidea_sinapis_SWE_SWE96_179
Leptidea_sinapis_SWE_SWE96_178
Leptidea_sinapis_SWE_SWE96_177
Leptidea_sinapis_SWE_SWE96_176
Leptidea_sinapis_SWE_SWE96_175
Leptidea_sinapis_SWE_SWE96_174
Leptidea_sinapis_SWE_SWE96_173
Leptidea_sinapis_SWE_SWE96_172
Leptidea_sinapis_SWE_SWE96_171
Leptidea_sinapis_SWE_SWE96_170
Leptidea_sinapis_SWE_SWE96_169
Leptidea_sinapis_SWE_SWE96_168
Leptidea_sinapis_SWE_SWE96_167
Leptidea_sinapis_SWE_SWE96_166
Leptidea_sinapis_SWE_SWE96_165
Leptidea_sinapis_SWE_SWE96_164
Leptidea_sinapis_SWE_SWE96_163
Leptidea_sinapis_SWE_SWE96_162
Leptidea_sinapis_SWE_SWE96_161
Leptidea_sinapis_SWE_SWE96_160
Leptidea_sinapis_SWE_SWE96_159
Leptidea_sinapis_SWE_SWE96_158
Leptidea_sinapis_SWE_SWE96_157
Leptidea_sinapis_SWE_SWE96_156
Leptidea_sinapis_SWE_SWE96_155
Leptidea_sinapis_SWE_SWE96_154
Leptidea_sinapis_SWE_SWE96_153
Leptidea_sinapis_SWE_SWE96_152
Leptidea_sinapis_SWE_SWE96_151
Leptidea_sinapis_SWE_SWE96_150
Leptidea_sinapis_SWE_SWE96_149
Leptidea_sinapis_SWE_SWE96_148
Leptidea_sinapis_SWE_SWE96_147
Leptidea_sinapis_SWE_SWE96_146
Leptidea_sinapis_SWE_SWE96_145
Leptidea_sinapis_SWE_SWE96_144
Leptidea_sinapis_SWE_SWE96_143
Leptidea_sinapis_SWE_SWE96_142
Leptidea_sinapis_SWE_SWE96_141
Leptidea_sinapis_SWE_SWE96_140
Leptidea_sinapis_SWE_SWE96_139
Leptidea_sinapis_SWE_SWE96_138
Leptidea_sinapis_SWE_SWE96_137
Leptidea_sinapis_SWE_SWE96_136
Leptidea_sinapis_SWE_SWE96_135
Leptidea_sinapis_SWE_SWE96_134
Leptidea_sinapis_SWE_SWE96_131
Leptidea_sinapis_SWE_SWE96_130
Leptidea_sinapis_SWE_SWE96_129
Leptidea_sinapis_SWE_SWE96_123
Leptidea_sinapis_SWE_SWE96_122
Leptidea_sinapis_SWE_SWE96_118
Leptidea_sinapis_SWE_SWE96_116
Leptidea_sinapis_SWE_SWE96_115
Leptidea_sinapis_SWE_SWE96_114
Leptidea_sinapis_SWE_SWE96_113
Leptidea_sinapis_SWE_SWE96_112
Leptidea_sinapis_SWE_SWE96_111
Leptidea_sinapis_SWE_SWE96_110
Leptidea_sinapis_SWE_SWE96_109
Leptidea_sinapis_SWE_SWE96_108
Leptidea_sinapis_SWE_SWE96_107
Leptidea_sinapis_SWE_SWE96_106
Leptidea_sinapis_SWE_SWE96_105
Leptidea_sinapis_SWE_SWE96_104
Leptidea_sinapis_SWE_SWE96_103
Leptidea_sinapis_SWE_SWE96_102
Leptidea_sinapis_SWE_SWE96_101
Leptidea_sinapis_SWE_Lars_10X_104
Leptidea_sinapis_SWE_Lars_10X_103
Leptidea_sinapis_SWE_FD_SWE_Swe−sin−92C
Leptidea_sinapis_SWE_FD_SWE_Swe−sin−91C
Leptidea_sinapis_SWE_FD_SWE_Swe−sin−62C
Leptidea_sinapis_SWE_FD_SWE_Swe−sin−61C
Leptidea_sinapis_SWE_FD_SWE_Swe−sin−32C
Leptidea_sinapis_SWE_FD_SWE_Swe−sin−31C
Leptidea_sinapis_SWE_FD_SWE_Swe−sin−2C
Leptidea_sinapis_SWE_FD_SWE_Swe−sin−1C
Leptidea_sinapis_SWE_FD_SWE_Swe−sin−102C
Leptidea_sinapis_SWE_FD_SWE_Swe−sin−101C
Leptidea_sinapis_CAT_Lars_10X_106
Leptidea_sinapis_CAT_Lars_10X_105
Leptidea_sinapis_CAT_FD_CAT_47
Leptidea_sinapis_CAT_FD_CAT_46
Leptidea_sinapis_CAT_FD_CAT_44
Leptidea_sinapis_CAT_FD_CAT_43
Leptidea_sinapis_CAT_FD_CAT_42
Leptidea_sinapis_CAT_FD_CAT_41
Leptidea_sinapis_CAT_FD_CAT_40
Leptidea_sinapis_CAT_FD_CAT_39
Leptidea_sinapis_CAT_FD_CAT_38
Leptidea_sinapis_CAT_FD_CAT_37
Leptidea_reali_CAT_Lars_10X_116
Leptidea_reali_CAT_Lars_10X_115
Leptidea_reali_CAT_CATrea_36
Leptidea_reali_CAT_CATrea_35
Leptidea_reali_CAT_CATrea_34
Leptidea_reali_CAT_CATrea_33
Leptidea_reali_CAT_CATrea_32
Leptidea_reali_CAT_CATrea_31
Leptidea_reali_CAT_CATrea_30
Leptidea_reali_CAT_CATrea_29
Leptidea_reali_CAT_CATrea_27
Leptidea_reali_CAT_CATrea_26
Leptidea_morsei_CHI_LEP_outgroups_108
Leptidea_morsei_CHI_LEP_outgroups_107
Leptidea_lactea_CHI_LEP_outgroups_106
Leptidea_lactea_CHI_LEP_outgroups_105
Leptidea_juvernica_SWE_SWE96_186
Leptidea_juvernica_SWE_SWE96_133
Leptidea_juvernica_SWE_SWE96_132
Leptidea_juvernica_SWE_SWE96_128
Leptidea_juvernica_SWE_SWE96_127
Leptidea_juvernica_SWE_SWE96_126
Leptidea_juvernica_SWE_SWE96_125
Leptidea_juvernica_SWE_SWE96_124
Leptidea_juvernica_SWE_SWE96_121
Leptidea_juvernica_SWE_SWE96_120
Leptidea_juvernica_SWE_SWE96_119
Leptidea_juvernica_SWE_SWE96_117
Leptidea_juvernica_SWE_Lars_10X_112
Leptidea_juvernica_SWE_Lars_10X_111
Leptidea_amurensis_MON_LEP_outgroups_102
Leptidea_amurensis_MON_LEP_outgroups_101
Comb_ID_MSG
0
2500000
5000000
7500000
Start
Dosage
10.0
7.5
5.0
2.5
0.0

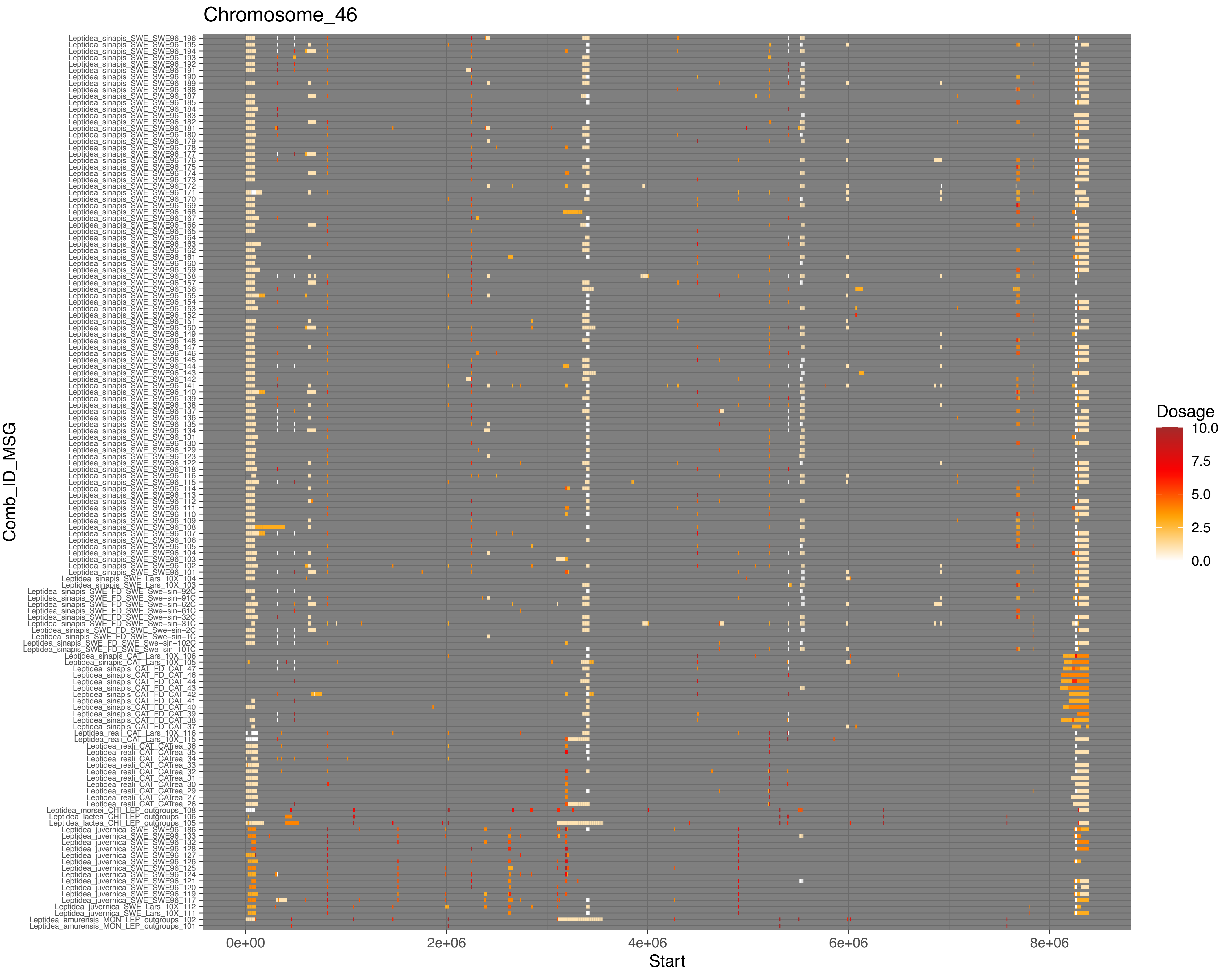

Chromosome_46
Comb_ID_MSG
Leptidea_sinapis_SWE_SWE96_196
Leptidea_sinapis_SWE_SWE96_195
Leptidea_sinapis_SWE_SWE96_194
Leptidea_sinapis_SWE_SWE96_193
Leptidea_sinapis_SWE_SWE96_192
Leptidea_sinapis_SWE_SWE96_191
Leptidea_sinapis_SWE_SWE96_190
Leptidea_sinapis_SWE_SWE96_189
Leptidea_sinapis_SWE_SWE96_188
Leptidea_sinapis_SWE_SWE96_187
Leptidea_sinapis_SWE_SWE96_185
Leptidea_sinapis_SWE_SWE96_184
Leptidea_sinapis_SWE_SWE96_183
Leptidea_sinapis_SWE_SWE96_182
Leptidea_sinapis_SWE_SWE96_181
Leptidea_sinapis_SWE_SWE96_180
Leptidea_sinapis_SWE_SWE96_179
Leptidea_sinapis_SWE_SWE96_178
Leptidea_sinapis_SWE_SWE96_177
Leptidea_sinapis_SWE_SWE96_176
Leptidea_sinapis_SWE_SWE96_175
Leptidea_sinapis_SWE_SWE96_174
Leptidea_sinapis_SWE_SWE96_173
Leptidea_sinapis_SWE_SWE96_172
Leptidea_sinapis_SWE_SWE96_171
Leptidea_sinapis_SWE_SWE96_170
Leptidea_sinapis_SWE_SWE96_169
Leptidea_sinapis_SWE_SWE96_168
Leptidea_sinapis_SWE_SWE96_167
Leptidea_sinapis_SWE_SWE96_166
Leptidea_sinapis_SWE_SWE96_165
Leptidea_sinapis_SWE_SWE96_164
Leptidea_sinapis_SWE_SWE96_163
Leptidea_sinapis_SWE_SWE96_162
Leptidea_sinapis_SWE_SWE96_161
Leptidea_sinapis_SWE_SWE96_160
Leptidea_sinapis_SWE_SWE96_159
Leptidea_sinapis_SWE_SWE96_158
Leptidea_sinapis_SWE_SWE96_157
Leptidea_sinapis_SWE_SWE96_156
Leptidea_sinapis_SWE_SWE96_155
Leptidea_sinapis_SWE_SWE96_154
Leptidea_sinapis_SWE_SWE96_153
Leptidea_sinapis_SWE_SWE96_152
Leptidea_sinapis_SWE_SWE96_151
Leptidea_sinapis_SWE_SWE96_150
Leptidea_sinapis_SWE_SWE96_149
Leptidea_sinapis_SWE_SWE96_148
Leptidea_sinapis_SWE_SWE96_147
Leptidea_sinapis_SWE_SWE96_146
Leptidea_sinapis_SWE_SWE96_145
Leptidea_sinapis_SWE_SWE96_144
Leptidea_sinapis_SWE_SWE96_143
Leptidea_sinapis_SWE_SWE96_142
Leptidea_sinapis_SWE_SWE96_141
Leptidea_sinapis_SWE_SWE96_140
Leptidea_sinapis_SWE_SWE96_139
Leptidea_sinapis_SWE_SWE96_138
Leptidea_sinapis_SWE_SWE96_137
Leptidea_sinapis_SWE_SWE96_136
Leptidea_sinapis_SWE_SWE96_135
Leptidea_sinapis_SWE_SWE96_134
Leptidea_sinapis_SWE_SWE96_131
Leptidea_sinapis_SWE_SWE96_130
Leptidea_sinapis_SWE_SWE96_129
Leptidea_sinapis_SWE_SWE96_123
Leptidea_sinapis_SWE_SWE96_122
Leptidea_sinapis_SWE_SWE96_118
Leptidea_sinapis_SWE_SWE96_116
Leptidea_sinapis_SWE_SWE96_115
Leptidea_sinapis_SWE_SWE96_114
Leptidea_sinapis_SWE_SWE96_113
Leptidea_sinapis_SWE_SWE96_112
Leptidea_sinapis_SWE_SWE96_111
Leptidea_sinapis_SWE_SWE96_110
Leptidea_sinapis_SWE_SWE96_109
Leptidea_sinapis_SWE_SWE96_108
Leptidea_sinapis_SWE_SWE96_107
Leptidea_sinapis_SWE_SWE96_106
Leptidea_sinapis_SWE_SWE96_105
Leptidea_sinapis_SWE_SWE96_104
Leptidea_sinapis_SWE_SWE96_103
Leptidea_sinapis_SWE_SWE96_102
Leptidea_sinapis_SWE_SWE96_101
Leptidea_sinapis_SWE_Lars_10X_104
Leptidea_sinapis_SWE_Lars_10X_103
Leptidea_sinapis_SWE_FD_SWE_Swe−sin−92C
Leptidea_sinapis_SWE_FD_SWE_Swe−sin−91C
Leptidea_sinapis_SWE_FD_SWE_Swe−sin−62C
Leptidea_sinapis_SWE_FD_SWE_Swe−sin−61C
Leptidea_sinapis_SWE_FD_SWE_Swe−sin−32C
Leptidea_sinapis_SWE_FD_SWE_Swe−sin−31C
Leptidea_sinapis_SWE_FD_SWE_Swe−sin−2C
Leptidea_sinapis_SWE_FD_SWE_Swe−sin−1C
Leptidea_sinapis_SWE_FD_SWE_Swe−sin−102C
Leptidea_sinapis_SWE_FD_SWE_Swe−sin−101C
Leptidea_sinapis_CAT_Lars_10X_106
Leptidea_sinapis_CAT_Lars_10X_105
Leptidea_sinapis_CAT_FD_CAT_47
Leptidea_sinapis_CAT_FD_CAT_46
Leptidea_sinapis_CAT_FD_CAT_44
Leptidea_sinapis_CAT_FD_CAT_43
Leptidea_sinapis_CAT_FD_CAT_42
Leptidea_sinapis_CAT_FD_CAT_41
Leptidea_sinapis_CAT_FD_CAT_40
Leptidea_sinapis_CAT_FD_CAT_39
Leptidea_sinapis_CAT_FD_CAT_38
Leptidea_sinapis_CAT_FD_CAT_37
Leptidea_reali_CAT_Lars_10X_116
Leptidea_reali_CAT_Lars_10X_115
Leptidea_reali_CAT_CATrea_36
Leptidea_reali_CAT_CATrea_35
Leptidea_reali_CAT_CATrea_34
Leptidea_reali_CAT_CATrea_33
Leptidea_reali_CAT_CATrea_32
Leptidea_reali_CAT_CATrea_31
Leptidea_reali_CAT_CATrea_30
Leptidea_reali_CAT_CATrea_29
Leptidea_reali_CAT_CATrea_27
Leptidea_reali_CAT_CATrea_26
Leptidea_morsei_CHI_LEP_outgroups_108
Leptidea_lactea_CHI_LEP_outgroups_106
Leptidea_lactea_CHI_LEP_outgroups_105
Leptidea_juvernica_SWE_SWE96_186
Leptidea_juvernica_SWE_SWE96_133
Leptidea_juvernica_SWE_SWE96_132
Leptidea_juvernica_SWE_SWE96_128
Leptidea_juvernica_SWE_SWE96_127
Leptidea_juvernica_SWE_SWE96_126
Leptidea_juvernica_SWE_SWE96_125
Leptidea_juvernica_SWE_SWE96_124
Leptidea_juvernica_SWE_SWE96_121
Leptidea_juvernica_SWE_SWE96_120
Leptidea_juvernica_SWE_SWE96_119
Leptidea_juvernica_SWE_SWE96_117
Leptidea_juvernica_SWE_Lars_10X_112
Leptidea_juvernica_SWE_Lars_10X_111
Leptidea_amurensis_MON_LEP_outgroups_102
Leptidea_amurensis_MON_LEP_outgroups_101
0e+00
2e+06
4e+06
6e+06
8e+06
Start
Dosage
10.0
7.5
5.0
2.5
0.0

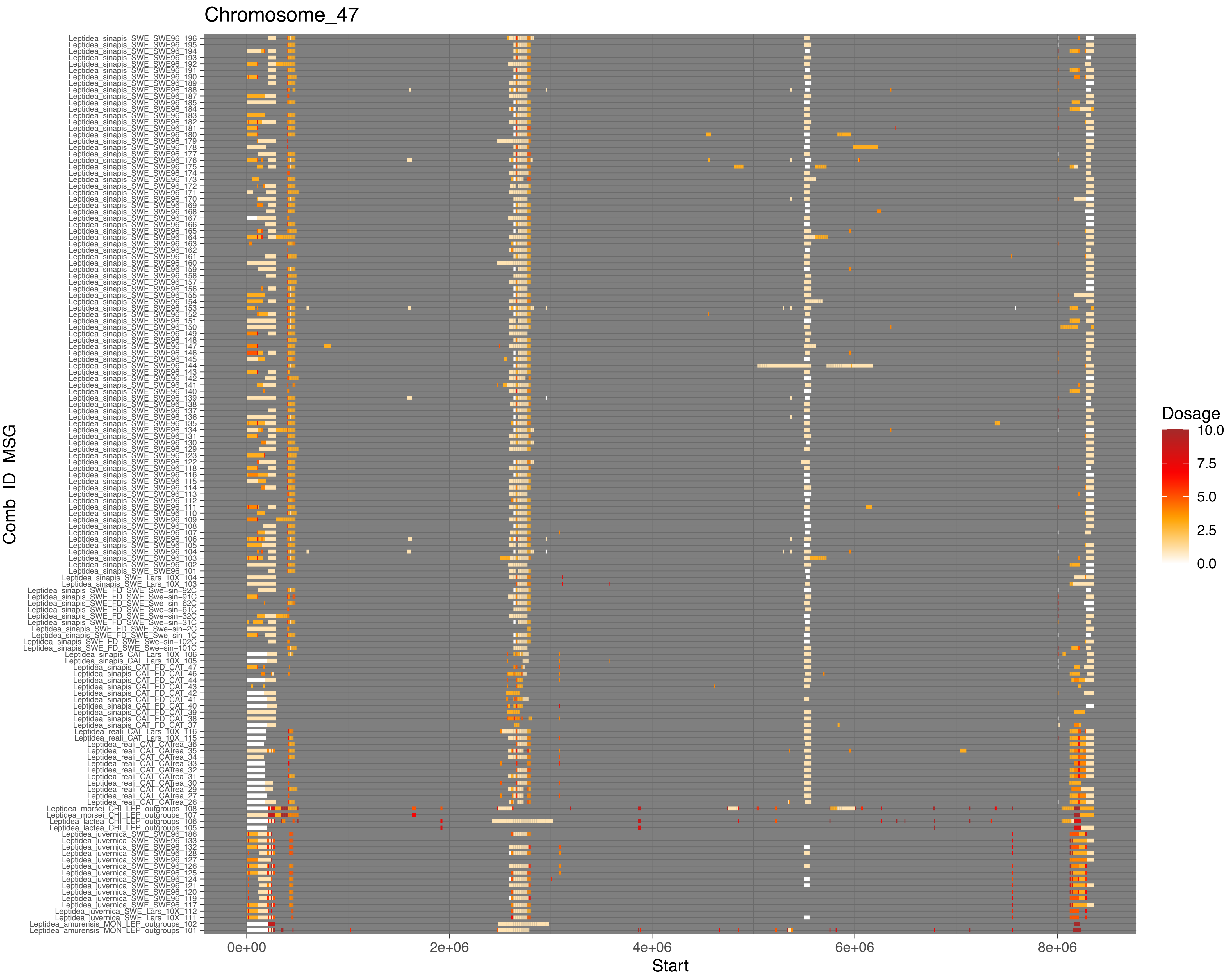
Chromosome_47
Comb_ID_MSG
Leptidea_sinapis_SWE_SWE96_196
Leptidea_sinapis_SWE_SWE96_195
Leptidea_sinapis_SWE_SWE96_194
Leptidea_sinapis_SWE_SWE96_193
Leptidea_sinapis_SWE_SWE96_192
Leptidea_sinapis_SWE_SWE96_191
Leptidea_sinapis_SWE_SWE96_190
Leptidea_sinapis_SWE_SWE96_189
Leptidea_sinapis_SWE_SWE96_188
Leptidea_sinapis_SWE_SWE96_187
Leptidea_sinapis_SWE_SWE96_185
Leptidea_sinapis_SWE_SWE96_184
Leptidea_sinapis_SWE_SWE96_183
Leptidea_sinapis_SWE_SWE96_182
Leptidea_sinapis_SWE_SWE96_181
Leptidea_sinapis_SWE_SWE96_180
Leptidea_sinapis_SWE_SWE96_179
Leptidea_sinapis_SWE_SWE96_178
Leptidea_sinapis_SWE_SWE96_177
Leptidea_sinapis_SWE_SWE96_176
Leptidea_sinapis_SWE_SWE96_175
Leptidea_sinapis_SWE_SWE96_174
Leptidea_sinapis_SWE_SWE96_173
Leptidea_sinapis_SWE_SWE96_172
Leptidea_sinapis_SWE_SWE96_171
Leptidea_sinapis_SWE_SWE96_170
Leptidea_sinapis_SWE_SWE96_169
Leptidea_sinapis_SWE_SWE96_168
Leptidea_sinapis_SWE_SWE96_167
Leptidea_sinapis_SWE_SWE96_166
Leptidea_sinapis_SWE_SWE96_165
Leptidea_sinapis_SWE_SWE96_164
Leptidea_sinapis_SWE_SWE96_163
Leptidea_sinapis_SWE_SWE96_162
Leptidea_sinapis_SWE_SWE96_161
Leptidea_sinapis_SWE_SWE96_160
Leptidea_sinapis_SWE_SWE96_159
Leptidea_sinapis_SWE_SWE96_158
Leptidea_sinapis_SWE_SWE96_157
Leptidea_sinapis_SWE_SWE96_156
Leptidea_sinapis_SWE_SWE96_155
Leptidea_sinapis_SWE_SWE96_154
Leptidea_sinapis_SWE_SWE96_153
Leptidea_sinapis_SWE_SWE96_152
Leptidea_sinapis_SWE_SWE96_151
Leptidea_sinapis_SWE_SWE96_150
Leptidea_sinapis_SWE_SWE96_149
Leptidea_sinapis_SWE_SWE96_148
Leptidea_sinapis_SWE_SWE96_147
Leptidea_sinapis_SWE_SWE96_146
Leptidea_sinapis_SWE_SWE96_145
Leptidea_sinapis_SWE_SWE96_144
Leptidea_sinapis_SWE_SWE96_143
Leptidea_sinapis_SWE_SWE96_142
Leptidea_sinapis_SWE_SWE96_141
Leptidea_sinapis_SWE_SWE96_140
Leptidea_sinapis_SWE_SWE96_139
Leptidea_sinapis_SWE_SWE96_138
Leptidea_sinapis_SWE_SWE96_137
Leptidea_sinapis_SWE_SWE96_136
Leptidea_sinapis_SWE_SWE96_135
Leptidea_sinapis_SWE_SWE96_134
Leptidea_sinapis_SWE_SWE96_131
Leptidea_sinapis_SWE_SWE96_130
Leptidea_sinapis_SWE_SWE96_129
Leptidea_sinapis_SWE_SWE96_123
Leptidea_sinapis_SWE_SWE96_122
Leptidea_sinapis_SWE_SWE96_118
Leptidea_sinapis_SWE_SWE96_116
Leptidea_sinapis_SWE_SWE96_115
Leptidea_sinapis_SWE_SWE96_114
Leptidea_sinapis_SWE_SWE96_113
Leptidea_sinapis_SWE_SWE96_112
Leptidea_sinapis_SWE_SWE96_111
Leptidea_sinapis_SWE_SWE96_110
Leptidea_sinapis_SWE_SWE96_109
Leptidea_sinapis_SWE_SWE96_108
Leptidea_sinapis_SWE_SWE96_107
Leptidea_sinapis_SWE_SWE96_106
Leptidea_sinapis_SWE_SWE96_105
Leptidea_sinapis_SWE_SWE96_104
Leptidea_sinapis_SWE_SWE96_103
Leptidea_sinapis_SWE_SWE96_102
Leptidea_sinapis_SWE_SWE96_101
Leptidea_sinapis_SWE_Lars_10X_104
Leptidea_sinapis_SWE_Lars_10X_103
Leptidea_sinapis_SWE_FD_SWE_Swe−sin−92C
Leptidea_sinapis_SWE_FD_SWE_Swe−sin−91C
Leptidea_sinapis_SWE_FD_SWE_Swe−sin−62C
Leptidea_sinapis_SWE_FD_SWE_Swe−sin−61C
Leptidea_sinapis_SWE_FD_SWE_Swe−sin−32C
Leptidea_sinapis_SWE_FD_SWE_Swe−sin−31C
Leptidea_sinapis_SWE_FD_SWE_Swe−sin−2C
Leptidea_sinapis_SWE_FD_SWE_Swe−sin−1C
Leptidea_sinapis_SWE_FD_SWE_Swe−sin−102C
Leptidea_sinapis_SWE_FD_SWE_Swe−sin−101C
Leptidea_sinapis_CAT_Lars_10X_106
Leptidea_sinapis_CAT_Lars_10X_105
Leptidea_sinapis_CAT_FD_CAT_47
Leptidea_sinapis_CAT_FD_CAT_46
Leptidea_sinapis_CAT_FD_CAT_44
Leptidea_sinapis_CAT_FD_CAT_43
Leptidea_sinapis_CAT_FD_CAT_42
Leptidea_sinapis_CAT_FD_CAT_41
Leptidea_sinapis_CAT_FD_CAT_40
Leptidea_sinapis_CAT_FD_CAT_39
Leptidea_sinapis_CAT_FD_CAT_38
Leptidea_sinapis_CAT_FD_CAT_37
Leptidea_reali_CAT_Lars_10X_116
Leptidea_reali_CAT_Lars_10X_115
Leptidea_reali_CAT_CAtrea_36
Leptidea_reali_CAT_CAtrea_35
Leptidea_reali_CAT_CAtrea_34
Leptidea_reali_CAT_CAtrea_33
Leptidea_reali_CAT_CAtrea_32
Leptidea_reali_CAT_CAtrea_31
Leptidea_reali_CAT_CAtrea_30
Leptidea_reali_CAT_CAtrea_29
Leptidea_reali_CAT_CAtrea_27
Leptidea_reali_CAT_CAtrea_26
Leptidea_morsei_CHI_LEP_outgroups_108
Leptidea_morsei_CHI_LEP_outgroups_107
Leptidea_lactea_CHI_LEP_outgroups_106
Leptidea_lactea_CHI_LEP_outgroups_105
Leptidea_juvernica_SWE_SWE96_186
Leptidea_juvernica_SWE_SWE96_133
Leptidea_juvernica_SWE_SWE96_132
Leptidea_juvernica_SWE_SWE96_128
Leptidea_juvernica_SWE_SWE96_127
Leptidea_juvernica_SWE_SWE96_126
Leptidea_juvernica_SWE_SWE96_125
Leptidea_juvernica_SWE_SWE96_124
Leptidea_juvernica_SWE_SWE96_121
Leptidea_juvernica_SWE_SWE96_120
Leptidea_juvernica_SWE_SWE96_119
Leptidea_juvernica_SWE_SWE96_117
Leptidea_juvernica_SWE_Lars_10X_112
Leptidea_juvernica_SWE_Lars_10X_111
Leptidea_amurensis_MON_LEP_outgroups_102
Leptidea_amurensis_MON_LEP_outgroups_101
0e+00
2e+06
4e+06
6e+06
8e+06
Start
Dosage
10.0
7.5
5.0
2.5
0.0

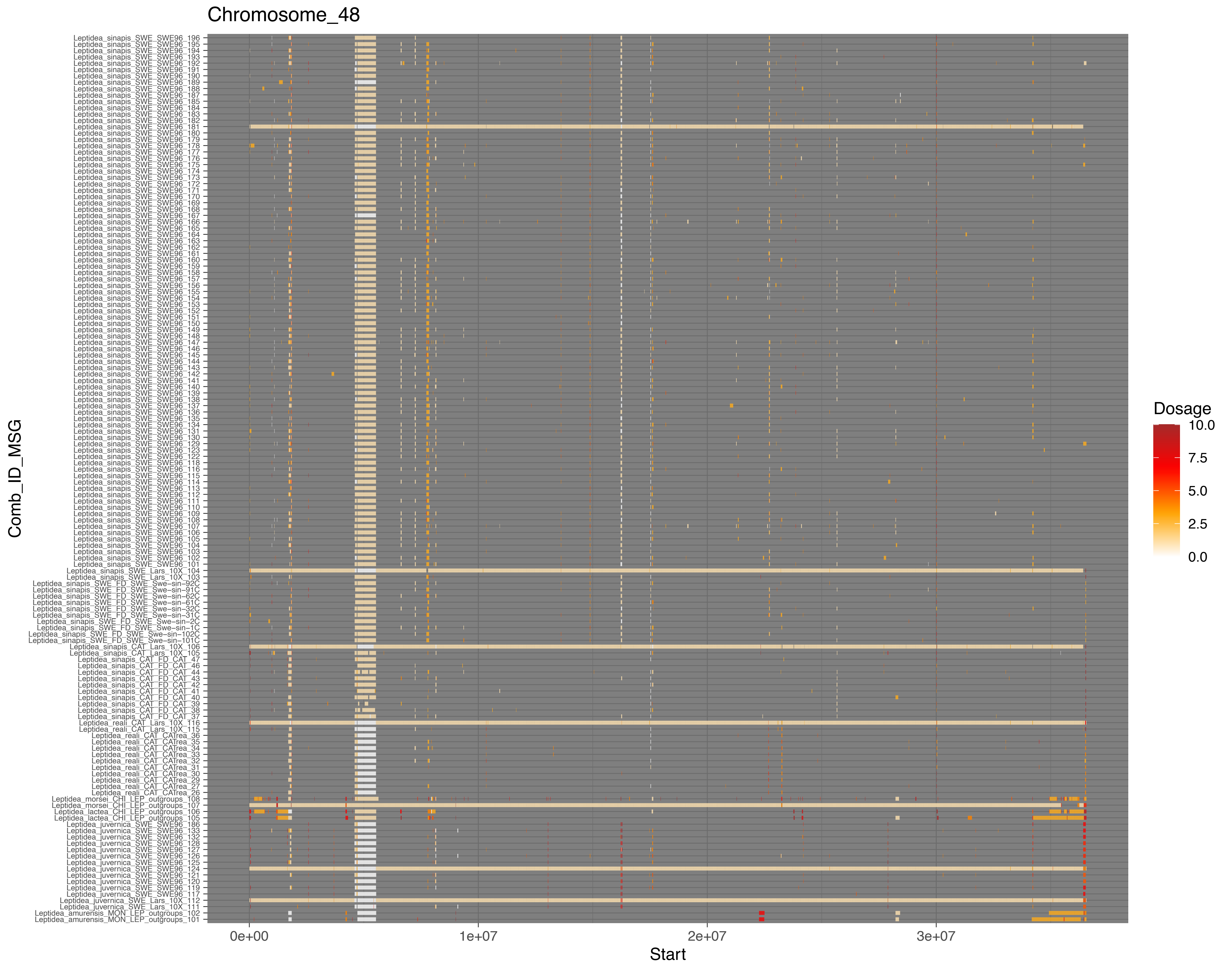
Chromosome_48
Comb_ID_MSG
Start
0e+00
1e+07
2e+07
3e+07
Dosage
10.0
7.5
5.0
2.5
0.0
